# Decoding the human PBMC isonome: Isoform-level resolution with single-cell long-read transcriptomics

**DOI:** 10.1101/2025.10.16.682832

**Authors:** Patricia Hayes Doyle, Madeline L. Page, J. Anthony Brandon, Bernardo Aguzzoli-Heberle, Brendan J. White, Ann M. Stowe, Mark T. W. Ebbert

## Abstract

Long-read single-cell RNA sequencing provides an opportunity to understand human health and disease at a level difficult to resolve with bulk or short-read methods. This approach enables isoform-level investigation of cellular diversity and disease mechanisms and definition of cell-types, rather than using genes alone. Using a modified, microfluidic-free PIPseq workflow and computational pipeline adapted for Oxford Nanopore long-read sequencing, we generated the largest long-read single-cell dataset of human peripheral blood mononuclear cells (PBMCs) to date. This study profiled isoform usage across immune cells, integrating marker expression and isoform discovery. We identified 128 novel isoforms from known and new genes, several with distinct cell-type-specific patterns, and characterized marker gene isoform expression across cell-types. Non-canonical protein-coding variants of *GZMB* and *CD3G* were enriched in unexpected cell-types, including megakaryocytes and monocyte-derived populations. We also discovered novel transcripts from *CMC1* and *LYAR* with cell-type-specific signatures that were the predominantly expressed transcript within the gene. This study expands versatility of long-read single-cell studies to not only relay changes in isoform signatures, but to position them within the functional context of the biology they impact. These results demonstrate the power of long-read single-cell sequencing for mapping the isoform landscape—the isonome—across tissues and disease contexts.

## Introduction

To resolve the etiology of complex diseases and develop diagnostics and treatments, researchers pursue various genetic and genomic approaches, including DNA and RNA sequencing and proteomics. Any one of these avenues often fails to reflect the true spectrum of disease on its own, but combined, we can obtain a much clearer picture. To this end, there has long been a strong interest in the genomics community in understanding various alterations in transcriptional regulation, such as post-translational modification and alternative splicing, to improve our understanding of the underlying regulatory mechanisms of disease. Alternative splicing events affect nearly 85% of all protein-coding genes in the human genome^1^, and aberrant events have been associated with various disease states, including cancer and neurodegenerative disease^2,3^. This connection indicates the importance of characterizing the diversity of RNA isoforms, which we hereby refer to as the “isonome”, and determining their role across human health and disease.

Long-read sequencing technologies, such as Oxford Nanopore Technologies (ONT), enable researchers to detect both gene and isoform expression patterns because longer reads can more accurately measure full-length transcripts^4^. This is an improvement over standard short-read technologies, which struggle to measure isoform-level expression^5^. Long-read sequencing has shown great promise and was named the Method of the Year by Nature Methods in 2023^6^. Several labs recently used bulk long-read sequencing to discover thousands of new RNA isoforms^7–9^, among other findings, including our own work in a small cohort of Alzheimer’s disease (AD) cases and controls, where we identified 53 novel isoforms from disease-associated genes and demonstrated the importance of performing differential expression at the isoform level, not just the gene level^9^.

While bulk RNA sequencing has great value, being able to characterize and quantify expression within individual cells via single-cell RNA sequencing (scRNA-seq), provides crucial context into cell-specific mechanisms. Researchers have used short-read scRNA-seq for many years, and it has become a powerful tool for disease research, enabling gene-level analyses at single-cell resolution. These studies offer direct insight into downstream cellular mechanisms for specific, critical cell populations unidentifiable via bulk approaches.

scRNA-seq typically uses short-read sequencing, however, which collapses expression measurements across all RNA isoforms for a given gene into a single gene expression measurement, ignoring isoform differences that may provide critical insight into molecular dysfunction^1,9^. Combining scRNA-seq and long-read RNA sequencing approaches allows a precise cellular study of isoform expression, offering unprecedented clarity into genomics, disease mechanisms, and potential drug targets. The integration of single-cell and long-read technologies has been performed in a relatively few number of studies^10–27^. The 10X Genomics droplet-based system is the most established and common among single-cell long-read studies^12–24^, but new benchtop-friendly approaches make single-cell approaches more accessible, deviating from the expensive microfluidic machinery that is prone to clogging by cell clumps or debris^26^ and reducing the cost per sample. These newer approaches, including those from Fluent Biosciences (recently acquired by Illumina)^28^ and Parse Biosciences^26^, utilize benchtop-friendly microfluidic-free approaches.

Although long reads have many advantages, short reads are often used to supplement long reads in single-cell sequencing because the short reads: (1) correct sequencing errors in the long reads (the higher error rate in long-read sequencing makes confidently identifying barcodes challenging); and (2) help identify cluster cell-types with the increased depth. The read depth can be a challenge with long-read single-cell approaches due to cost. Read depth from long reads is not necessarily insufficient when analyzing data at the gene level (i.e., collapsing all isoforms into a single expression measurement), but it becomes a challenge at the isoform level. Thus, the same number of reads is getting split across a dramatically higher number of dimensions (i.e., expression values). For example, a recent study by Page et al.^1^, using data from Glinos et al.^7^, demonstrated that 13,236 unique genes are regularly expressed in the cerebellum, resulting in 22,522 uniquely expressed isoforms. Indeed, to truly grasp biology at both the single-cell and isoform level will require extremely deep long-read sequencing.

Here, we present: (1) a single-cell protocol that has been adapted for use with ONT long reads, (2) a bespoke computational approach (see **Data and code availability**) to identify barcodes and maximize read recovery in long-read single-cell approaches using ONT, and (3) proof-of-principle results demonstrating the value of this approach using data that are biologically relevant to peripheral blood mononuclear cells (PBMCs). Specifically, we adapted the V4.0PLUS PIPseq protocol originally developed for short-read sequencing by Fluent Biosciences^28^ to be compatible with long-read nanopore sequencing, combining the strengths of single-cell approaches with long-read sequencing in a benchtop-friendly design; this, in part, required small but essential modifications to the protocol and our custom computational approach to identify barcodes and maximize read recovery that is conceptually simple but computationally challenging. In doing so, we enable isoform expression analysis with cell-type specificity. For this study, we chose to use PBMCs to validate our novel adaptation of long-read PIPseq, given their common use as a clinical sample, cellular diversity, ease of collection, and the absence of well-powered studies on the single-cell long-read level.

## Results

### Adapted PIPseq protocol enables bench-top long-read single-cell sequencing

In this study, we adapted the PIPseq^28^ T20 3’ Single Cell RNA Kit developed by Fluent Biosciences (now marketed as Illumina Single Cell 3’ RNA Prep, with updated chemistry) for long-read sequencing using ONT (**Fig. 1**) and performed sequencing on two PBMC technical replicates (three PromethION flow cells per replicate) across the largest number of cells in any single-cell long-read study to date. We loaded 60,000 PBMCs per technical replicate to target 30,000 PBMCs (50% capture rate; **Fig. 1a**), the upper limit of the PIPseq T20 chemistry. The PBMCs were isolated from whole blood immediately after collection to preserve RNA integrity and minimize transcriptomic changes. We systematically modified the PIPseq protocol through several key steps to maintain mRNA and cDNA read length (**Fig. 1b,c**) and ensure compatibility with downstream ONT sequencing. Complete details are included in the **Methods**, but briefly, this was done through a series of key modifications: (1) where possible, we replaced standard pipette tips with wide-bore tips to reduce shear forces while handling fluids; and (2) we eliminated vortexing steps where possible, opting for gentler mixing methods including tube inversion, flicking, or careful pipetting. These steps were especially utilized after barcoded cDNA was removed from the protective PIP environment. To bridge cDNA produced from the modified PIPseq protocol with an improved ONT single-cell protocol, we introduced custom oligos designed for compatibility with both. The samples were then prepared for ONT sequencing using the PCR-cDNA Sequencing Kit V14 with a modified single-cell preparation optimized for preserving read length, as described above (**Fig. 1c**).

**Figure 1.**
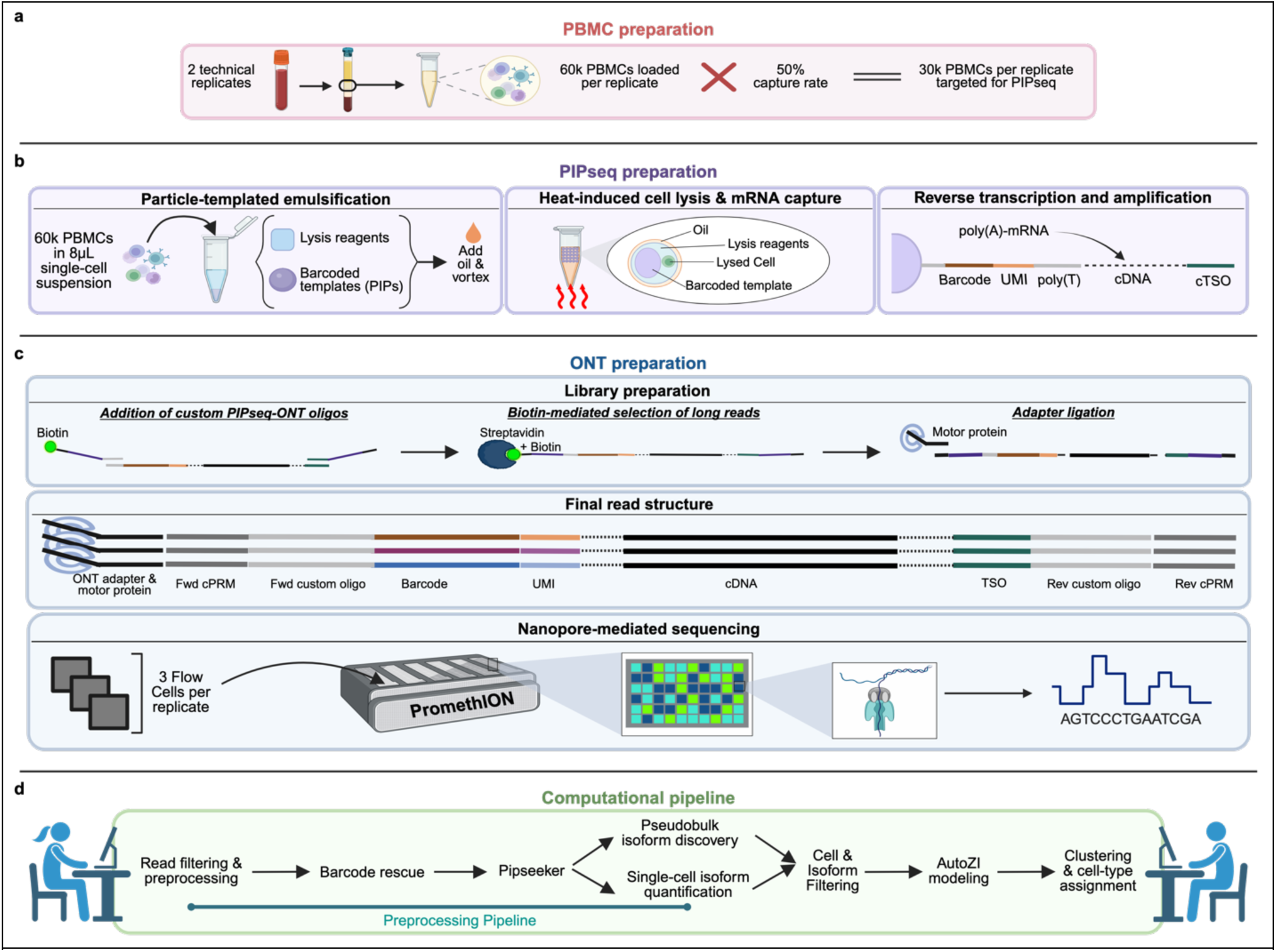
Adapted PIPSeq protocol enables bench-top long-read single-cell sequencing. **(a)** Buffy coat from whole blood was processed and 30k cells per replicate were targeted for capture. **(b)** Single cells were encapsulated in droplets with barcoded templates, lysed, and mRNA captured for reverse transcription and whole-transcriptome amplification to preserve cell-of-origin information with barcoded cDNA. (**c)** Barcoded cDNA underwent quality assurance, adapter addition, biotin-mediated enrichment of long fragments, and preparation for PromethION sequencing. **(d)** Reads underwent quality filtering, barcodes were rescued, and reads were processed with PIPseeker and fed into pipelines for bulk-level isoform discovery and single-cell isoform quantification. Clustering and cell-type analyses were then performed. *Figure created using BioRender.com*.

Following sequencing, we processed the reads for quality control and demultiplexing into individual cell expression matrices (**Fig. 1d**). Briefly, we performed the following steps in our preprocessing pipeline: (1) filtering out low-quality reads (quality score < 9) and rescuing full-length reads with *pychopper*; (2) correcting and rescuing barcodes using a modified PIPseeker workflow optimized for long-reads; (3) generating pseudo paired-end reads for filtering and demultiplexing using PIPseeker; and (4) rescuing reads with imperfect barcodes from sequencing error.

### Bulk-level characterization reveals isoform expression landscape in PBMCs

Before performing single-cell analyses, we first characterized the gene and isoform expression landscape in PBMCs to better understand the nature of the peripheral immune cells at a more profound level, including which isoforms for key PBMC genes are driving the underlying biology for these critical cells in the human immune system. During this process, we also discovered new isoforms and new gene bodies using Bambu^29^, and identified genes where individual isoforms appear to be specific to different cell types.

Sequencing generated a total of 275.8M and 216M raw reads for each respective sample (**Table 1**), yielding the largest and most extensive long-read PBMC dataset to date^18,30^. After quality filtering and alignment, approximately 121.8M and 80.0M reads remained, enabling high-confidence isoform detection. The mean read length N50s were 658.7 and 664.7, respectively, while the median read lengths were 476.7 and 501.3, respectively.

**Table 1.** Read metrics from PBMC samples collected across 3 flow cells per sample. Values signify metrics from ONT sequencing, filtering, and alignment aggregated across the three flow cells used to sequence each PBMC replicate. Full metrics after each step can be found in **Supplemental** Table 1. Abbreviations: ONT, Oxford Nanopore Technologies; MAPQ, mapping quality score; nt, nucleotides.

|  | PBMC1 | PBMC2 |
| --- | --- | --- |
| N reads (total) | 275,827,438 | 215,987,216 |
| N reads with quality score >9 (ONT) | 228,920,664 | 163,153,766 |
| N trimmed and processed passed reads | 153,038,291 | 99,777,656 |
| N reads aligned (MAPQ>10) | 121,867,484 | 80,033,261 |
| N50 FASTQ (mean across flow cells, nt) | 658.7 | 664.7 |
| Median read length (nt) | 476.7 | 501.3 |

Our initial analyses characterized RNA isoform expression characteristics; this included performing isoform discovery using Bambu^29^ across all cells for both samples to maximize evidence of new isoforms. Afterward, we performed rigorous cell and isoform filtering (see **Methods**), ultimately resulting in 11,352 and 21,666 cells for PBMC1 and PBMC2, respectively.

We identified 128 new isoforms in total, where 59 were from previously annotated gene bodies (new from known), and 67 were from previously unidentified gene bodies (new from new). Genes were identified as “new” when Bambu identified consistent expression at loci that were not associated with known Ensembl^31^ Gene IDs. Similarly, new isoforms were defined as those that did not associate with known Ensembl Transcript IDs. Of these 128 isoforms, 18 were previously identified by other groups^7–9^ (**Fig. S1**) and 7 were predicted to be protein-coding, according to analyses using the Coding Potential Calculator 2^32,33^(**Supplemental File 1**).

Many multi-isoform genes expressed only two isoforms, however 50.6% expressed ≥5 isoforms and 15.5% expressed ≥10 isoforms (**Fig. 2a**). New transcripts from known genes were relatively short, with a median of 453 nucleotides (**Fig. 2b**). For reference, the overall median length for all isoforms in our dataset was 1737nt across both samples (**Fig S2**). Although all the new isoforms from known genes exhibited multiple exons in the novel transcript, ∼64% of these transcripts contained just two exons (**Fig. 2c**).

**Figure 2.**
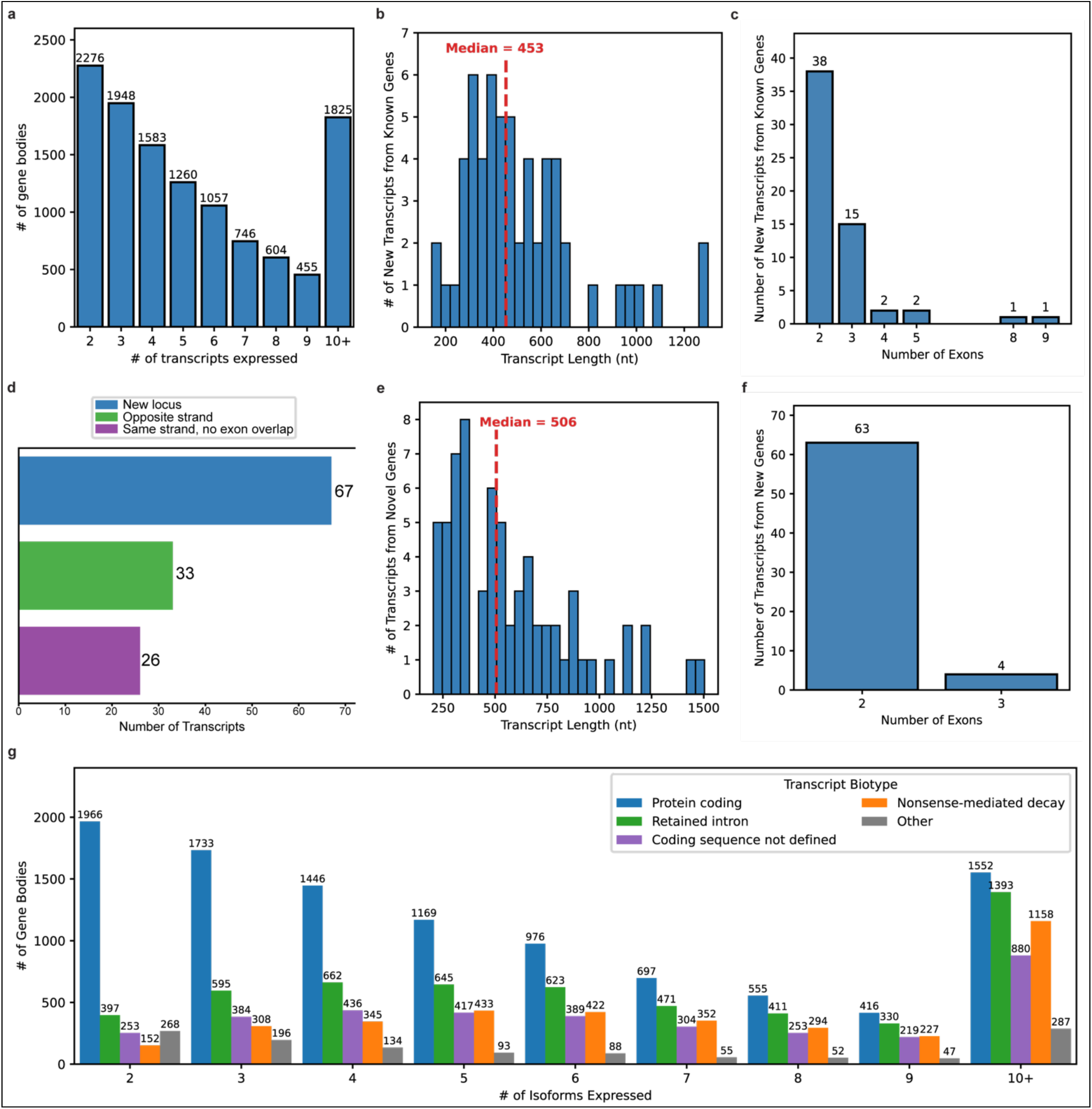
Bulk-level characterization of transcripts from long-read single-cell PBMC data after QC. **(a)** Number of gene bodies expressing more than one isoform. **(b)** Transcript length distribution (nt) for new isoforms found in known genes (median = 453nt, range = 142 to 1300). **(c)** Exon number distribution for new isoforms from known genes. **(d)** Bar plot demonstrating genomic position of new transcripts in relation to known genes and transcripts. **(e)** Transcript length distribution for new isoforms from new genes (median = 506nt, range = 204 to 1504). **(f)** Exon number distribution for new isoforms from new genes. **(g)** Bar plot of gene bodies by isoform number, as in 2a, subdivided by transcript biotype. The “Other” Category includes biotypes such as long non-coding RNAs and nonsense mediated decay.

We also investigated the genomic position of new transcripts from known genes and found that 55.9% of the new isoforms resided on the opposite strand of the known gene, while the remaining 44.1% resided on the same strand with no exonic overlap (**Fig. 2d**). Our 67 new gene bodies, however, were identified as entirely new loci, which could be important for better understanding peripheral immune cells. These new transcripts from previously unannotated genes had a median transcript length of 506nt (**Fig. 2e**). Of these new loci, ∼94% contained just two loci, consistent with the trend that we identified in our novel isoforms from known genes. Given the relatively short length of many of our new isoforms and our short median read length (**Table 1**), it is possible some may be incomplete fragments of a larger isoform.

We identified that most of the transcripts expressed in the PBMCs were annotated as protein-coding, according to Ensembl (HG38 release 113)^31^ regardless of the number of isoforms expressed in the gene body (**Fig. 2g**). However, as the number of isoforms expressed within a single gene increases across the distribution, we noticed that fewer protein-coding isoforms are expressed in favor of a higher proportion of other transcript biotypes, such as those with retained introns or annotated as nonsense-mediated decay. Several transcripts also appear to have undefined coding sequences (e.g *CD33*, *AHR*), reflecting a remaining gap in our understanding of functional isoform expression.

### Canonical marker genes exhibit high isoform diversity

After performing isoform discovery in the bulk-level analysis, we proceeded to perform the single-cell analyses. We utilized AutoZI to adjust for zero inflation, which occurs when many genes or isoforms exhibit zero counts more often than expected due to limited read depth or capture. This phenomenon is more common in single-cell data sets since reads are split across individual cells rather than pooled, as in bulk-level data, making it difficult to directly measure expression consistently across the cells. Unlike other methods, AutoZI identifies genes susceptible to zero inflation (**Table S3**) and applies a zero-inflated negative binomial (ZINB) distribution to estimate the true expression values for individual genes and isoforms^34^. After zero-inflation correction, we constructed a K-nearest neighbor (KNN) graph with 20 neighbors, performed unsupervised clustering with the Leiden algorithm (resolution: 0.06), and visualized clusters using UMAP. Using known markers (**Table S4** and **Methods**), we identified five clusters with major immune cell types at both the gene (**Fig. 3a**) and isoform levels (**Fig. 3b**); in order of abundance, these are T cells (24,757 cells), natural killer (NK) cells (4,623), B cells (2,231), monocyte-derived cells (1,339), and megakaryocytes (68). While canonically considered rare for megakaryocytes to be found in the circulation, except in the instance of bone marrow-related diseases, we identified a very small population through genetic markers (*GP1BA*, *ITGA2B*, and *MPL*) distinct from circulating neutrophils; our findings concur with previous scRNA-seq work that observed megakaryocytes in circulation.

**Figure 3.**
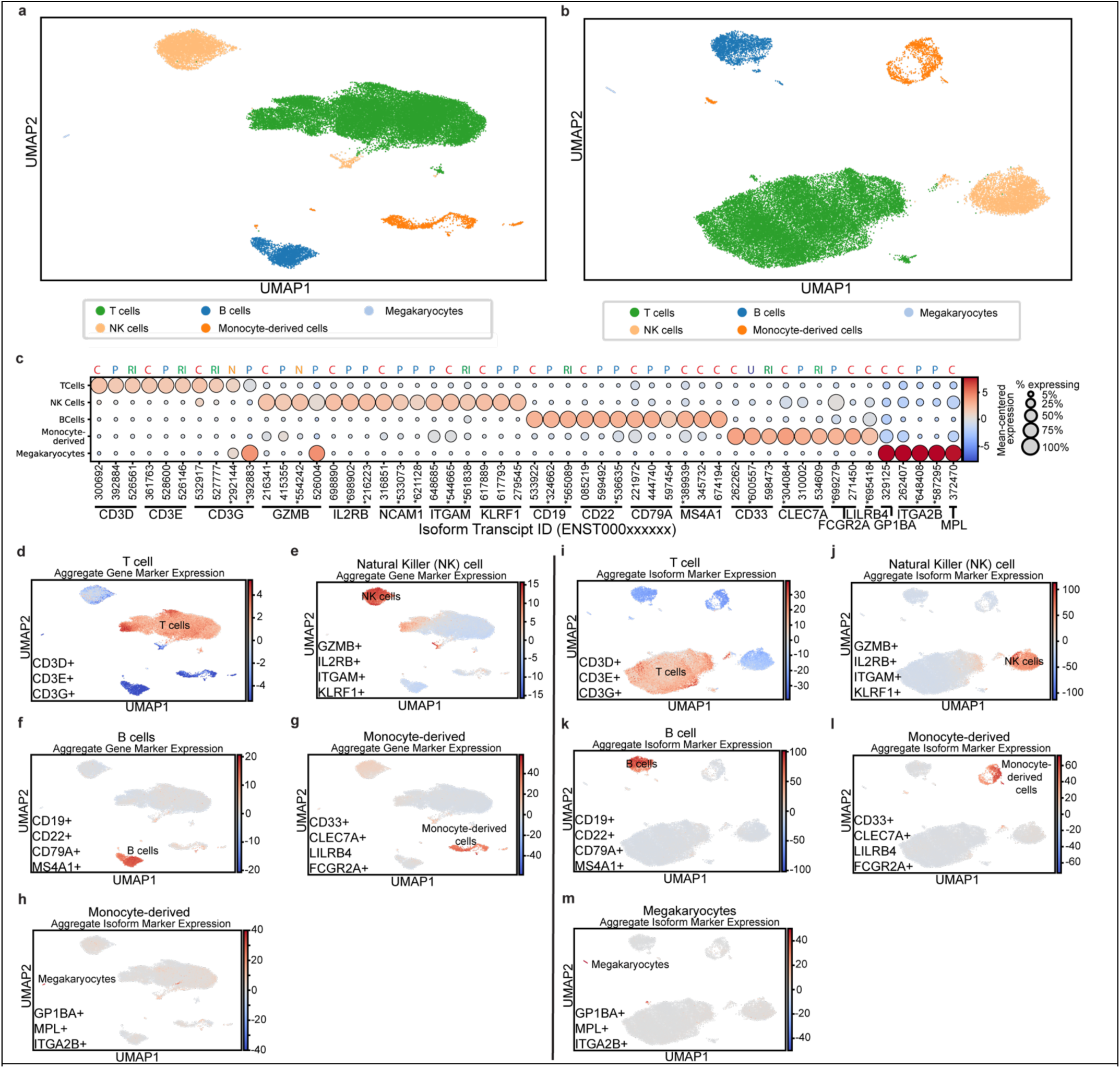
Immune cell clustering and marker-based cell-type assignment at the gene- and isoform-level. (**a-b**) UMAP projections of cell clustering, colored by assigned cell-type: (**a**) gene-level clustering and (**b**) isoform-level clustering. (**c**) Dot plot of up to three representative isoforms per gene, selected as the most highly expressed isoforms by raw counts (**Supplemental File 2**), and including any isoforms enriched in a non-canonical cell-type (e.g. *GZMB* expression in megakaryocytes). The x-axis represents isoform transcript ID (ENST00000xxxxxx), with the last 6 digits of the isoform ID printed to differentiate isoforms and their respective dot. See **Figure S3 f**or all isoforms expressed. Genes marked with an asterisk (*) indicate isoforms not recovered in unique-count analysis alone, which may reflect read counts split between multiple similar isoforms or insufficient full-length reads to resolve isoform ambiguity. Circle size indicates the percentage of cells in each cluster expressing the marker, and circle color indicates fold change from average expression. Transcript type is marked above each transcript (C = Canonical Protein; P = Alternative Protein; N = nonsense-mediated decay; RI = Retained intron, noncoding, U = Unknown coding status). (**d-h**) Gene-level UMAPs of aggregate marker expression for (**d**) T cells (*CD3D*, *CD3E*, *CD3G*); (**e**) Natural Killer (NK) cells (*GZMB*, *IL2RB*, ITGAM, *KLRF1*); (**f**) B cells (*CD19*, *CD22*, *CD79A*, *MS4A1*); (**g**) Monocyte-derived cells (*CD33*, *CLEC7A*, *FCGR2A*, *LILRB4*); and (**h**) Megakaryocytes (*GP1BA*, *ITGA2B*, *MPL*). (**i-m**) Isoform-level UMAPs of aggregate marker expression for (**i**) T cells, (**j**) NK cells, (**k**) B cells, (**l**) Monocyte-derived cells, and (**m**) Megakaryocytes.

We then assessed isoform expression differences by cell-type across marker genes. To understand biology at the isoform level, we first characterized the landscape by defining cell types at the RNA isoform level (i.e., marker isoforms) rather than solely at the gene level (i.e., marker genes). The ultimate goal is to eventually determine which specific protein isoforms drive cellular biology for each cell type and state; this will have important implications for molecular studies (e.g., those that rely on antibodies).

As expected, marker genes and their individual isoforms were either enriched in or specific to their respective cell types (**Fig. 3c**), but we were surprised by the number of unique isoforms expressed by each marker gene. Of the 18 primary cell-type marker genes used in this study, 15 actively expressed multiple unique isoforms, where half expressed ≥5 isoforms and three expressed ≥10 (**Figure S3, S6**; **Supplemental File 2)**. Additionally, there were two marker genes (*CD3G* & *GZMB*) that are considered cell-type specific, but each had an isoform enriched in a different cell type (megakaryotes; **Fig. 3c**). *CD3G* is a canonical marker for T cells that encodes the gamma subunit of the CD3 protein complex, a crucial receptor in T cell receptor signaling. *GZMB*, on the other hand, is a marker gene for NK cells and cytotoxic T cells that encodes the Granzyme B serine protease (a cytotoxic secretory molecule associated with the induction of apoptosis in other cells). Both genes exhibited enrichment of an alternative protein-coding isoform (*CD3G*: ENST00000392883; *GZMB*: ENST000005260004) in megakaryocytes. Whether the individual isoforms for each marker gene are performing a unique function (even if subtle) is not yet known, but understanding when, where, and why individual isoforms are expressed will be essential to better understanding gene function and cellular behavior.

We also generated aggregate “gene scores” for each set of marker genes, which clearly indicate each cluster (**Fig. 3d-h**). Specifically, the aggregate gene scores are generated by combining expression values for each set of marker genes into a single expression measurement to maximize cluster contrast (e.g., *CD3G*, *CD3D*, & *CD3E* for T cells). Similarly, we aggregated all marker isoforms into “isoform scores” to assign a cluster to their respective cell identity (**Fig. 3i-m**).

### Cell-type-specific isoforms of *GZMB* and *CD3G* suggest functional divergence or loss of function of non-canonical proteins from canonical functions

As proof of principle for the importance of understanding isoform differences, we compared the protein sequences for the *GZMB* and *CD3G* isoforms expressed in megakaryocytes to their canonical forms. The differences between proteins suggest potential functional divergence or loss of function.

For *GZMB*, the canonical protein sequences of (GZMB-201, ENST00000216341) and GZMB-204 (ENST000005260004; enriched in megakaryocytes) are identical through the 69^th^ amino acid (aa) in exon 3, after which GZMB-204 diverges, resulting in a much shorter protein sequence (247aa vs. 90aa). Structural studies into the catalytic mechanism underlying the Granzyme B function have determined the utilization of a catalytic triad (His59, Asp103, Ser198)^35^, all of which are required for canonical activation of the protease. InterProScan (v5.75-106.0)^36,37^ and Sherbrooke Alternative Protein Feature IdentificatoR (SAPFIR)^38^ analyses suggested that both isoforms contain a trypsin histidine active site encoded in exon 2 (**Fig. 4a,b**). GZMB-204, however, lacks exons 4 and 5 (**Fig. 4a,b**), which encode for the trypsin serine active site (197aa-208aa), a domain functionally responsible for targeting the peptide bond of a target protein^35^. Although not highlighted by InterProScan, the truncated GZMB-204 also lacks the Asp103 active site, which is critical for stabilization of the protonation state^35^. Because GZMB-204 is missing Asp103 and Ser198, the catalytic triad would likely not be formed and would be catalytically inactive. AlphaFold2-based predictions^39,40^ also suggest major differences in the protein structure, with two β-barrels in GZMB-201(**Fig. 4c**), and only one in GZMB-204 (**Fig. 4d**). As the juncture of these two β-barrels is where substrates are bound, we expect this change would significantly impact fuction. However, with an intact signal peptide, a region responsible for localization to the endoplasmic reticulum for packaging into the secretory system, the GZMB-204 protein could still be secreted, but it would not be able to degrade extracellular matrices and cell membranes. Why GZMB-204 is preferentially expressed in this rare population of circulating megakaryocytes, or why it is ever expressed, along with any potential function, is unknown. This is a clear example of the need to understand biology at the next level of isoforms.

**Figure 4.**
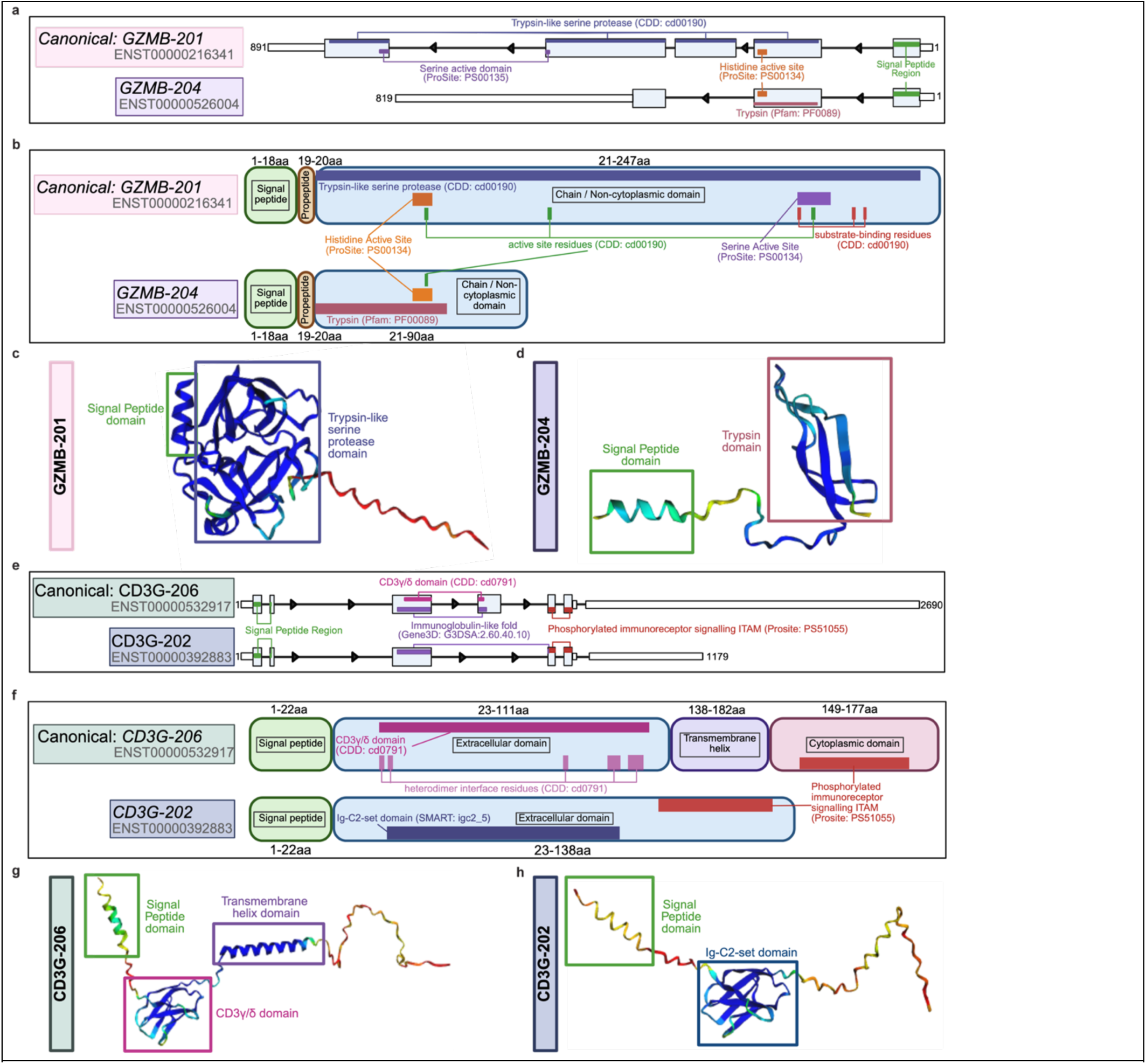
Protein-coding region significance and predicted structures of GZMB and CD3G isoforms. **(a)** GZMB-201 and GZMB-204 RNA isoform structure on the negative strand, based on results from SAPFIR^38^. Both isoforms encode an N-terminal signal peptide region, while GZMB-201 contains the entire trypsin-like serine protease domain with histidine and serine active sites. GZMB-204 lacks exons 2 and 5, removing the serine active domain and changing the specificity of the trypsin domain. **(b)** Domain architecture of the proteins resulting from GZMB-201 and GZMB-204 according to InterProScan^36,37^, including histidine and serine active sites and conserved residues. GZMB-204 exhibits loss of active site and substrate-binding residues in the mature functional polypeptide (chain region). **(c-d)** Predicted protein structure of GZMB-201 **(c)** and GZMB-204, missing the serine active site and reducing domain specificity **(d)**, modeled by AlphaFold, with confidence scores colored from blue (high) to red (low). AlphaFold output files for these structures including .PDB of the final structure can be found on Zenodo^49^, as described in **Data and Code Availability**. GZMB-204 exhibits the loss of protease domain integrity. **(e)** CD3G-206 and CD3G-202 RNA isoform structures, encoding domains including the signal peptide, immunoglobulin-like fold, and immunoreceptor tyrosine-based active motif (ITAM) site. Loss of exon 4 in CD3G-202 changes immunoglobulin-like specificity, replacing the CD3γ/δ domain with a less specific immunoglobulin-like fold. **(f)** Domain architecture of CD3G-206 and CD3G-202 proteins, including heterodimer interface residues (extracellular domain), transmembrane helix, and ITAM (cytoplasmic domain). Loss of the transmembrane helix region exhibited in CD3G-202 alters ITAM region localization relative to the membrane. **(g-h)** Predicted protein structure of CD3G-206 **(g)** and CD3G-202, missing the transmembrane helix domain **(h)**, modeled by AlphaFold using the protein sequences in **Supplemental Table 6**, with confidence scores colored from blue (high) to red (low) and highlighting isoform-specific changes in protein folding. AlphaFold output files including .PDB of the final structure can be found on Zenodo^49^, as described in **Data and Code Availability.**

*GZMB* and other immune response genes have been reportedly expressed in megakaryocytes during immune challenges, such as in sepsis or COVID-19^41^, so this may provide some context for why we have identified GZMB in our small megakaryocyte population. Additionally, secreted alternative proteins without canonical functionality have been identified in the transmembrane receptor RAGE^42,43^ and receptor kinase VEGFR1^44^, which act as dominant-negative decoys to bind substrates. Truncated non-receptor protein-coding isoforms that lose their canonical functional domains have also been reported as immune regulators; for example, the STING (TMEM173) protein produces isoforms lacking an N-terminal transmembrane domain, disrupting proper cellular localization and negatively regulating canonical STING signaling^45^. Additionally, NK cells showed enrichment for several GZMB isoforms with differing structures, including GZMB-203 (ENST00000415355) (**Figure 3C**) and GZMB-202 (ENST00000382540) (**Figure S3**). GZMB-203 retains a functional domain identical to that of the canonical isoform, but is missing a signal peptide. Therefore, this form of GZMB would be unlikely to be secreted and may be degraded by the proteasome. NK cells also express GZMB-202, an isoform missing exon 3, which contains Asp103, which stabilizes and activates the catalytic triad, leaving the protein likely to be catalytically inactive. While still likely secreted, GZMB-204 and -202 could contribute to the extracellular granzyme pool, limiting tissue damage without contributing to cytotoxic activity. Thus, potential functions of GZMB-204- and GZMB-202-derived proteins could be (1) to compete with canonical GZMB-201 in the secretory machinery, potentially slowing the protease’s release from NK cells; (2) to act as a decoy to bind substrates or target sites competitively, modulating protease activity in extracellular sites; or (3) nonfunctional.

We next examined *CD3G*, a canonical T cell gene, which also preferentially expressed a specific isoform. As expected, the canonical isoform (CD3G-206, ENST00000532917) is enriched in T cells, but an alternative protein-coding isoform (CD3G-202, ENST00000392883) is enriched in megakaryocytes. The canonical isoform (CD3G-206) contains a signal peptide region, an extracellular IgC1 domain (Conserved Domain Database^46^ (CDD): IgC1_CD3_gamma_delta), a transmembrane helix domain, and a cytoplasmic tail containing an immunoreceptor tyrosine-based activation (ITAM) motif (**Fig. 4e,f**). Based on InterScanPro and CDD, the protein sequence produced from the CD3G-202 isoform matches a sequence motif associated with a generic immunoglobulin-like fold, appearing to be missing the heterodimer interface (CDD: cd07691) and Ig strands A, C’, and D (**Fig. 4f**). CD3G-202 is missing the transmembrane domain, and the ITAM motif is in the extracellular domain, indicating this may be secreted rather than a membrane-bound form of the protein. These predictions were supported by results from DeepTMHMM (v1.0.44)^47^, which predicted a transmembrane helix in CD3G-206 (aa 118-136), but none in CD3G-202. The physical structure predicted by AlphaFold2 appears to support these conclusions (**Fig. 4g,h**), based on a well-defined structure in CD3G-206; however, the predicted structure of CD3G-202 lacks the predicted transmembrane helix. As postulated for GZMB-204 and - 202, the release of a secreted form of *CD3G* may function as a decoy isoform or as an immune regulator, a biological mechanism that has been proposed for other genes as potential biomarkers and treatments in diseases such as SARS-CoV-2 infection^48^. These examples underscore the importance of isoform-level analyses and are critical for understanding endogenous modulation of immune responses in humans.

### T cell subtype characterization demonstrates subtype-dependent alternative splicing and isoform usage

With the added resolution provided by the large number of cells and the use of long reads, we next focused on T cells for subtype clustering. T cells are the most abundant cellular population in our dataset and are known to be biologically heterogeneous, exhibiting diverse roles in the adaptive immune system^50^, making them a logical choice to use to evaluate the heterogeneity in our data as a proof-of-principle.

We reanalyzed the T cell population using a Leiden resolution of 0.26 and identified three major subtypes based on canonical marker genes (**Table S4; Fig. 5a,b**): effector CD8^+^ T cells (*CD8A*, *CD8B*, *KLRB1*, *GATA3*, *CCL5*), effector CD4^+^ T cells (*CD4*, *IL2RA*, *TNF*, *AHR*, *GATA3*), and memory T cells (*TCF7*, *CCR7*, *SELL*). The CD8^+^ effector cluster was identified as such, as opposed to a general CD8^+^ T cell population, due to their expression of known effector and activation genes, *GATA3*^40^ and *CCL5*^51^. We identified CD4^+^ and CD8^+^ expression within portions of the memory T cell cluster; however, resolution was insufficient to separate these into sub-groups (**Fig. 5c**). In some genes (*AHR* and *KLRB1)*, only a single isoform was expressed, though *AHR* has 7 known isoforms. Overall, however, we identified that most of our T cell subtype marker genes expressed multiple isoforms in our dataset, with 6 expressing at least 5 isoforms, and 2 of these genes expressing over 10 isoforms (**Supplemental File 2**). It is possible that alternative splicing may be dynamically regulated across cell-type or activation state, resulting in preferential isoform expression within a specific context.

**Figure 5.**
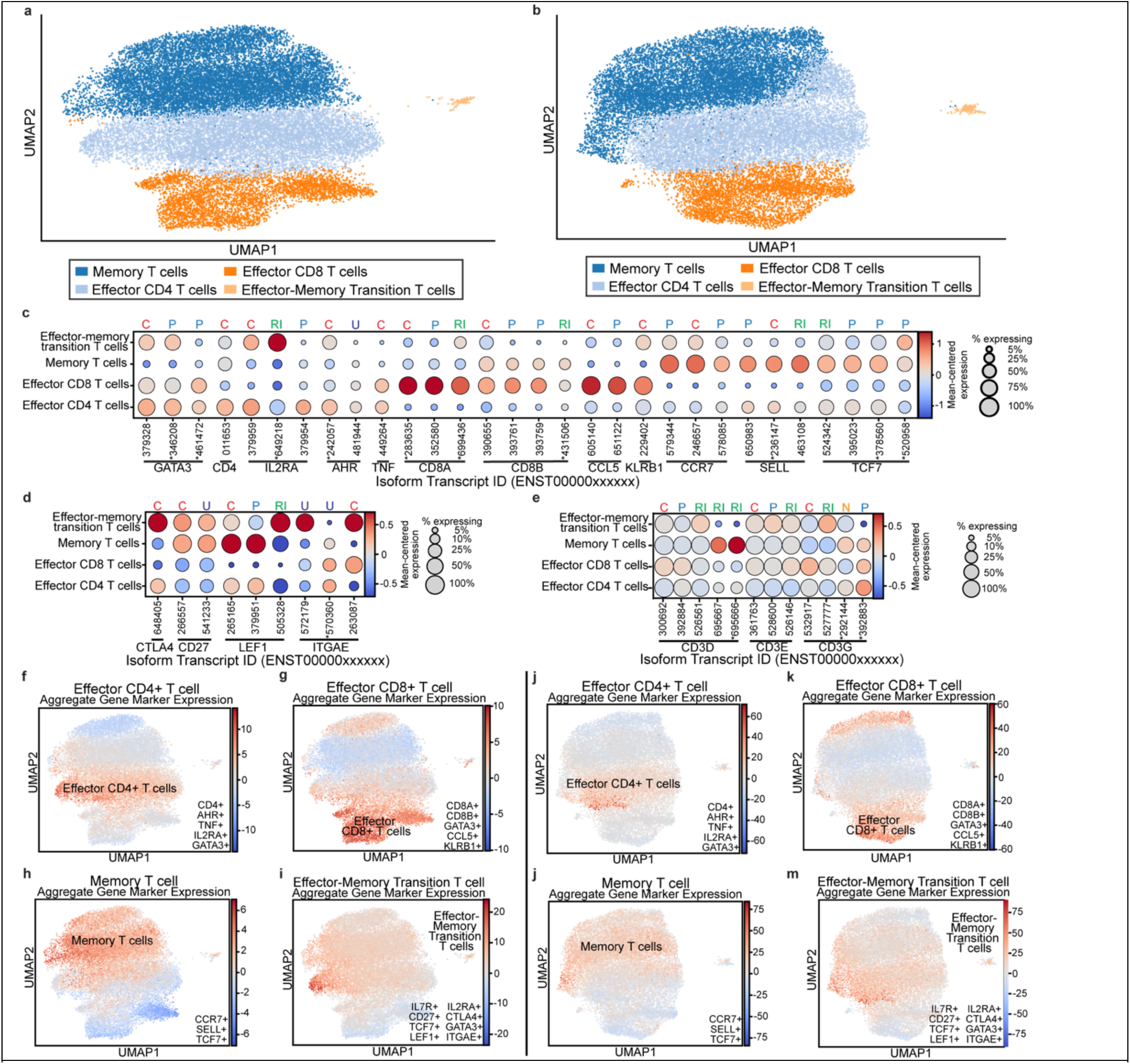
T cell subtype clustering by canonical markers at the gene- and isoform-level. **(a)** Gene-level UMAP projection of T cell subtype clusters. **(b)** Isoform-level UMAP projection of T cell subtype clusters. **(c-e)** Dot plots of up to three representative isoforms per gene, selected as the most highly expressed isoforms by raw counts (**Fig. S4**), and including any isoforms enriched in a non-canonical cell-type. X axis represents isoform transcript ID (ENST00000xxxxxx) with the last 6 digits of the isoform ID printed to differentiate isoforms and their respective dot. See **Supplementary** Figure 6 for all isoforms expressed. Colored boxes correspond to cluster-defining markers. Genes marked with an asterisk (*) indicate isoforms not recovered in unique-count analysis alone, which may reflect read counts split between multiple similar isoforms or insufficient full-length reads to resolve isoform ambiguity. Circle size indicates the percentage of cells in each cluster expressing the marker; circle color indicates fold change relative to average expression. Transcript type is indicated above each transcript (C = Canonical Protein; P = Alternative Protein; N = nonsense-mediated decay; RI = Retained intron, noncoding, U = Unknown coding status). Plots are split up by **(c)** Main T cell-type markers (Memory T cells, Effector CD4 and CD8 T cells), **(d)** markers specific to effector-memory transition T cells, and **(e)** general T cell markers. **(f-i)** Gene-level expression aggregation on the single-cell level in **(f)** CD4+ effector T cells, **(g)** cytotoxic/effector CD8+ T cells, **(h)** memory T cells, and **(i)** effector-memory transition T cells. **(j-m)** Isoform-level expression aggregation on the single-cell level in **(j)** CD4+ effector T cells, **(k)** cytotoxic/CD8+ effector T cells, **(l)** memory T cells, and **(m)** effector-memory transition T cells.

*CD8B* is an intriguing case, where a non-canonical isoform (CD8B-206, ENST00000431506) encodes a nearly identical protein to that of the canonical isoform (CD8B-203, ENST00000390655), except for a single amino acid change: a phenylalanine (F) in CD8B-203 is replaced by a leucine (L) in CD8B-206 within the cytoplasmic tail region (208aa). Notably, CD8B-206 is more highly expressed in memory T cells than in cytotoxic T cells (**Fig. 5c**). Although the protein-coding sequences are nearly identical, these two mRNA isoforms differ in the 3’ untranslated region (3’UTR), where CD8B-206 completely lacks a 3’UTR, according to UTRdb^52^ (**Figure S4**). Studies suggest the 3’ UTR region regulates mRNA localization, translation, and stability, and shows cell-type-specific patterns of 3’UTR length changes upon activation of signaling pathways^53^. 3’UTRs often contain motifs bound by RNA-binding proteins and microRNAs that mediate RNA decay or transcriptional regulation; therefore, the loss of this region in CD8B-206 may indicate enhanced stability and reduced likelihood of regulation. The presence of a *CD8B* mRNA isoform lacking a 3’UTR in CD8^+^ memory T cells may help sustain expression of this protein in memory T cells, which tend to be long-lived. Consistent with this, prior gene-level studies have shown 3’UTR shortening in proliferating T cells, with CD8^+^ effector memory T cells exhibiting the most pronounced shortening^54^. These isoform-specific patterns would not be detectable at the gene level or in bulk long-read sequencing, highlighting the unique value of long-read single-cell approaches in providing crucial information on transcriptional patterns in a cell-type-dependent manner.

In addition to these three main clusters, we identified a smaller cluster of cells adjacent to the main T cell population that expressed a mixture of effector/activation markers (*IL2RA, CTLA4, GATA3*) and markers for memory persistence, stem-like, and transition states (*IL7R, CD27, TCF7, & LEF1;* **Fig. 5d**), with isoform specificity. These cells do not express Natural Killer-T cell markers (*NCAM1, GZMB*), exhibit unremarkable doublet scores, and show low mitochondrial gene expression (**Fig S5**), which supports their validity as a distinct cluster, rather than an artifact. We conclude this group is likely a transitional effector-memory T cell cluster. The expression of *ITGAE* (CD103; **Fig. 5d**), also enriched in this cluster, is associated with tissue-resident T cells and may indicate that these cells have tissue-resident memory potential.

A distinct cell-type-specific pattern emerged in our effector-memory transition cluster, where marker genes exhibited limited numbers of isoforms compared to other T cell subtypes. Unlike other clusters that typically highly express most isoforms of their defining marker genes for that cluster, this transition cluster only highly expressed 1-2 isoforms per marker gene. For example, while *IL2RA* is highly expressed in effector CD4^+^ T cells among all isoforms, but only the canonical isoform (IL2RA-201, ENST00000379959) and an isoform with a retained intron (IL2RA-206, ENST00000649218) was highly expressed in the transition cluster (**Fig. 5c**). Similarly, marker genes of this transition state showed preferential expression of an isoform annotated as noncoding in *LEF1* (LEF1-210; ENST00000505328), instead of the diverse array of isoforms seen in other clusters (**Fig. 5d**). *ITGAE* expression was likewise enriched in this cluster for the canonical form (ITGAE-201; ENST00000263087) and one with an unknown coding sequence (ITGAE-207; ENST00000572179) (**Fig. 5d**). This selective isoform usage may reflect a specialized transcriptional control required for the transitional state between effector and memory conditions.

Interestingly, we also noticed that two large non-coding isoforms of CD3D are enriched in memory T cells (**Fig. 5e**). Although the memory subgroup exhibits overlapping expression patterns on the aggregated gene level (**Fig. 5f-i**), our isoform-level analyses (**Fig. 5j-m**) exhibit clear expression differences, highlighting the importance and utility of isoform-level studies to better understand the underlying biology.

### New isoforms exhibit distinct cell-type specificity in expression and structural changes from known isoforms

To identify trends of expression patterns for our newly discovered isoforms from new and known genes, we examined inter-gene clustering for our new isoforms to see which exhibited similar expression patterns (**Fig. 6a,b**). Several of our new genes demonstrated enrichment in specific cell types, with dendrograph clustering appearing fairly distinct for genes enriched in NK cells, megakaryocytes, B cells, monocyte-derived cells, and effector-memory transition T cells (**Fig. 6a**). Although some genes were enriched in the memory, effector CD4^+^, and effector CD8^+^ T cell clusters, very few exhibited strong, specific enrichment.

**Figure 6.**
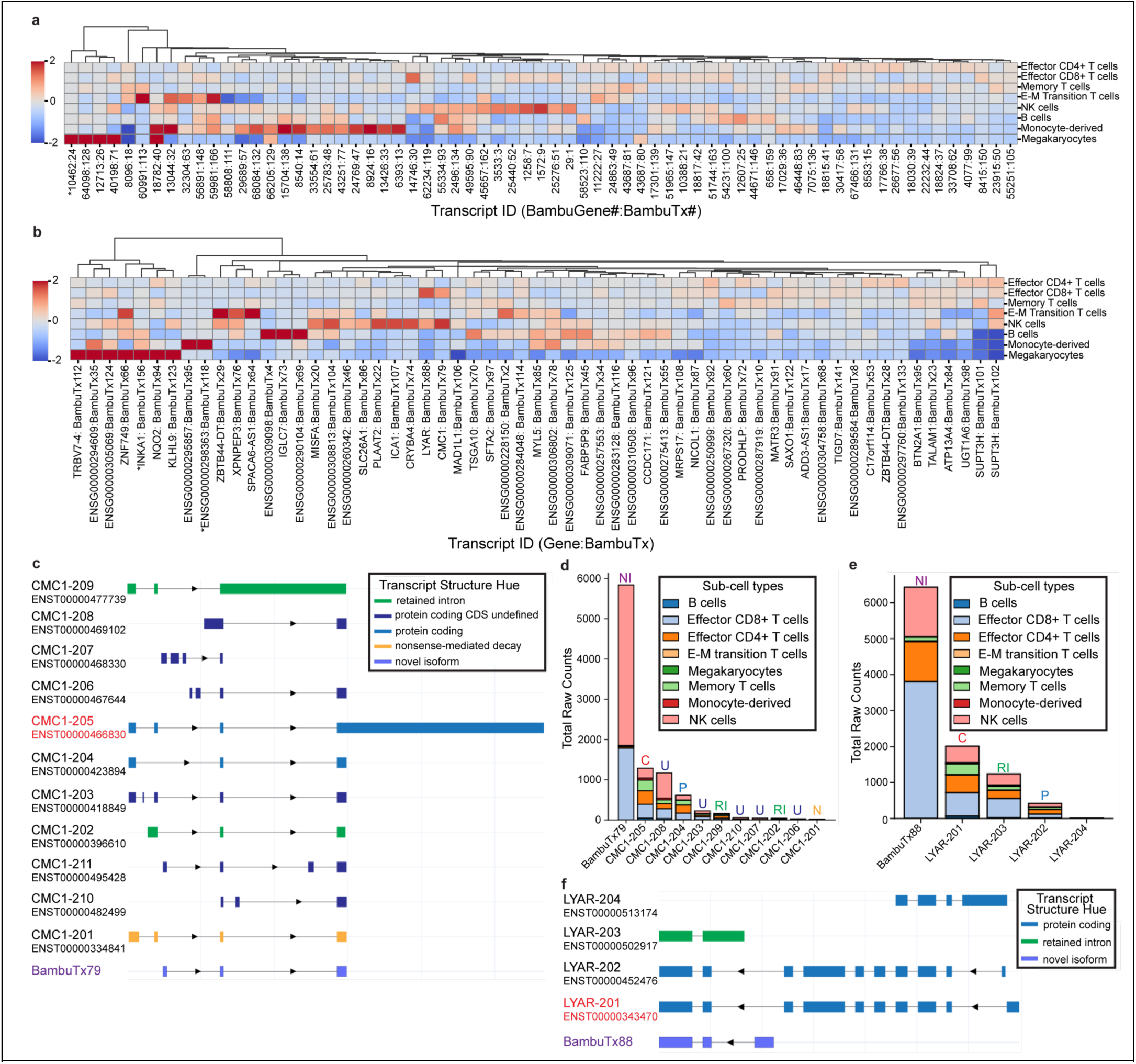
New isoforms from new and known genes exhibit cell-type specific expression patterns and distinct isoform structure. Isoform-level diversity of previously unannotated (a-b) Heatmaps showing normalized expression (Z-score) of novel isoforms across major cell-types and T cell sub-types in (a) previously unannotated genes and (b) known genes. (c) Transcript structure models generated with *RNAPysoforms* for *CMC1* isoforms, including the novel transcript BambuTx79 (indicated in purple) and canonical isoform CMC1-205 (indicated in red). (d-e) Raw expression of *CMC1* (d) and *LYAR* (e) isoforms across major cell-types and T cell sub-types, showing that novel transcripts are the most highly expressed variants in both genes. (f) Transcript structure models for *LYAR* isoforms, including novel BambuTx88.

Of our 58 known genes in which we identified a new isoform, 26 were RNA genes and two were pseudogenes, while the remaining 30 were protein-coding genes. Among these, two of the RNA genes (*TALAM1*, *ADD3-AS1*), though noncoding, have reported regulatory roles in gene expression and cellular signaling, sometimes affecting their neighboring genes, such as in the case of *TALAM1*^55,56^. According to GeneCaRNA^57,58^ (v5.25), five of the RNA genes are known antisense transcripts, a role that is predicted to regulate the sense gene^59,60^. Similarly, two are known sense intronic transcripts, which originate within a gene’s intron and can be precursors for small non-coding RNAs or can regulate its host gene^61,62^. These observations highlight the utility of RNA genes often ignored in genomics research, which require further study to truly understand their role.

Intriguingly, for eight known genes, the most highly expressed isoform was the new transcript (**Supplemental File 2**). For two of these genes, *SUPT3H* and *ZBTB44-DT*, two new isoforms were identified, and both had greater overall raw counts than known isoforms (**Supplemental File 2**). These new isoforms, however, did not show the same expression patterns, despite being in the same gene. Although the new isoforms for *SUPT3H* (BambuTx101 and 102) clustered together on the dendrograph (**Fig. 6b**) and showed consistent enrichment in CD4^+^ and CD8^+^ effector T cells, BambuTx102 was also enriched in NK cells and effector-memory transition T cells, while BambuTx101 was enriched in Memory T cells. Additionally, in *ZBTB44-DT*, BambuTx28 showed distinct isoform expression patterns from its sister isoform, BambuTx29, diverging on the dendrograph by several clades and showing enrichment in different cell-types (**Fig. 6b**). BambuTx28 demonstrated enrichment in effector CD4^+^ T cells, while BambuTx29 was enriched in effector-memory Transition T cells and NK cells.

Another gene that exhibits a new transcript as its most highly expressed isoform is *CMC1*, which encodes a highly conserved mitochondrial protein that contributes to the construction of cytochrome c oxidase (complex IV), the last stop in the electron transport chain^63^. Transcript modeling of *CMC1* isoforms using *RNApysoforms*^64^ confirmed a novel exon confirmation in BambuTx79, showing a structure maintaining an intact exon 2 and 3, but a missing exon 1 and truncated exon 4 (**Fig. 6c**). Interestingly, BambuTx79 also exhibits a distinctive expression pattern from the other isoforms within the gene, displaying enrichment in CD8^+^ T cells and NK cells alone, whereas the other isoforms of *CMC1* have a broader expression pattern across cell-types (**Fig. 6d**). It appears that there is still more to learn about *CMC1*, in part since we notice many transcripts that are marked as protein-coding, but whose protein-coding sequences are unknown (**Fig. 6d**). We cannot definitively predict this isoform’s function since we do not know whether it is protein-coding and therefore what the amino acid sequence is. However, it exhibits enrichment in cytotoxic cell-types, cells whose activation is associated with increased metabolic activity^65^, so a novel isoform in this gene may play a role in this change.

Another gene with a new highly expressed isoform is *LYAR* (**Fig. 6e**), a protein-coding gene with a broad range of roles, including transcriptional regulation, DNA-binding transcription factor binding activity, and regulation of immune responses (e.g. NFκB) ^66,67^. Our new isoform of *LYAR*, BambuTx88, exhibits similar expression patterns to its other isoforms, with most raw counts originating from effector CD8^+^ T cells, with the next largest proportion of reads originating from NK cells, then effector CD4^+^ T cells (**Fig. 6e**). When we modeled the *LYAR* isoform structures, we noticed that the end structure of BambuTx88 matched with exons 9 and 10 seen in LYAR-201 and LYAR-202 but appeared to have an exon 8 that does not align with any other isoform in that location, as the others have a larger intronic region before exon 8, supporting its novelty (**Fig. 6f**). Since the functions of *LYAR* are wide-ranging, this newly identified isoform could have a myriad of different functions that would need to be examined experimentally.

Together, these analyses reveal several previously unannotated isoforms that are not only highly expressed but also have distinct isoform structures from previously annotated isoforms and exhibit defined cell-type-specific expression patterns. Future and deeper studies will be necessary to clarify the functions of the individual isoforms across these important genes.

## Discussion

### Long-read scRNA-seq unlocks new biological insights in human PBMCs

Gene-level scRNA-seq has proven an important tool for understanding transcriptomic profiles in disease and for characterizing heterogeneity across a range of biological processes^68^, including immune function^69^. However, isoform-level variation within a single gene can vastly impact protein structure and functionality, a detail that is crucial to understanding the underlying biology that has been overlooked because of short-read approaches. Our adapted PIPseq long-read single-cell approach, including our bespoke read recovery pipeline, bridges this gap by directly capturing canonical and non-canonical isoform expression in a cell-type-specific manner, uncovering potential functional variation that cannot be identified via standard gene-level approaches.

In PBMCs, we identified several striking and unexpected trends, including that most of our marker genes simultaneously expressed multiple isoforms in the same cell type (in some cases over 10), while other marker genes appeared to preferentially express different isoforms in different cell types. Of the 35 marker genes used in this study, only six expressed just a single isoform (*TNF, KLRB1, CTLA4, MPL, LILRB4, GP1BA),* whereas the rest averaged ∼5.4 isoforms (**Fig S3, S6, Supplemental File 2**). Notably, in megakaryocytes, we identified enrichment for two non-canonical isoforms: (1) an abbreviated *GZMB* isoform missing a domain required for its canonical proteolytic activity, and (2) a secreted form of *CD3G*; both isoforms are overlooked in gene-level data. Additionally, we noticed that for many of our marker genes (*ITGAM, FCGR2A, CCR7, SELL, TCF7*, & *ITGAE*) and genes with newly identified isoforms, the most abundant isoform was not the canonical transcript (**Supplemental File 2**), underscoring the sometimes-arbitrary nature of canonical definitions and the importance of isoform-level studies to understand the underlying biology. These findings reveal the extensive isoform diversity of immune genes and show how long-read data can provide functional insight into different proteins from a single gene.

While not directly validated here, numerous examples exist demonstrating that different isoforms for the same gene provide valuable insights into human health and disease. For example, isoform-level differences have shown clinical relevance in Alzheimer’s disease, where a tissue-specific isoform of tau, a characteristic pathological marker, outperforms total tau as a predictive blood biomarker of the disease^70^. Similarly, *GZMB* has been identified as a potential drug target to slow brain aging^71^, showing up in genetic association studies. It has also appeared in neurons and T cells from individuals with neurodegenerative disease and may be involved in the apoptosis of diseased or injured cells, according to murine and in vitro studies^72,73^. This emphasizes the clinical potential of isoform studies to identify specific biomarkers.

### Long-read single-cell approaches reveal functional transcriptomic and proteomic differences overlooked in conventional gene-level analyses

While gene-level RNA sequencing provides important insights into trends regarding gene expression in disease, due to the many factors that drive protein abundance and function (e.g. alternative splicing, translation efficiency, protein folding), it often does not reflect the proteins generated. A major criticism of transcriptomic studies has been their poor correlation to protein-level abundance and function, which limits biological and clinical translatability. Isoform-level transcriptomics may help to bridge the gap between transcriptomics and proteomics, allowing disease studies to be more efficient and increasingly translatable.

mRNA diversity within individual genes can be vast, with isoforms exhibiting a range of protein-coding potential. Individual isoforms may exhibit truncated UTRs, premature stop codons, missing exons with crucial functional domains, or absent signal peptides required for proper cellular localization. By resolving individual isoforms using long-read sequencing, we can better determine which isoforms are expressed across tissues and cell types, potentially providing more accurate predicted protein expression. Additionally, combining isoform analyses with open-source computational tools (e.g. InterPro, AlphaFold2, SAPFIR), enables prediction of differences between structure, function, localization, translational efficiency, and their combined impact on cell types of interest. Our ability to predict changes in functionality on the protein level based on the isoforms expressed, as shown in *GZMB*, may allow us to better predict how elevated gene expression in disease reflects changes in function, and which isoforms would be more likely to be degraded. By performing these analyses, our understanding is enhanced not only regarding the function of protein isoforms within a single gene but also how these isoforms are regulated. Of course, all results from such studies would need to be validated experimentally.

As discussed in our *CD8B* isoform analyses, proteins that have seemingly the same protein sequence may exhibit different expression patterns between cell types. Similarly, our identification of a truncated *GZMB* isoform and secreted *CD3G* isoform suggests that isoforms within the same gene may have different regulatory patterns and functions. These examples highlight the new hypotheses afforded by long-read scRNA sequencing; these tools provide mechanistic insights into the relationship between isoform structure and protein function, addressing a critical gap in our understanding of transcriptomic regulation.

Lastly, we identified several isoforms previously unannotated that were not only expressed in a cell-type-specific manner but were the predominant isoform within their gene. These new isoform analyses support the notion that long-read single-cell sequencing combined with isoform discovery informs our knowledge of the isonome on a cell-type-specific basis, which would remain hidden in gene-level analyses.

### Limitations and Future Improvements

Despite the large number of reads generated per sample, the primary limitation of this study is read depth, given the large number of cells captured (three R10 flow cells across 30,000 cells, per sample). While our achieved read depth was sufficient to analyze the more highly expressed marker genes and associated isoforms, it was not sufficient for broader analyses across genes and isoforms with lower expression. Due to the high cell count targeted and subsequent low read depth per cell, nearly all genes and approximately half of isoforms (∼99% of genes, ∼48.7% of isoforms, **Table S2**) exhibited technical zero inflation, requiring modeling using the zero-inflated negative binomial (ZINB) distribution. According to similar studies that obtained greater depth, 10,000-20,000 reads per cell would be considered sufficient for long-read analyses with normal corrective measures^10^. However, the rate of zero-inflated genes and isoforms in these datasets that may be better modeled by ZINB would be worthy of investigation.

To fully leverage the benefits of this method, future studies using this methodology should aim for greater sequencing depth, either by using fewer cells or more flow cells. In future experiments using the same number of flow cells, we recommend reducing the number of cells targeted to 5,000, which we predict will result in a targeted depth of ∼15,000 reads per cell when utilizing 3 flow cells per sample. These recommendations will change if throughput per flow cell increases over time. Our future estimated depth is in line with our initial estimate of a sufficient number of reads for our planned analyses and has previously been sufficient for single-cell long-read analyses in human tumors^10^.

Our knowledge of the isonome and its impact on the proteome remains limited, as evidenced by the genes who produce multiple protein-coding isoforms whose translated sequences and functions are uncharacterized. While previous long-read scRNA-seq have primarily focused on cataloging isoform diversity, our work extends these efforts by assessing structural differences between isoforms of cell-type-specific markers and linking these to cell-type enrichment and potential functional differences across immune populations. To fully understand the influence of distinct isoforms on biology, researchers must continue to explore innovative approaches to capturing these complex signatures and testing them experimentally. The adapted microfluidic-free approach and accompanying computational pipeline described here provide a benchtop-accessible preparation for long-read single-cell sequencing, generating an extensive view of the isonomic landscape of PBMCs with the greatest number of reads and cells produced in a single-cell long-read study.

## Methods

For a full list of reagents and materials used, see **Supplemental Information**.

### Workspace preparation and cleanup

Unless otherwise stated, before and after each day of the protocol, UV light was used in the cell culture hood for 30 minutes and workspace and micropipettes were cleaned with a 70% ethanol solution and RNase decontaminant solution (*Invitrogen,* #AM9782). Since all steps were performed within an open system, it was imperative to perform all possible steps in this sterile environment to minimize RNase contamination.

### Day 1: PBMC isolation

PBMCs for this proof-of-principle study were isolated from a voluntarily donated blood sample from a 43-year-old healthy Caucasian male. A total of 20mL of venous blood was collected into two 10mL K2EDTA anticoagulant tubes (*VWR* #BDAM367525) and transported to the laboratory at room-temperature. Tubes were placed on a rocker for approximately 10 minutes to resuspend blood cells. Ficoll-Paque PLUS (*Millipore Sigma* #17-1440-020) was prepared according to directions from manufacturer. PBMCs were isolated using an optimized version of the Sepmate protocol (V2.0.0; *Stemcell Technologies*, #85450). Briefly, room-temperature Ficoll-Paque PLUS was mixed by inversion, and 15mL was aliquoted into the lower section of a 50mL Sepmate Tube. Whole Blood was diluted 1:1 with room-temperature PBMC Medium (1xPBS + 2% FBS) and layered onto the Ficoll-Paque PLUS in the Sepmate tube before centrifugation (20 min, 1200xg, 20°C, max acceleration and break). The top layer (containing PBMCs and plasma) was decanted into a new tube and brought to a volume of 50mL using PBMC Medium. Samples were centrifuged (10min, 300xg, 4°C, acceleration 9, break 9) and the supernatant was aspirated to 5mL to remove the plasma layer. The bottom of the tube was gently flicked to break up the cell pellet, and PBMC Medium was added to 50mL and centrifuged (8 min, 200xg, 4°C, acceleration 9, break 1) before removing as much supernatant as possible without disturbing the cells. Red blood cell lysis was performed for 5 minutes according to manufacturer’s instructions (*Invitrogen* #00-4300-54), then washed with 15mL 1xPBS (6 min, 300xg, 4°C, max acceleration and break).

### PIPseq scRNA-seq preparation

We modified the PIPseq T20 V4.0PLUS Single Cell RNA Kit User Guide (Revision 8.9) protocol for adaptation to long-read sequencing. Vortexing to mix cell suspensions and sample mixtures is abundant in this protocol; however, since we have found that vortexing is a major source of shearing, we minimized vortexing after cDNA isolation from PIPs to maximize fragment length. Unless otherwise stated, any time a wash occurred or a cell pellet was formed from centrifugation, gentle tapping of the tube was used to dissociate the pellet before resuspending and pipetting to mix gently and slowly with a wide-bore pipette tip. The PIPseq protocol was performed over 3 days with our noted alterations, as indicated below. The base protocol includes 5 stop points ranging from overnight to 96 hours where users can stop if needed; however, we chose to utilize only 3 and perform on consecutive days to minimize RNA/cDNA degradation from extended periods of storage. We also did not store cDNA at temperatures below 4°C until after sequencing to avoid shearing during multiple freeze-thaw cycles.

As recommended in the protocol, when it was noted that we “spin down” samples, this was done on a benchtop minifuge reaching speeds up to 20,000RPM. All plastic tubes and reagents described in the PIPseq sections below were included in the kit, unless noted. A swinging bucket rotor was used for PIPseq to minimize damage to cells, RNA, and cDNA, as suggested. Unless otherwise mentioned, all reagents and plastics in the PIPseq section of the protocol originated from one of the following kits that make up the T20: (PIPseq T20 3’ Single Cell RNA Ambient Kit v4.0PLUS (*Fluent* #FBS-SCR-T20-4-V4.05-1), PIPseq T20 3’ Single Cell RNA 4°C Kit v3.0 or 4.0 (*Fluent* #FBS-SCR-T20-4-V3&V4-2), PIPseq T20 3’ Single Cell RNA 20°C Kit v4.0PLUS (*Fluent* #FBS-SCR-T20-4-V4.05-3), PIPseq T20 3’ Single Cell RNA -80°C Kit v4.0PLUS (*Fluent* #FBS-SCR-T20-4-V4.05-4), PIPseq T20 3’ Single Cell Consumables Kit v4.0PLUS (*Fluent* #FBS-SCR-T20-4-V4.05-6)).

### Day 1: PIPseq—Cell Preparation

Immediately after isolating PBMCs and washing cells in 1xPBS, rather than performing a single wash in warmed Cell Suspension Buffer, samples were washed twice in 1mL PIPseq Cell Suspension (one at room temperature, one at 4°C) to ensure the gentle removal of reagents inhibitory to PIPseq (e.g. FBS). Cells were centrifuged at a setting we optimized for most cell-types (6min, 300xg, 4°C) and supernatant was aspirated carefully. We recommend that users tailor the temperature and time of centrifugation to their cell type of interest based on the optimal conditions for survival. A swinging bucket with a 1mL adapter is gentler on the cells during this section, aiding in better viability. Between washes, the cell pellet was dissociated by gently tapping the bottom of the tube and cell suspension was mixed with slow wide-bore pipetting reduce mechanical strain. We found that mixing with a wide-bore tip alone, as recommended in the base protocol, was insufficient to obtaining a uniform cell suspension. Dissociating the pellet with gentle tapping replaced the recommendation need to increase pipetting force, which may increase mechanical strain on cells and lysis. After washes, the cell pellet was resuspended in 500µL of PIPseq Cell Suspension Buffer (instead of 200-400uL, again to reduce the effect of any residual FBS), as described above (gently tapping and wide-bore mixing). Cells were counted using trypan blue (*PromoKine* #PK-CA902-1209) to determine viability and cell concentration (**Formula 1**). Since PIPseq has an estimated 50% capture rate, to target 30,000 cells for sequencing, we identified that a total of 60,000 cells would be needed in a volume of 8uL to begin the PIPseq protocol. Thus, our Targeted Concentration was 7,500 cells/µL for loading. The Targeted Concentration was then used to determine the volume of Cell Suspension Buffer to add to the sample for the desired concentration (**Formula 2**). If the value produced from **Formula 2** was negative, we removed that volume of supernatant rather than adding.

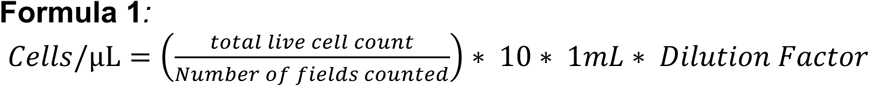

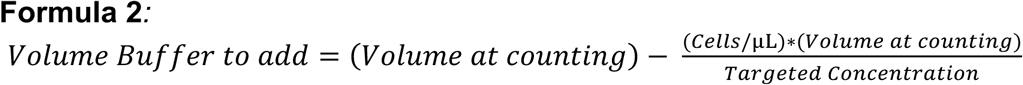

### Day 1: PIPseq—Cell Capture and Lysis

To adjust cell suspension concentration, samples were centrifuged at our optimized setting (6 min, 300xg 4°C, max acceleration and break). Volume was adjusted according to the value calculated with **Formula 2** and placed on ice. Cell counts (**Formula 1**) were repeated to ensure accurate concentration and >90% viability before loading. As in the protocol, the cell suspension was gently pipette-mixed with a wide-bore tip before transferring 8µL of cell suspension and 2µL of SUPERase•In RNase Inhibitor (20U/µL) (*Invitrogen* # AM2696) into PIP mixture. The cell:PIP mixture was very slowly pipette-mixed using a standard-bore P200 pipette as recommended by the protocol. We ensured that very slow, even strokes were used, that the emulsion is released along the inner side wall of the tube, and that the pipette was not pushed to the second stop until mixing is complete. Although this protocol recommends the use of a multi-channel pipette while mixing the cell:PIP mixture, we found using a single-channel pipette to better reduce the amount of bubbles and foam produced during this step, a crucial problem given how viscous the solution is. When mixing the cells into the PIPs, This solution is extremely viscous and prone to foaming. Although a small number of bubbles are alright, if excessive foaming occurred, we briefly centrifuged on the minifuge to remove. Excessive centrifugation can cause removal of cells from PIPS and was used sparingly. On the Fluent Website, a video protocol of this step was previously included at the beginning of the Cell Capture step in the “Legacy: PIPseq T20 V4.0PLUS Single Cell RNA Kit User Guide” (Revision 8.9). This video tutorial can now be found on the Illumina website under “Illumina Single Cell 3’ RNA Prep, T20 Workflow” (https://support.illumina.com/sequencing/sequencing_kits/illumina-single-cell-prep/training.html). We highly recommend users watch this video and take note of the images throughout the protocol before applying the changes made for long reads.

The remainder of this section follows the standard v4.0PLUS PIPseq protocol as written. Briefly, Partitioning Reagent was added before vortexing on the PIPseq rotating vortexer. After vortexing, the bottom phase of the sample (Partitioning Reagent Waste) was removed to the lowest line of the blue PIPseq tube stand. Partitioning reagent and Chemical Lysis Buffer 3 (CLB3) were combined and vortexed before adding to the PIP emulsion. The PIP emulsion was then gently inverted to mix ten times. Cells were lysed using setting A (**Table 2**) in the PIPseq Dry Bath (*Fluent* #FBS-SCR-PDB) and held at 15-25°C overnight:

**Table 2.**
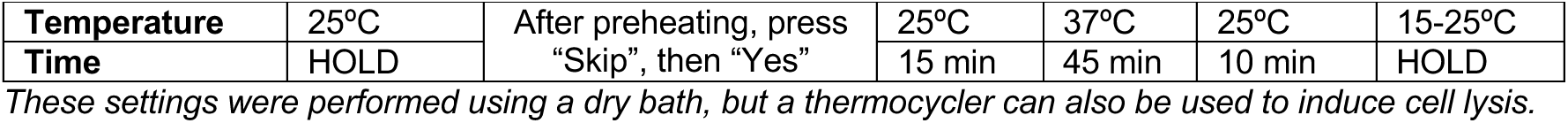
PIPseq Dry Bath Setting A parameters for heat-induced cell lysis.

### Day 2: PIPseq—mRNA isolation

This section was performed according to the base protocol. Briefly, after incubation, the bottom phase was removed to the middle line on the blue PIPseq tube stand. Breaking Buffer and De-partitioning reagent were both added to the sample and the sample was inverted gently ten to twenty times, then spun down briefly. Although the PIP formulation is designed to protect samples from damage during vortexing, our experience adapting this protocol suggests that this protection is inconsistent for long strands. The bottom, pink-colored waste layer was removed carefully and PIPs were transferred to an aliquot of Washing Buffer. A total of three washes were performed as follows: 1) The pellet was washed with 12mL Washing Buffer, 2) cell pellet was dispersed with gentle tapping and tube inversion, 3) centrifugation with a swinging bucket rotor (2 min, 750xg, 4°C, low break and max acceleration), 4) Slow aspiration of supernatant without disturbing pellet. The PIP mixture was transferred to a pre-weighed 1.5mL tube, weighed, and the volume to remove to normalize the mixture to 250µL was calculated (**Formula 3**). Samples were spun down ∼30 seconds before carefully removing the determined volume from **Formula 3**.

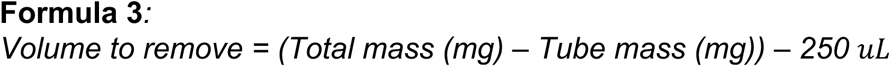

### Day 2: PIPseq—cDNA Synthesis

A master mix was prepared with the following recipe per sample: 232.2µL RT Additive Mix, 21.6µL TSO, 16.2µL RT Enzyme Mix. The pellet was dissociated by gently tapping the bottom of the tube, 250µL of the master mix was added, and the tube was gently inverted twenty times to mix. Protocol “C” on the PIPseq Dry Bath was used for cDNA synthesis (lid=105°) (**Table 3**).

**Table 3.** PIPseq Dry Bath Setting “C” parameters for cDNA Synthesis.

| Temperature | 25°C | 42°C | 85°C | 4°C |
| --- | --- | --- | --- | --- |
| Time | 30 min | 90 min | 10 min | HOLD |

### Day 2: PIPseq—cDNA Amplification

Rather than storing the samples at 4°C overnight, after cDNA synthesis, we proceeded directed to the cDNA Amplification steps. Samples were spun down for 30 seconds and 300µL of supernatant was removed. PIPs were washed three times in 0.5X Wash Buffer according to the standard protocol, including brief vortexing (5sec, 3000RPM, PIPseq vortexer). Aspiration volume to normalize each mixture volume to 250µL was again calculated (**Formula 3**). Samples were spun down for 30 seconds, and the designated volume of supernatant was removed. WTA Mastermix was generated according to the following recipe per sample: 94.4µL 4X PCR Master Mix and 1.87µL WTA Primer. 85.7µL of WTA Mastermix was added to each sample and sample mixtures were vortexed briefly (5sec, 3000RPM, PIPseq vortexer). WTA reaction mixture was distributed into eight 42µL aliquots across a PCR tube strip. Samples underwent amplification (lid = 105°C), then the samples were stored at 4°C in the thermocycler overnight (**Table 4**). 15 cycles for the step that includes both annealing and extending were chosen according to the recommended conditions for 30,000 primary cells.

**Table 4.** Thermocycler settings for cDNA Amplification.

| Temperature | Time | # of cycles |
| --- | --- | --- |
| 95°C | 3 min | 1 |
| 98°C<br>69°C | 15s<br>4 min 20 sec | **15** |
| 72°C | 5 min | 1 |
| 4°C | hold | - |

### Day 3: PIPseq—Isolate cDNA from PIPs

To split open PIPs, 75µL of CE Buffer was added to each WTA reaction and quickly tapped 5 times on the vortex (3000RPM) and spun down 30 seconds, instead of 5 seconds, to ensure tight pelleting of PIPs off tube walls. WTA products were pooled by sample and tap-vortexed again for 30 seconds. cDNA-containing supernatant was collected and prepared for SPRI bead enrichment at a magnetic bead ratio of 0.8x. Rather than vortexing samples and incubating for 5 minutes as stated in the base protocol, samples were gently flicked to mix and incubated on the hula mixer for 10 minutes at room temperature. This change was made to minimize shearing on the isolated cDNA, while ensuring samples are thoroughly mixed and beads do not settle. After incubation, the sample tubes were bound to a magnet (*Invitrogen* #12321D) for 5 minutes and washed twice with 85% ethanol for 30 seconds each. Beads containing cDNA were resuspended in room-temperature IDTE (pH 8.0), gently mixed by pipetting with a wide-bore tip, and incubated on a hula mixer for 5 minutes. Sample tubes were bound to the magnet for 2 minutes and, using a wide-bore pipette tip, cDNA was collected into a 1.5mL LoBind Eppendorf tube.

The PIPseq protocol utilizes the included TE Buffer (pH 8.0) to collect cDNA from beads, however we decided to utilize IDTE (pH 8.0) instead. IDTE has a lower concentration of EDTA, which can act as a chelating agent capable of inhibiting PCR, so ONT recommends ensuring that input cDNA have low levels of similar contaminants. For this reason, we chose IDTE over TE buffer.

### Quality Control (QC)

QC Checks were performed after the completion of the PIPseq (Day 3) and ONT (Day 4) portions of the protocol. The 1X dsDNA Qubit High Sensitivity Kit (*ThermoFisher* #Q33231) was used to determine cDNA concentration, with the following deviations: (1) 1uL of the sample was diluted in 9uL Qubit Working Solution before being sonicated for 3 minutes to break up long strands of cDNA for accurate concentration measurement; (2) 190 µL of Qubit Working Solution was added to each Qubit sample tube (*Invitrogen #Q33252*) for a final volume of 200 µL, then vortexed for 3 to 5 seconds; (3) the sample was spun in the benchtop minifuge briefly before incubating at room temperature for 10 minutes. After incubation, the standard protocol for the kit was used to determine concentration using the Qubit (*ThermoFisher Scientific* #Q33327). Based on the concentration, the cDNA was diluted to fall below 500pg/µL in alignment with the maximum detectable concentration for the standard Femto Pulse gDNA 165kb protocol (*Agilent*, #FP-1002-0275). The Femto Pulse gDNA 165kb protocol was used to determine that fragments are largely intact and an average amplicon size >500bp (**Table S6**). The Standard PIPseq protocol utilizes a TapeStation 4200 for the QC assessment, however, we have not tested the reliability of this method for our described preparation.

### Custom oligos for adaptation of PIPseq cDNA for ONT sequencing

Custom DNA oligos were designed to provide overlapping sequences that allow connecting sequences primed with the PIPseq WTA Primer to the ONT-specific cDNA primer (cPRM). These oligos bridged the PIPseq protocol to the ONT Single-Cell Protocol and allowed the nanopore sequencing of cDNA produced via PIPseq. Oligos with the below sequences were ordered from Integrated DNA Technologies (IDT) at 100nM (dry) with HPLC Purification. Once received, oligos were reconstituted in TE Buffer (pH 8.0) at a stock concentration of 100µM and stored at -20°C. Stock concentrations were diluted in TE Buffer to a working concentration of 10uM for the protocol. Oligo sequences below are color-coded according to the portion of the sequence it aligns to.

**Red** indicates the overhang sequence, **black** indicates the ONT-specific cDNA primer (cPRM), and **blue** indicates the PIPseq WTA Primer. The Forward primer has a Biotin tag on the 5’ end.

### Oligo Sequences

- [Btn]Fwd_3580_partial_read1_PIP sequence: 5’-/5Biosg/CAGCACTTGCCTGTCGCTCTATCTTCCTCTTTCCCTACACGACGCTC -3’
- Rev_PR2_partial_TSO_PIP sequence: 5’-CAGCTTTCTGTTGGTGCTGATATTGCAAGCAGTGGTATCAACGCAGAG -3’

### Days 4 and 5: ONT library preparation and sequencing

This section was adapted from two protocols provided by Oxford Nanopore Technologies (ONT): “*Single-cell transcriptomics with cDNA prepared using 10X Genomics*” and “*cDNA-PCR Sequencing V14 (SQK-PCS114)*”. Because of a legal agreement with ONT, we are unable to disclose the specific steps of these protocols. For clarity, these will be referred to as “*Single-cell ONT protocol*” and “*V14 protocol*”, respectively, throughout this manuscript.

Briefly, this protocol used PCR to attach custom oligos, which ensured compatibility of the PIPseq-generated cDNA with the ONT cPRM. Biotin-mediated size selection was conducted to ensure the isolation of reads with the custom oligos. PCR was again performed to attach the isolated transcripts to the ONT cDNA Primer (cPRM) before performing a quality check, as described in the Quality Control (QC) section of this pape**r**. The next day (Day 5), the adapter containing the motor protein was attached to each strand and the sample was fed into flow cells in the PromethION. See **Figure 1C** (middle box) for the final structure of reads before sequencing.

We made the following changes to these published protocols: Any time either ONT protocol called for the cDNA sample to be vortexed, we instead pipetted very gently 10-20 times with a wide-bore pipette tip to mix. The ONT single-cell method was designed to be used with single-cell barcoded cDNA produced with the 10X Genomics Next GEM Single Cell 3’ Kit (V3.1). Our modifications to this protocol utilize cDNA amplicons produced using the long-read PIPseq (V4.0PLUS) protocol above. As such, we recommend the modified sequences for the custom oligos described in the “Custom Oligos for Adaptation of PIPseq cDNA for ONT Sequencing” methods section, which were designed to bridge the ONT and PIPseq sections of the protocol to ensure compatibility. This protocol was designed with the updated ONT R10 chemistry in mind. Sequencing preparation for the ONT protocol was intended to be performed using the PCR-cDNA Sequencing Kit (*ONT, #SQK-PCS111*) with PromethION R9.4.1 flow cells (*ONT #FLO-PRO002*). Our protocol utilizes the updated ONT PCR-cDNA Sequencing Kit V14 (*ONT #SQK-PCS114*) with PromethION R10 flow cells (*ONT, #FLO-PRO0000*). The standard protocol uses a AMPure XP bead ratio of 0.8X for both the “Pre-Pull-Down” and “Post-Pull-Down” bead enrichment steps, however our lab found that using a ratio of 0.7X enriches for larger fragments. Bead washing for the “Pull-down” section of the protocol (steps 1-7 in single-cell ONT protocol, “Pull-down” section), was performed during the pre-pull-down PCR cycling to increase efficiency. Aside from these alterations, the biotin-tagging reaction and pre-pull-down PCR sections were performed as written in the published ONT single-cell protocol. Targeting three flow cells, we aimed to recover >210ng cDNA before sequencing. After Post-Pull-Down, QC was performed and concentration was assessed using the standard protocols for the Qubit High Sensitivity Kit and Femto Pulse gDNA 165kb kit, as described in the **Quality Control** methods section.

The next day, once fragment quality had been confirmed, the sequencing preparation was resumed. Starting at the “Adapter Addition” section of the ONT single-cell protocol, we followed the adapter addition, priming, and flow cell loading steps contained within the “cDNA-PCR Sequencing V14 (SQK-PCS114)” protocol. The average amplicon size from the Agilent Femto Pulse Bioanalyzer was used to calculate the required sample volume for 70 fmol and diluted into 31µL of Elution Buffer (EB). Just before loading the samples into the flow cells, the sample tubes were gently flicked to mix. We waited a minimum of 20 minutes after loading the library into the flow cells before initiating sequencing. Sample cDNA libraries were sequenced continuously until the end of the flow cell life. Data were collected using MinKNOW (23.11.4) and .fast5 files were basecalled using the Dorado (7.2.13+fba8e8925) graphics processing unit (GPU) basecaller. Any remaining sample was placed at -80°C for long-term storage.

### Computing power used for analyses

Analyses were performed using the University of Kentucky Morgan Computing Cluster (MCC), a batch-processing cluster with AMD EPYC Rome processer cores distributed across 182 compute nodes. Each node is configured with 512GB to 4TB of memory. The MCC has a shared GPFS parallel filesystem with 3PB of disk storage for user home, project, and scratch directories. Additionally, the cluster has a high-speed 10Gb data transfer node for efficient data movement to and from external servers. Analyses were performed in Jupyter Notebook^74,75^ using Python3 v3.10.13.

### Read preprocessing and Barcode Rescue

After sequencing, the resulting files were moved off the PromethION Data Acquisition Units (*ONT, #PRO-PRCA100*) to our storage units. The sequencing summary .txt files and the passing FASTQ files were copied onto one of the University of Kentucky’s computing clusters. All FASTQ files generated from Dorado (V7.2.13+fba8e8925) were processed in batches of 500 files separated by sample and flowcell, then concatenated into a single file by flow cell. Using a Python 3.10.13 script, reads with a mean base quality less than nine were removed (See **Data and code availability** section). The resulting filtered FASTQ was run through pychopper (2.7.10) using custom primers: The Fwd_3580_partial_read1_PIP sequence minus the overhang sequence and the cPRM from the Rev_PR2_partial_TSO_PIP sequence. The rescued fused reads from pychopper were concatenated with the regular reads output from pychopper. The number of reads was counted and saved, and the files were split into multiple files of 8,000 reads each.

Due to the higher error rate in long-read data, simply running the paired-end reads through PIPseeker was insufficient for barcode rescue. PIPseeker ^28^uses Hamming distance for barcode matching and error correction, which accounts only for base substitutions, the most common error in short-read sequencing. While insertion and deletion errors are rare in short-read sequencing, long-read sequencing frequently produces insertions and deletions, with an expected insertion rate ∼0.46%, deletion rate of ∼0.62%, and mismatch rate ∼2.1% for ONT-based sequencing^76^. This may shift the sequence of the barcode, so with the tiered structure of the barcode, a more complex pre-processing approach was needed before plugging the data into PIPseeker. Unlike Hamming distance, Levenshtein distance accounts for insertions, deletions, and substitutions, rather than just substitutions alone. Adopting this approach enhances the accuracy and reliability of barcode recovery.

Java code (openjdk 17.0.13) was used on the pychopper output to rescue barcodes and convert them into pseudo-paired-end read files for PIPseeker^28^, as described below. During the rescue process, each read was examined to find the tiered barcode used by PIPseeker. We then calculated the Levenshtein distance of each tier to each of the whitelist options for that tier. If an exact match for the tier was not found, (i.e., a Levenshtein distance of 0) we performed one of three actions: (1) If the lowest Levenshtein distance was greater than two, the read was discarded, as we could not determine the correct sequence for the tier; (2) if two or more correct sequences shared the lowest Levenshtein distance in the tier, the barcode was labeled as ambiguous and sent it to a separate file to attempt rescuing later; (3) we replaced the tier with the sequence from the whitelist with the lowest Levenshtein distance.

PIPseeker^28^ is designed for use with short-read data that sequences libraries using paired-end reads. As required, the read was converted into pseudo-paired-end read files, with a Read 1 (R1) file containing the corrected barcode sequence indicating that of an individual cell and a molecular index sequence (UMI) unique to each mRNA molecule. A Read 2 (R2) file contained the cDNA contents from the mRNA captured in each PIP. A list of all full-length barcodes in the R1 file was kept to use as a sample-specific master whitelist of barcodes seen in the sample. All barcode whitelists across batches were combined into a single master whitelist, split by sample. This whitelist was then fed into a second script (java) to rescue barcodes, which compared the barcodes in the reads with ambiguous barcodes with these sample whitelists. If using this whitelist made the barcode unambiguous, the read was kept and converted to a pseudo-paired-end read as described above, otherwise, the read was discarded.

We then ran the pseudo-paired-end FASTQs through PIPseeker (v02.01.04)^28^, which replaced barcodes with a shorter version and applied additional read filtering. Software documentation can be read for full details; however, briefly, the TSO and poly-A sequences were removed from the R2 files, and a FASTQ file was exported from reads >20bp with barcodes and UMI sequences in the read header. Reads were then concatenated by sample and flowcell into one FASTQ file (no longer paired-end) to use later for isoform discovery. Reads were also demultiplexed into separate FASTQs (no longer paired-end) by barcode and isoforms with at least 100 reads per cell were kept to use for single-cell isoform quantification.

### Pseudo-bulk alignment and isoform discovery

Our process for genomic alignment and quality control was fundamentally the same as that described in Heberle, *et al* ^9^, in the section titled “Read preprocessing, genomic alignment and quality control”. Deviations from the described process included earlier use of pychopper, as described above, and the use of updated versions of minimap2 (v2.28-r1209) and SAMtools^77^ (v1.19.2). Please see **Data and Code Availability** section for more information.

Our process for isoform discovery and quantification was also similar to that reported in the first paragraph of the section titled “Transcript discovery and quantification” in Heberle, *et al* ^9^. The changes were listed here. ERCC spike-ins were not used, since these samples were within the same batch. We used the Ensembl HG38 release 113 annotation file. An updated version of Bambu^29^ (v3.8.1) was used instead of v3.0.5. We used the New Discovery Rate (NDR) recommended by the Bambu machine learning model, which was 0.051 for this dataset. Using the outputted extended annotation file, we ran gffcompare^78^ (v0.12.6) against the Glinos et al.^7^, Leung et al.^8^, and Heberle et al.^9^ annotations.

### Single-cell alignment and isoform quantification

Barcode-separated FASTQ files were aligned using minimap2^79^, and the aligned BAMs were filtered using SAMtools^77^, as in the pseudo-bulk alignment. However, discovery to find new isoforms was not performed, instead we used the extended annotation from the pseudo-bulk isoform discovery as the annotation passed to Bambu to quantify isoforms. We also batched the FASTQs into groups of 300 files and ran those through the quantification process. After, we used the Polars^80^ Python package to merge the matrices outputted by Bambu into a single matrix per output type (gene counts, transcript counts, transcript CPM, full length counts, and unique counts).

### Data processing and bioinformatic quality control

To deal with file management, initial filtering utilized a java script that automatically removed of cells with <300 reads, and genes expressed in <10 cells. After initial filtering to remove empty droplets, supervised filtering was performed to identify appropriate thresholds for low-quality or stressed cells. Cell counts were taken initially and after filtering. QC metrics were computed, and the distribution of each measure was plotted. Ranges for quality threshold were determined based on recommended values^81^, however, final thresholds were determined according to the data distribution, when the violin plot begins to thin at the top and bottom. Different sample datasets within the same study underwent QC filtering individually before concatenation. Hemoglobin and mitochondrial genes were identified according to gene prefix, and the following QC metrics were enforced: cells per gene/isoform, total number of genes/isoforms per cell, total transcript counts in a cell, log-transformed total counts for genes/isoforms, percent counts from mitochondrial genes, and percent counts from hemoglobin genes. These metrics (**Table S2)** were employed to identify and dispose of low-quality, stressed, or dying cells. For example, we applied a mitochondrial upper threshold of 15%, adjusted based on the distribution observed in violin plots of mitochondrial gene percentages. We tailored this threshold for both datasets to ensure that only healthy cells are retained for analysis.

Datasets were converted to AnnData (v0.11.3)^82,83^ files and combined with an “inner” join to keep only genes common between the datasets. After combining datasets, as a secondary measure to remove doublets, the Scrublet package V0.2.3^84^ was used to estimate and remove possible doublets. Automatic thresholds for the doublet score for each dataset were calculated at 0.648 (gene) and 0.642 (isoform), however, a more stringent threshold of 0.3 (based on the point of distribution curve plateauing) was used to maximize doublet. Summary statistics by sample after filtering can be found in **Table S7**.

### Model building and AutoZI

The AutoZI model from the scVI-tools library (V1.2.0)^34,85^ was used to analyze the concatenated data, with batch correction applied using sample metadata. The model was initialized with 20 latent dimensions and a dropout rate of 0.3. The model was trained on 90% of the data for a maximum of 300 epochs at a learning rate of 0.01. To minimize over-training, early stopping was enabled (patience = 70) and a weight decay rate of 0.0005 was utilized to prevent the overuse of any one neuron during modeling. The remaining 10% of the dataset was used for training validation. This sampling was performed randomly. After modeling, the latent representations and denoised data were extracted, normalized per 10,000 reads, and stored in the AnnData object to correct for zero inflation. The zero-inflated probabilities were identified and printed for the user’s evaluation, to show the extent of correction needed **(Table S3).**

### Cell clustering and cell-type assignment

With the denoised data from the AutoZI model, clustering was performed using a k-nearest neighbor (kNN)^86^ graph, built with 20 neighbors per cell. Uniform manifold approximation and projection (UMAP)^87^ was applied for dimensionality reduction, and clusters were identified using the Leiden^88^ algorithm. The Leiden algorithm (flavor = igraph^89–91^) was run at multiple resolutions (0.06 for cell-type, 0.26 for sub-cell-type) to identify clusters and determine the resolution that best represents the biological dataset.

After clustering, cell-type clusters were assigned using known cell-type-specific flow cytometry markers, including surface, secreted, and transcription factor markers. Megakaryocytes, Monocytes, T cells, B cells, and Natural Killer (NK) Cells were detected based on cluster-specific expression of the markers in **Table S4** and explained further in **Supplemental Information**. For cell sub-types, these are detected in clusters that express the broad cell markers (e.g., *CD3D* for T cells) along with the subtype-specific markers. The appropriate resolution was determined by cluster distinctness on UMAP figures and expression patterns of well-established PBMC markers used in flow cytometry that reflect both lineage and function, highly expressed in single clusters of interest. Since transcript expression does not always correlate with protein expression due to differences in transcriptional bursting and protein half-life, we tested a range of highly specific markers to sort cells. The final markers that we used to determine cell type based on high co-expression in a single cluster are in **Table S4**.

Resolutions with distinct clustering and multiple marker genes that fell within the confines of our biological relevance thresholds were deemed to be most relevant to the study. Markers were also plotted to demonstrate expression levels across the cluster, to ensure that they were widely expressed within the cluster of interest. Marker genes were plotted across clusters to check for high cluster specificity and expression within the cluster of interest, as well as on the pseudobulk level to ensure high raw read count (**Figure S4**).

### Protein Structure Prediction

Protein structures were predicted for isoforms according to amino acid sequence using the ColabFold v1.5.5^40,92,93^ running AlphaFold2-ptm^39^. Predicted structures were generated and ranked using the following settings: Multiple-sequence alignments (MSA) were generated using MMseq2^94^ with the mmseqs2_uniref_env option^95,96^. The pdb100 database^97,98^ was used to source structural templates from known homologs. Modeling was performed with dropout enabled across 8 random seeds, each producing 5 protein model predictions. Protein structures were refined via 12 recycling iterations (feeding the output back into the model) without early stopping (recycle_early_stop_tolerance = 0.0). A greedy pairing strategy was enabled to align taxonomically matching sequences, and paired MSAs were generated in unpaired_paired mode to allow independent and co-evolutionary alignment. A maximum number of paired:unpaired MSA sequences was set to 256:512. All 5 models per seed underwent AMBER relaxation^99^ after generation to improve predicted geometry. Model quality was assessed using the predicted local distance difference test (pLDDT) and predicted aligned error (PAE). The top-ranked model was visualized and colorized according to IDDT. Protein sequences used for modeling can be found in **Table S5** and outputs including .pdb file for 3-D visualization can be found on Zenodo^49^, as described in **Data and code availability**.

## Acknowledgements

The authors thank Fluent Biosciences for providing guidance on optimal techniques for the base protocol and collaborating with us on barcode retrieval from PIPseeker. We would also like to thank Bibi Broome for her assistance with collecting blood samples. Lastly, we thank the University of Kentucky Center for Computational Sciences (CCS) and Information Technology Services Research Computing for their support and use of the Morgan Compute Cluster (MCC) and associated research computing resources. This work was supported by the National Institutes of Health [5T32AG078110-02 to University of Kentucky Sanders-Brown Center on Aging, AG078110 to Doyle; R01AG068331 to Ebbert; GM138626 to Ebbert; NS088555 to Stowe]; Alzheimer’s Association [2019-AARG-644082 to Ebbert]; the Bright Focus Foundation [A2020161S to Ebbert]; and PhRMA Foundation Predoctoral Drug Discovery Fellowship to Heberle.

## Author Information

These authors contributed equally: Patricia Doyle and Madeline Page.

## Contributions

P.H.D., M.T.W.E., and J.A.B. conceived and designed the study. P.H.D. designed modifications to the long-read PIPseq protocol and performed all experiments. M.L.P., B.A.H., and B.J.W. designed the data preprocessing pipeline. M.L.P. implemented the preprocessing pipeline, performed all preprocessing, and carried out initial bulk-level isoform discovery. M.T.W.E. created a pre-filtering script. P.H.D. performed cell filtering, bulk- and single-cell-level analyses, and generated all figures. A.M.S. helped with data interpretation. P.H.D., M.T.W.E., M.L.P., and A.M.S. wrote the manuscript.

## Data and code availability

Raw long-read scRNA-seq data generated and utilized for this study will be publicly accessible once we find a solution to host the full dataset long-term. In the meantime, it is available upon request.

Reference files, annotations, and outputs from long-read scRNA-seq pipelines and ColabFold protein modeling are publicly available at Zenodo^49^: (http://doi.org/10.5281/zenodo.17341210).

All code used for these analyses is publicly accessible at: https://github.com/UK-SBCoA-EbbertLab/single_cell_analysis_PBMCs/tree/main.

HG38 reference genome sequence is available at: https://ftp.ensembl.org/pub/release-113/fasta/homo_sapiens/cdna/.

HG38 reference GFF3 annotation is available at: https://ftp.ensembl.org/pub/release-113/gff3/homo_sapiens/.

Transcript annotations utilized to determine the novelty of new transcripts and genes can be found at the following links:

Glinos *et al*.^7^: https://storage.googleapis.com/gtex_analysis_v9/long_read_data/flair_filter_transcripts.gtf.gz.

Leung *et al*.^8^: https://zenodo.org/record/7611814/preview/Cupcake_collapse.zip#tree_item12/HumanCTX.collapsed.gff.

Heberle *et al*.^9^: https://doi.org/10.5281/zenodo.8180677.

## Rigor and Reproducibility

This study was performed under the ethics oversight of the University of Kentucky Institutional Review Board. A singularity container was used for most analyses in this study, with the exception of web-based analyses SAPFIR, InterScanPro, DeepTMHMM, and ColabFold. Use of singularity containers enable the encasement of specific versions of software in a way that is reproducible. Instructions on accessing the singularity container used for this project can be found in the GitHub repository for this project. All raw data, output from preprocessing pipeline, output from AutoZI modeling, references and annotations are publicly available.

## Declaration of generative AI and AI-assisted technologies in the writing process

During the preparation of this work, the author(s) used ChatGPT to generate and troubleshoot errors in code and increase the efficiency of the analysis pipeline development process. After utilizing this tool, the author(s) thoroughly reviewed, refined, and validated all code and analyses, taking full responsibility for the final content of the published article.

