## Supplemental Information, Figures, and Tables for "Decoding the human PBMC isonome: Isoform-level resolution with single-cell long-read transcriptomics"

**Additional Cell Marker information:**
We identified T cells by the co-expression the $\gamma$, $\delta$, and $\varepsilon$ subunits of the CD3 complex (*CD3G*, *CD3D*, *CD3E*), a major co-receptor involved in activation of both CD4+ T cells and CD8+ T cells[1]. NK cells were defined by the co-expression of surface-level marker *KLRF1* (NKp80)[2] and marker for cytotoxicity, *GZMB*[1,2], a major function of NK cells. *ITGAM* (CD11b)[3], and *IL2RB* (CD122)[2], are not NK-specific, but reflect their status as innate lymphocytes that participate in cytokine signaling. Monocyte-derived cells were distinguished using a key myeloid lineage marker, *CD33*, an inhibitory marker enriched in monocyte-derived dendritic cells (*LILRB4*), and functional receptors for immune responses (*FCGR2A*/CD32A, and *CLEC7A*/Dectin-1); this mixture of disease-relevant monocyte and dendritic cell markers in the absence of markers of macrophages led us to categorize this cluster as monocyte-derived[4–7]. B cells were identified based on expression of cell-type-specific surface markers *CD22, CD79A*, *MS4A1* (CD20), and *CD19*. *CD79A* and *CD22* are key genes involved in B cell receptor signaling[1,8], while *MS4A1* (CD20) and *CD19* are involved with B cell activation[1,9]. Finally, small numbers of Megakaryocytes, the major platelet producers, were identified by high expression of *ITGA2B* (CD41) and *GP1BA (CD42b)*, two surface receptors involved in platelet aggregation, while *MPL* regulates megakaryocyte growth and development of platelets[1,10,11].

Memory T cells were recognized via expression of *TCF7*, a transcription factor associated with self-renewal and maintenance of memory function, as well as *CCR7* and *SELL* (L-selectin), two genes associated with connection to the lymphatic system that enable long-term immune surveillance. Effector CD8+ T cells were characterized by co-expression of *CD8A* and *CD8B*, which together encode for the CD8 co-receptor, a crucial protein involved in antigen recognition and T cell signaling. These cells were also classified by co-expression of *KLRB1*, a surface marker enriched in populations exhibiting cytotoxic activity as well as *GATA3* and *CCL5*, markers tied to effector T cell function. Effector CD4+ T cells expressed high levels of *CD4* as well as *IL2RA* (CD25) and TNF, which act as markers for T cell activation. These cells co-expressed transcription factors *AHR* and *GATA3*, tied to activation and cytokine signaling responses.

**Package Version Used:**
1. Python: 3.10.13
2. os: Linux 4.18.0-305.10.2.el8_4.x86_64
3. scanpy: 1.11.0
4. anndata: 0.11.3
5. polars: 1.24.0
6. pandas: 2.2.3
7. scipy: 1.15.2
8. matplotlib: 3.10.1
9. numpy: 2.1.3
10. skimage: 0.25.2

11. scvi-tools: 1.3.0
12. scikit-learn: 1.5.2
13. muon: 0.1.7
14. scvi-tools: 1.3.0
15. seaborn: 0.13.2
16. torch: 2.6.0
17. scrublet: 0.2.3
18. scikit-image: 0.25.2
19. pybiomart: 0.2.0
20. bioservices:1.12.1
21. Rdata: 0.11.2
22. adjustText: 1.3.0
23. leidenalg: 0.10.2
24. igraph: 0.11.6
25. statsmodels: 0.14.2
26. joblib: 1.4.2
27. gseapy: 1.1.3

#### Materials

1. **General Laboratory Equipment Required**
   Cell Culture Hood with UV, Centrifuge with swinging rotor and/or adapters suitable for 15mL and 1mL tubes (*Thermo Scientific* #755007210, #75005743 (1mL adapter), #75005743 (15mL adapter)), 2000xg Benchtop mini centrifuge compatible with 1.5mL and 0.5mL tubes (*Benchmark Scientific #C1012*), PCR Thermocycler (ProFlex PCR System, 3x32-well: Applied Biosystems #4484073), Hemacytometer and 10X Microscope for cell counting, Invitrogen Qubit Flex Fluorometer (*ThermoFisher Scientific* #Q33327), Digital Vortex Mixer (*Thermo Scientific* #88882009), Brandon Ultrasonics CPXH Series Ultrasonic Cleaning Bath (*Fisher Scientific* #15-336-125), Femto Pulse System (*Agilent*, #M5330AA), DynaMag-2 Magnet (*Invitrogen* #12321D), Laboratory balance (mg) (*FisherScientific* #01-919-149), HulaMixer Sample Mixer (*Invitrogen* #BB3362792), Motorized Serological Pipette Controller (*BrandTech* #26333), micropipettes (*Eppendorf* #3123000063 (P1000), #3123000055 (P200), #3123000039 (P20), #3123000020 (P10)), PromethION 24 (*ONT,* #PRO-SEQ024), PromethION Data Acquisition Unit (*ONT, #PRO-PRCA100*)

2. **Materials and Reagents Needed for PBMC Isolation (optional)**
   Blood Collection tube, BD Vacutainer® K2EDTA 10mL (*VWR* #BDAM367525), SepMate-50 (IVD) (*Stemcell Technologies*, #85450), Ficoll PAQUE Plus (*Millipore Sigma* #17-1440-020), Sterile 50-mL conical tubes (*Eppendorf* #0030122178), Serological Pipettes (*VWR* #75816-090 (25mL), *VWR* #75816-100) (10mL)), eBioscience 10X RBC Lysis Buffer (Multi-species) (*Invitrogen* #00-4300-54), Fetal Bovine Serum (FBS) (Corning #35-015-CV), Ultrapure 20X Phosphate Buffered Saline (PBS) pH 7.5 (VWR #E703-1L), Ultrapure H2O

3. **PIPseq Equipment Required**
   The following equipment is included in the PIPseq T20 3’ Single Cell Starter Equipment Kit (*Fluent* #FBS-SCR-STKIT): PIPseq Vortex Mixer (*Fluent #FBS-SCR-DVM*), PIPseq Dry Bath with heated lid (*Fluent* #FBS-SCR-PDB), PIPseq Dry Bath 1.5mL block (*Fluent* #FB0002498), PIPseq 4-tube stand, blue, for 1.5mL tubes (*Fluent* #FB0004722), PIPseq Rotating vortex base assembly (*Fluent* #FB0002100).

4. **Plastic Consumables Required for Long-Read PIPseq**
   a. **PIPseq Plastics:** The following consumables are included within the PIPseq T20 3’ Single Cell Consumables Kit v4.0PLUS (*Fluent* #FBS-SCR-T20-4-V4.05-6): 1.5mL Safe-Lock PCR Clean tubes (*Eppendorf* #022363212), 0.2mL PCR 8-tube strip without Cap (*Greiner Bio-One*, #373270), PCR 8-Cap strips, domed cap (*Greiner Bio-One*, #373270), 3mL syringe, 1mL, G22 blunt bottom syringe needle.
   b. **Other Plastics:** Standard low-retention pipette tips (*Genesee Scientific* #24-430 (1000uL), #24-412 (200uL), #26-404 (20uL), #24-401 (10uL)), Wide-bore low-retention pipette tips (*Genesee Scientific* #22-428 (1000uL), #22-425 (200uL)), 10mL serological pipettes (*VWR* #75816-100), Sterile DNase-, RNase-, DNA-free 15mL tubes (*Eppendorf,* #0030122160), 0.2mL thin-walled PCR tubes (*Genesee Scientific* #24-705), 1.5mL DNA LoBind tubes (*Eppendorf*, #022431021), Qubit Flex Assay Tube Strips (*Invitrogen #Q33252*)

5. **Kits Needed for Long-Read PIPseq**
   a. **PIPseq Kits:** PIPseq T20 3’ Single Cell RNA Ambient Kit v4.0PLUS (*Fluent* #FBS-SCR-T20-4-V4.05-1), PIPseq T20 3’ Single Cell RNA 4ºC Kit v3.0 or 4.0 (*Fluent* #FBS-SCR-T20-4-V3&V4-2), PIPseq T20 3’ Single Cell RNA -20ºC Kit v4.0PLUS (*Fluent* #FBS-SCR-T20-4-V4.05-3), PIPseq T20 3’ Single Cell RNA -80ºC Kit v4.0PLUS (*Fluent* #FBS-SCR-T20-4-V4.05-4)
   b. **Quality Control Kits:** Qubit 1X dsDNA High Sensitivity Assay Kit (*ThermoFisher* #Q33231), Genomic DNA 165kb Kit (*Agilent*, #FP-1002-0275)
   c. **ONT Kits:** The following reagents were used from ONT PCR-cDNA Sequencing Kit V14 (ONT #SQK-PCS114): Rapid Adapter (RA), Adapter Buffer (ADB), Flow Cell Tether (FCT), Flow Cell Flush (FCF), Sequencing Buffer (SB), Library Beads (LIB), Elution Buffer (EB).

6. **Additional Reagents and Materials Needed for Long-Read PIPseq**
   a. **PIPseq:** SUPERase•In RNase Inhibitor (20U/uL) (*Invitrogen* # AM2696), 100% Ethanol molecular biology grade (*Millipore sigma* #EX0276-4), Tris-EDTA (TE) Buffer 1X solution pH 8.0 (*Fisher Bioreagents* #BP2473-1), IDTE (pH 8.0), Ultrapure water
   b. **Quality Control (QC):** Trypan Blue Solution (*PromoKine* #PK-CA902-1209), RNaseZap RNase Decontamination Solution (*Invitrogen,* #AM9782)
   c. **Oxford Nanopore Technologies (ONT) Sequencing Preparation:** 100% Ethanol molecular biology grade (*Millipore sigma* #EX0276-4), Tris Base (*Fisher Bioreagents* #BP152-1), HCl 5.0N (*VWR* #BDH7419-1), NaCl (*VWR* #0241-500G), 0.5M EDTA pH 8.0 *(Invitrogen* #AM9260G), Tris-EDTA (TE) Buffer 1X

solution pH 8.0 (*Fisher Bioreagents* #BP2473-1), Ultrapure water, AMPure XP bead-based reagent (*Beckman* #A63880), M280 streptavidin 10ug/uL (*Invitrogen* #11205D), LongAmp Hot start Taq 2X Master Mix (*New England BioLabs* #M0533S), R10.4.1 flow cells (*ONT* #FLO-PRO114M)

#### Supplemental Tables:

**Table S1. Unfiltered pseudo-bulk read metrics throughout preprocessing from PBMC samples collected across 3 flow cells per sample.**

| | PBMC1 | PBMC2 |
|---|---|---|
| N reads (total) | 275,827,438 | 215,987,216 |
| N passed reads (ONT) | 228,920,664 | 163,153,766 |
| N Reads After Pychopper | 221,804,949 | 157,166,409 |
| N Reads After NanoporeConvert | 158,309,133 | 103,099,471 |
| N Reads After PIPseeker | 153,038,291 | 99,777,656 |
| N trimmed and processed passed reads | 153,038,291 | 99,777,656 |
| N reads aligned | 135,302,551 | 89,261,231 |
| N reads aligned (MAPQ≥10) | 121,867,484 | 80,033,261 |
| Reads mapping to positive strand (millions) | 70.5 M | 46.2 M |
| Reads mapping to negative strand (millions) | 51.4 M | 33.8 M |
| Non-splice reads (millions) | 58.9 M | 37.5 M |
| Splice reads (millions) | 63 M | 42.4 M |
| N50 FASTQ (mean across flow cells, nt) | 658.67 | 664.67 |
| Median read length (nt) | 476.67 | 501.33 |

**Table S2. Filtering criteria for cell quality.**

| | Lower Threshold (≥) | Upper Threshold (≤) |
|---|---|---|
| Mitochondrial Gene % | N/A | 15 |
| Hemoglobin Gene % | N/A | 5 |
| Total Counts | 600 | 11000 |
| Unique Gene Count | 250 | 1800 |
| Unique Isoform Count | 350 | 3000 |

**Table S3. Proportion of genes and isoforms predicted to be zero-inflated by AutoZI Modeling.**

| | Fraction |
|---|---|
| Predicted ZI Genes | 0.996 |
| Genes with average expression > 1.0 | 0.009 |
| Predicted ZI Genes with average expression > 1.0 | 0.575 |
| Predicted ZI Isoforms | 0.487 |
| Isoforms with Average Expression > 1.0 | 0.002 |
| Predicted ZI Genes with average expression > 1.0 | 0.565 |

*Fractions represent proportions of genes or isoforms meeting each criterion. Mean expression refers to average raw counts per feature across all cells. Values >1.0 indicate an average of at least 1 raw count per cell.*

**Table S4. Marker genes for cell-type and sub-cell-type cluster annotation.**

| | | Markers |
|---|---|---|
| T cells | | CD3D+, CD3E+, CD3G+ |
| | CD8+ Effector | CD8A+, CD8B+, GATA3, KLRB1+, CCL5+ |
| | CD4+ Effector | CD4+, IL2RA+, GATA3+, AHR+, TNF+ |
| | Memory | CCR7+, SELL+, TCF7+ |
| | Effector-Memory Transition | ITGAE+, LEF1+ IL2RA+ CTLA4+, GATA3+, IL7R+, CD27+, TCF7+ |
| Natural Killer (NK) | | GZMB+, KLRF1+, NCAM1+, ITGAM+, IL2RB+ |
| B cells | | CD22+, CD79A+, MS4A1+, CD19+ |
| Monocyte-derived | | FCGR2A+, CLEC7A+, CD33+, LILRB4+ |
| Megakaryocyte | | GP1BA+, MPL+, ITGA2B+ |

| Table S5. Protein sequences used for AlphaFold modeling, retrieved from Uniprot (v2025_03)[12]. | | |
|---|---|---|
| | | **Sequence:** |
| ***GZMB-201*** *(ENST00000 216341)* | Canonical protein-coding | **Known Protein Sequence (UniProt: P10144)** |
| | | MQPILLLLAFLLLPRADAGEIIGGHEAKPHSRPYMAYLMIWDQKSLKRCGGFLIRDDFVLTAAHCWGSSINVTLGAHNIKEQEPTQQFIPVKRPIPHPAYNPKNFSNDIMLLQLERKAKRTRAVQPLRLPSNKAQVKPGQTCSVAGWGQTAPLGKHSHTLQEVKMTVQEDRKCESDLRHYYDSTIELCVGDPEIKKTSFKGDSGGPLVCNKVAQGIVSYGRNNGMPPRACTKVSSFVHWIKKTMKRY |
| ***GZMB-204*** *(ENST00000 216341)* | Protein-coding | **Known Protein Sequence (UniProt: E9PRD7)** |
| | | MQPILLLLAFLLLPRADAGEIIGGHEAKPHSRPYMAYLMIWDQKSLKRCGGFLIRDDFVLTAAHCWGSWRERPSGPELCSPSGYLATRPR |
| **CD3G-206** (ENST00000 532917) | Canonical protein-coding | **Known Protein Sequence (UniProt: P09693)** |
| | | MEQGKGLAVLILAIILLQGTLAQSIKGNHLVKVYDYQEDGSVLLTCDAEAKNITWFKDGKMIGFLTEDKKKWNLGSNAKDPRGMYQCKGSQNKSKPLQVYYRMCQNCIELNAATISGFLFAEIVSIFVLAVGVYFIAGQDGVRQSRASDKQTLLPNDQLYQPLKDREDDQYSHLQGNQLRRN |
| **CD3G-202** (ENST00000 392883) | Protein-coding | **Known Protein Sequence (UniProt: A8MUH3)** |
| | | MEQGKGLAVLILAIILLQGTLAQSIKGNHLVKVYDYQEDGSVLLTCDAEAKNITWFKDGKMIGFLTEDKKKWNLGSNAKDPRGMYQCKGSQNKSKPLQVYYRTSDKQTLLPNDQLYQPLKDREDDQYSHLQGNQLRRN |

| Table S6. Study Protocol Quality Control (QC) metrics. | | |
|---|---|---|
| | PBMC1 | PBMC2 |
| PIPseq cDNA Yield (ng/μL) | 10.08 | 17.95 |
| Peak bp average after PIPseq Protocol | 1064.5 | 956 |
| ONT cDNA Yield (ng/μL) (input = 10ng) | 20.90 | 22.20 |
| Peak bp average after ONT Protocol | 1189.5 | 1142.5 |
| A260/A280 ratio after ONT Protocol (ng/μL) | 1.72 | 2.18 |

| Table S7. Summary Statistics After Filtering. | | | |
|---|---|---|---|
| | | PBMC1 | PBMC2 |
| Total Gene counts per cell | Min | 800.00 | 800.00 |
| | Median | 4558.5 | 1760.00 |
| | Mean | 4640.38 | 1906.00 |
| | Max | 9999.0 | 9707.00 |
| Transcripts per Gene | Min | 1.63 | 1.45 |
| | Median | 4.48 | 2.96 |
| | Mean | 4.51 | 3.00 |
| | Max | 8.59 | 6.45 |
| Unique Genes per Cell | Min | 350.00 | 350.00 |
| | Median | 1003.00 | 596.00 |
| | Mean | 997.33 | 630.06 |
| | Max | 1700.00 | 1694.0 |
| Mitochondrial Percentage | Min | 0.00 | 0.00 |
| | Median | 7.34 | 7.15 |
| | Mean | 7.63 | 7.50 |
| | Max | 15.00 | 14.99 |
| Total Isoform Counts per cell | Min | 800.11 | 800.01 |
| | Median | 4405.01 | 1689.94 |
| | Mean | 4504.40 | 1831.13 |
| | Max | 9991.86 | 9227.28 |
| Reads per Isoform per Cell | Min | 0.84 | 0.70 |
| | Median | 2.72 | 1.82 |
| | Mean | 2.76 | 1.84 |
| | Max | 5.49 | 3.77 |
| Unique Isoforms per cell | Min | 501 | 500 |
| | Median | 1596.0 | 936.0 |
| | Mean | 1603.89 | 1004.75 |
| | Max | 2995 | 2992 |

### Supplemental Figures

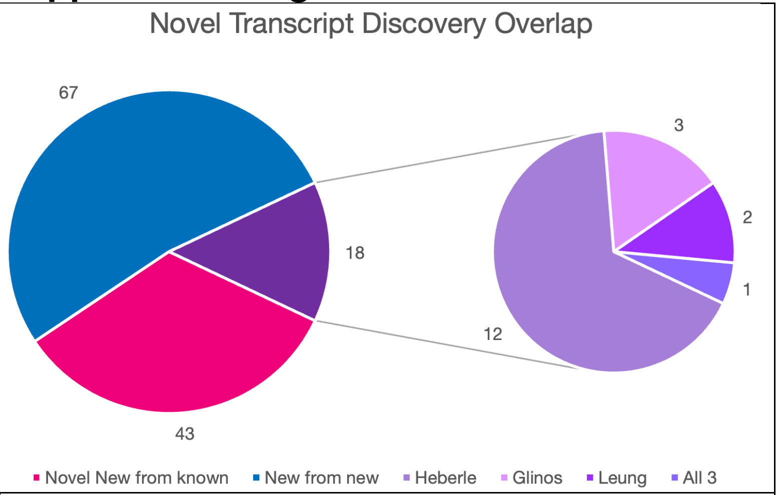


**Figure S1. Incidence of newly identified isoforms from known and new genes in existing datasets.** Pie charts showing overlap of new transcripts from this study. Of the 18 isoforms from our "new from known" list of isoforms, the highest majority (12) were from Heberle *et al.*[13]*,* followed by Glinos *et al.*[14], and Leung *et al.*[15] A single isoform was identified in all three datasets.

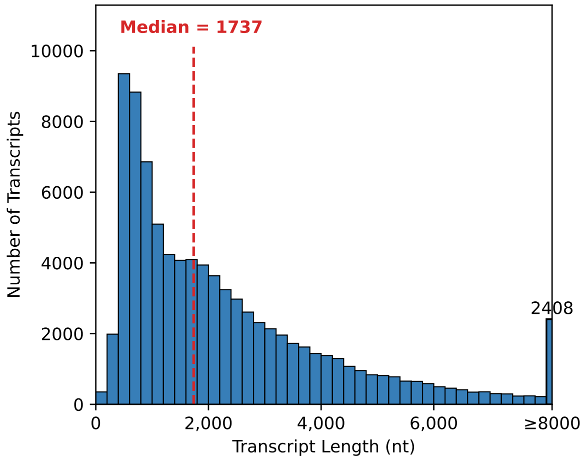


**Figure S2. Distribution of isoform length in PBMC datasets.** Histogram showing distribution of transcript lengths (in nucleotides) for all isoforms detected. The median is 1737nt, as highlighted by the red line.

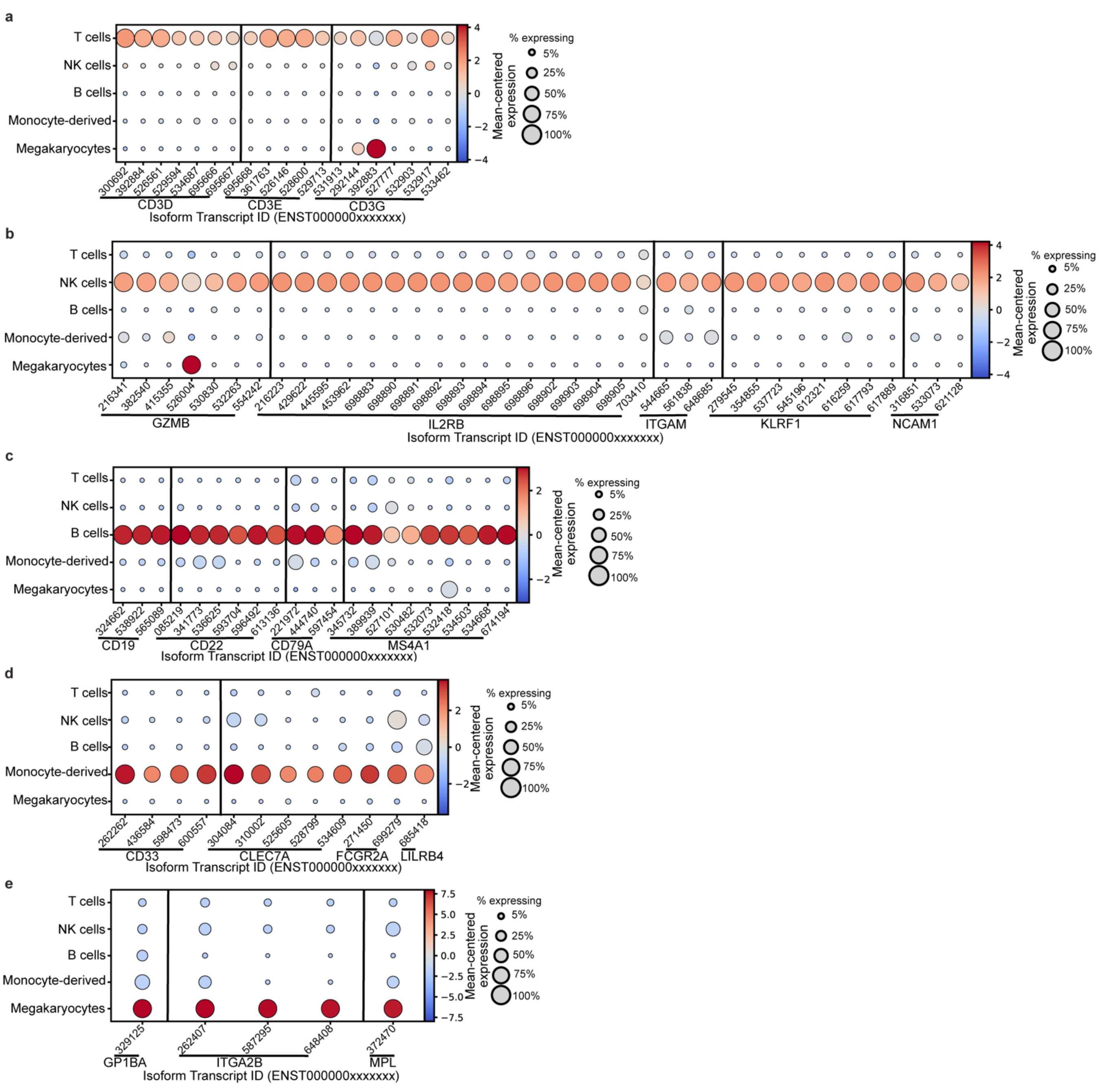


**Figure S3. Cell-type marker gene isoform expression across major PBMC cell types**. (a-e) Dotplots showing normalized expression (Mean-centered, color scale) and percentage of cells per cluster expressing each transcript (dot size) across five major PBMC cell-types: (a) T cells (*CD3D, CD3E, CD3G*), (b) Natural Killer (NK) cells (*GZMB, IL2RB, ITGAM, KLRF1, NCAM1),* (c) B cells (*CD19, CD22, CD79A, MS4A1*), (d) Monocyte-derived cells (*CD33, CLEC7A, FCGR2A, LILRB4*), and (e) Megakaryocytes (*GP1BA, ITGA2B, MPL*). Isoform transcript identifiers correspond to Ensembl ID, starting with “ENST00000” and showing the last 6 digits on the y-axis of the corresponding point.

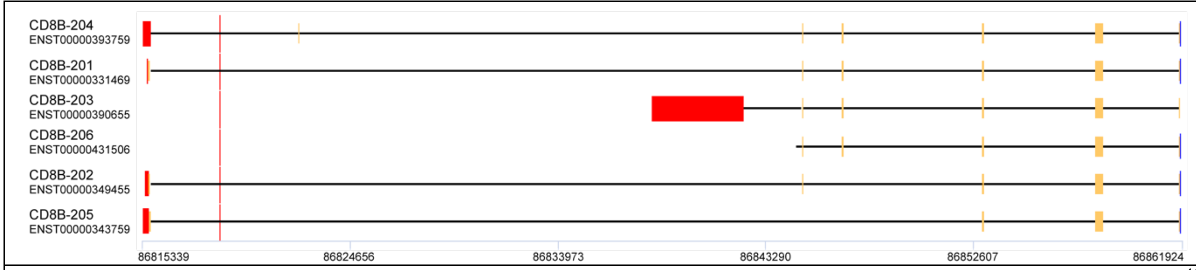


**Figure S4. Alternative untranslated region (UTR) usage among *CD8B* isoforms as visualized by UTRdb[16].** Screenshot from UTRdb showing predicted 3' (red) and 5'UTR (blue) segments, with coding exons shown as yellow boxes. Each horizontal track represents a distinct isoform of CD8B, with black lines representing introns.

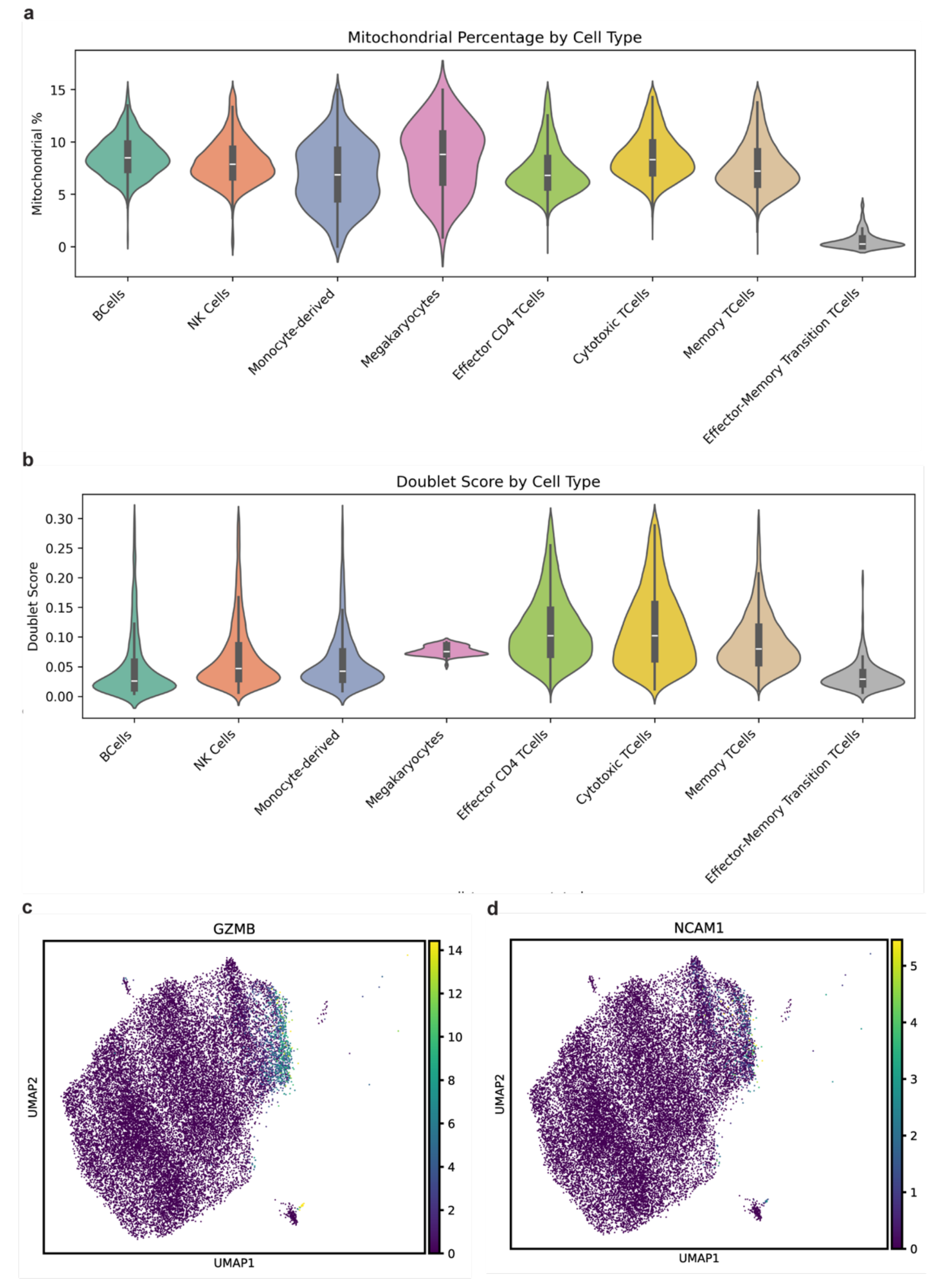


**Figure S5. Quality control metrics confirm effector-memory transition T cells form a distinct cluster from NK-T cells. (a)** Distribution by cell-type of mitochondrial read percentage per cell. **(b)** Distribution by cell-type of doublet score as calculated by Scrublet[17] python package. **(c)** UMAP colored by GZMB expression in T cell datasets. **(d)** UMAP colored by NCAM1 expression in T cell datasets.

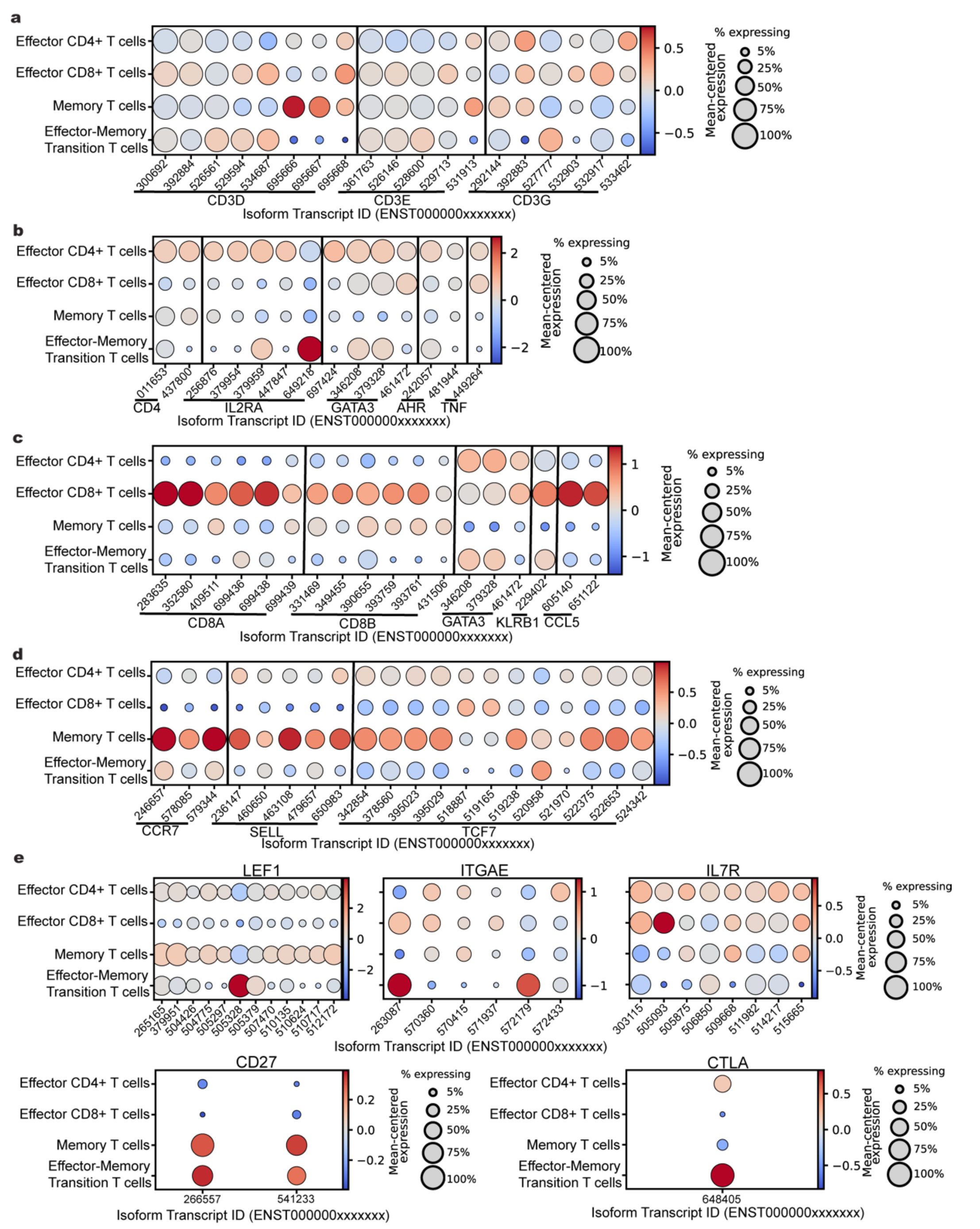


**Figure S6. Isoform-specific expression patterns among T cell subtypes. (a-e)** Dot plots showing all isoforms from marker genes expressed in our dataset and their respective normalized expression relative to mean expression (color scale) and percentage of cells expressing that isoform (dot size), across all T cell subtypes. Isoform transcript identifiers correspond to Ensembl ID, starting with "ENST00000" and showing the last 6 digits on the y-axis of the corresponding point. Isoforms express enrichment of marker genes for **(a)** general T cell markers (*CD3D, CD3E, CD3G*), **(b)** Effector CD4$^{+}$ T cells (*CD4, IL2RA, GATA3, AHR, TNF*), **(c)** Effector CD8$^{+}$ T cells (*CD8A, CD8B, GATA3, KLRB1, CCL5*), **(d)** Memory T cells (*CCR7, SELL, TCF7*), **(e)** Effector-Memory Transition T cells (*LEF1, ITGAE, IL7R, CD27, CTLA*)
