## Supplemental File 2 for "Decoding the human PBMC isonome: Isoform-level resolution with single-cell long-read transcriptomics"

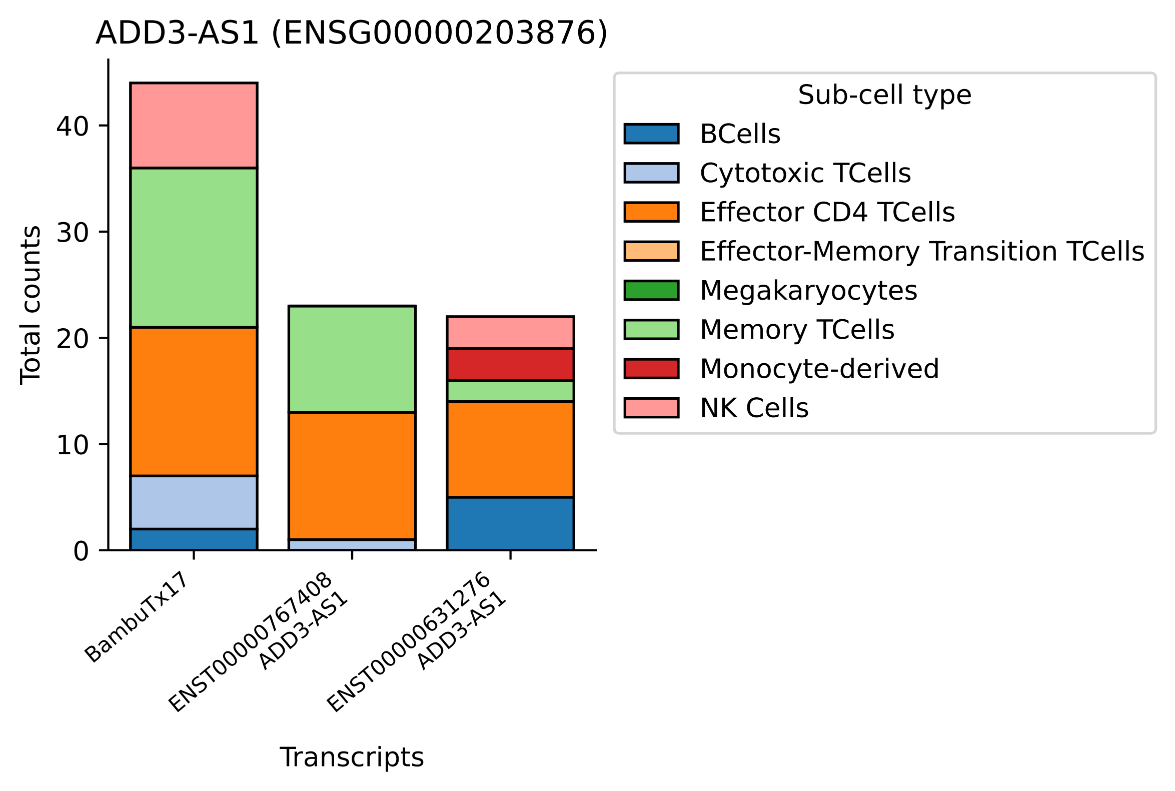

ADD3-AS1 (ENSG00000203876)
Total counts
0
10
20
30
40
BambuTx17
ENST00000767408
ADD3-AS1
ENST00000631276
ADD3-AS1
Transcripts
Sub-cell type
BCells
Cytotoxic TCells
Effector CD4 TCells
Effector-Memory Transition TCells
Megakaryocytes
Memory TCells
Monocyte-derived
NK Cells

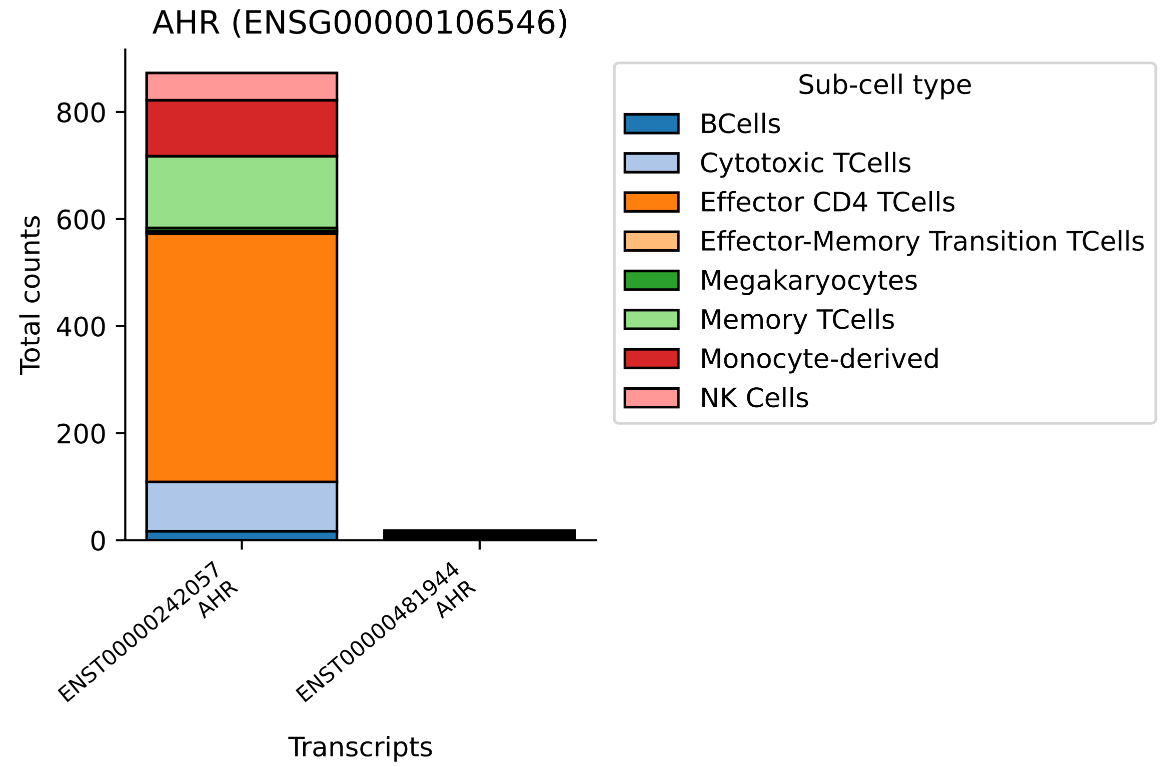

AHR (ENSG00000106546)
Total counts
0
200
400
600
800
ENST00000242057
AHR
ENST00000481944
AHR
Transcripts
Sub-cell type
BCells
Cytotoxic TCells
Effector CD4 TCells
Effector-Memory Transition TCells
Megakaryocytes
Memory TCells
Monocyte-derived
NK Cells

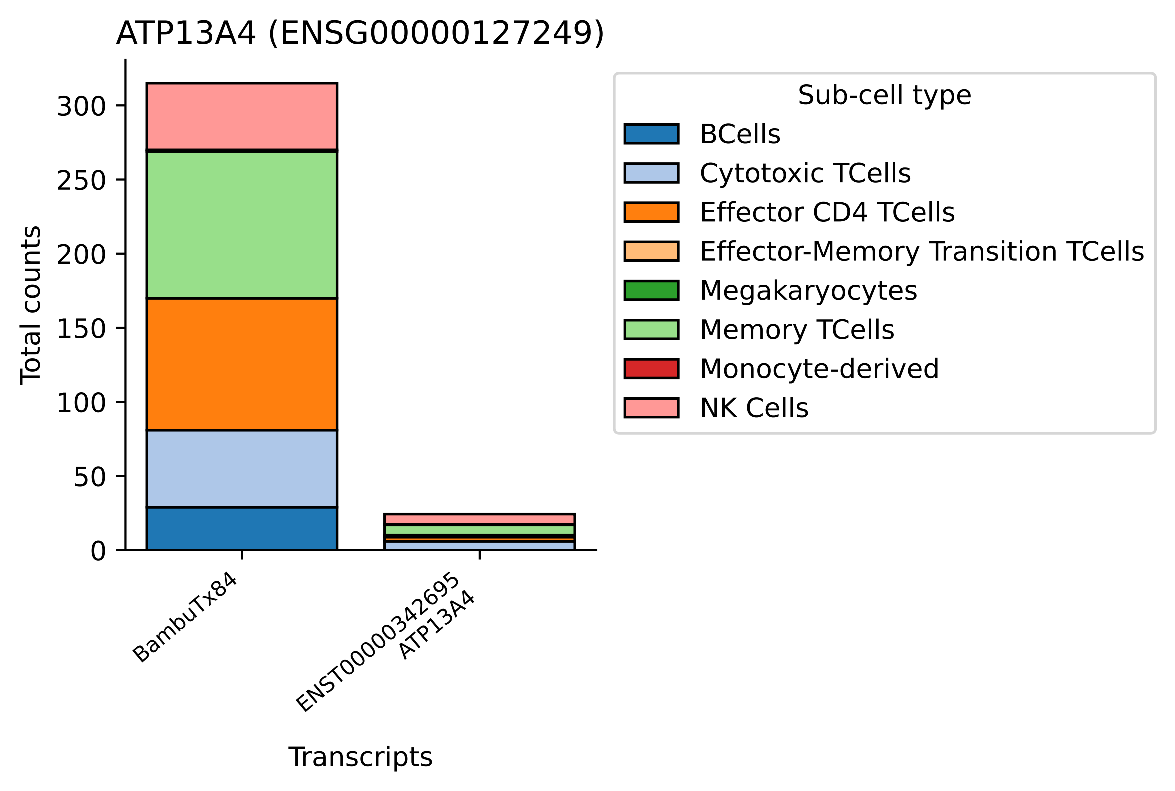

ATP13A4 (ENSG00000127249)
Total counts
300
250
200
150
100
50
0
BambuTx84
ENST00000342695
ATP13A4
Transcripts
Sub-cell type
BCells
Cytotoxic TCells
Effector CD4 TCells
Effector-Memory Transition TCells
Megakaryocytes
Memory TCells
Monocyte-derived
NK Cells

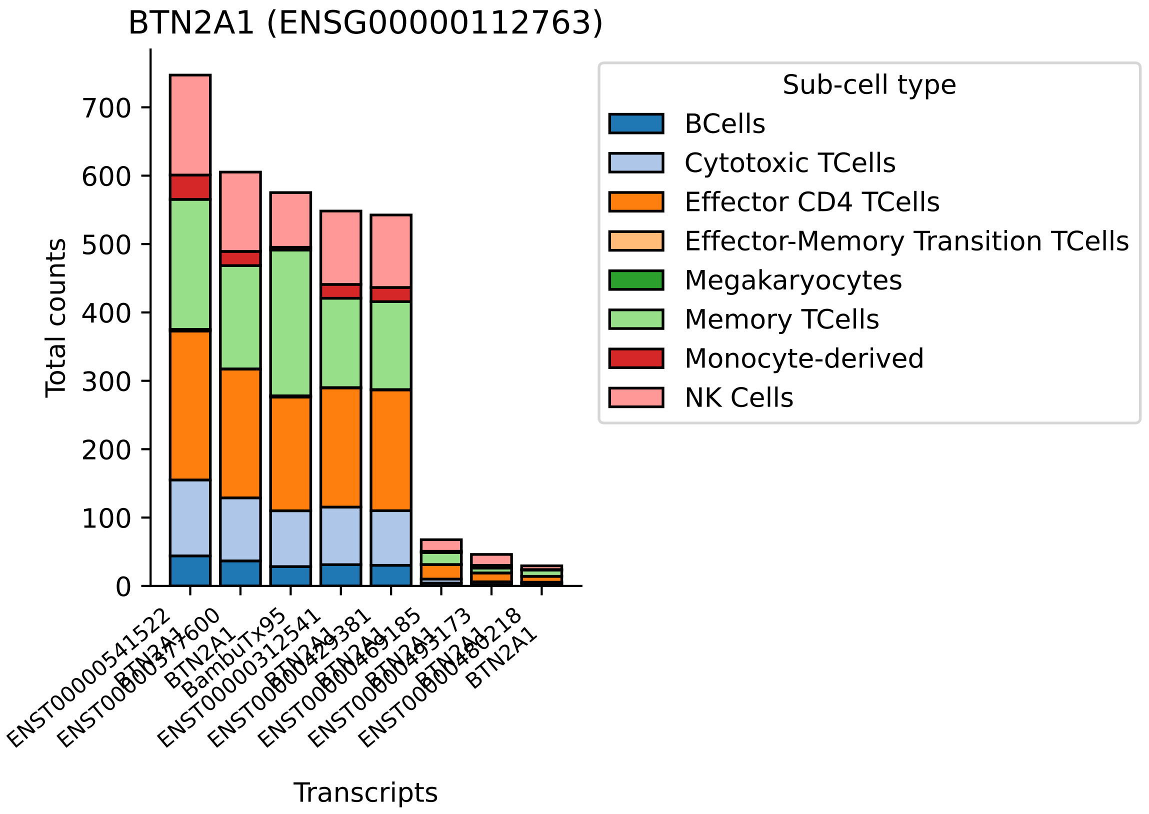

BTN2A1 (ENSG00000112763)
Total counts
0
100
200
300
400
500
600
700
Transcripts
ENST00000541522
BTN2A1
BambuTx95
ENST00000312541
Sub-cell type
BCells
Cytotoxic TCells
Effector CD4 TCells
Effector-Memory Transition TCells
Megakaryocytes
Memory TCells
Monocyte-derived
NK Cells

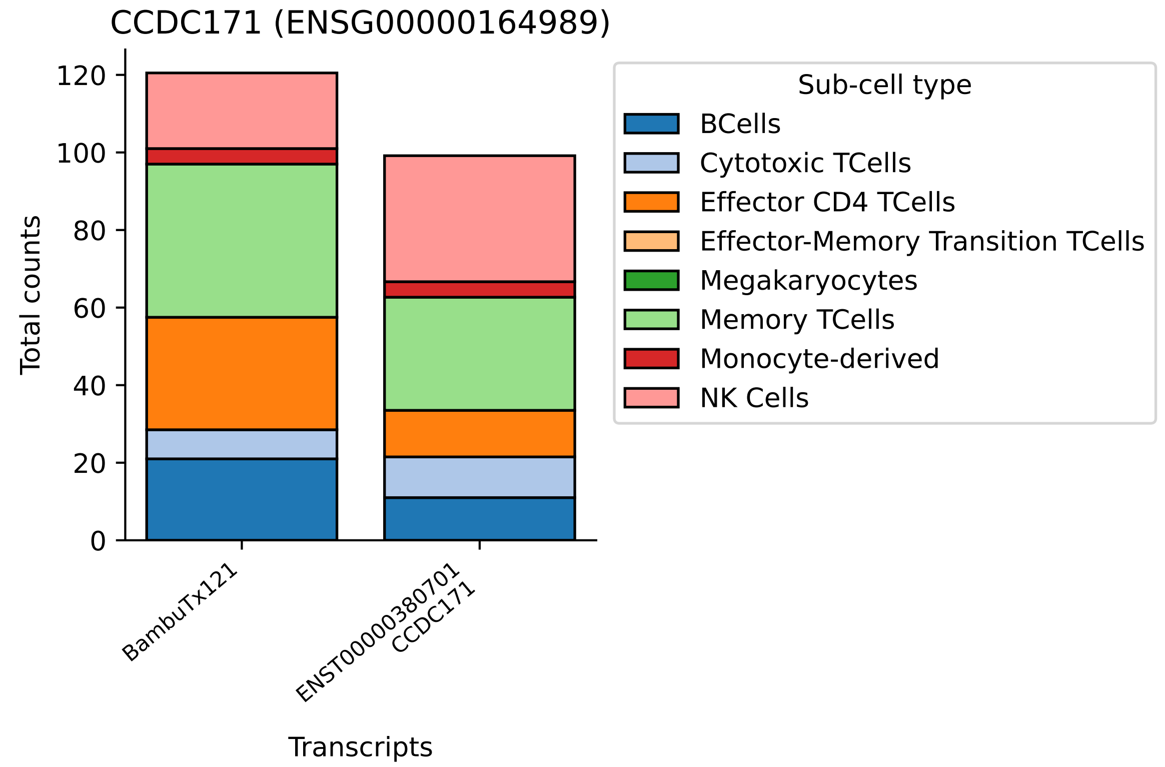

CCDC171 (ENSG00000164989)
Total counts
0
20
40
60
80
100
120
BambuTx121
ENST00000380701
CCDC171
Transcripts
Sub-cell type
BCells
Cytotoxic TCells
Effector CD4 TCells
Effector-Memory Transition TCells
Megakaryocytes
Memory TCells
Monocyte-derived
NK Cells

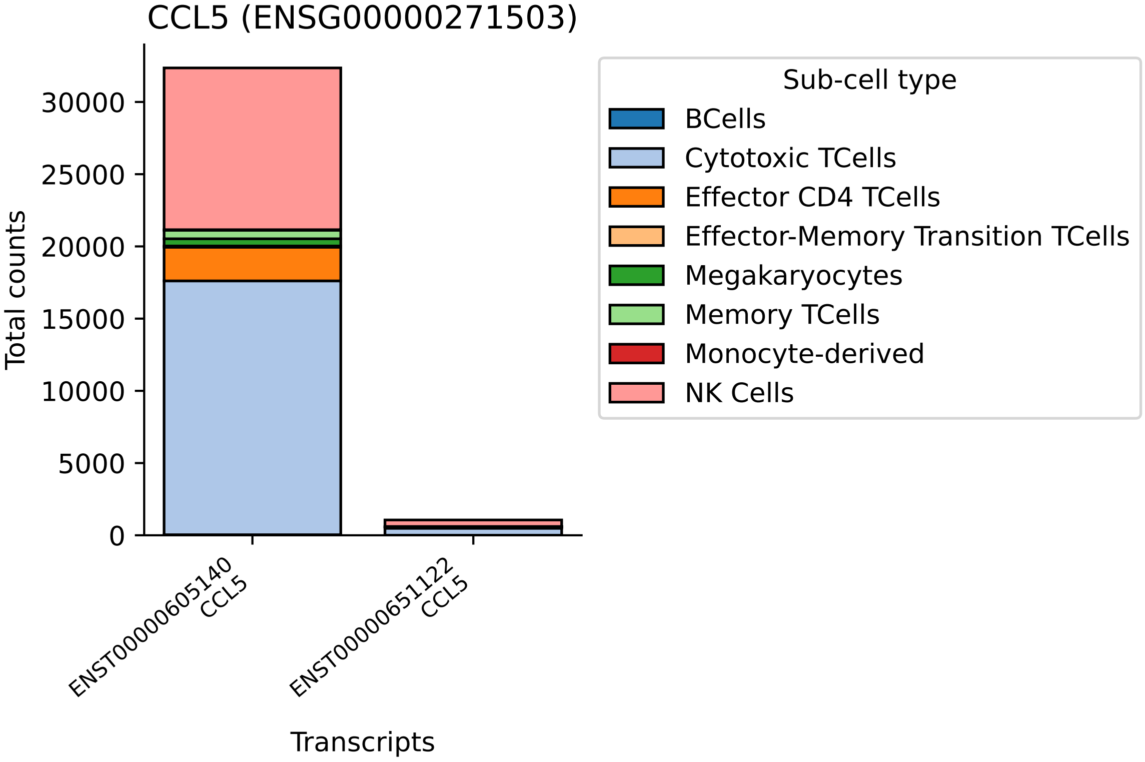

CCL5 (ENSG00000271503)
Total counts
0
5000
10000
15000
20000
25000
30000
ENST00000605140
CCL5
ENST00000651122
CCL5
Transcripts
Sub-cell type
BCells
Cytotoxic TCells
Effector CD4 TCells
Effector-Memory Transition TCells
Megakaryocytes
Memory TCells
Monocyte-derived
NK Cells

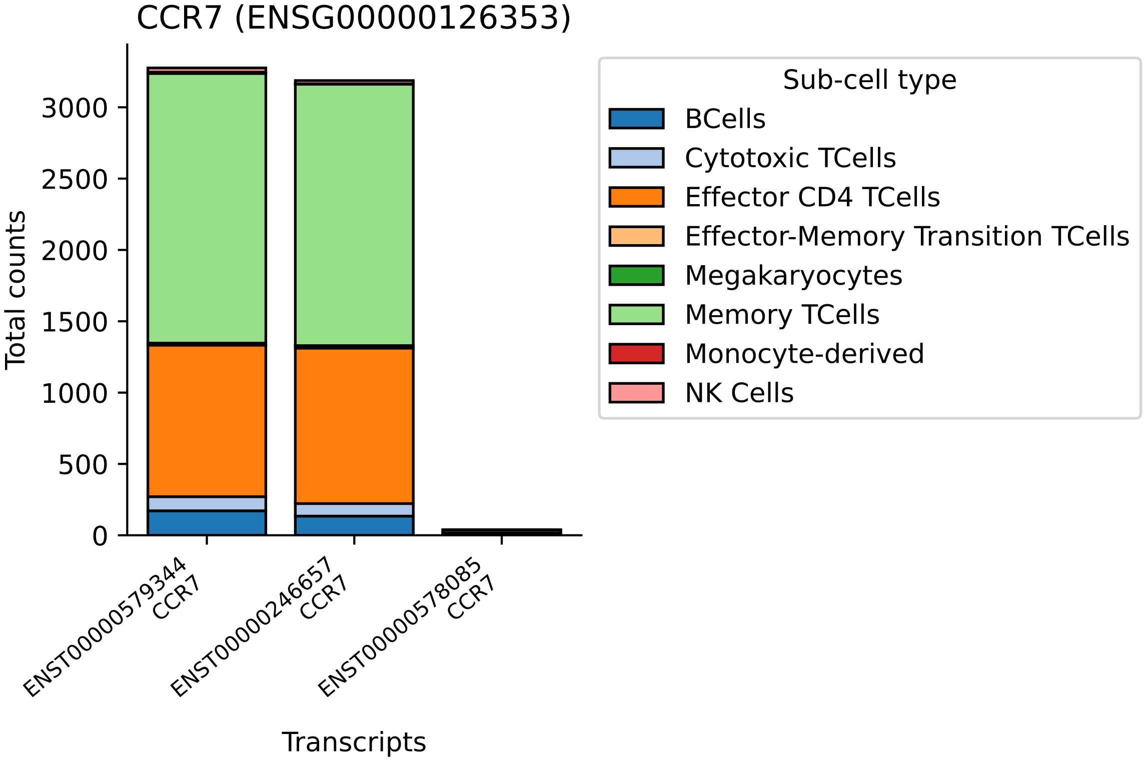
CCR7 (ENSG00000126353)
Total counts
3000
2500
2000
1500
1000
500
0
ENST00000579344
CCR7
ENST00000246657
CCR7
ENST00000578085
CCR7
Transcripts
Sub-cell type
BCells
Cytotoxic TCells
Effector CD4 TCells
Effector-Memory Transition TCells
Megakaryocytes
Memory TCells
Monocyte-derived
NK Cells

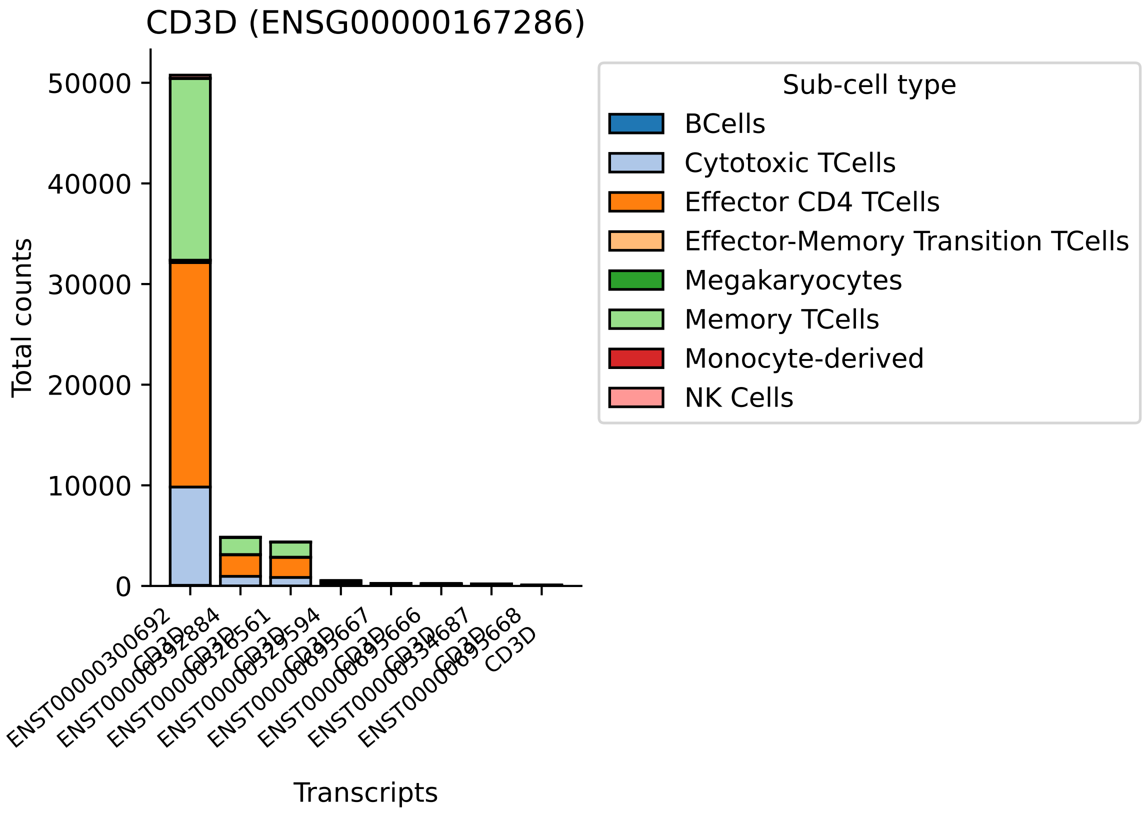

CD3D (ENSG00000167286)
Total counts
0
10000
20000
30000
40000
50000
Transcripts
ENST00000300692
CD3D
Sub-cell type
BCells
Cytotoxic TCells
Effector CD4 TCells
Effector-Memory Transition TCells
Megakaryocytes
Memory TCells
Monocyte-derived
NK Cells

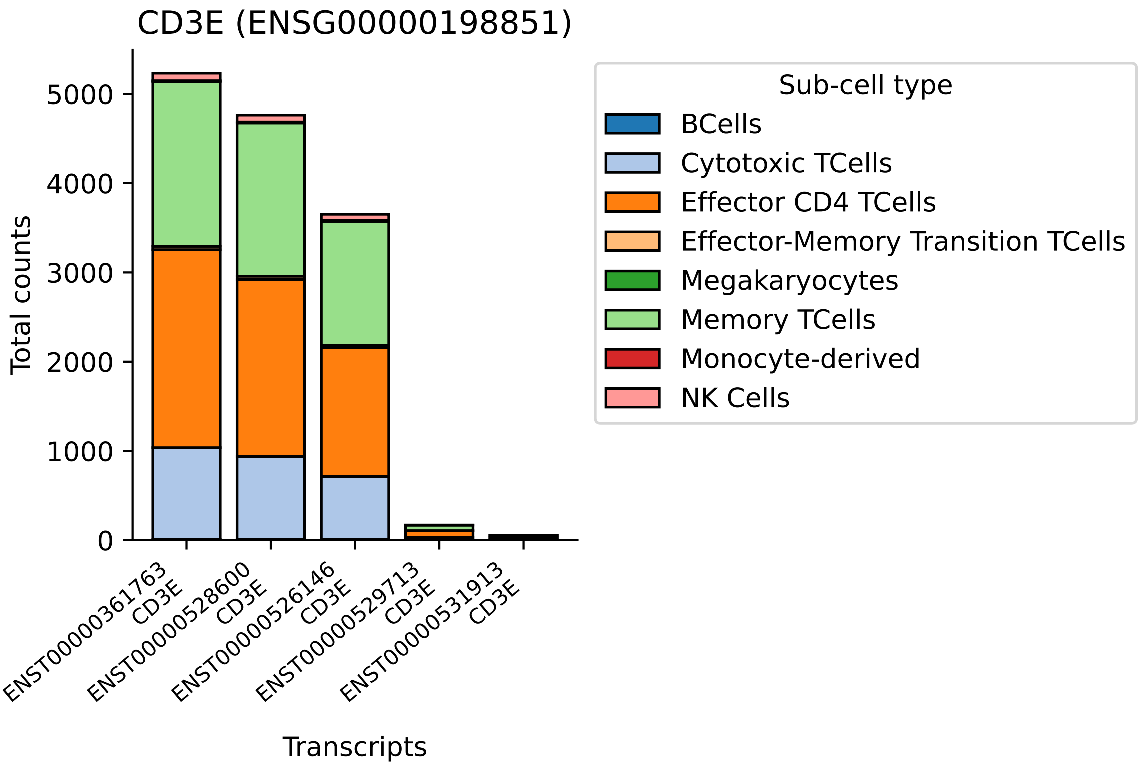

CD3E (ENSG00000198851)
Total counts
0
1000
2000
3000
4000
5000
ENST00000361763 CD3E
ENST00000528600 CD3E
ENST00000526146 CD3E
ENST00000529713 CD3E
ENST00000531913 CD3E
Transcripts
Sub-cell type
BCells
Cytotoxic TCells
Effector CD4 TCells
Effector-Memory Transition TCells
Megakaryocytes
Memory TCells
Monocyte-derived
NK Cells

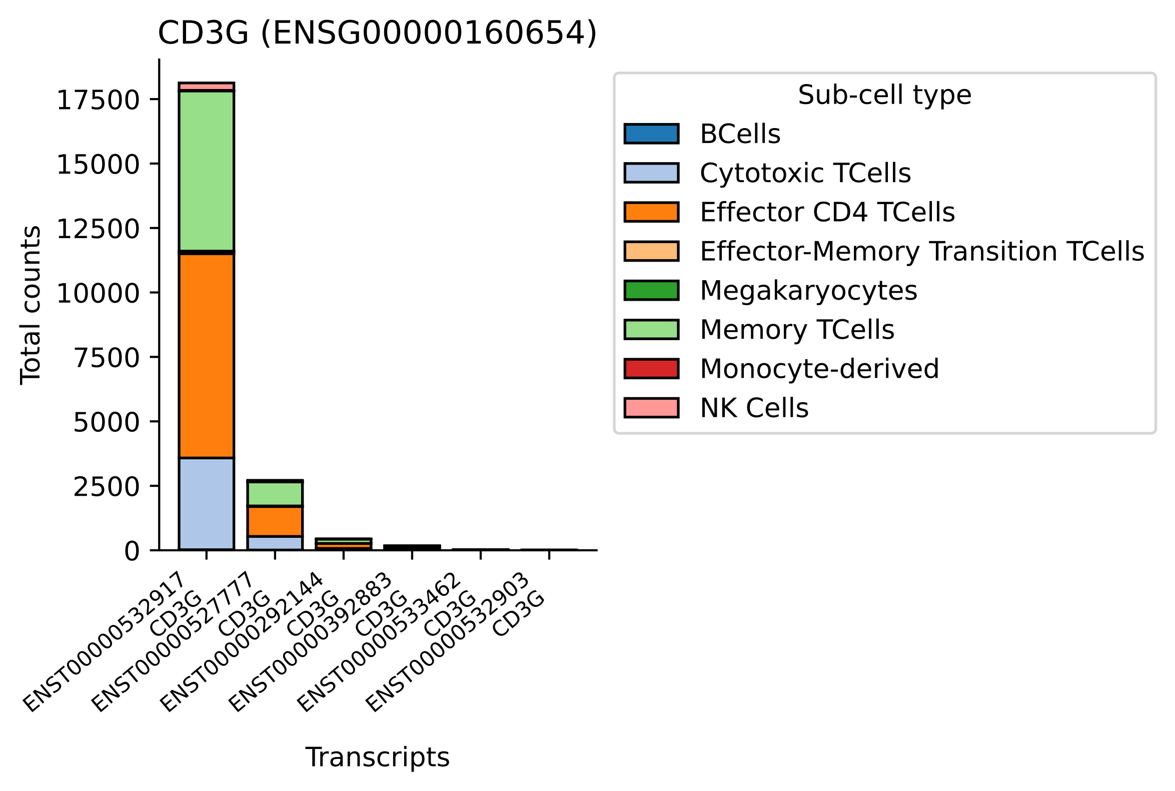

CD3G (ENSG00000160654)
Total counts
0
2500
5000
7500
10000
12500
15000
17500
ENST00000532917 CD3G
ENST00000527777 CD3G
ENST00000292144 CD3G
ENST00000392883 CD3G
ENST00000533462 CD3G
ENST00000532903 CD3G
Transcripts
Sub-cell type
BCells
Cytotoxic TCells
Effector CD4 TCells
Effector-Memory Transition TCells
Megakaryocytes
Memory TCells
Monocyte-derived
NK Cells

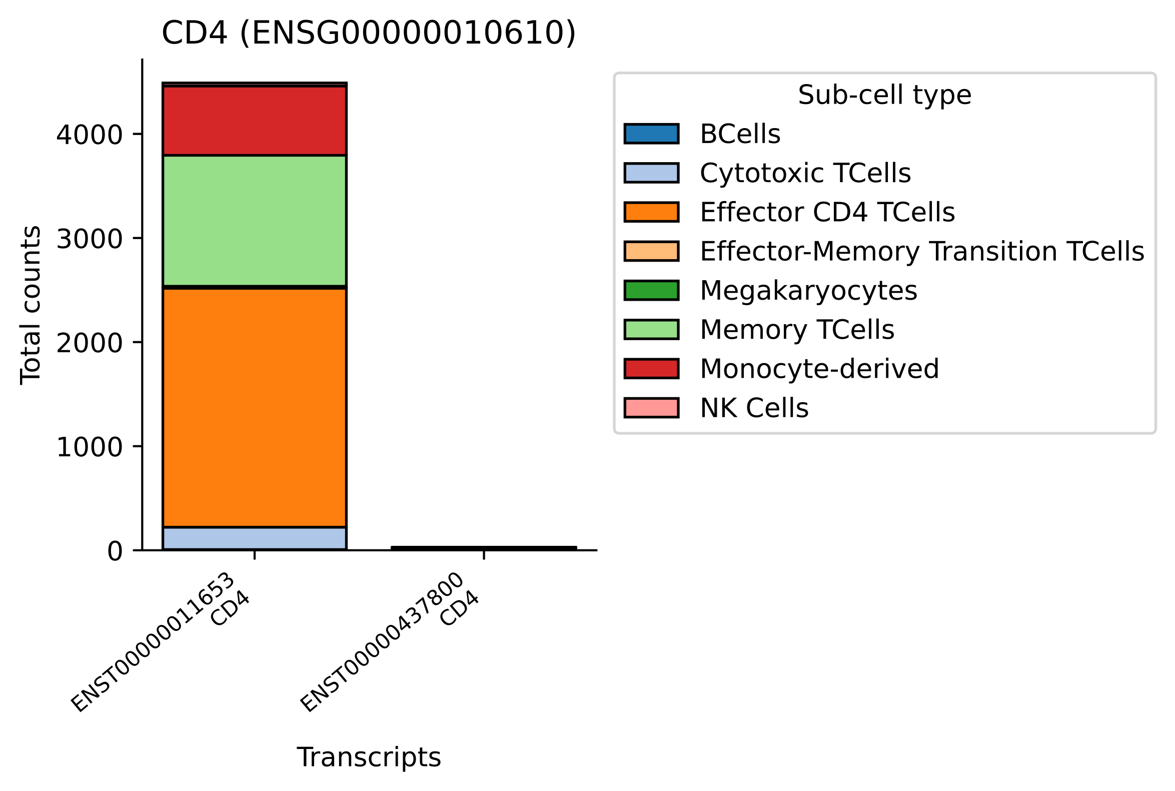
CD4 (ENSG00000010610)
Total counts
0
1000
2000
3000
4000
ENST00000011653
CD4
ENST00000437800
CD4
Transcripts
Sub-cell type
BCells
Cytotoxic TCells
Effector CD4 TCells
Effector-Memory Transition TCells
Megakaryocytes
Memory TCells
Monocyte-derived
NK Cells

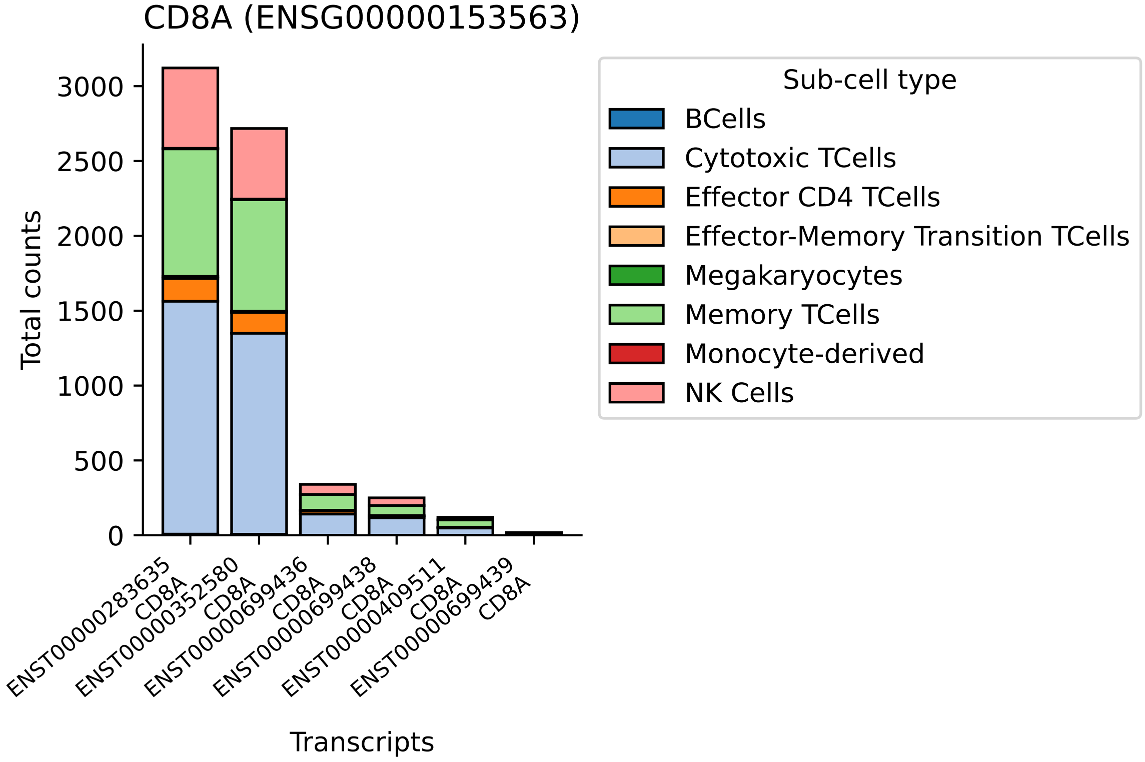
CD8A (ENSG00000153563)
Total counts
0
500
1000
1500
2000
2500
3000
Transcripts
ENST00000283635 CD8A
ENST00000352580 CD8A
ENST00000699436 CD8A
ENST00000699438 CD8A
ENST00000409511 CD8A
ENST00000699439 CD8A
Sub-cell type
BCells
Cytotoxic TCells
Effector CD4 TCells
Effector-Memory Transition TCells
Megakaryocytes
Memory TCells
Monocyte-derived
NK Cells

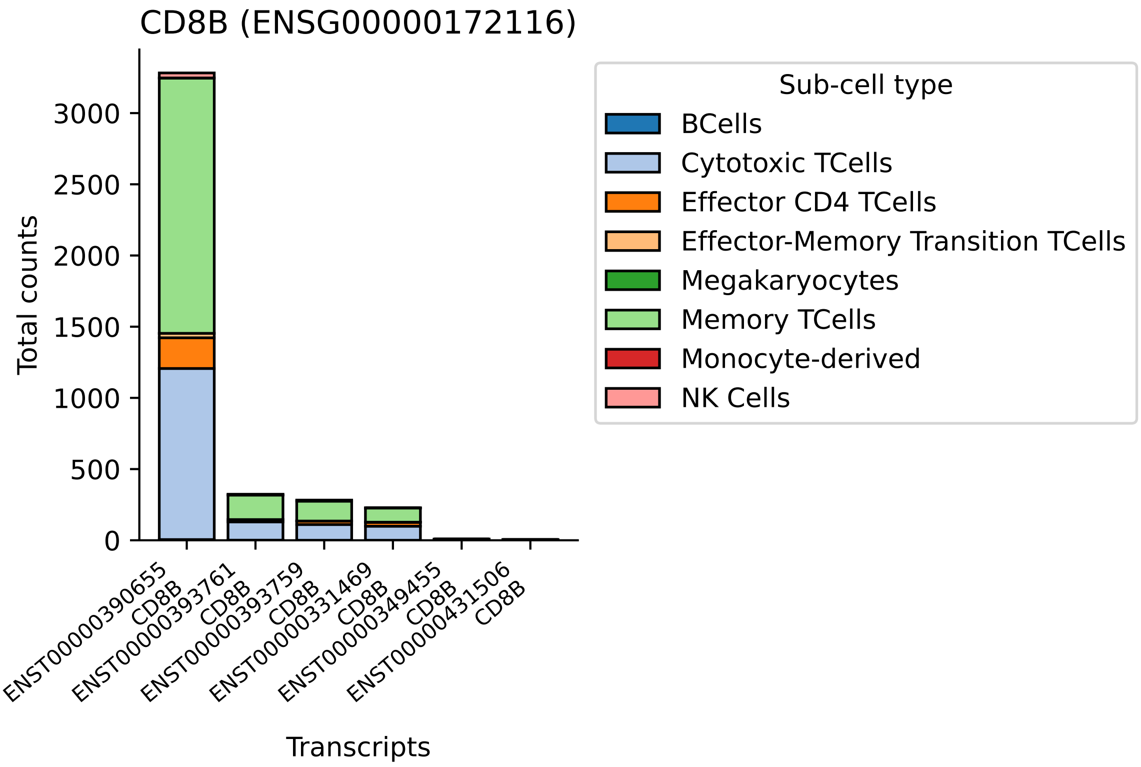

CD8B (ENSG00000172116)
Total counts
0
500
1000
1500
2000
2500
3000
ENST00000390655
CD8B
ENST00000393761
CD8B
ENST00000393759
CD8B
ENST00000331469
CD8B
ENST00000349455
CD8B
ENST00000431506
CD8B
Transcripts
Sub-cell type
BCells
Cytotoxic TCells
Effector CD4 TCells
Effector-Memory Transition TCells
Megakaryocytes
Memory TCells
Monocyte-derived
NK Cells

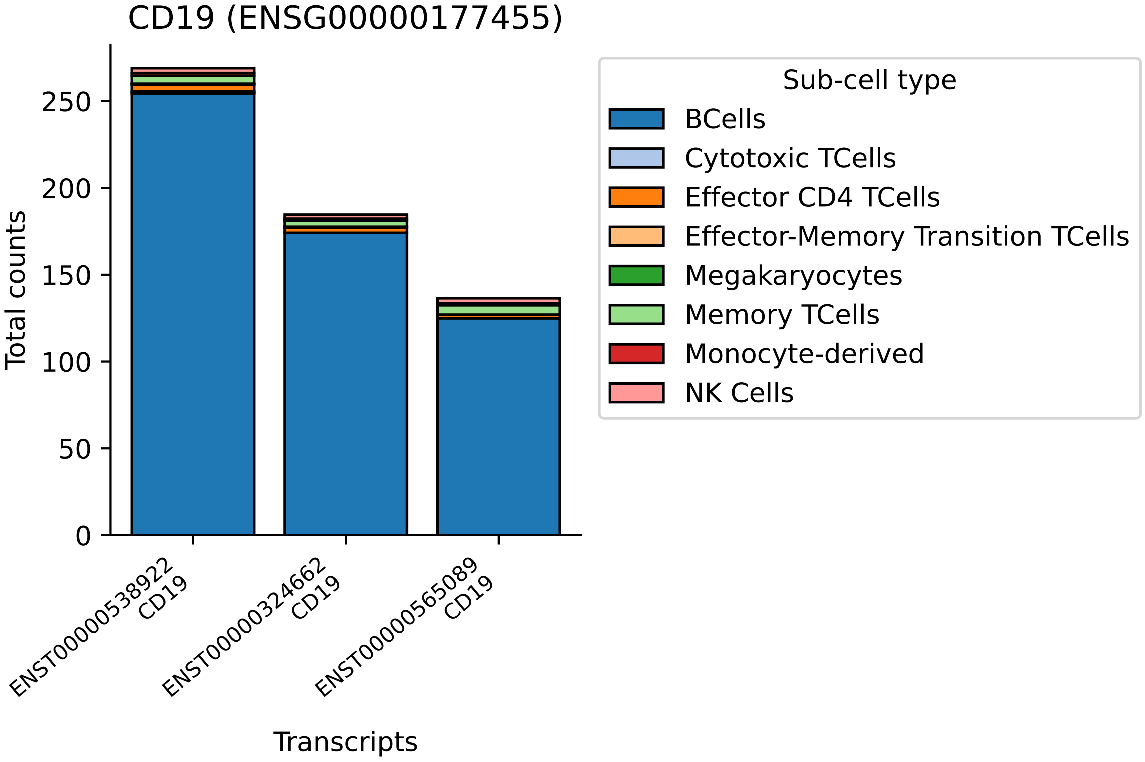

CD19 (ENSG00000177455)
Total counts
0
50
100
150
200
250
ENST00000538922
CD19
ENST00000324662
CD19
ENST00000565089
CD19
Transcripts
Sub-cell type
BCells
Cytotoxic TCells
Effector CD4 TCells
Effector-Memory Transition TCells
Megakaryocytes
Memory TCells
Monocyte-derived
NK Cells

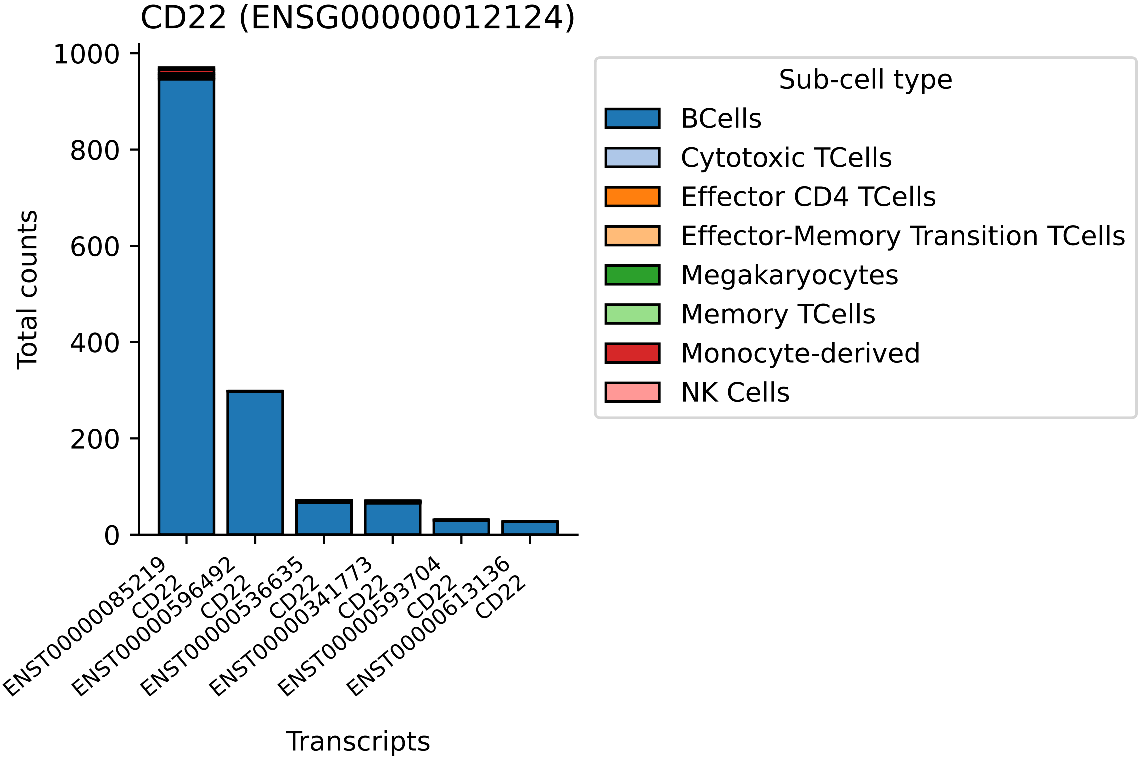

CD22 (ENSG00000012124)
Total counts
1000
800
600
400
200
0
ENST00000085219 CD22
ENST00000596492 CD22
ENST00000536635 CD22
ENST00000341773 CD22
ENST00000593704 CD22
ENST00000613136 CD22
Transcripts
Sub-cell type
BCells
Cytotoxic TCells
Effector CD4 TCells
Effector-Memory Transition TCells
Megakaryocytes
Memory TCells
Monocyte-derived
NK Cells

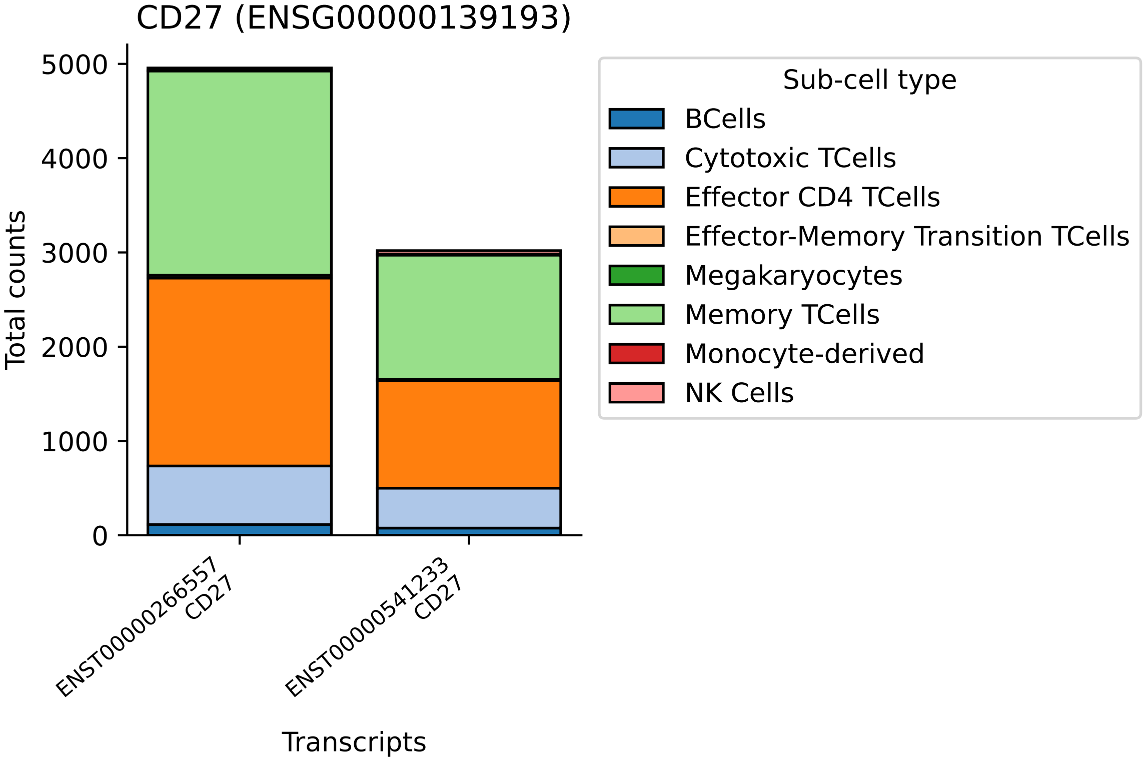

CD27 (ENSG00000139193)
Total counts
0
1000
2000
3000
4000
5000
ENST00000266557
CD27
ENST00000541233
CD27
Transcripts
Sub-cell type
BCells
Cytotoxic TCells
Effector CD4 TCells
Effector-Memory Transition TCells
Megakaryocytes
Memory TCells
Monocyte-derived
NK Cells

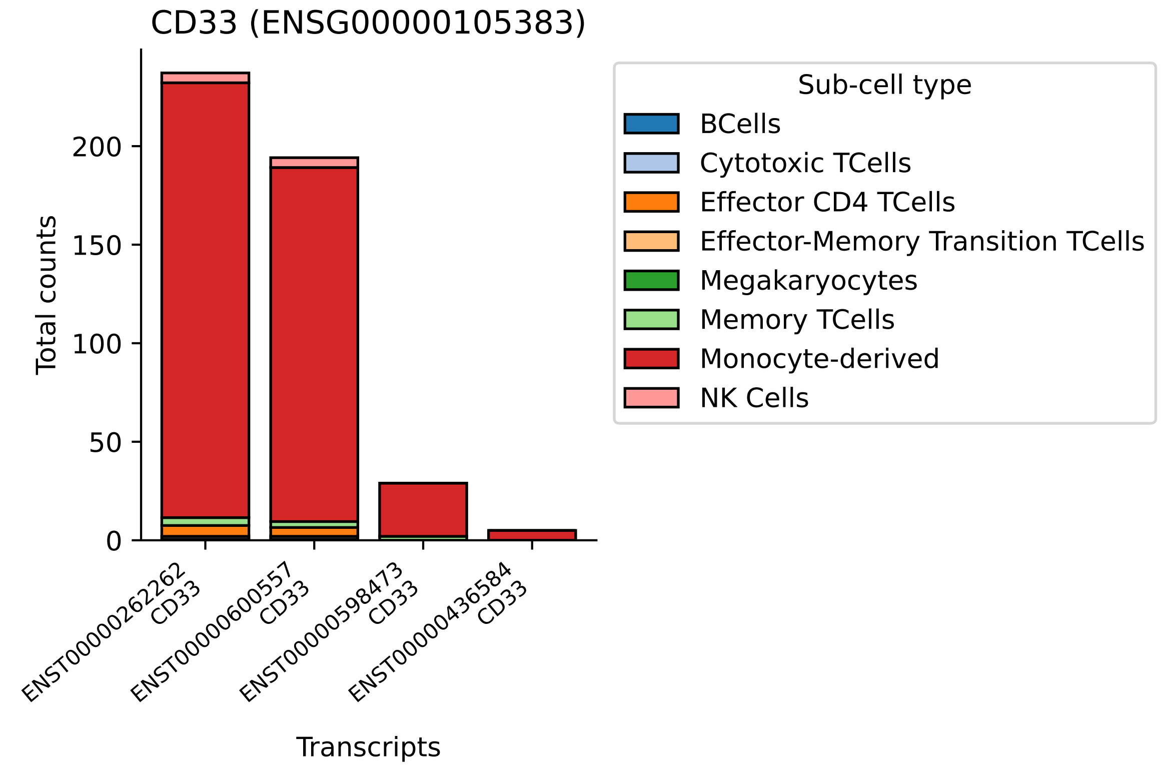
CD33 (ENSG00000105383)
Total counts
0
50
100
150
200
ENST00000262262
CD33
ENST00000600557
CD33
ENST00000598473
CD33
ENST00000436584
CD33
Transcripts
Sub-cell type
BCells
Cytotoxic TCells
Effector CD4 TCells
Effector-Memory Transition TCells
Megakaryocytes
Memory TCells
Monocyte-derived
NK Cells

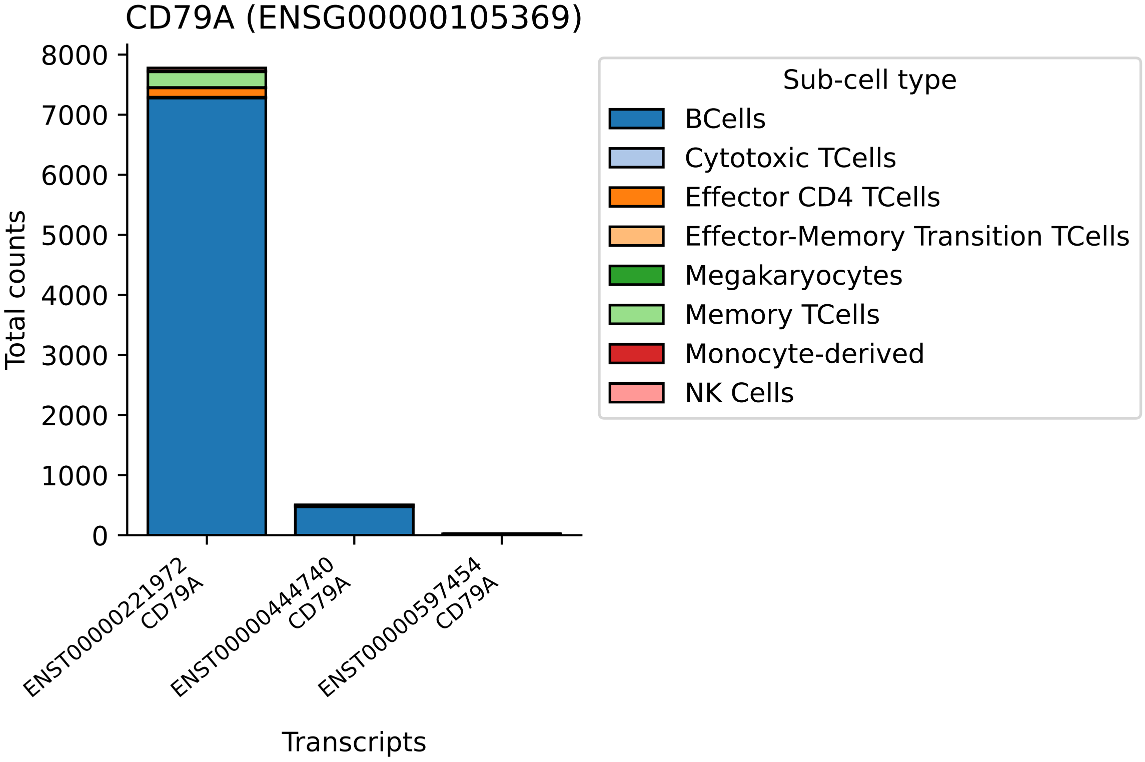

CD79A (ENSG00000105369)
Total counts
0
1000
2000
3000
4000
5000
6000
7000
8000
ENST00000221972
CD79A
ENST00000444740
CD79A
ENST00000597454
CD79A
Transcripts
Sub-cell type
BCells
Cytotoxic TCells
Effector CD4 TCells
Effector-Memory Transition TCells
Megakaryocytes
Memory TCells
Monocyte-derived
NK Cells

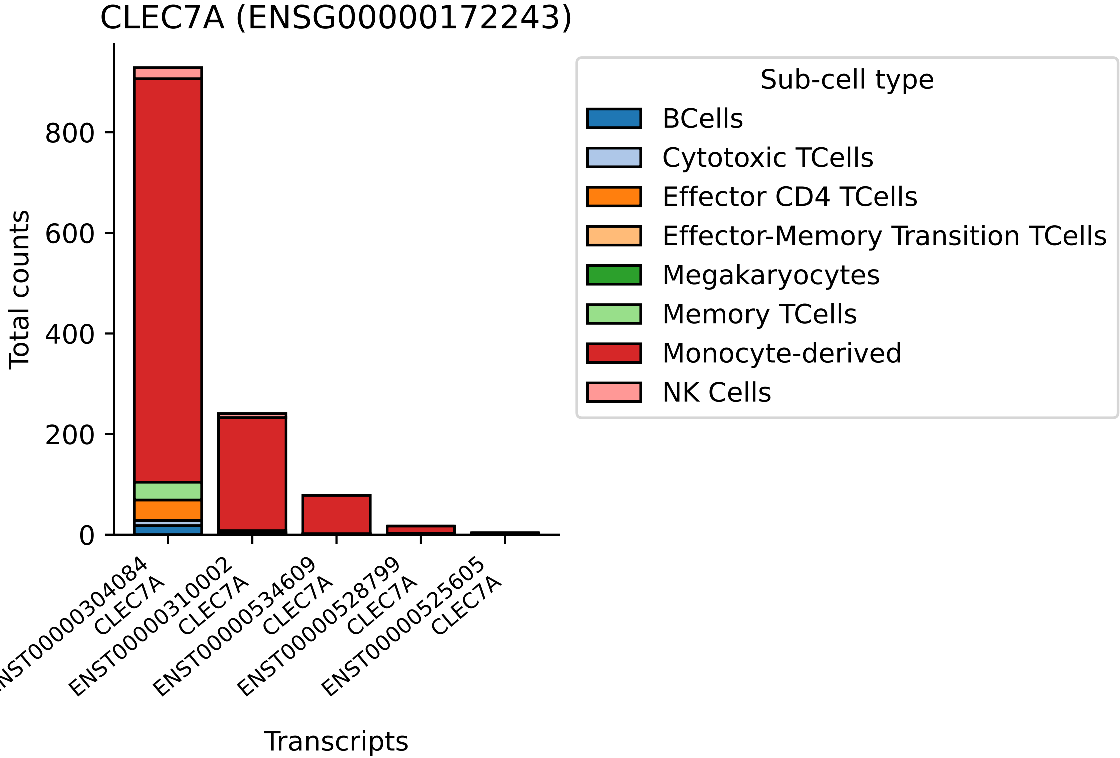
CLEC7A (ENSG00000172243)
Total counts
0
200
400
600
800
ENST00000304084 CLEC7A
ENST00000310002 CLEC7A
ENST00000534609 CLEC7A
ENST00000528799 CLEC7A
ENST00000525605 CLEC7A
Transcripts
Sub-cell type
BCells
Cytotoxic TCells
Effector CD4 TCells
Effector-Memory Transition TCells
Megakaryocytes
Memory TCells
Monocyte-derived
NK Cells

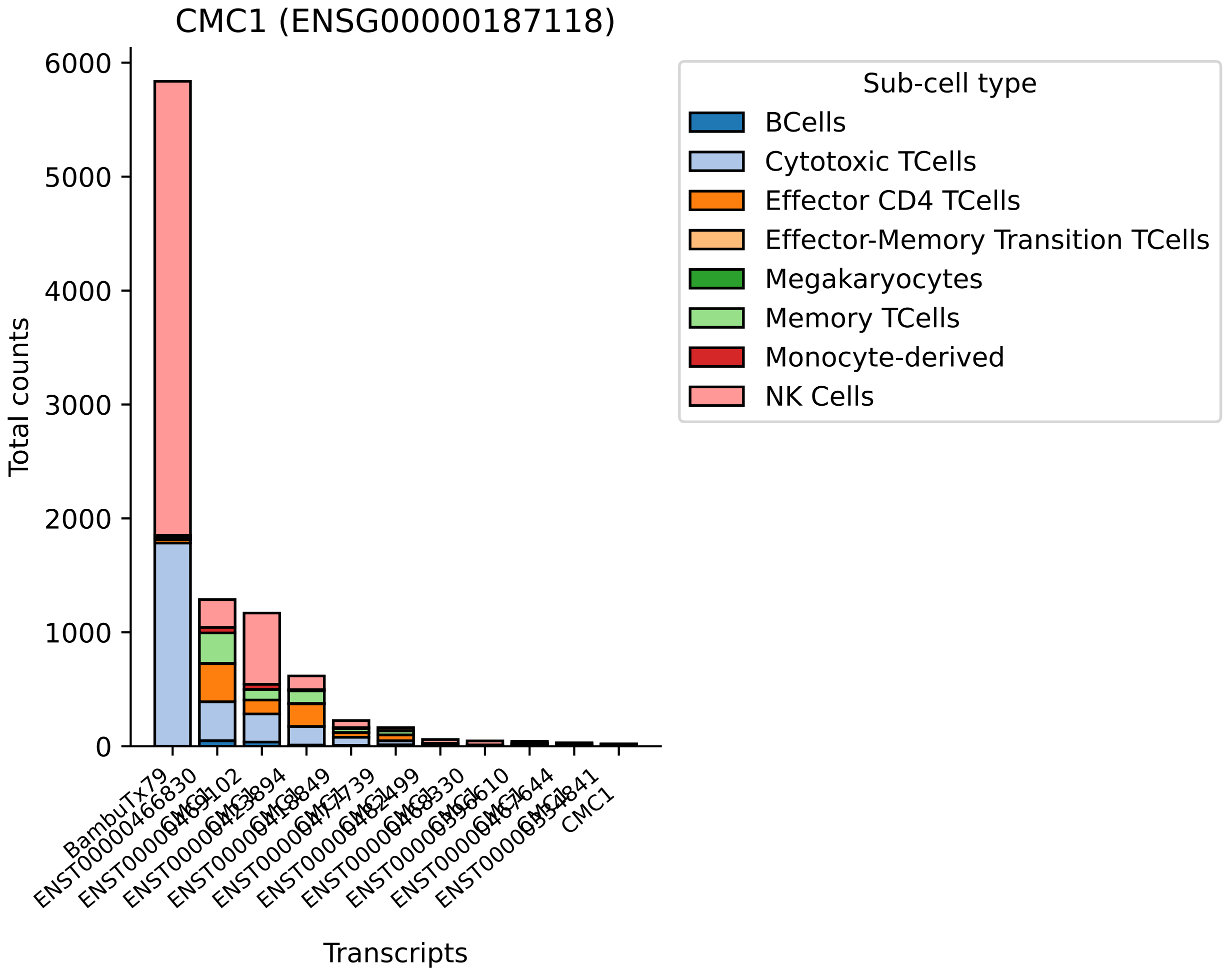

CMC1 (ENSG00000187118)
Total counts
6000
5000
4000
3000
2000
1000
0
Transcripts
BambuTx79
CMC1
Sub-cell type
BCells
Cytotoxic TCells
Effector CD4 TCells
Effector-Memory Transition TCells
Megakaryocytes
Memory TCells
Monocyte-derived
NK Cells

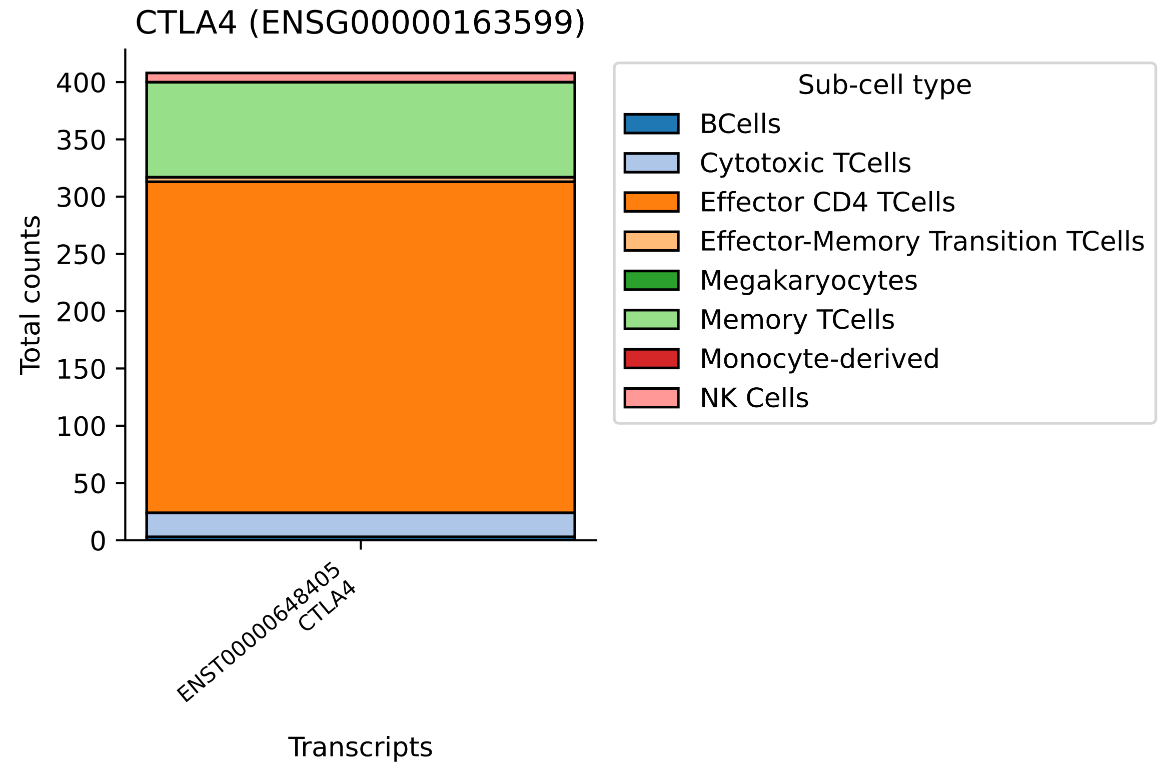

CTLA4 (ENSG00000163599)
Total counts
0
50
100
150
200
250
300
350
400
ENST00000648405
CTLA4
Transcripts
Sub-cell type
BCells
Cytotoxic TCells
Effector CD4 TCells
Effector-Memory Transition TCells
Megakaryocytes
Memory TCells
Monocyte-derived
NK Cells

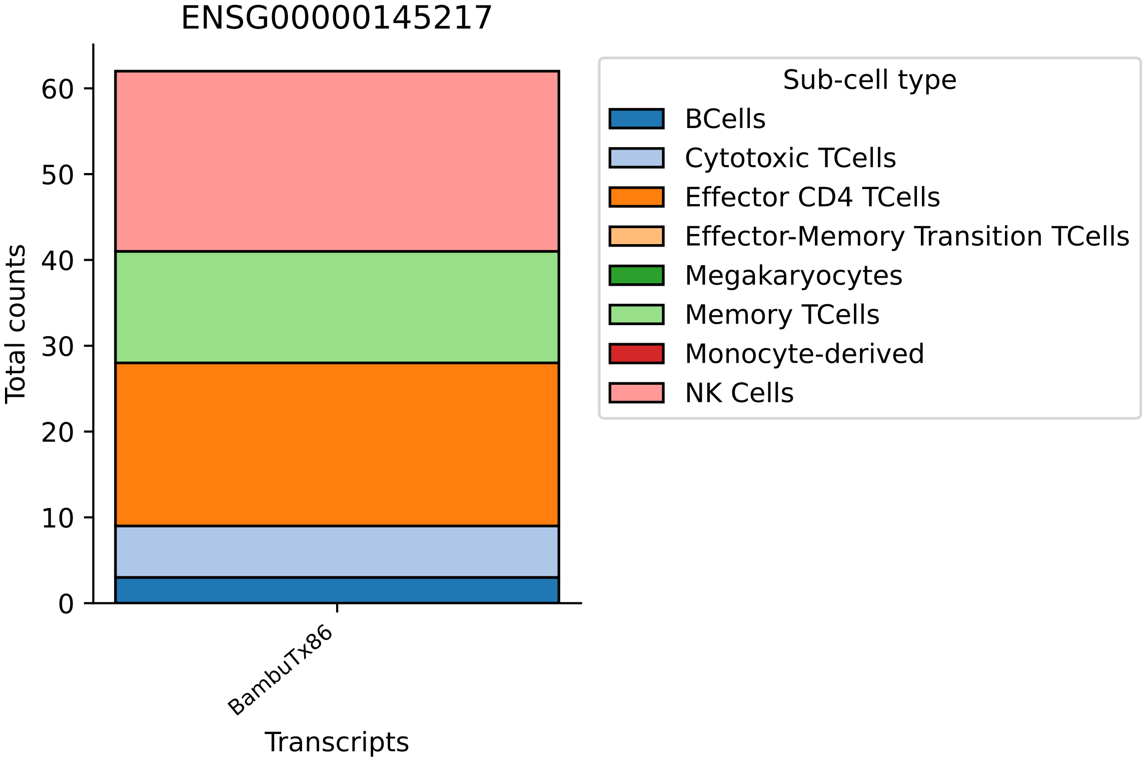

ENSG00000145217
Total counts
0
10
20
30
40
50
60
BambuTx86
Transcripts
Sub-cell type
BCells
Cytotoxic TCells
Effector CD4 TCells
Effector-Memory Transition TCells
Megakaryocytes
Memory TCells
Monocyte-derived
NK Cells

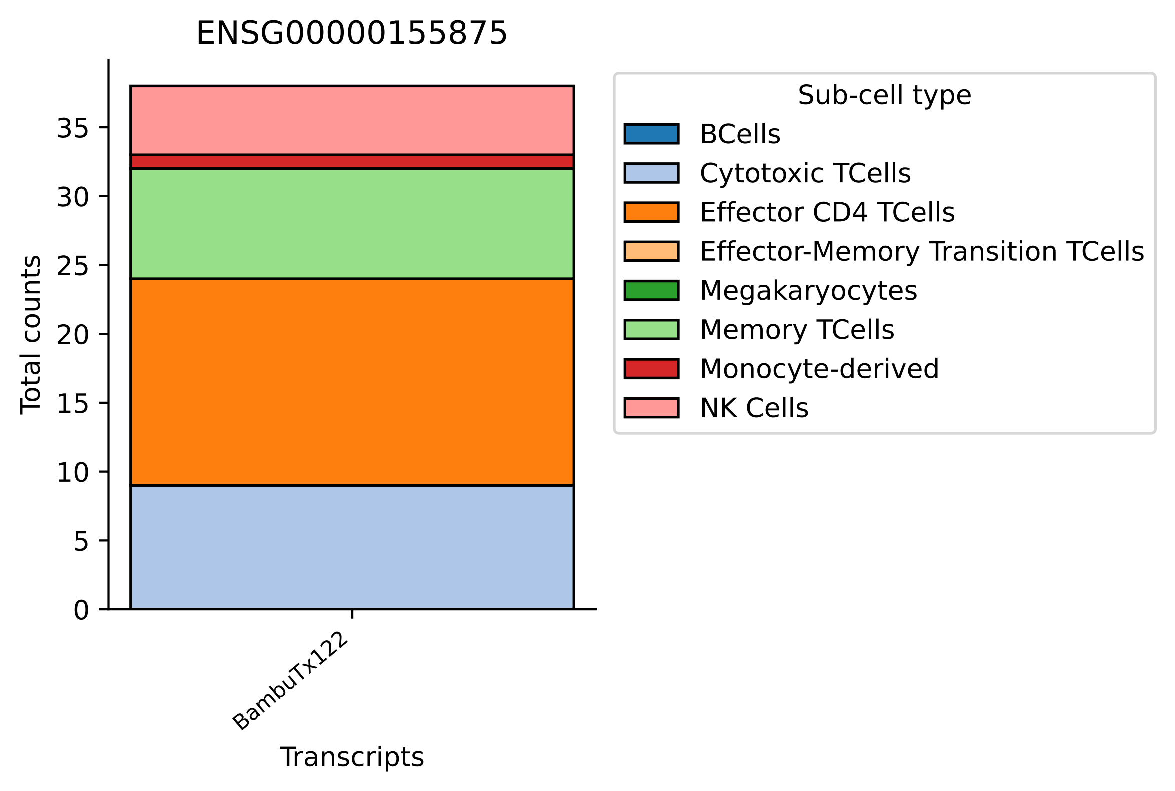
ENSG00000155875
Total counts
0
5
10
15
20
25
30
35
BambuTx122
Transcripts
Sub-cell type
BCells
Cytotoxic TCells
Effector CD4 TCells
Effector-Memory Transition TCells
Megakaryocytes
Memory TCells
Monocyte-derived
NK Cells

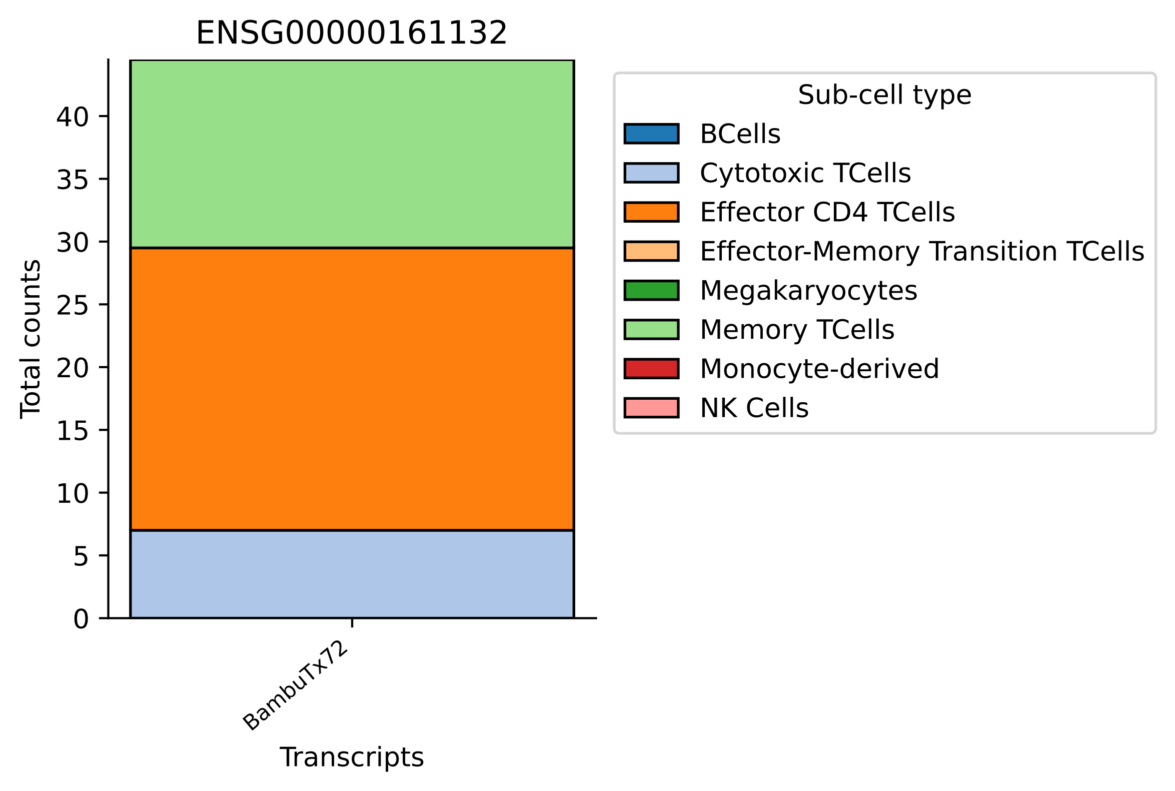

ENSG00000161132
Total counts
0
5
10
15
20
25
30
35
40
BambuTx72
Transcripts
Sub-cell type
BCells
Cytotoxic TCells
Effector CD4 TCells
Effector-Memory Transition TCells
Megakaryocytes
Memory TCells
Monocyte-derived
NK Cells

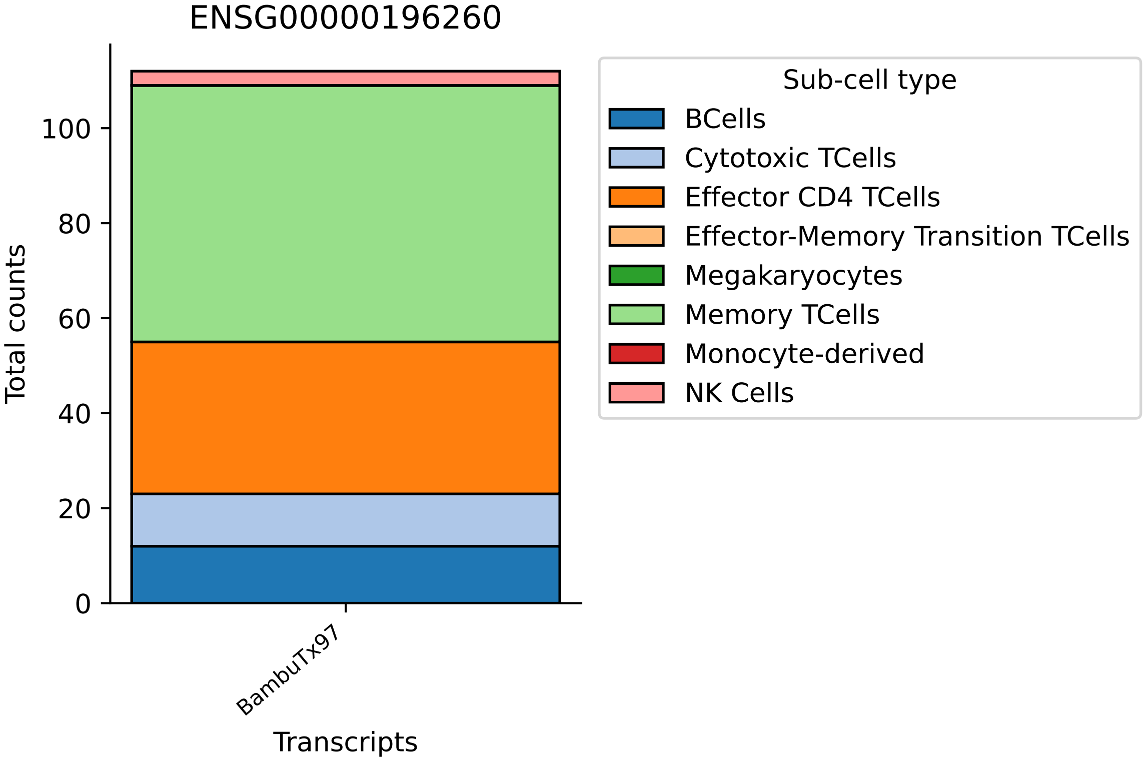

ENSG00000196260
Total counts
0
20
40
60
80
100
BambuTx97
Transcripts
Sub-cell type
BCells
Cytotoxic TCells
Effector CD4 TCells
Effector-Memory Transition TCells
Megakaryocytes
Memory TCells
Monocyte-derived
NK Cells

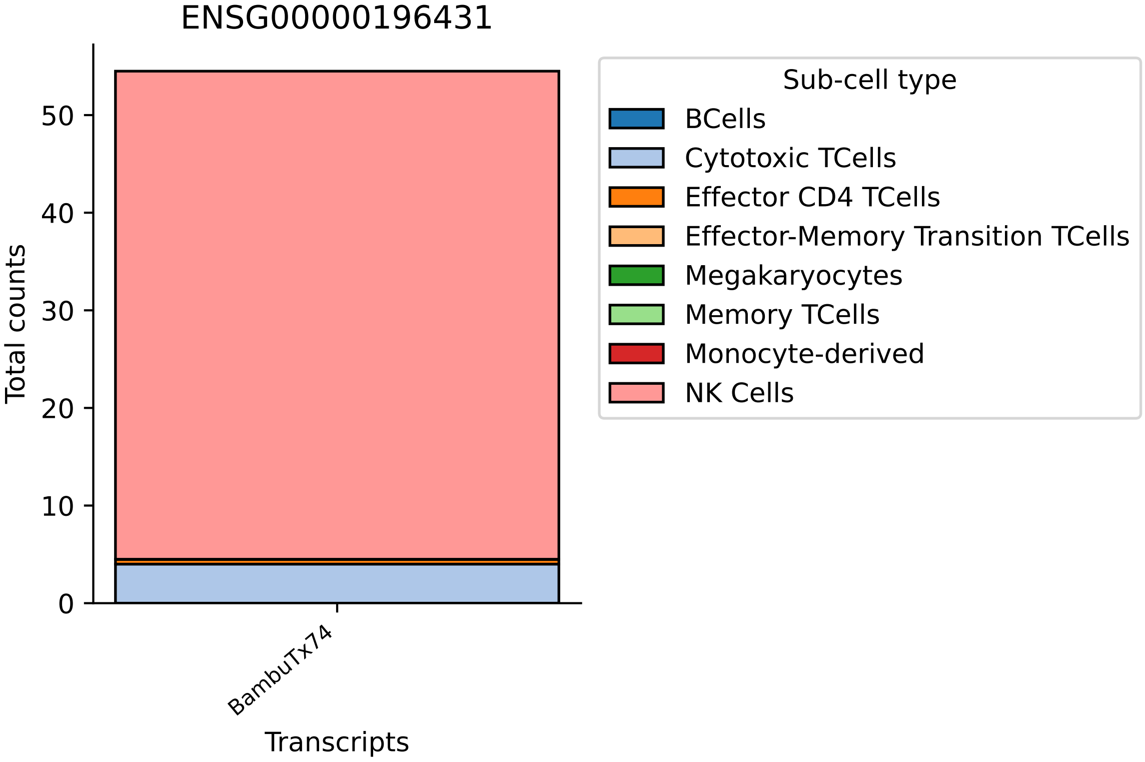

ENSG00000196431
Total counts
0
10
20
30
40
50
BambuTx74
Transcripts
Sub-cell type
BCells
Cytotoxic TCells
Effector CD4 TCells
Effector-Memory Transition TCells
Megakaryocytes
Memory TCells
Monocyte-derived
NK Cells

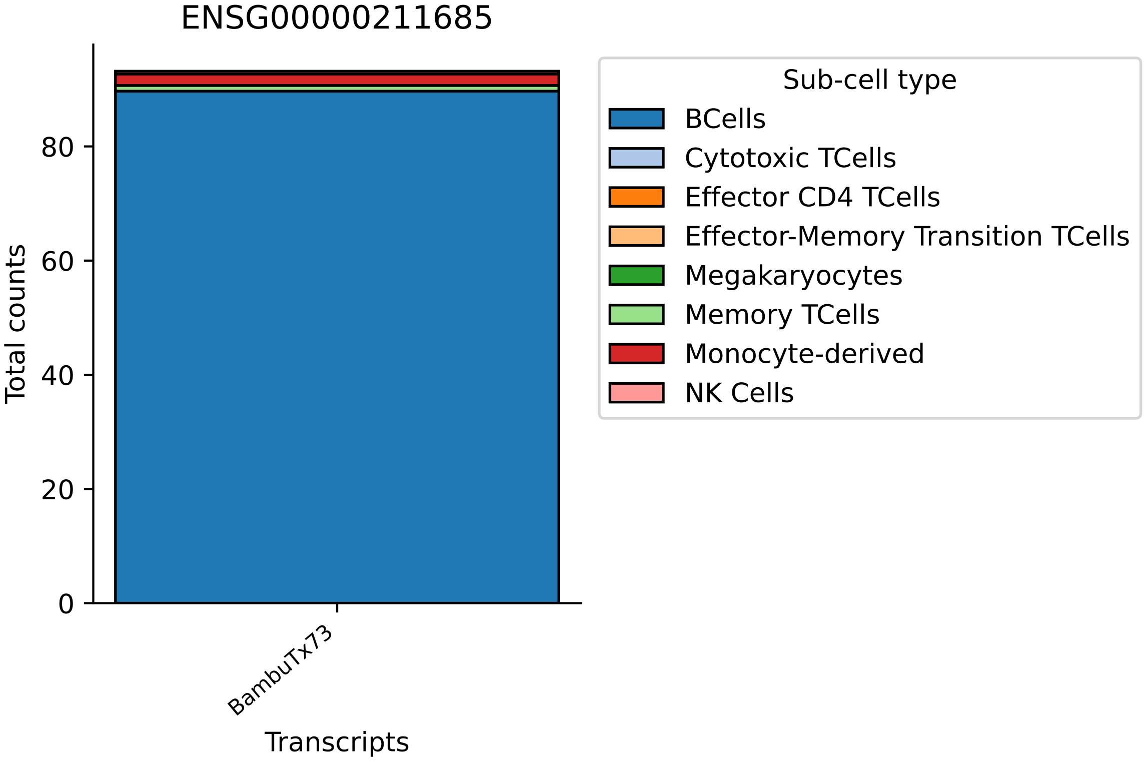

ENSG00000211685
Total counts
0
20
40
60
80
BambuTx73
Transcripts
Sub-cell type
BCells
Cytotoxic TCells
Effector CD4 TCells
Effector-Memory Transition TCells
Megakaryocytes
Memory TCells
Monocyte-derived
NK Cells

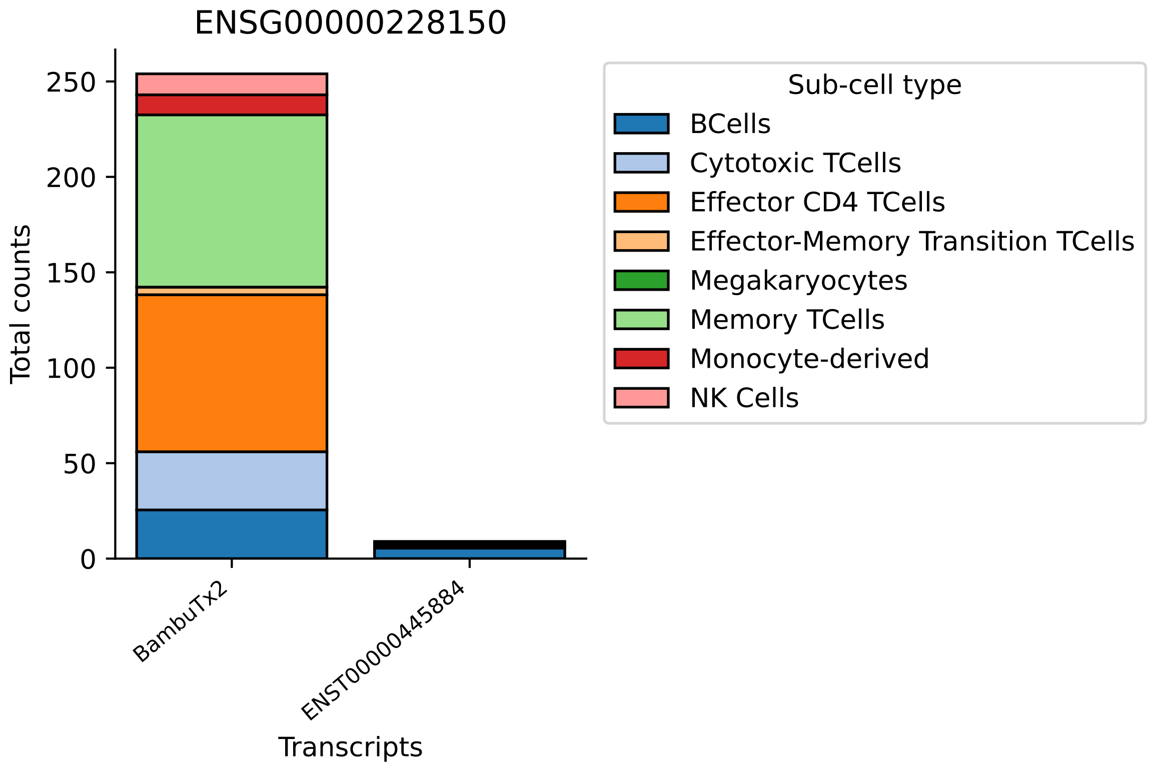
ENSG00000228150
Total counts
0
50
100
150
200
250
BambuTx2
ENST00000445884
Transcripts
Sub-cell type
BCells
Cytotoxic TCells
Effector CD4 TCells
Effector-Memory Transition TCells
Megakaryocytes
Memory TCells
Monocyte-derived
NK Cells

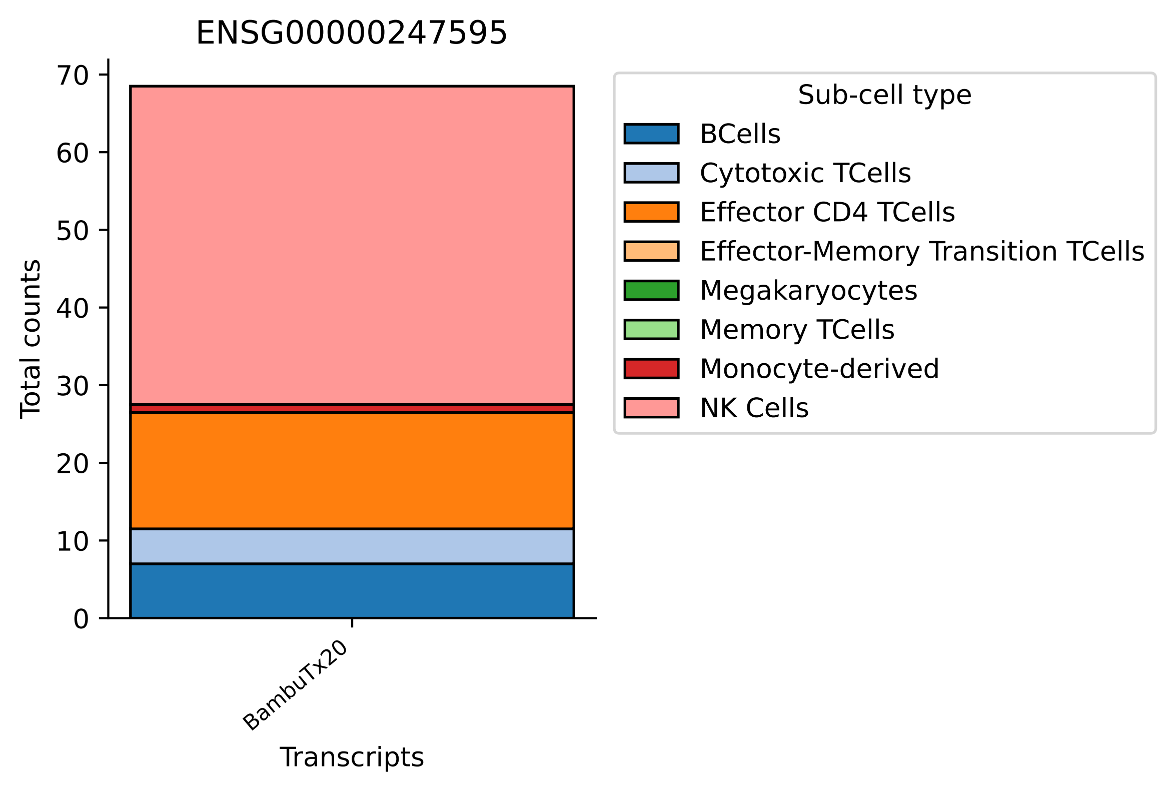
ENSG00000247595
Total counts
0
10
20
30
40
50
60
70
BambuTx20
Transcripts
Sub-cell type
BCells
Cytotoxic TCells
Effector CD4 TCells
Effector-Memory Transition TCells
Megakaryocytes
Memory TCells
Monocyte-derived
NK Cells

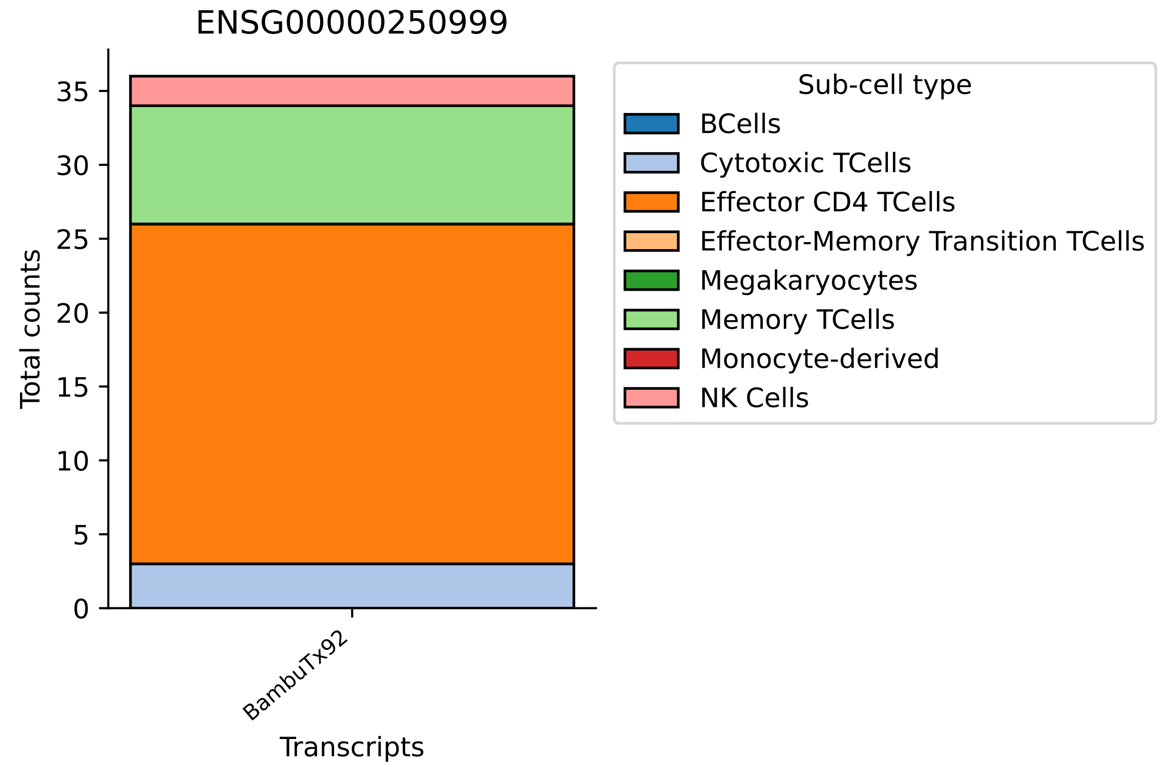

ENSG00000250999
Total counts
0
5
10
15
20
25
30
35
BambuTx92
Transcripts
Sub-cell type
BCells
Cytotoxic TCells
Effector CD4 TCells
Effector-Memory Transition TCells
Megakaryocytes
Memory TCells
Monocyte-derived
NK Cells

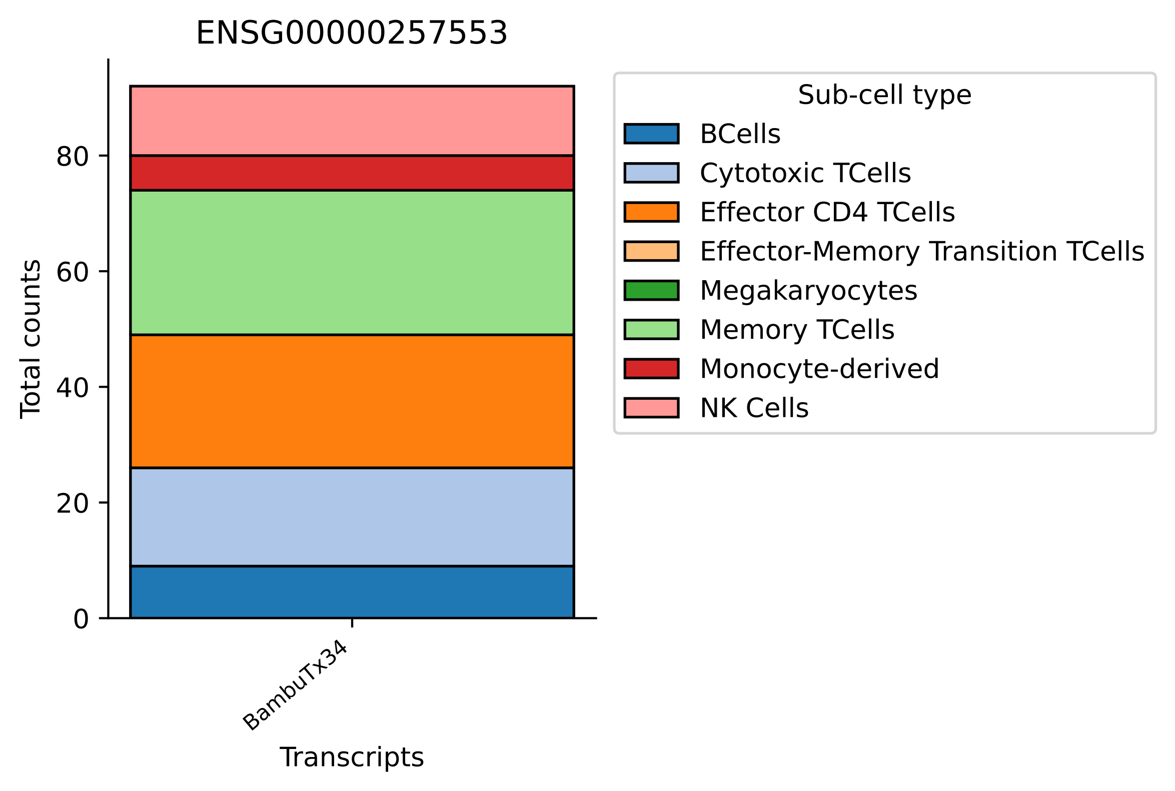
ENSG00000257553
Total counts
0
20
40
60
80
BambuTx34
Transcripts
Sub-cell type
BCells
Cytotoxic TCells
Effector CD4 TCells
Effector-Memory Transition TCells
Megakaryocytes
Memory TCells
Monocyte-derived
NK Cells

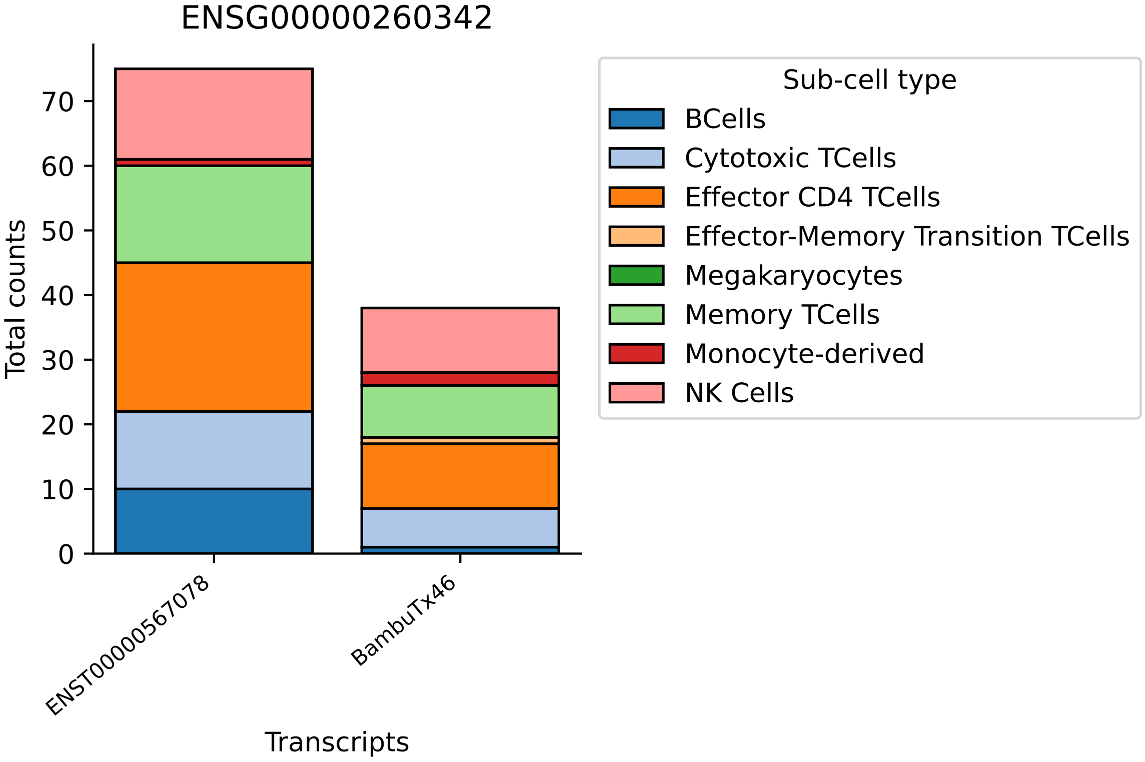

ENSG00000260342
Total counts
0
10
20
30
40
50
60
70
ENST00000567078
BambuTx46
Transcripts
Sub-cell type
BCells
Cytotoxic TCells
Effector CD4 TCells
Effector-Memory Transition TCells
Megakaryocytes
Memory TCells
Monocyte-derived
NK Cells

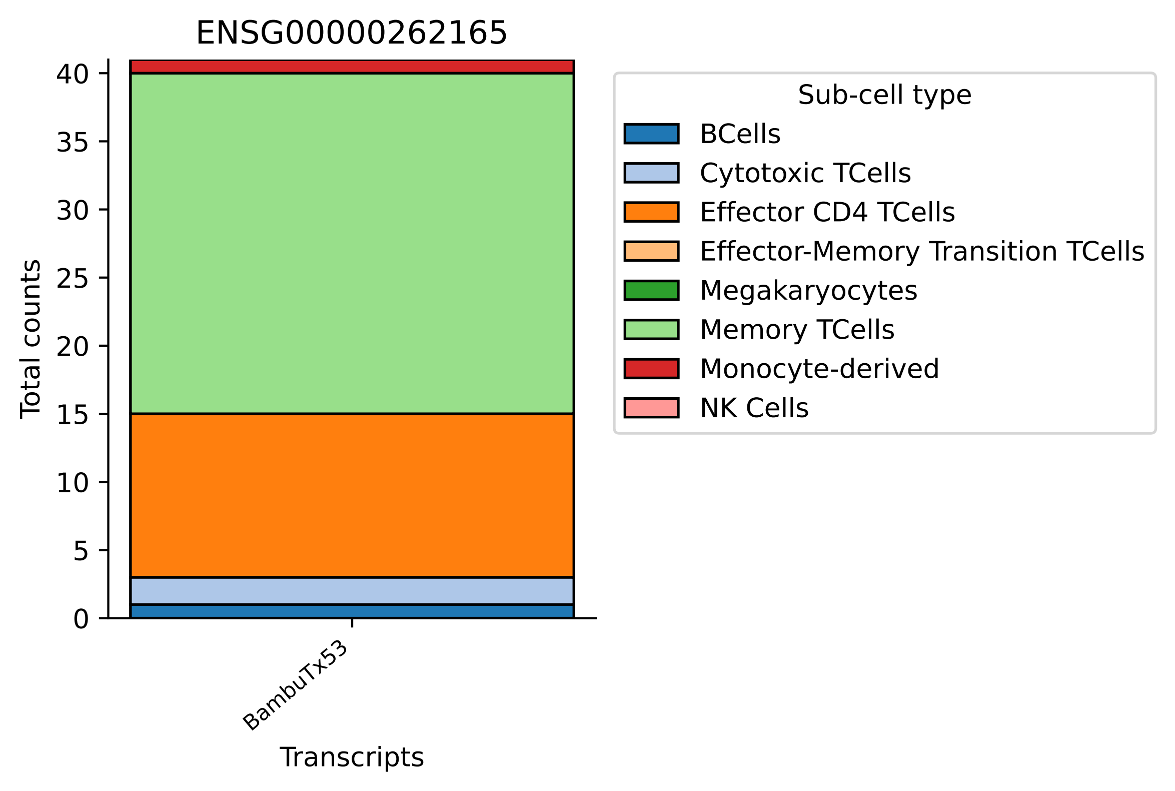

ENSG00000262165
Total counts
0
5
10
15
20
25
30
35
40
BambuTx53
Transcripts
Sub-cell type
BCells
Cytotoxic TCells
Effector CD4 TCells
Effector-Memory Transition TCells
Megakaryocytes
Memory TCells
Monocyte-derived
NK Cells

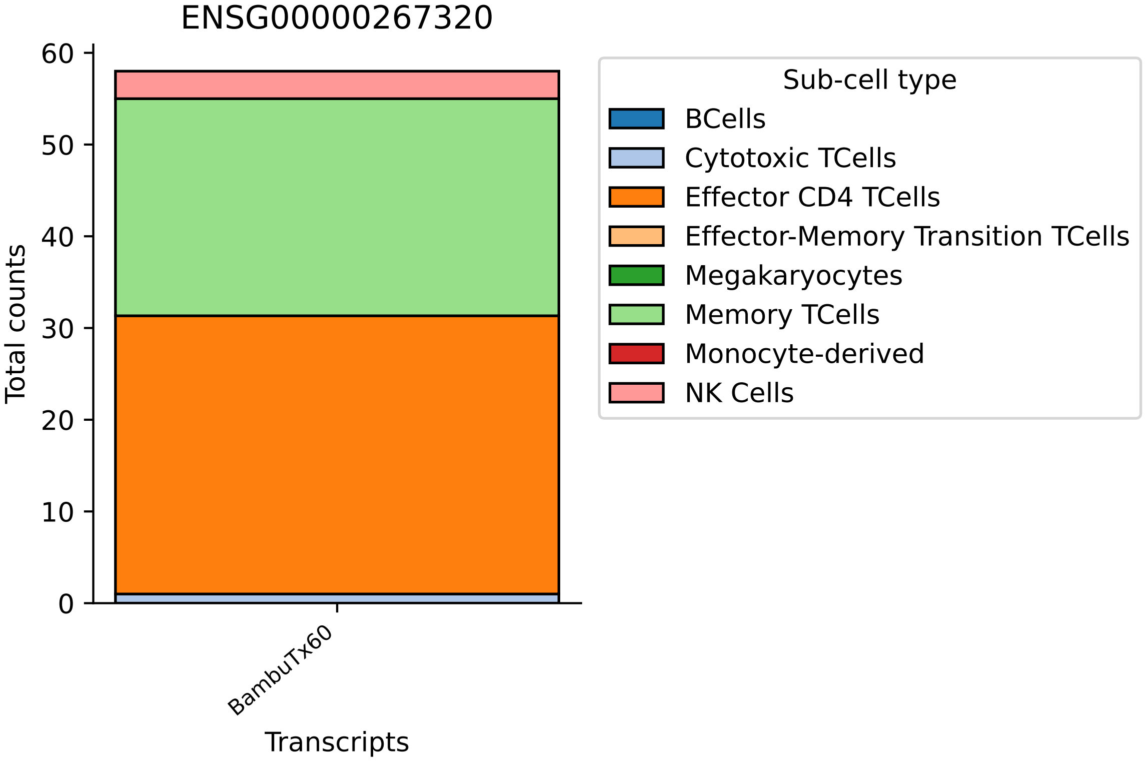
ENSG00000267320
Total counts
0
10
20
30
40
50
60
BambuTx60
Transcripts
Sub-cell type
BCells
Cytotoxic TCells
Effector CD4 TCells
Effector-Memory Transition TCells
Megakaryocytes
Memory TCells
Monocyte-derived
NK Cells

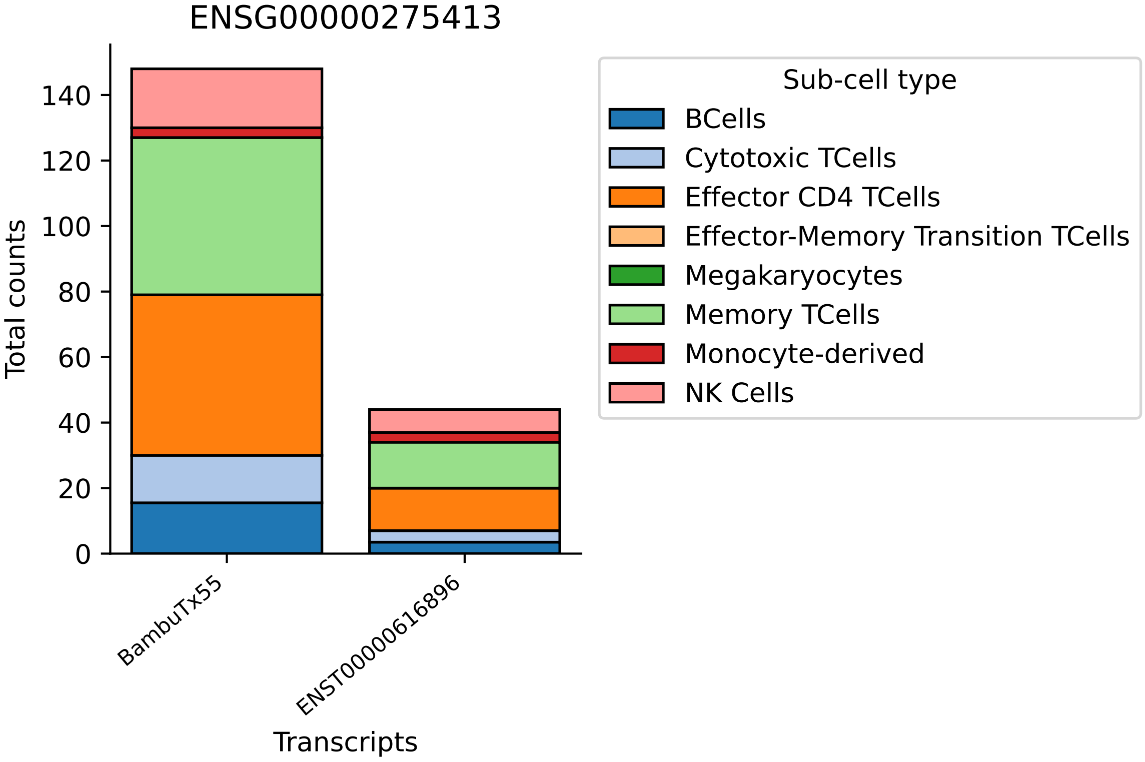
ENSG00000275413
Total counts
0
20
40
60
80
100
120
140
BambuTx55
ENST00000616896
Transcripts
Sub-cell type
BCells
Cytotoxic TCells
Effector CD4 TCells
Effector-Memory Transition TCells
Megakaryocytes
Memory TCells
Monocyte-derived
NK Cells

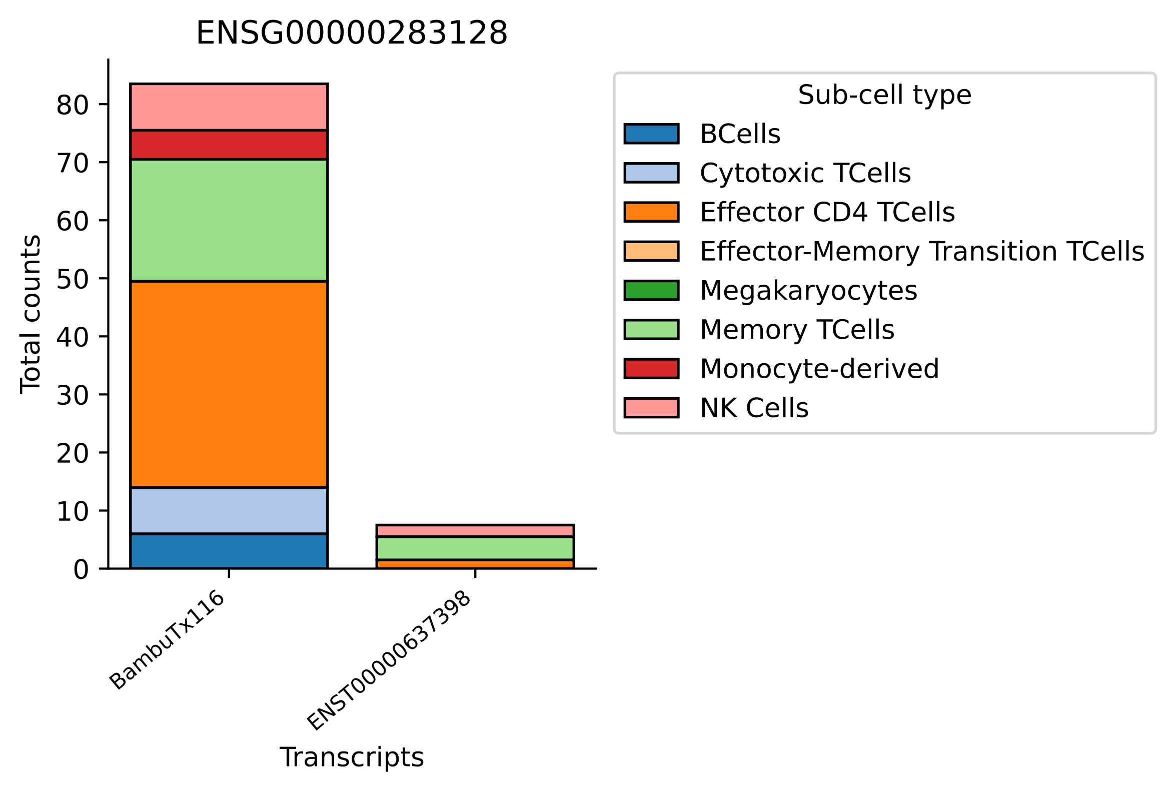

ENSG00000283128
Total counts
0
10
20
30
40
50
60
70
80
BambuTx116
ENST00000637398
Transcripts
Sub-cell type
BCells
Cytotoxic TCells
Effector CD4 TCells
Effector-Memory Transition TCells
Megakaryocytes
Memory TCells
Monocyte-derived
NK Cells

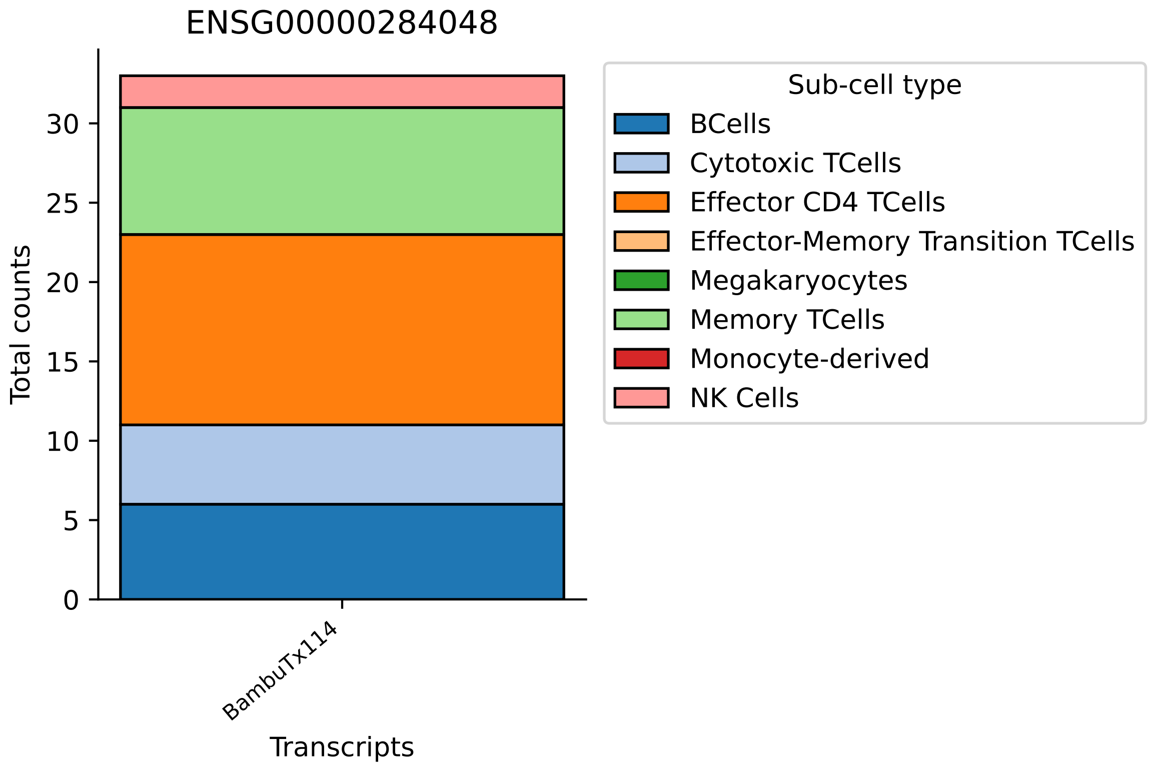

ENSG00000284048
Total counts
0
5
10
15
20
25
30
BambuTx114
Transcripts
Sub-cell type
BCells
Cytotoxic TCells
Effector CD4 TCells
Effector-Memory Transition TCells
Megakaryocytes
Memory TCells
Monocyte-derived
NK Cells

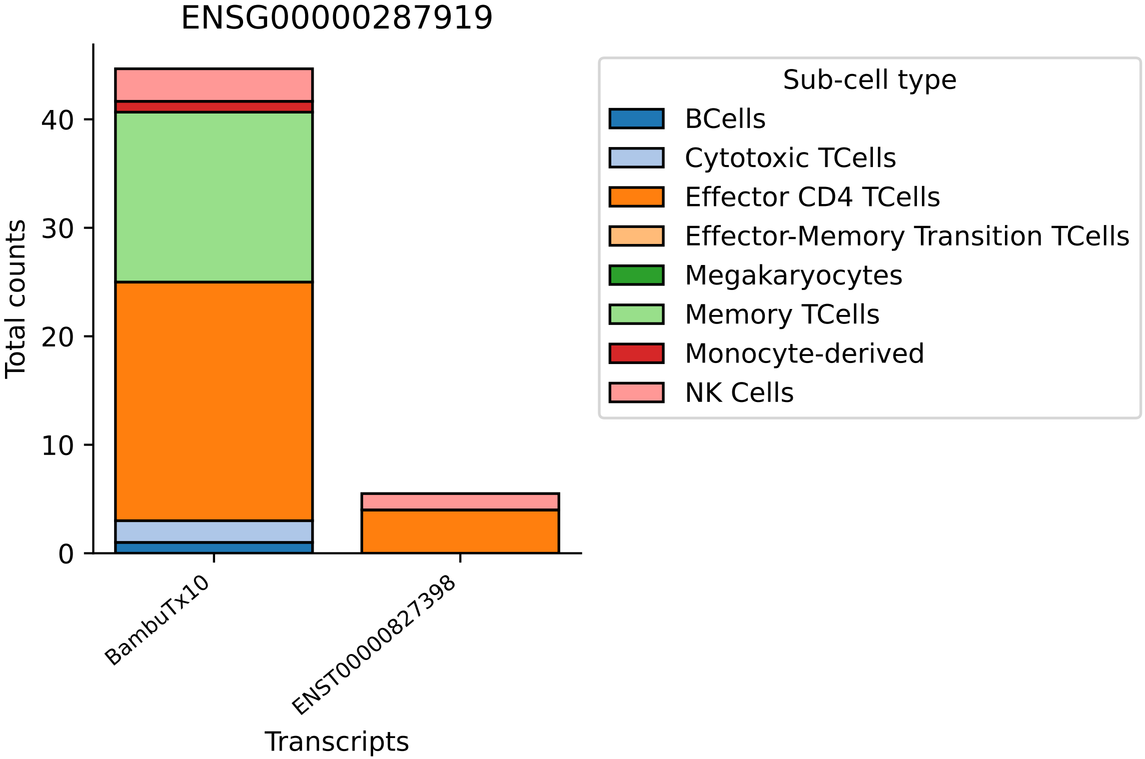
ENSG00000287919
Total counts
0
10
20
30
40
BambuTx10
ENST00000827398
Transcripts
Sub-cell type
BCells
Cytotoxic TCells
Effector CD4 TCells
Effector-Memory Transition TCells
Megakaryocytes
Memory TCells
Monocyte-derived
NK Cells

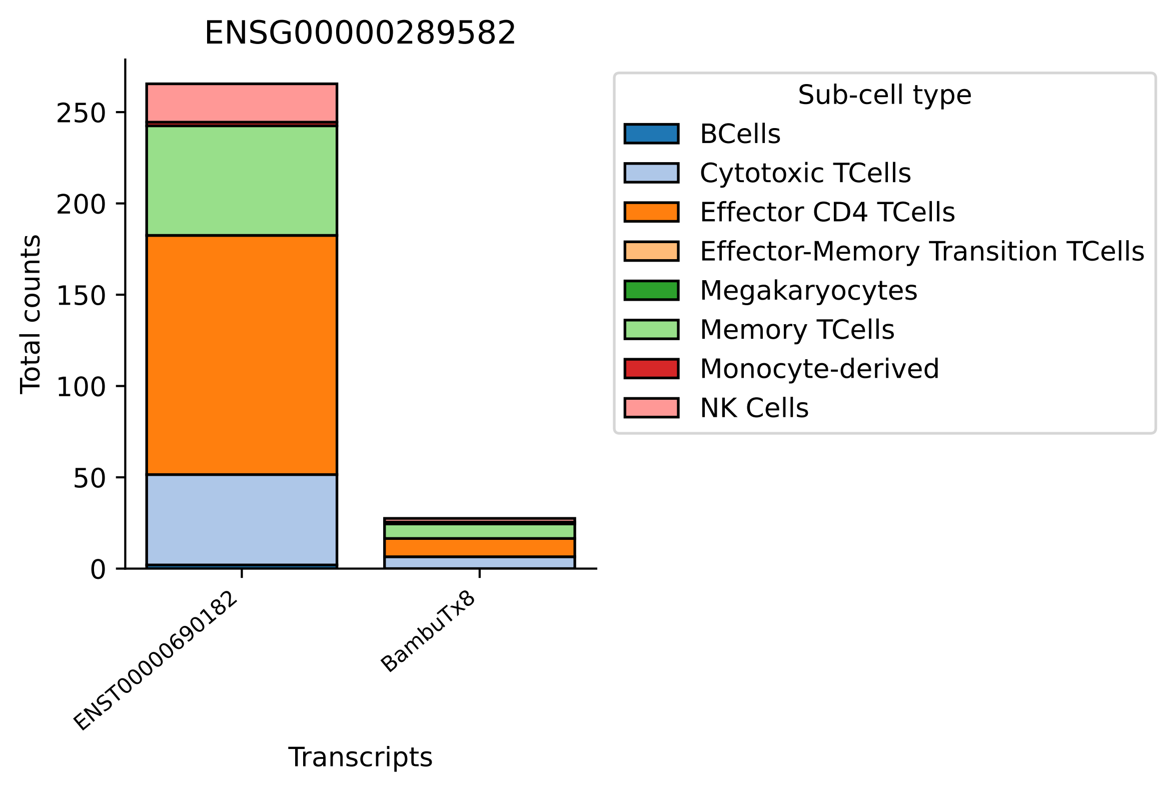
ENSG00000289582
Total counts
0
50
100
150
200
250
ENST00000690182
BambuTx8
Transcripts
Sub-cell type
BCells
Cytotoxic TCells
Effector CD4 TCells
Effector-Memory Transition TCells
Megakaryocytes
Memory TCells
Monocyte-derived
NK Cells

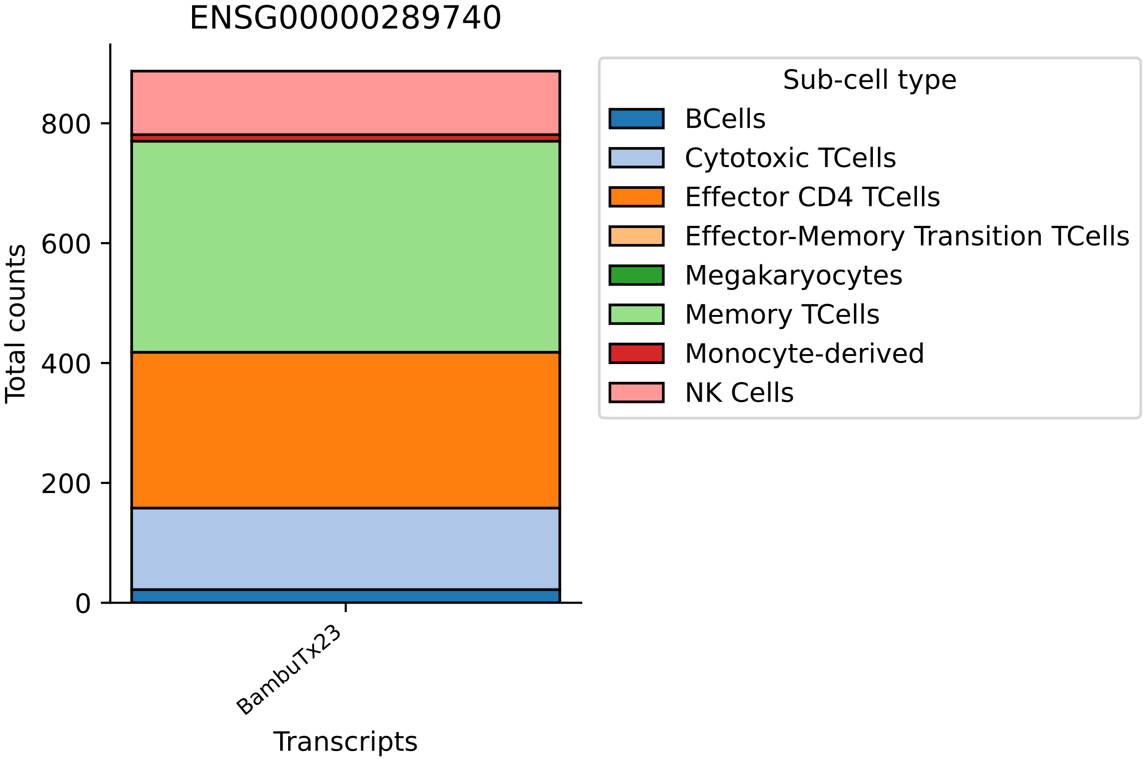
ENSG00000289740
Total counts
0
200
400
600
800
BambuTx23
Transcripts
Sub-cell type
BCells
Cytotoxic TCells
Effector CD4 TCells
Effector-Memory Transition TCells
Megakaryocytes
Memory TCells
Monocyte-derived
NK Cells

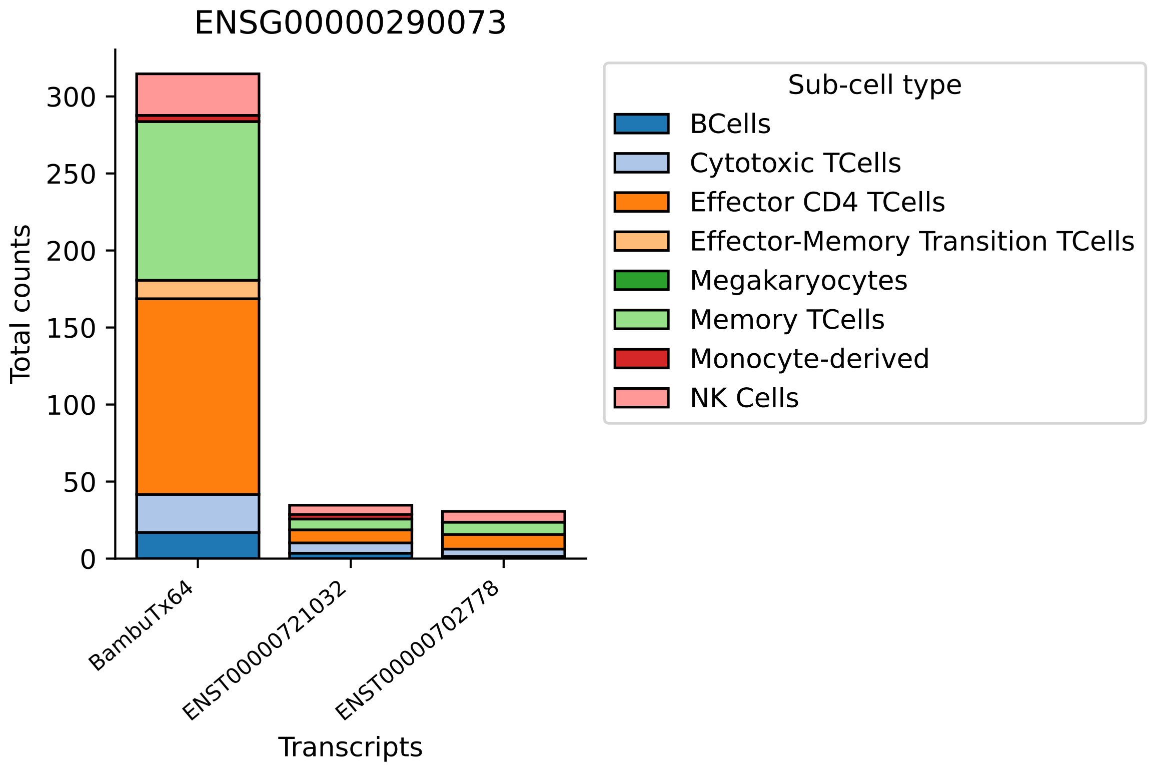
ENSG00000290073
Total counts
0
50
100
150
200
250
300
BambuTx64
ENST00000721032
ENST00000702778
Transcripts
Sub-cell type
BCells
Cytotoxic TCells
Effector CD4 TCells
Effector-Memory Transition TCells
Megakaryocytes
Memory TCells
Monocyte-derived
NK Cells

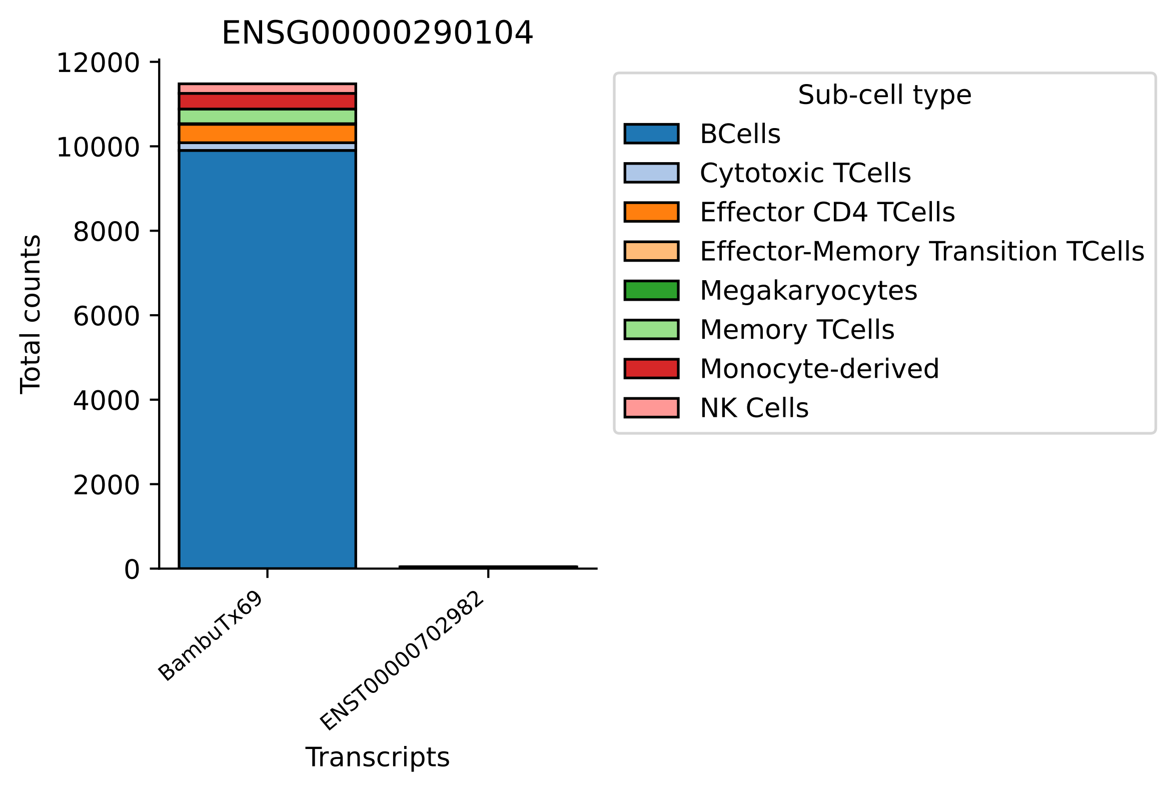
ENSG00000290104
Total counts
12000
10000
8000
6000
4000
2000
0
BambuTx69
ENST00000702982
Transcripts
Sub-cell type
BCells
Cytotoxic TCells
Effector CD4 TCells
Effector-Memory Transition TCells
Megakaryocytes
Memory TCells
Monocyte-derived
NK Cells

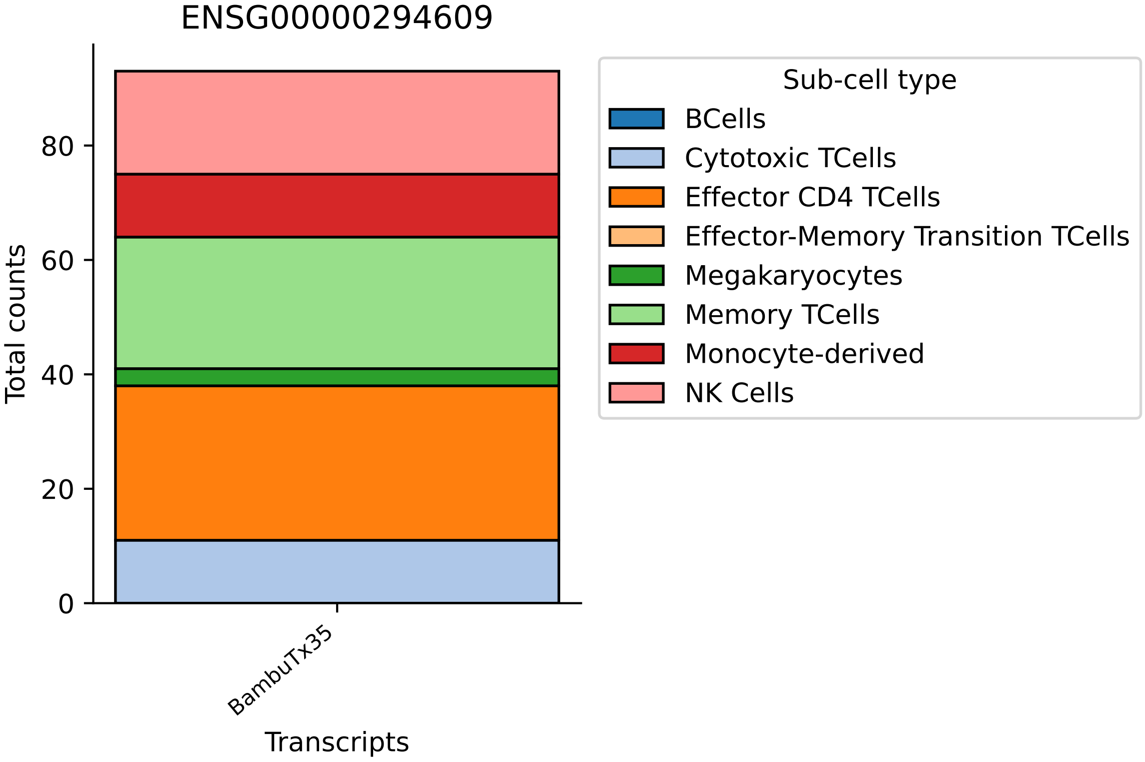
ENSG00000294609
Total counts
0
20
40
60
80
BambuTx35
Transcripts
Sub-cell type
BCells
Cytotoxic TCells
Effector CD4 TCells
Effector-Memory Transition TCells
Megakaryocytes
Memory TCells
Monocyte-derived
NK Cells

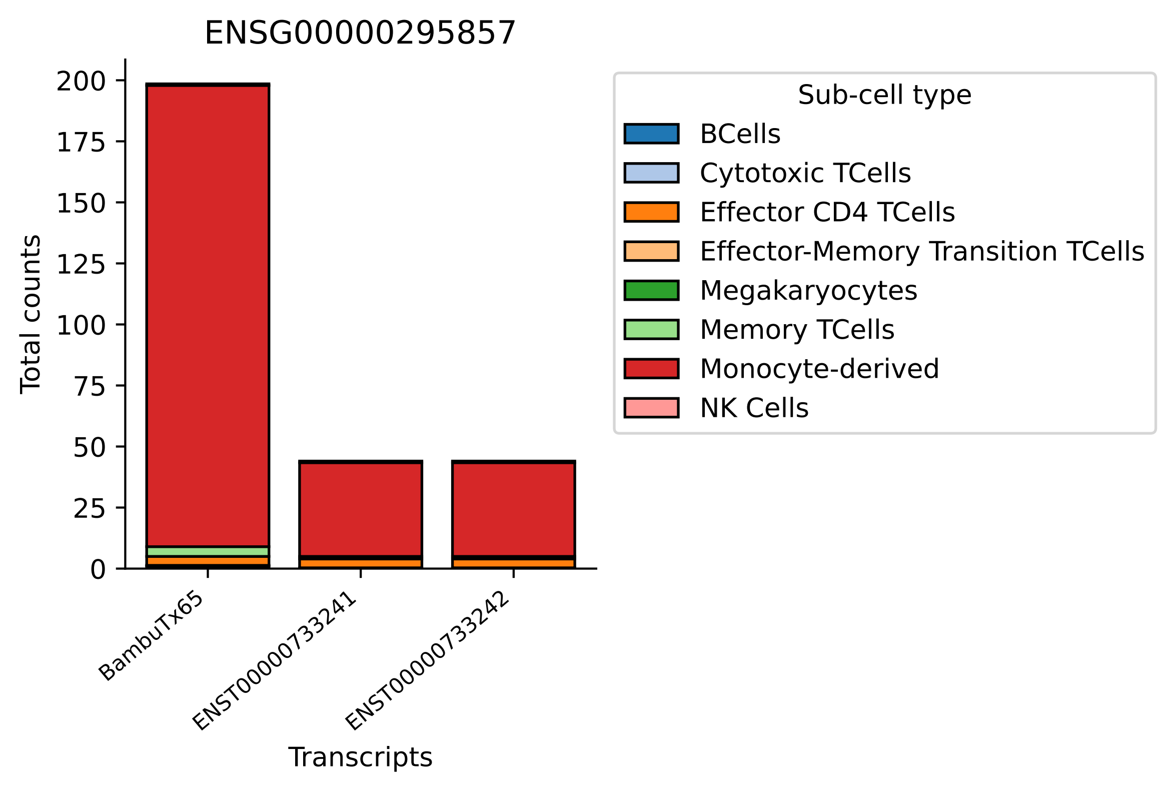
ENSG00000295857
Total counts
200
175
150
125
100
75
50
25
0
BambuTx65
ENST00000733241
ENST00000733242
Transcripts
Sub-cell type
BCells
Cytotoxic TCells
Effector CD4 TCells
Effector-Memory Transition TCells
Megakaryocytes
Memory TCells
Monocyte-derived
NK Cells

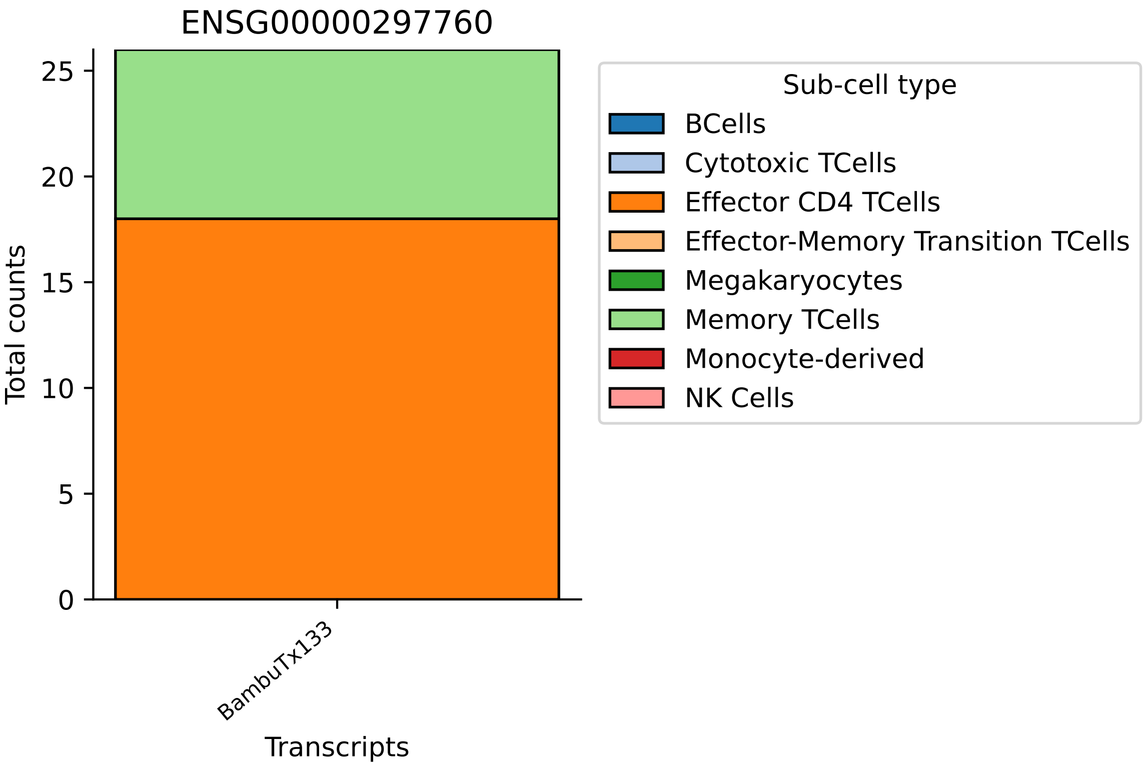

ENSG00000297760
Total counts
0
5
10
15
20
25
BambuTx133
Transcripts
Sub-cell type
BCells
Cytotoxic TCells
Effector CD4 TCells
Effector-Memory Transition TCells
Megakaryocytes
Memory TCells
Monocyte-derived
NK Cells

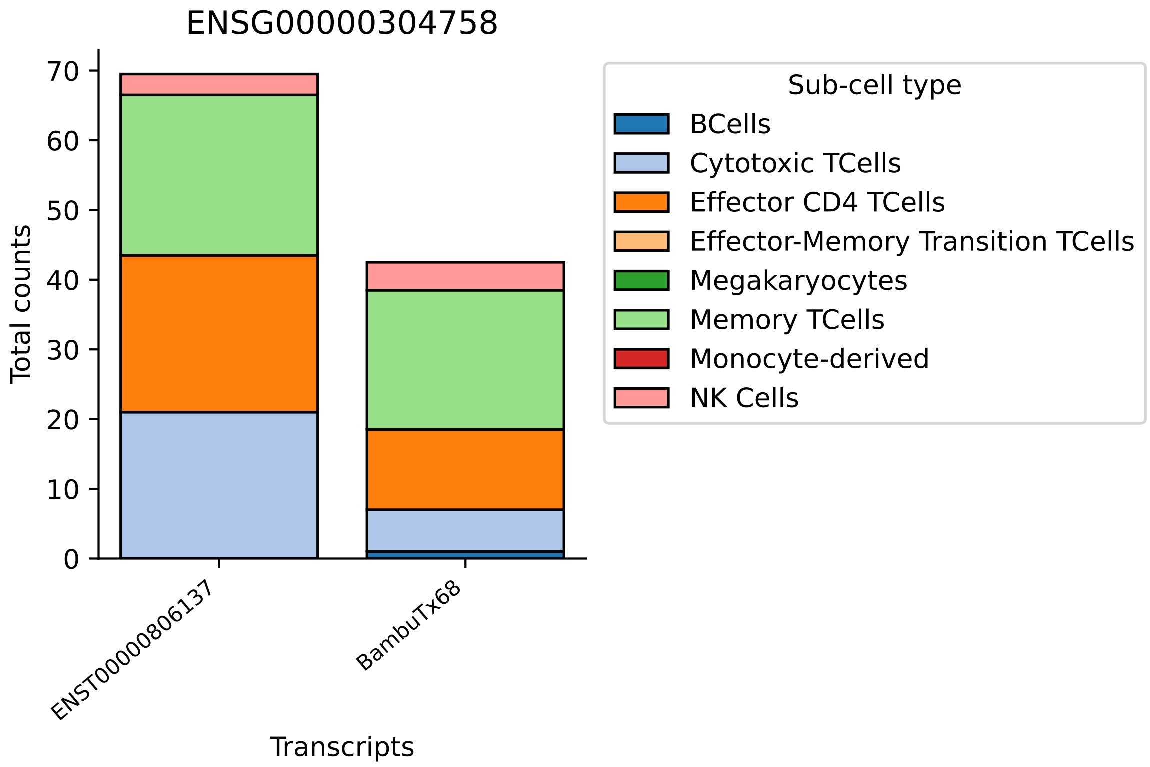

ENSG00000304758
Total counts
0
10
20
30
40
50
60
70
ENST00000806137
BambuTx68
Transcripts
Sub-cell type
BCells
Cytotoxic TCells
Effector CD4 TCells
Effector-Memory Transition TCells
Megakaryocytes
Memory TCells
Monocyte-derived
NK Cells

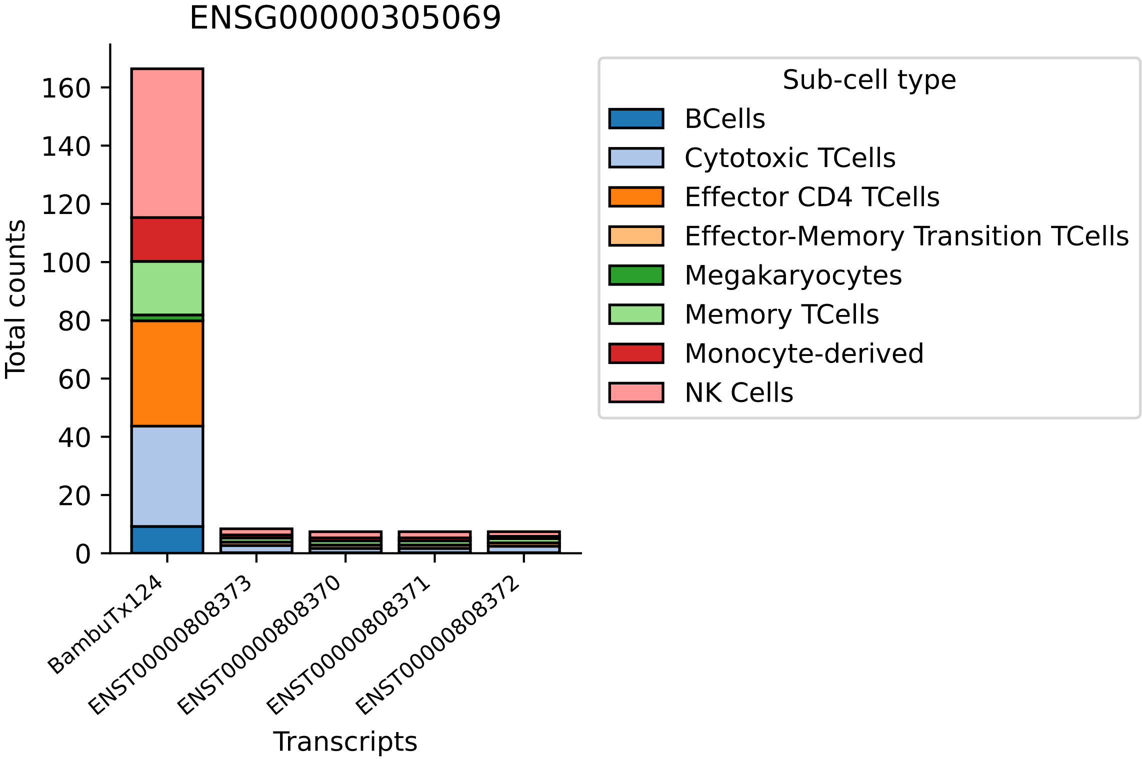

ENSG00000305069
Total counts
0
20
40
60
80
100
120
140
160
BambuTx124
ENST00000808373
ENST00000808370
ENST00000808371
ENST00000808372
Transcripts
Sub-cell type
BCells
Cytotoxic TCells
Effector CD4 TCells
Effector-Memory Transition TCells
Megakaryocytes
Memory TCells
Monocyte-derived
NK Cells

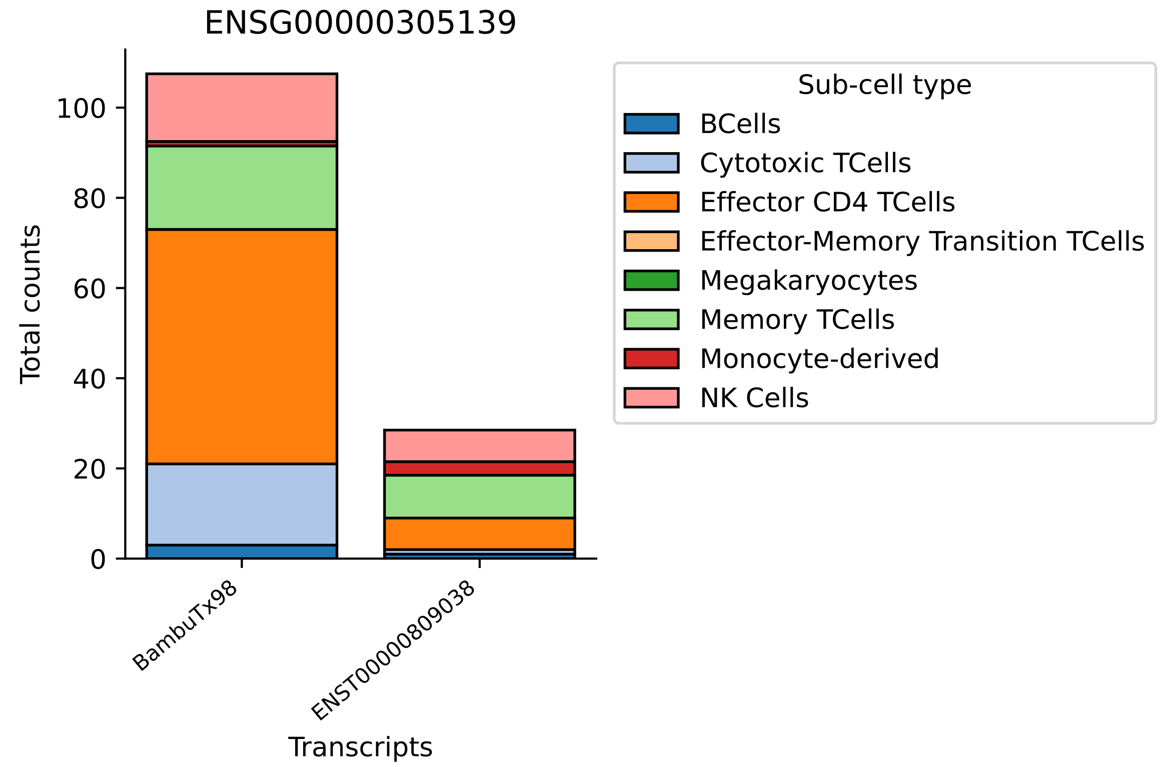
ENSG00000305139
Total counts
0
20
40
60
80
100
BambuTx98
ENST00000809038
Transcripts
Sub-cell type
BCells
Cytotoxic TCells
Effector CD4 TCells
Effector-Memory Transition TCells
Megakaryocytes
Memory TCells
Monocyte-derived
NK Cells

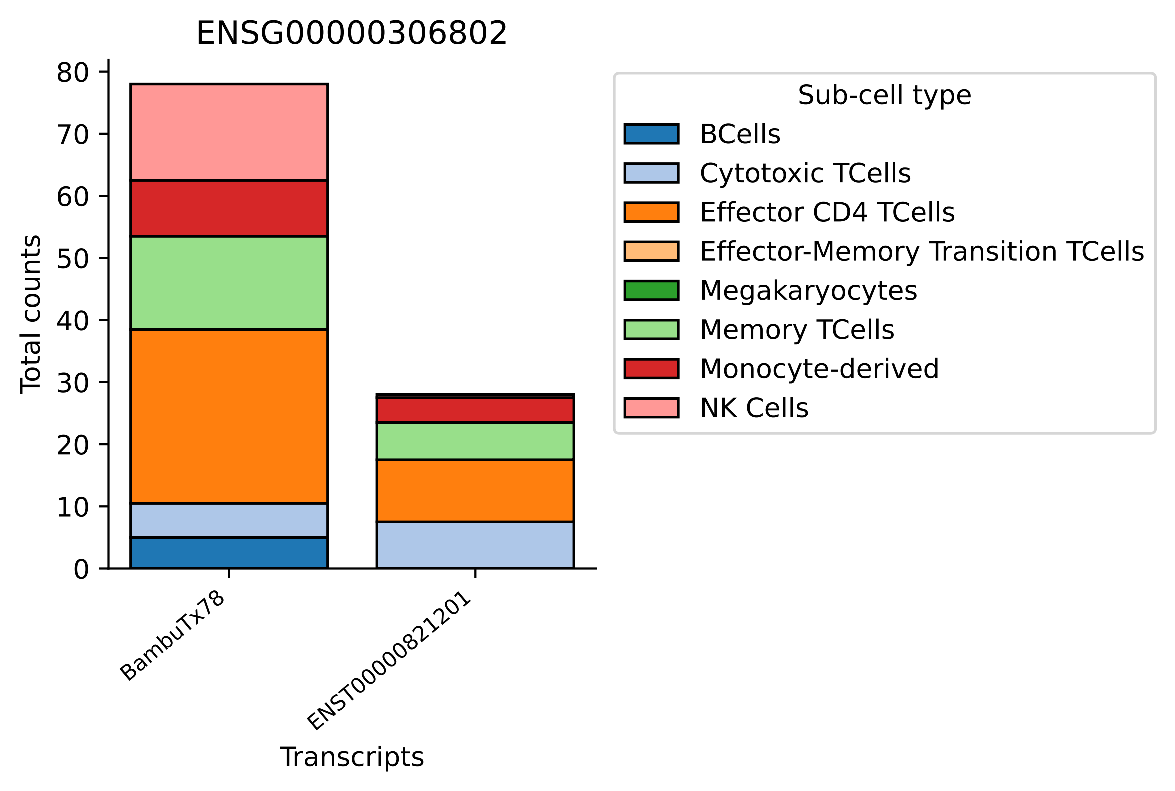
ENSG00000306802
Total counts
0
10
20
30
40
50
60
70
80
BambuTx78
ENST00000821201
Transcripts
Sub-cell type
BCells
Cytotoxic TCells
Effector CD4 TCells
Effector-Memory Transition TCells
Megakaryocytes
Memory TCells
Monocyte-derived
NK Cells

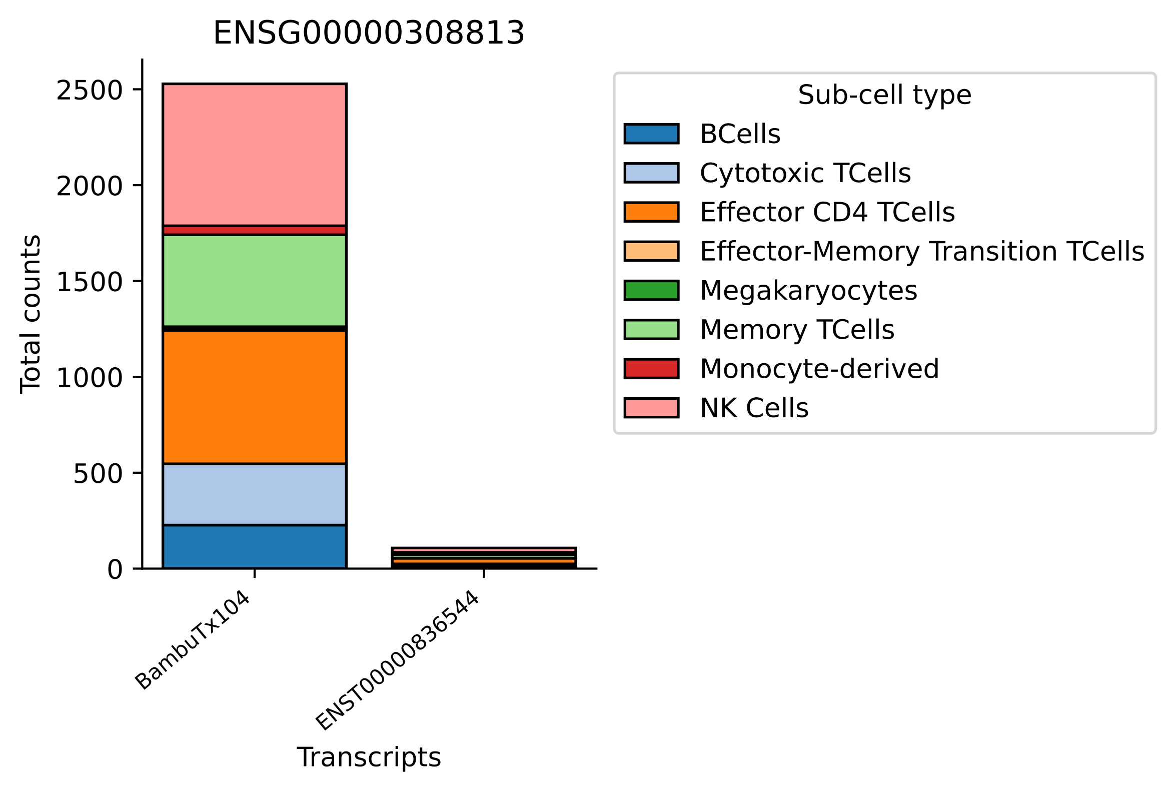

ENSG00000308813
Total counts
0
500
1000
1500
2000
2500
BambuTx104
ENST00000836544
Transcripts
Sub-cell type
BCells
Cytotoxic TCells
Effector CD4 TCells
Effector-Memory Transition TCells
Megakaryocytes
Memory TCells
Monocyte-derived
NK Cells

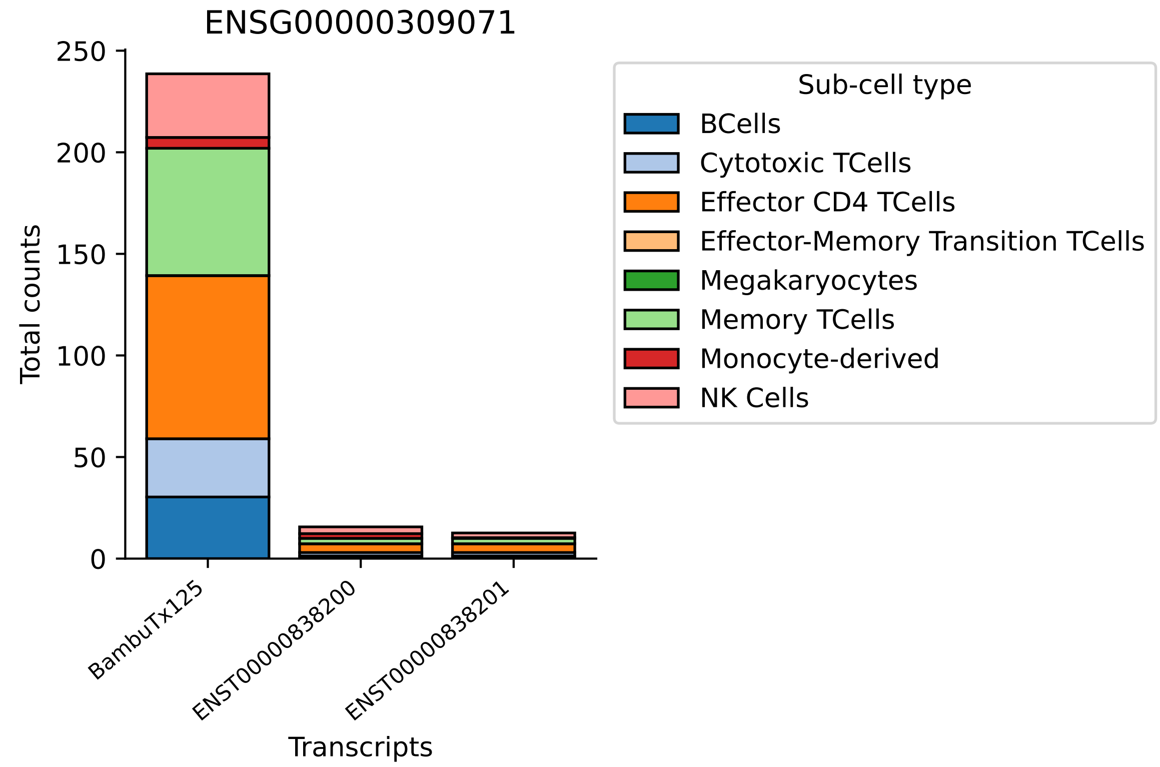
ENSG00000309071
Total counts
250
200
150
100
50
0
BambuTx125
ENST00000838200
ENST00000838201
Transcripts
Sub-cell type
BCells
Cytotoxic TCells
Effector CD4 TCells
Effector-Memory Transition TCells
Megakaryocytes
Memory TCells
Monocyte-derived
NK Cells

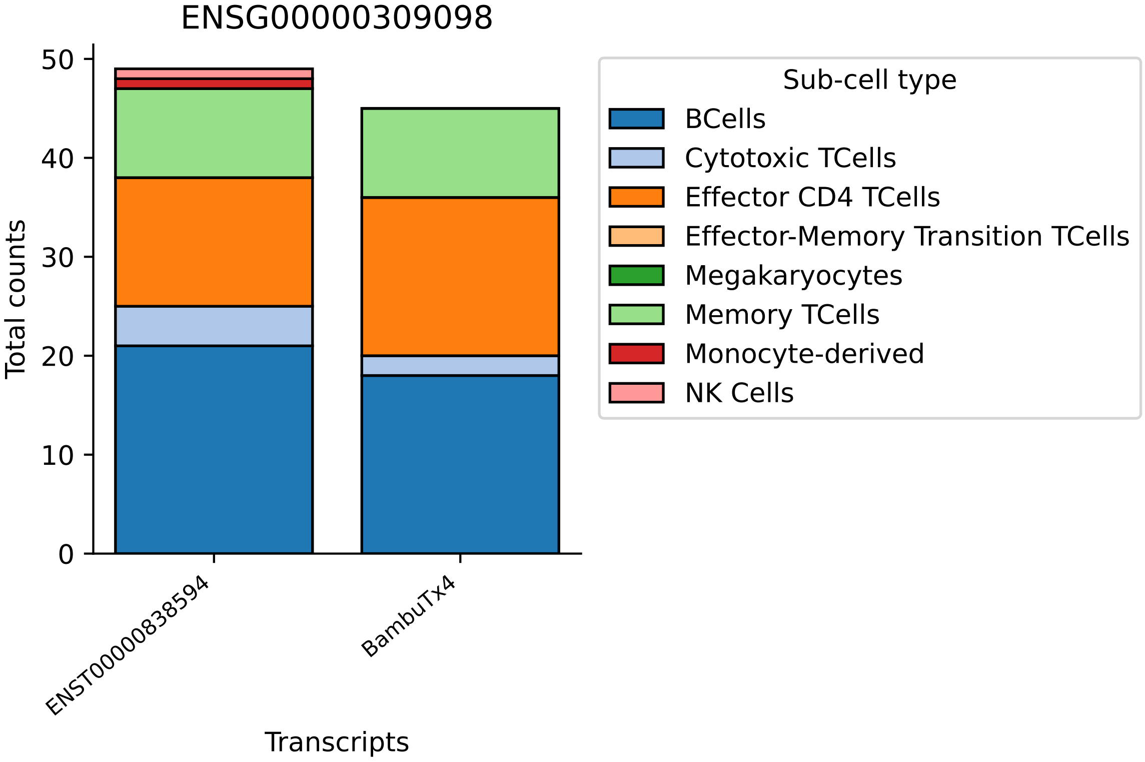
ENSG00000309098
Total counts
0
10
20
30
40
50
ENST00000838594
BambuTx4
Transcripts
Sub-cell type
BCells
Cytotoxic TCells
Effector CD4 TCells
Effector-Memory Transition TCells
Megakaryocytes
Memory TCells
Monocyte-derived
NK Cells

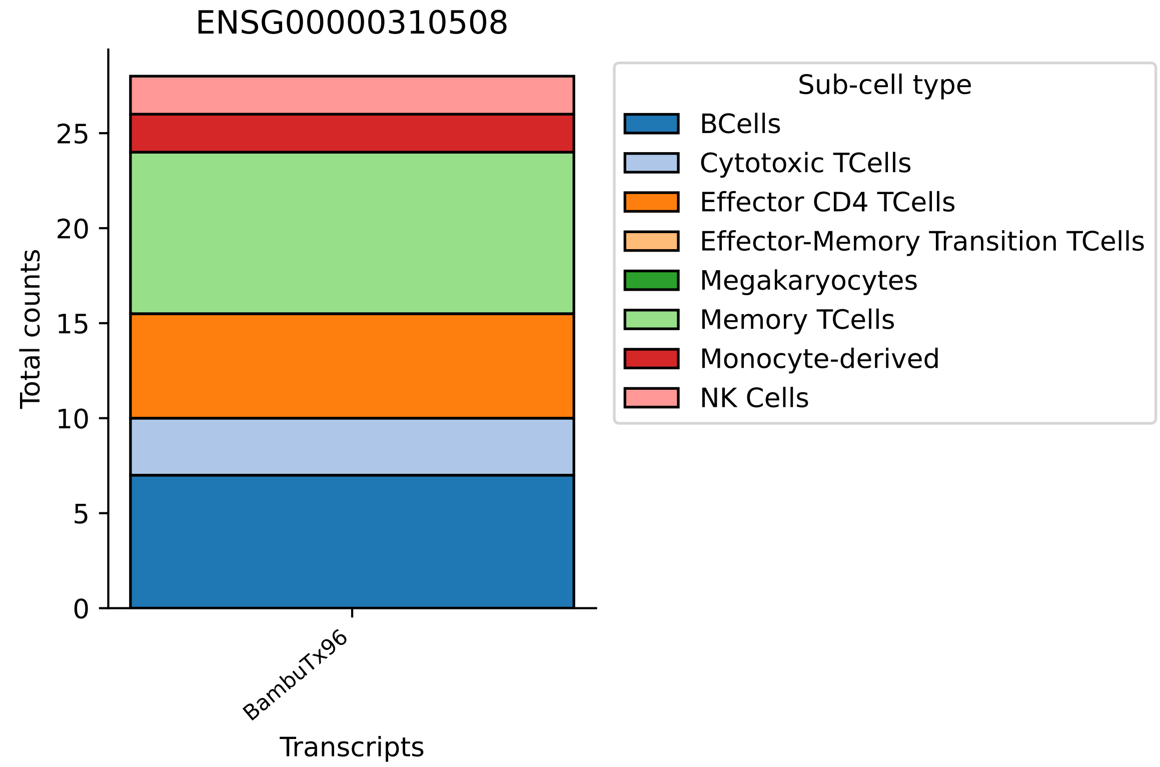
ENSG00000310508
Total counts
0
5
10
15
20
25
BambuTx96
Transcripts
Sub-cell type
BCells
Cytotoxic TCells
Effector CD4 TCells
Effector-Memory Transition TCells
Megakaryocytes
Memory TCells
Monocyte-derived
NK Cells

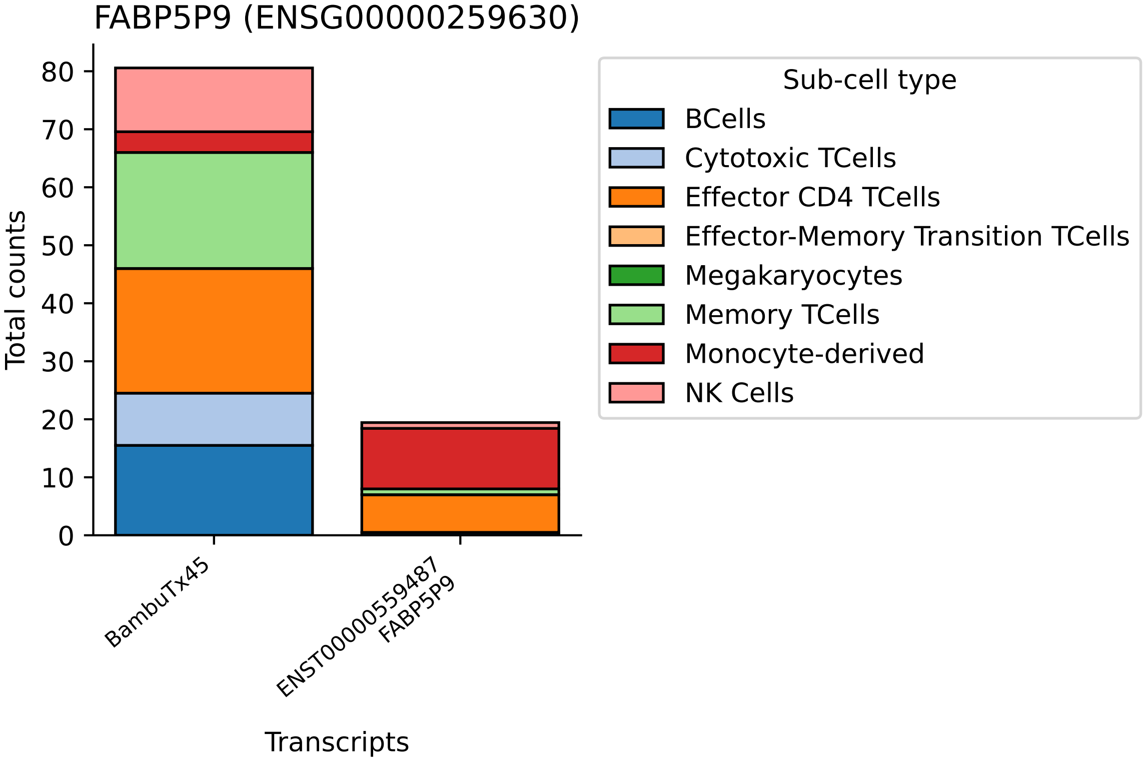

FABP5P9 (ENSG00000259630)
Total counts
0
10
20
30
40
50
60
70
80
BambuTx45
ENST00000559487
FABP5P9
Transcripts
Sub-cell type
BCells
Cytotoxic TCells
Effector CD4 TCells
Effector-Memory Transition TCells
Megakaryocytes
Memory TCells
Monocyte-derived
NK Cells

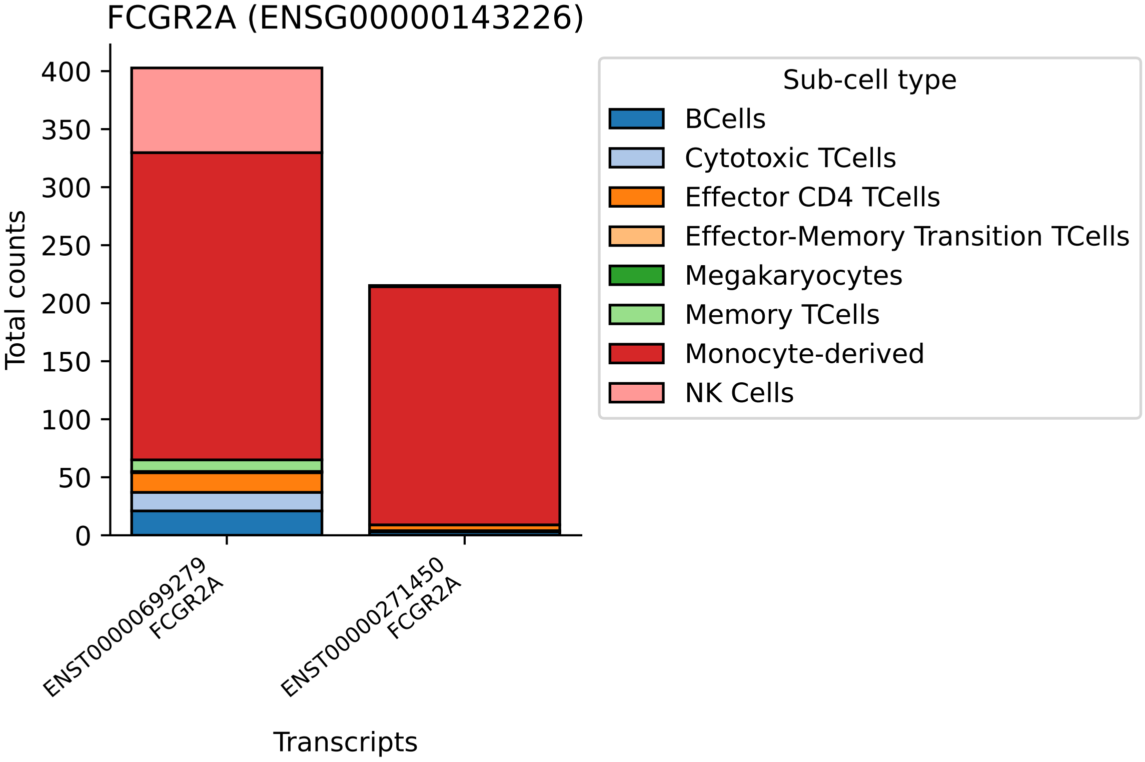

FCGR2A (ENSG00000143226)
Total counts
0
50
100
150
200
250
300
350
400
ENST00000699279
FCGR2A
ENST00000271450
FCGR2A
Transcripts
Sub-cell type
BCells
Cytotoxic TCells
Effector CD4 TCells
Effector-Memory Transition TCells
Megakaryocytes
Memory TCells
Monocyte-derived
NK Cells

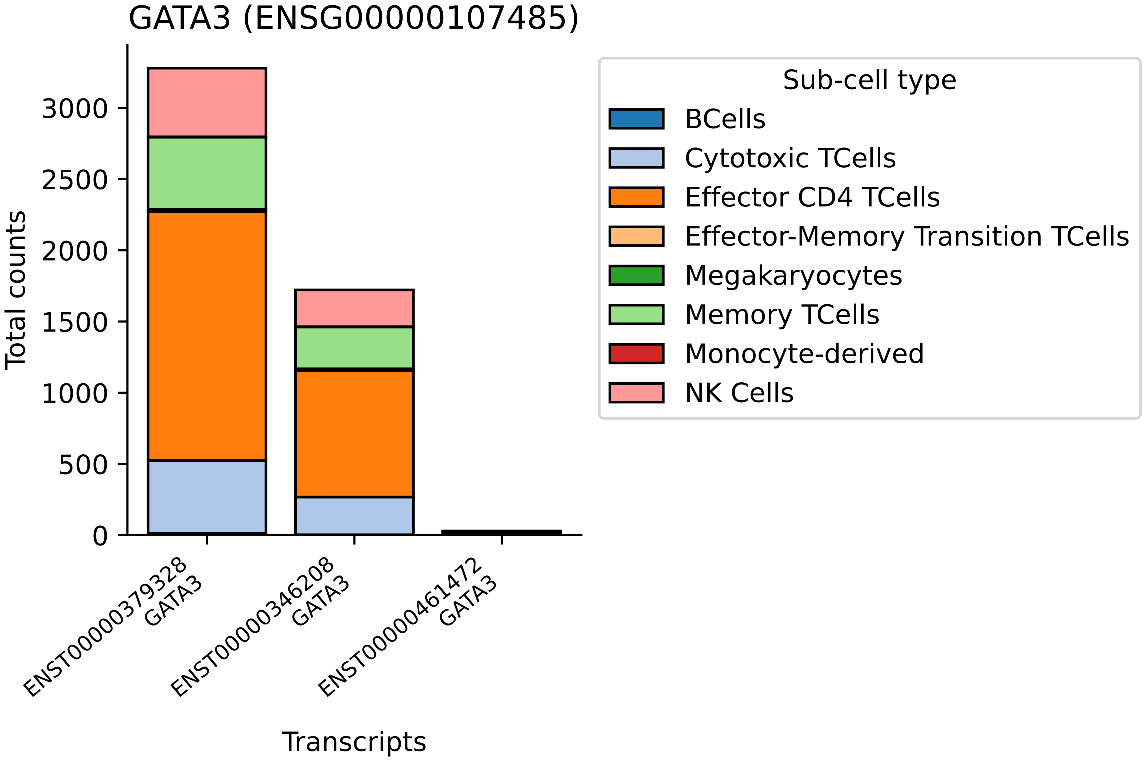
GATA3 (ENSG00000107485)
Total counts
3000
2500
2000
1500
1000
500
0
ENST00000379328
GATA3
ENST00000346208
GATA3
ENST00000461472
GATA3
Transcripts
Sub-cell type
BCells
Cytotoxic TCells
Effector CD4 TCells
Effector-Memory Transition TCells
Megakaryocytes
Memory TCells
Monocyte-derived
NK Cells

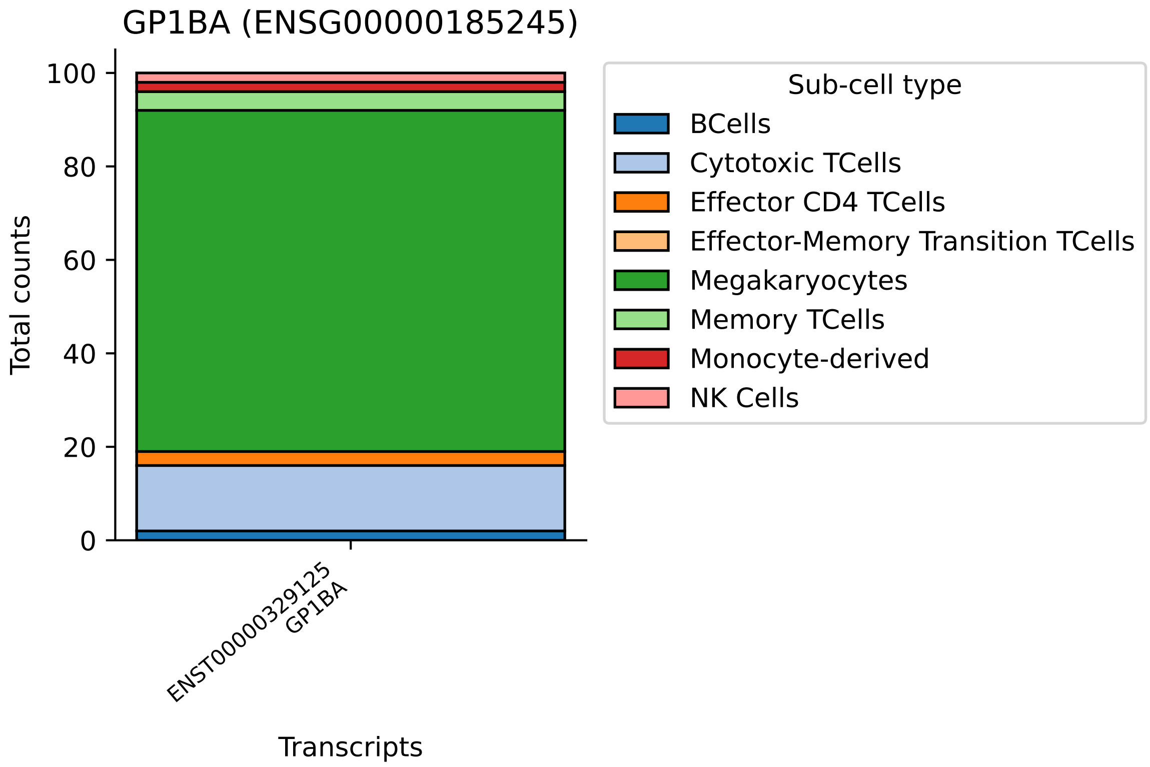
GP1BA (ENSG00000185245)
Total counts
100
80
60
40
20
0
ENST00000329125
GP1BA
Transcripts
Sub-cell type
BCells
Cytotoxic TCells
Effector CD4 TCells
Effector-Memory Transition TCells
Megakaryocytes
Memory TCells
Monocyte-derived
NK Cells

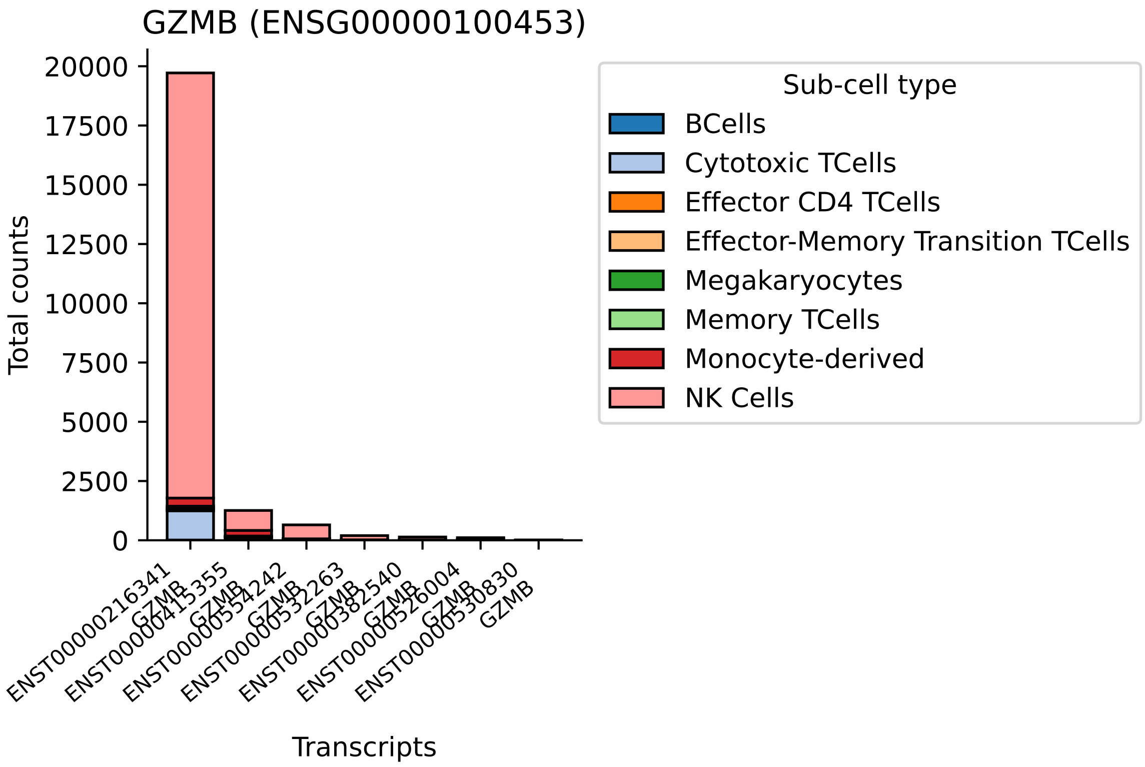

GZMB (ENSG00000100453)
Total counts
20000
17500
15000
12500
10000
7500
5000
2500
0
ENST00000216341 GZMB
ENST00000415355 GZMB
ENST00000554242 GZMB
ENST00000532263 GZMB
ENST00000382540 GZMB
ENST00000526004 GZMB
ENST00000530830 GZMB
Transcripts
Sub-cell type
BCells
Cytotoxic TCells
Effector CD4 TCells
Effector-Memory Transition TCells
Megakaryocytes
Memory TCells
Monocyte-derived
NK Cells

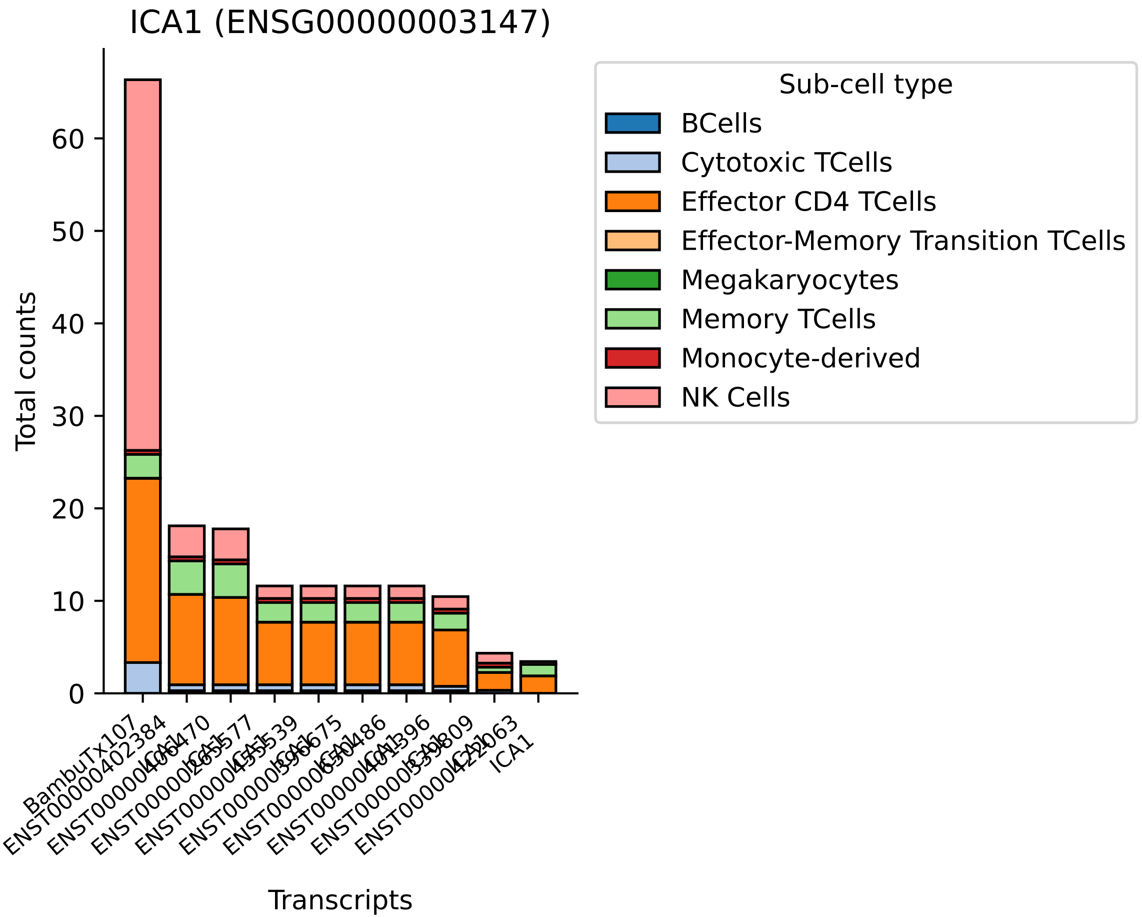

ICA1 (ENSG00000003147)
Total counts
0
10
20
30
40
50
60
Transcripts
BambuTx107
ICA1
Sub-cell type
BCells
Cytotoxic TCells
Effector CD4 TCells
Effector-Memory Transition TCells
Megakaryocytes
Memory TCells
Monocyte-derived
NK Cells

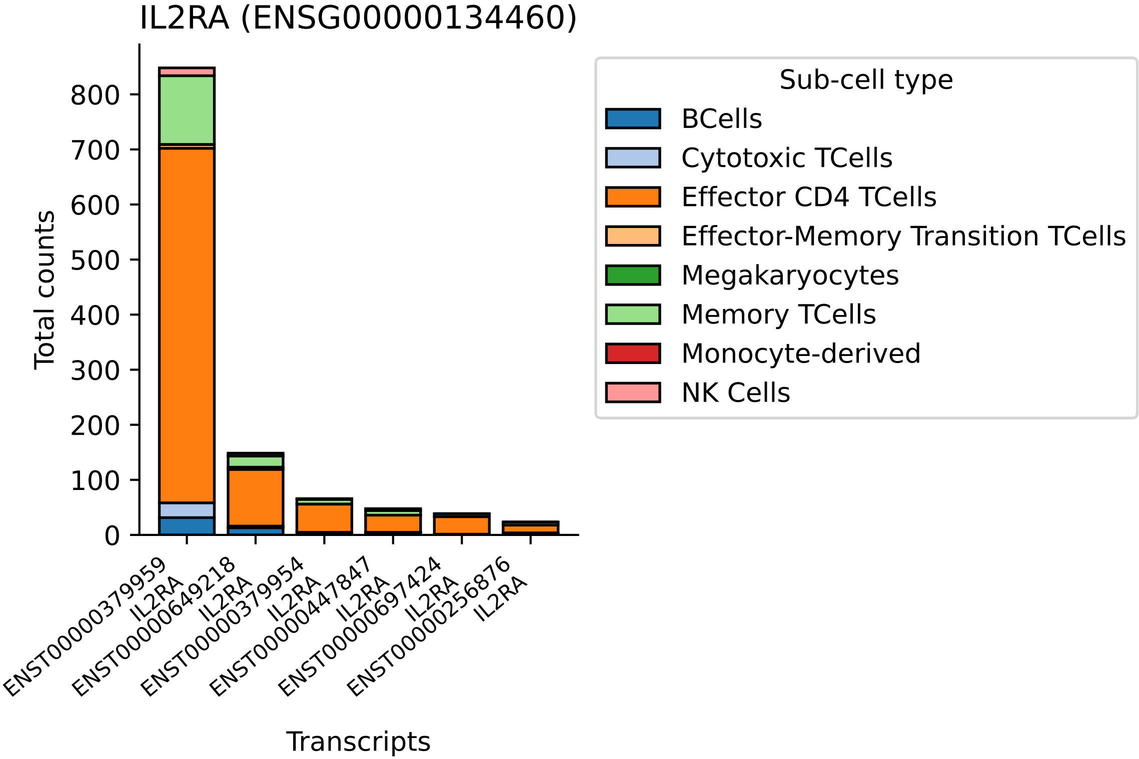

IL2RA (ENSG00000134460)
Total counts
0
100
200
300
400
500
600
700
800
ENST00000379959 IL2RA
ENST00000649218 IL2RA
ENST00000379954 IL2RA
ENST00000447847 IL2RA
ENST00000697424 IL2RA
ENST00000256876 IL2RA
Transcripts
Sub-cell type
BCells
Cytotoxic TCells
Effector CD4 TCells
Effector-Memory Transition TCells
Megakaryocytes
Memory TCells
Monocyte-derived
NK Cells

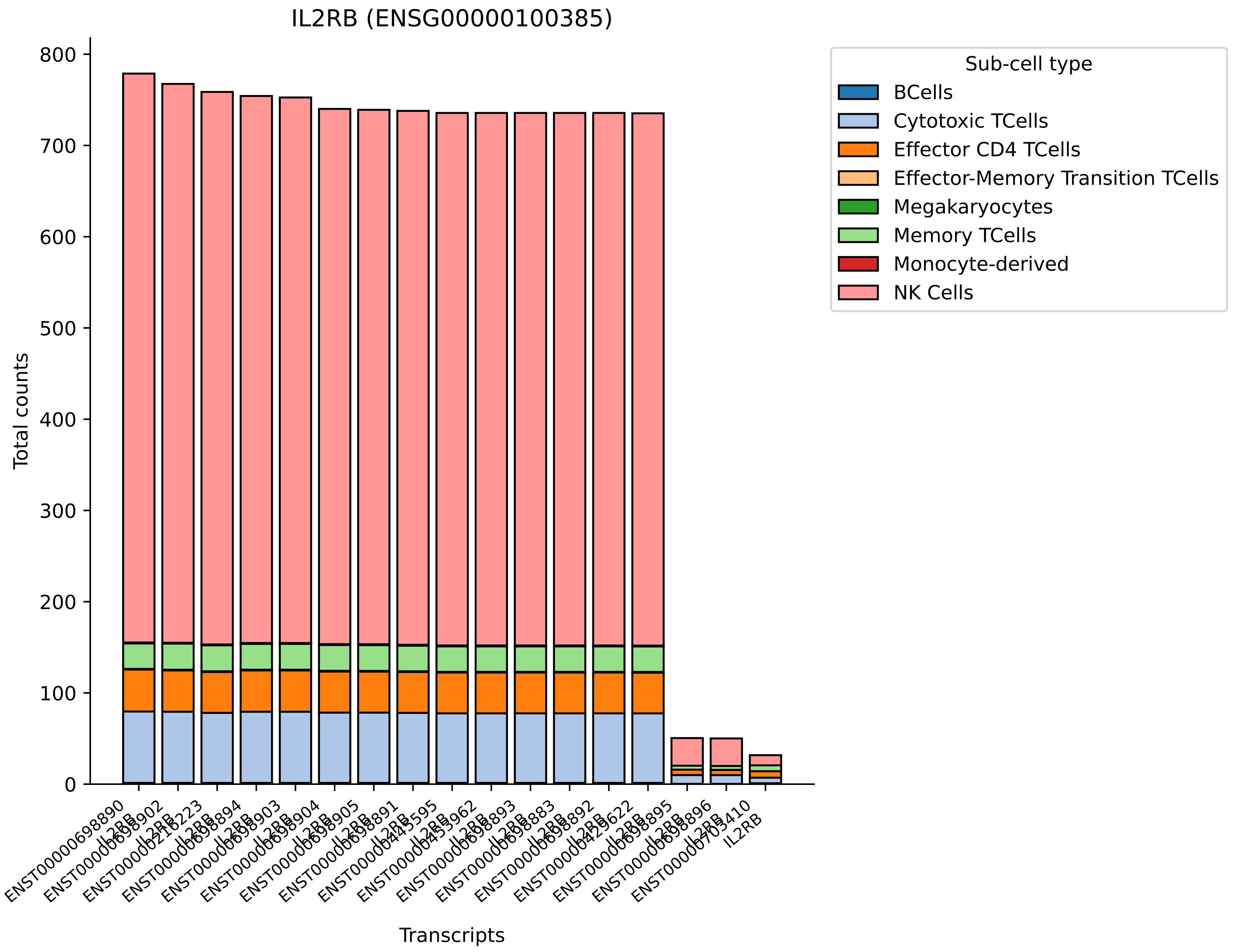

IL2RB (ENSG00000100385)
Total counts
0
100
200
300
400
500
600
700
800
Transcripts
ENST00000698890
IL2RB
Sub-cell type
BCells
Cytotoxic TCells
Effector CD4 TCells
Effector-Memory Transition TCells
Megakaryocytes
Memory TCells
Monocyte-derived
NK Cells

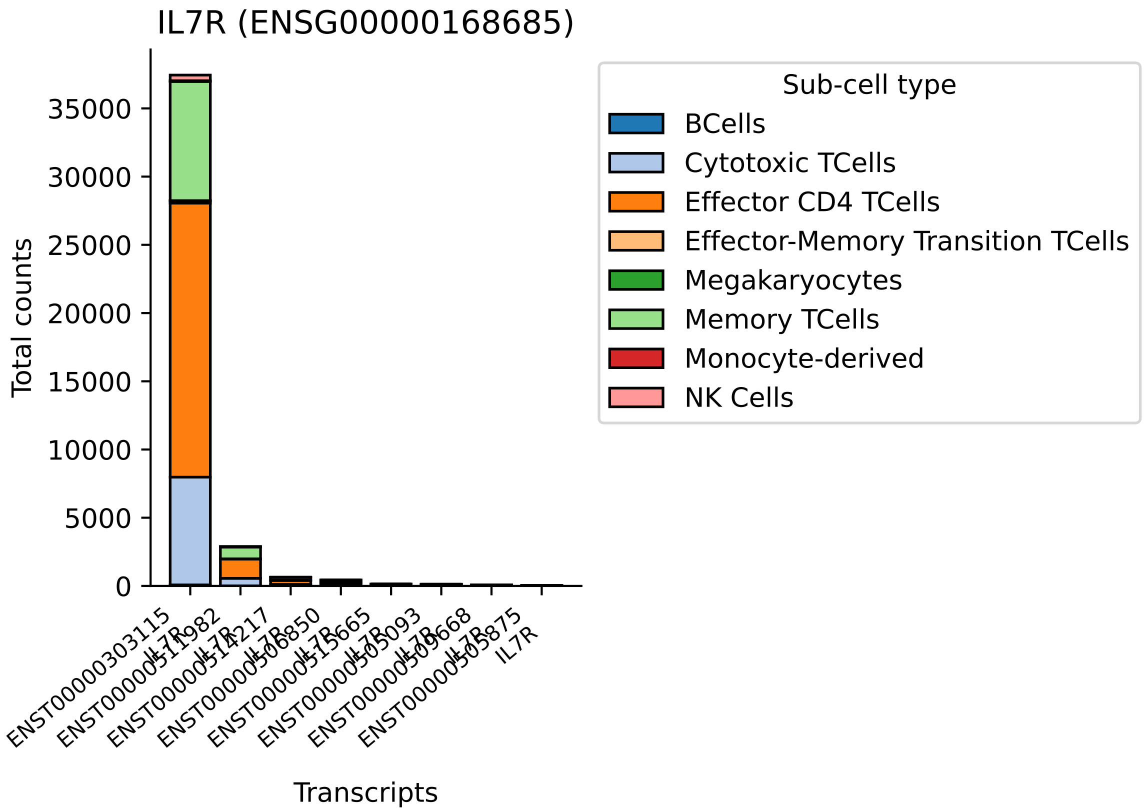
IL7R (ENSG00000168685)
Total counts
0
5000
10000
15000
20000
25000
30000
35000
Transcripts
ENST00000303115
IL7R
ENST00000511982
IL7R
ENST00000514217
IL7R
ENST00000506850
IL7R
ENST00000515665
IL7R
ENST00000505093
IL7R
ENST00000509668
IL7R
ENST00000505875
IL7R
Sub-cell type
BCells
Cytotoxic TCells
Effector CD4 TCells
Effector-Memory Transition TCells
Megakaryocytes
Memory TCells
Monocyte-derived
NK Cells

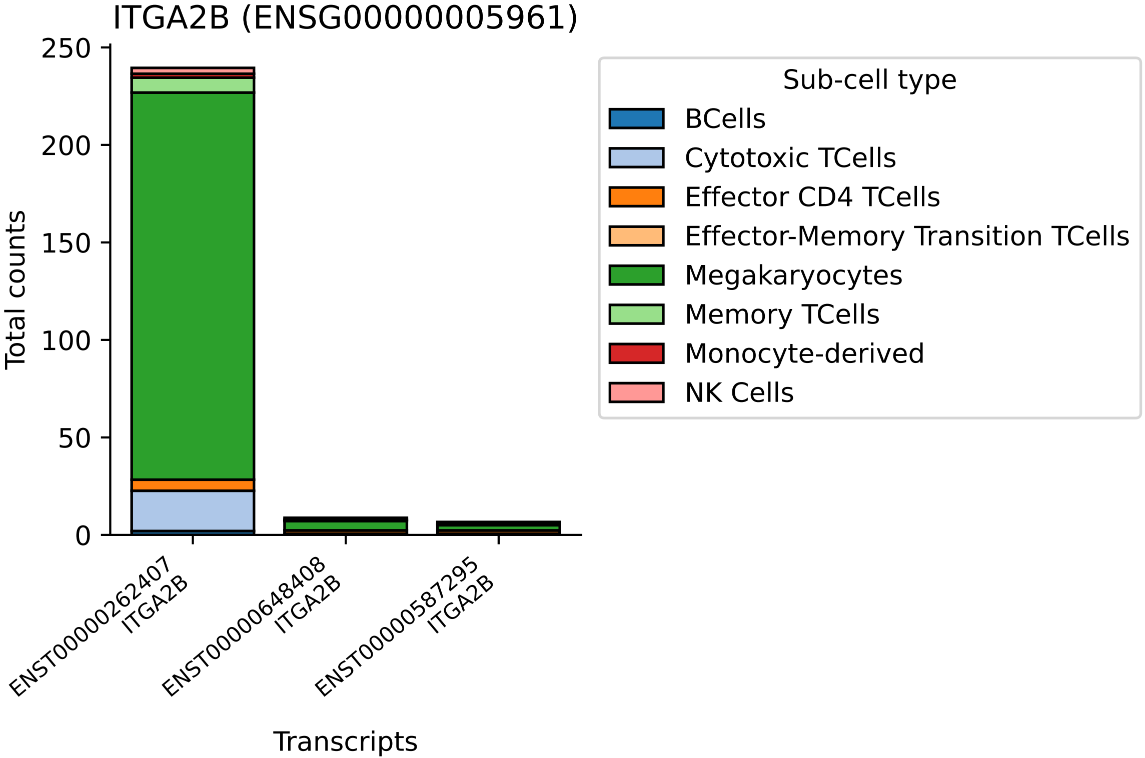

ITGA2B (ENSG00000005961)
Total counts
0
50
100
150
200
250
ENST00000262407
ITGA2B
ENST00000648408
ITGA2B
ENST00000587295
ITGA2B
Transcripts
Sub-cell type
BCells
Cytotoxic TCells
Effector CD4 TCells
Effector-Memory Transition TCells
Megakaryocytes
Memory TCells
Monocyte-derived
NK Cells

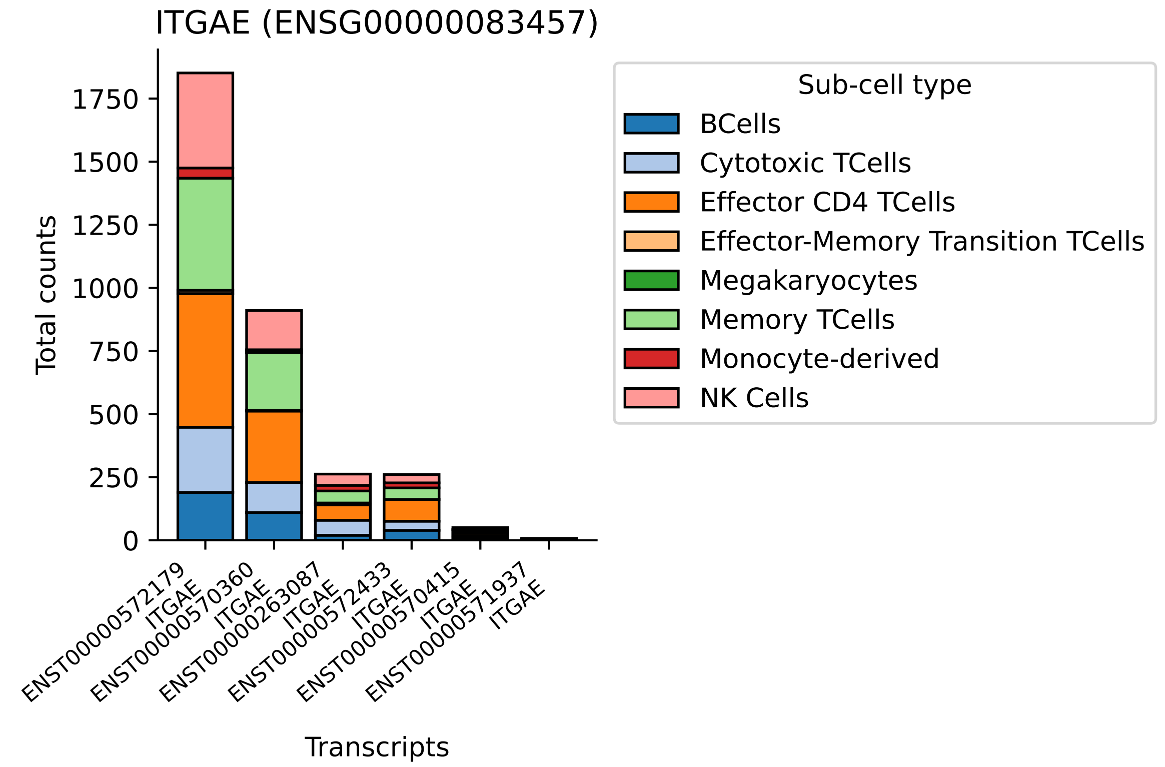

ITGAE (ENSG00000083457)
Total counts
0
250
500
750
1000
1250
1500
1750
Transcripts
ENST00000572179 ITGAE
ENST00000570360 ITGAE
ENST00000263087 ITGAE
ENST00000572433 ITGAE
ENST00000570415 ITGAE
ENST00000571937 ITGAE
Sub-cell type
BCells
Cytotoxic TCells
Effector CD4 TCells
Effector-Memory Transition TCells
Megakaryocytes
Memory TCells
Monocyte-derived
NK Cells

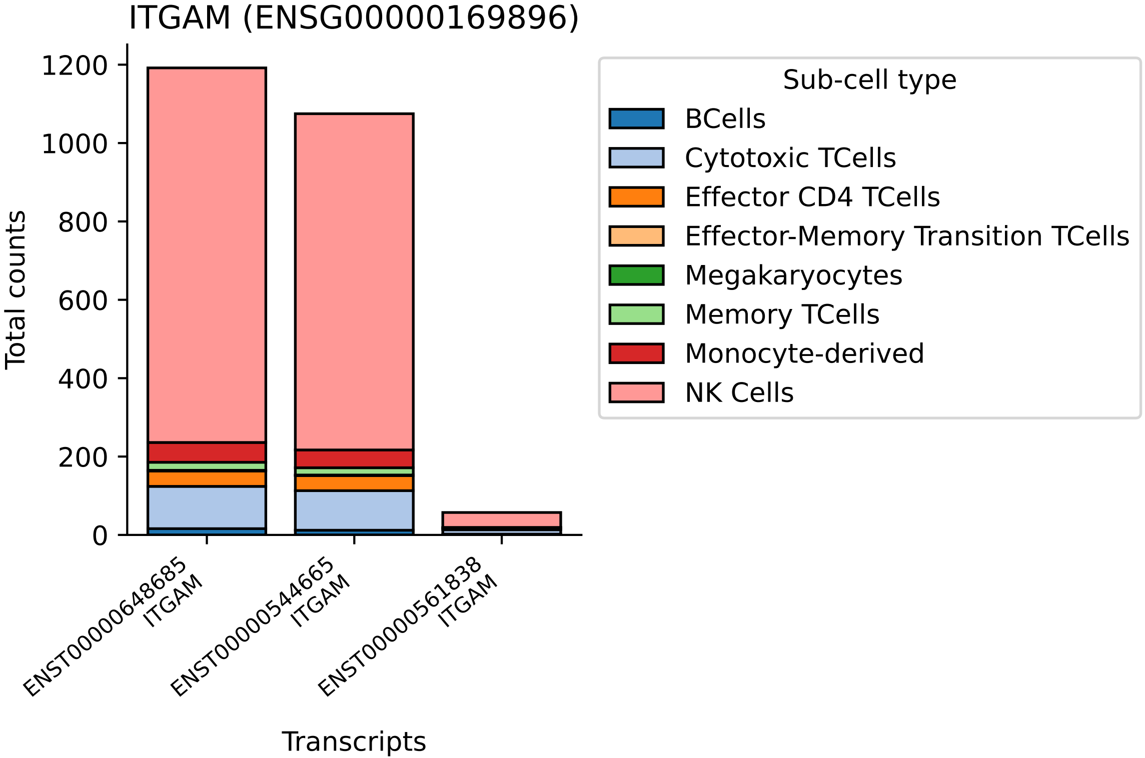

ITGAM (ENSG00000169896)
Total counts
1200
1000
800
600
400
200
0
ENST00000648685
ITGAM
ENST00000544665
ITGAM
ENST00000561838
ITGAM
Transcripts
Sub-cell type
BCells
Cytotoxic TCells
Effector CD4 TCells
Effector-Memory Transition TCells
Megakaryocytes
Memory TCells
Monocyte-derived
NK Cells

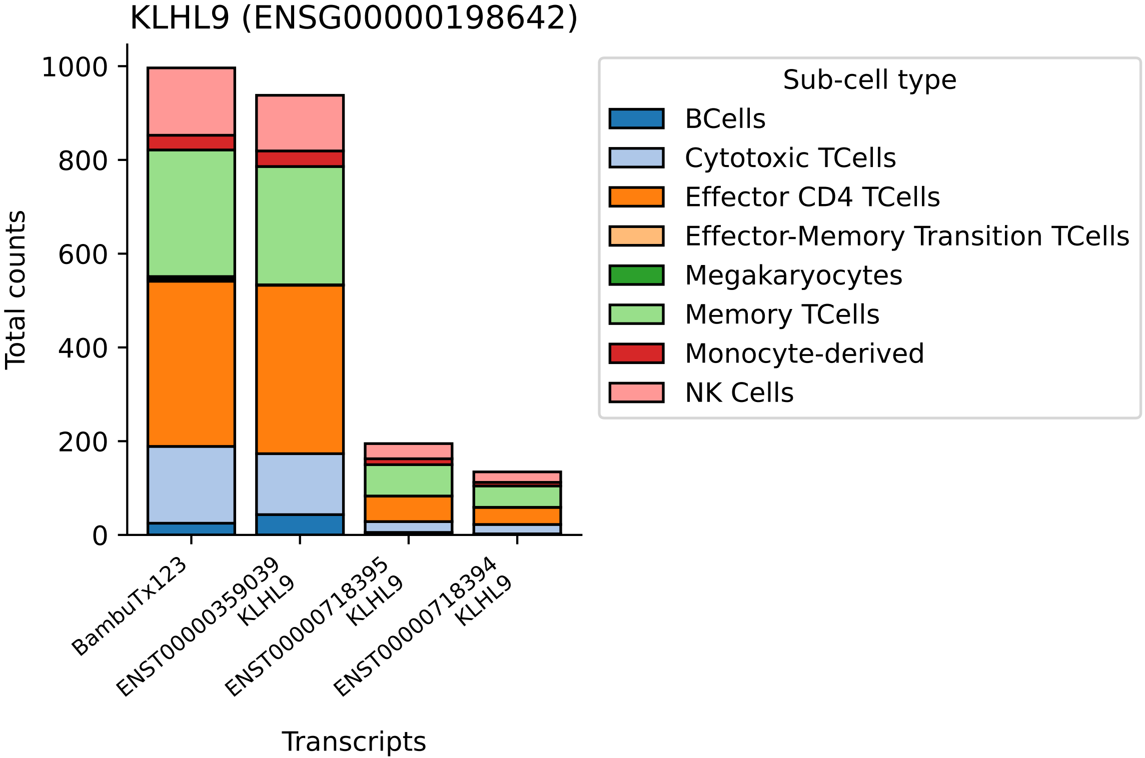

KLHL9 (ENSG00000198642)
Total counts
1000
800
600
400
200
0
Transcripts
BambuTx123
ENST00000359039 KLHL9
ENST00000718395 KLHL9
ENST00000718394 KLHL9
Sub-cell type
BCells
Cytotoxic TCells
Effector CD4 TCells
Effector-Memory Transition TCells
Megakaryocytes
Memory TCells
Monocyte-derived
NK Cells

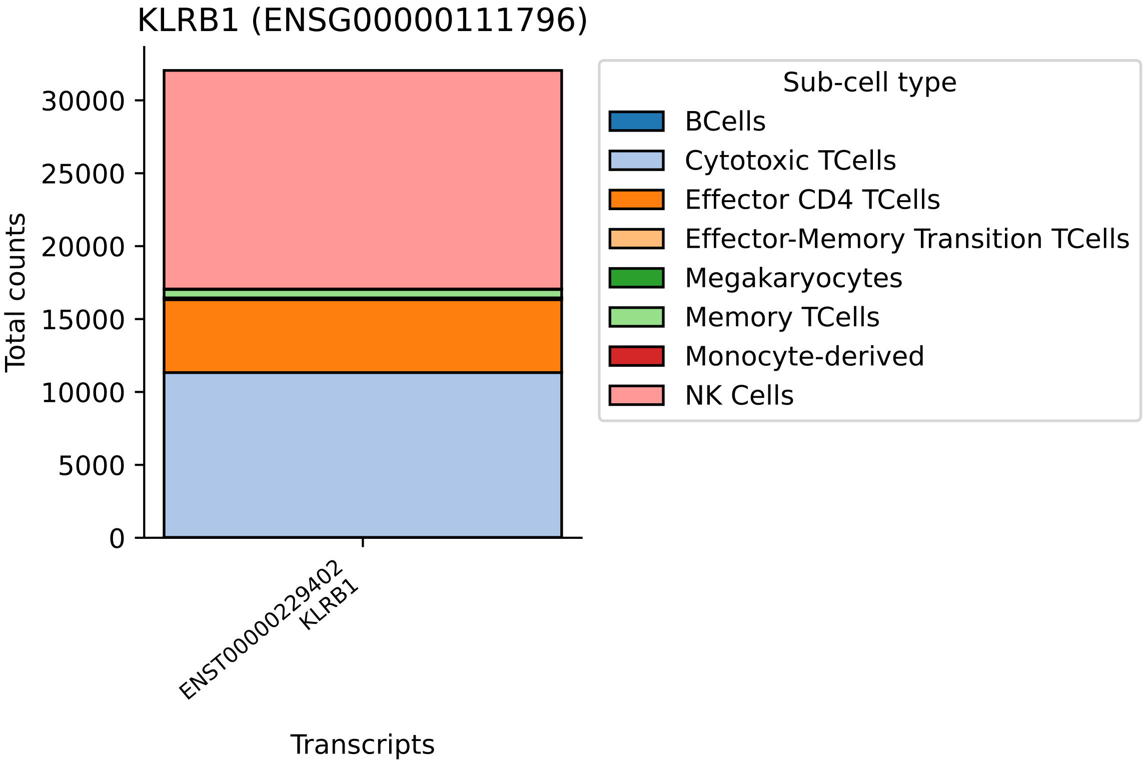

KLRB1 (ENSG00000111796)
Total counts
0
5000
10000
15000
20000
25000
30000
ENST00000229402
KLRB1
Transcripts
Sub-cell type
BCells
Cytotoxic TCells
Effector CD4 TCells
Effector-Memory Transition TCells
Megakaryocytes
Memory TCells
Monocyte-derived
NK Cells

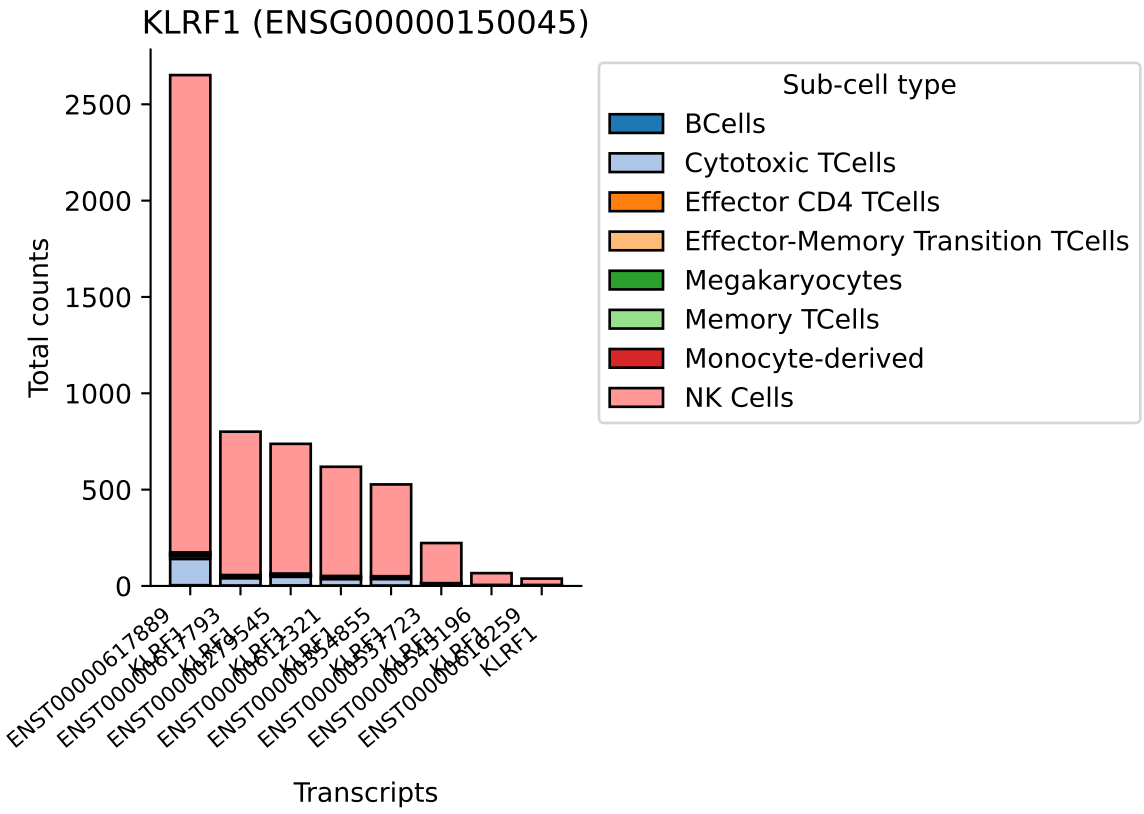
KLRF1 (ENSG00000150045)
Total counts
0
500
1000
1500
2000
2500
Transcripts
ENST00000617889
KLRF1
Sub-cell type
BCells
Cytotoxic TCells
Effector CD4 TCells
Effector-Memory Transition TCells
Megakaryocytes
Memory TCells
Monocyte-derived
NK Cells

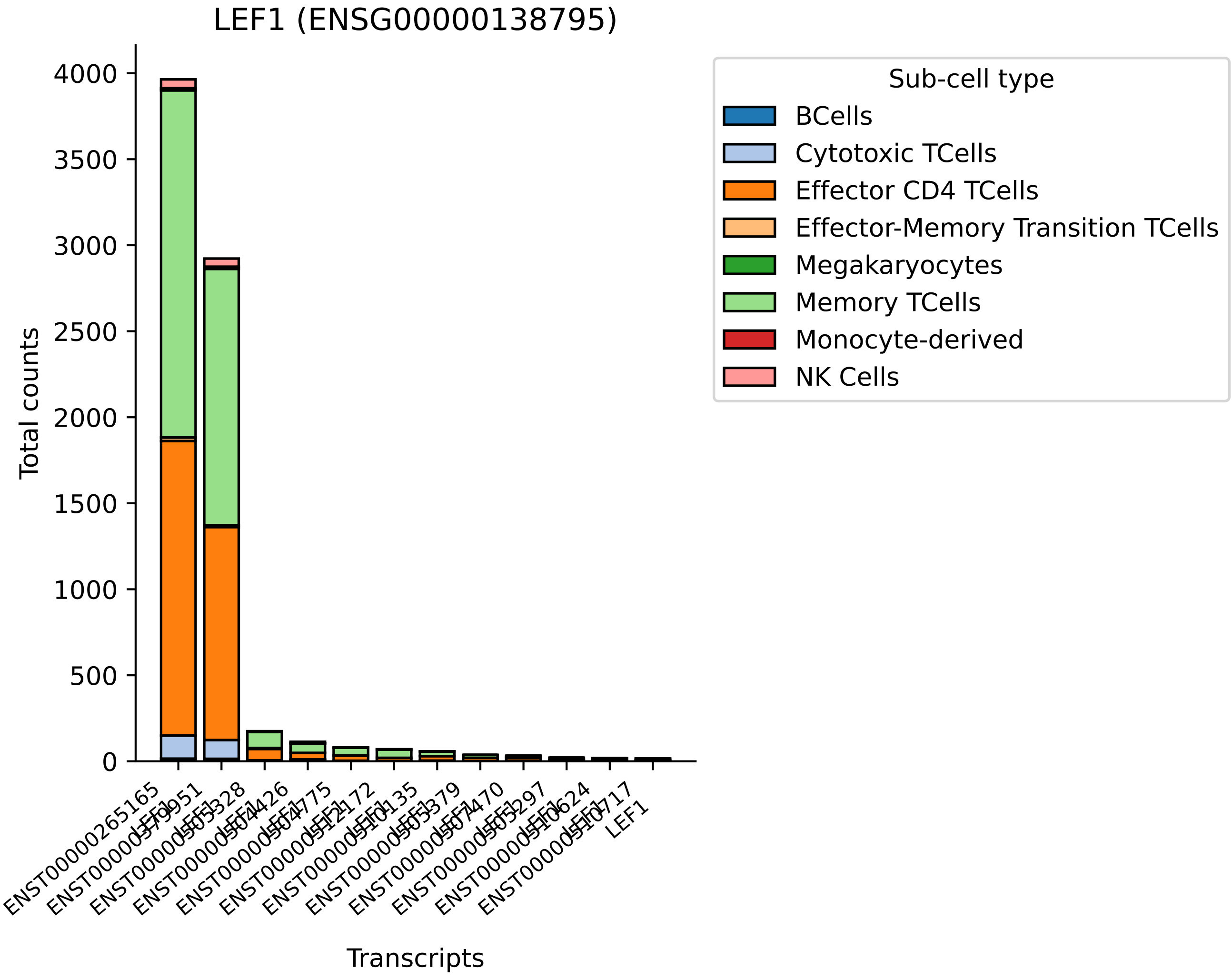
LEF1 (ENSG00000138795)
Total counts
0
500
1000
1500
2000
2500
3000
3500
4000
Transcripts
LEF1
Sub-cell type
BCells
Cytotoxic TCells
Effector CD4 TCells
Effector-Memory Transition TCells
Megakaryocytes
Memory TCells
Monocyte-derived
NK Cells

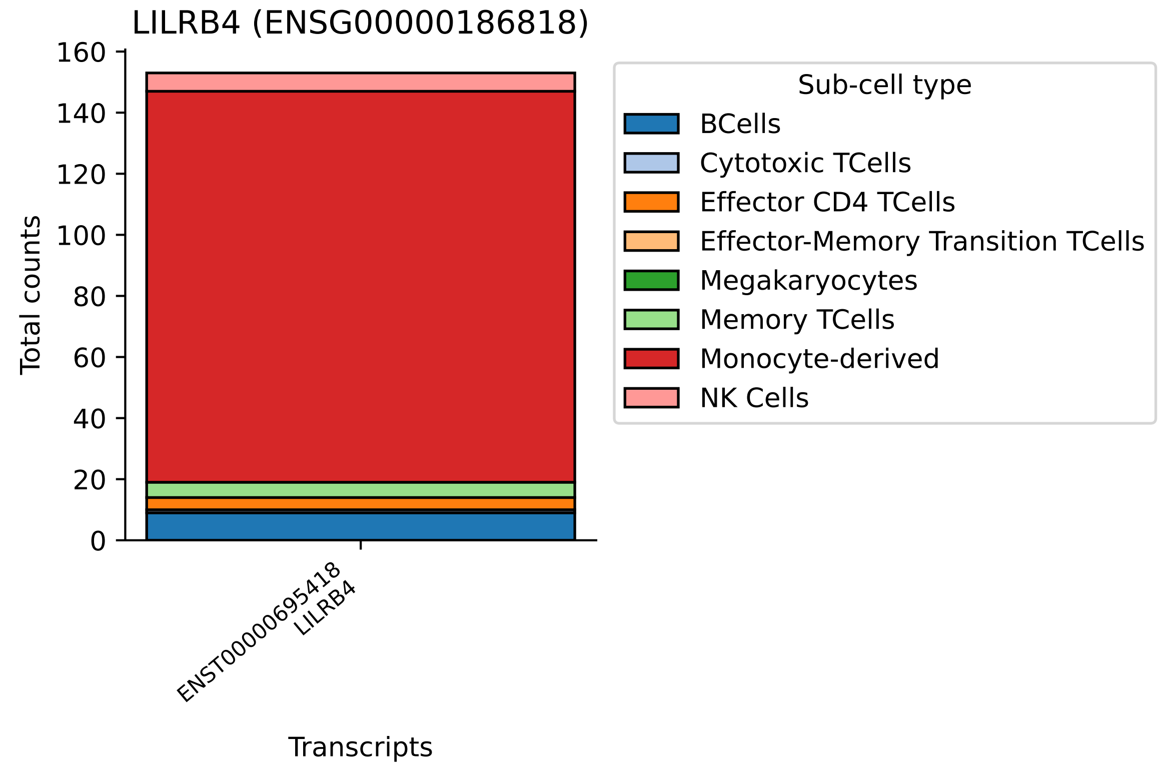
LILRB4 (ENSG00000186818)
Total counts
0
20
40
60
80
100
120
140
160
ENST00000695418
LILRB4
Transcripts
Sub-cell type
BCells
Cytotoxic TCells
Effector CD4 TCells
Effector-Memory Transition TCells
Megakaryocytes
Memory TCells
Monocyte-derived
NK Cells

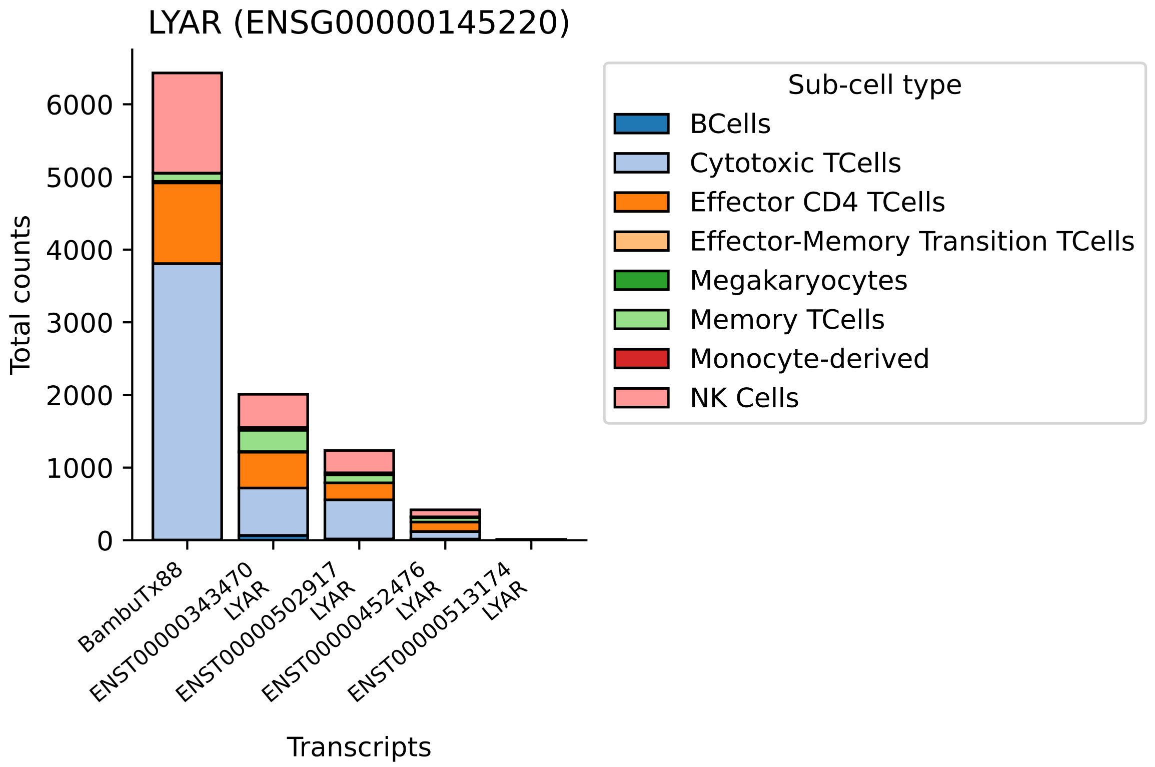
LYAR (ENSG00000145220)
Total counts
0
1000
2000
3000
4000
5000
6000
BambuTx88
ENST00000343470
LYAR
ENST00000502917
LYAR
ENST00000452476
LYAR
ENST00000513174
LYAR
Transcripts
Sub-cell type
BCells
Cytotoxic TCells
Effector CD4 TCells
Effector-Memory Transition TCells
Megakaryocytes
Memory TCells
Monocyte-derived
NK Cells

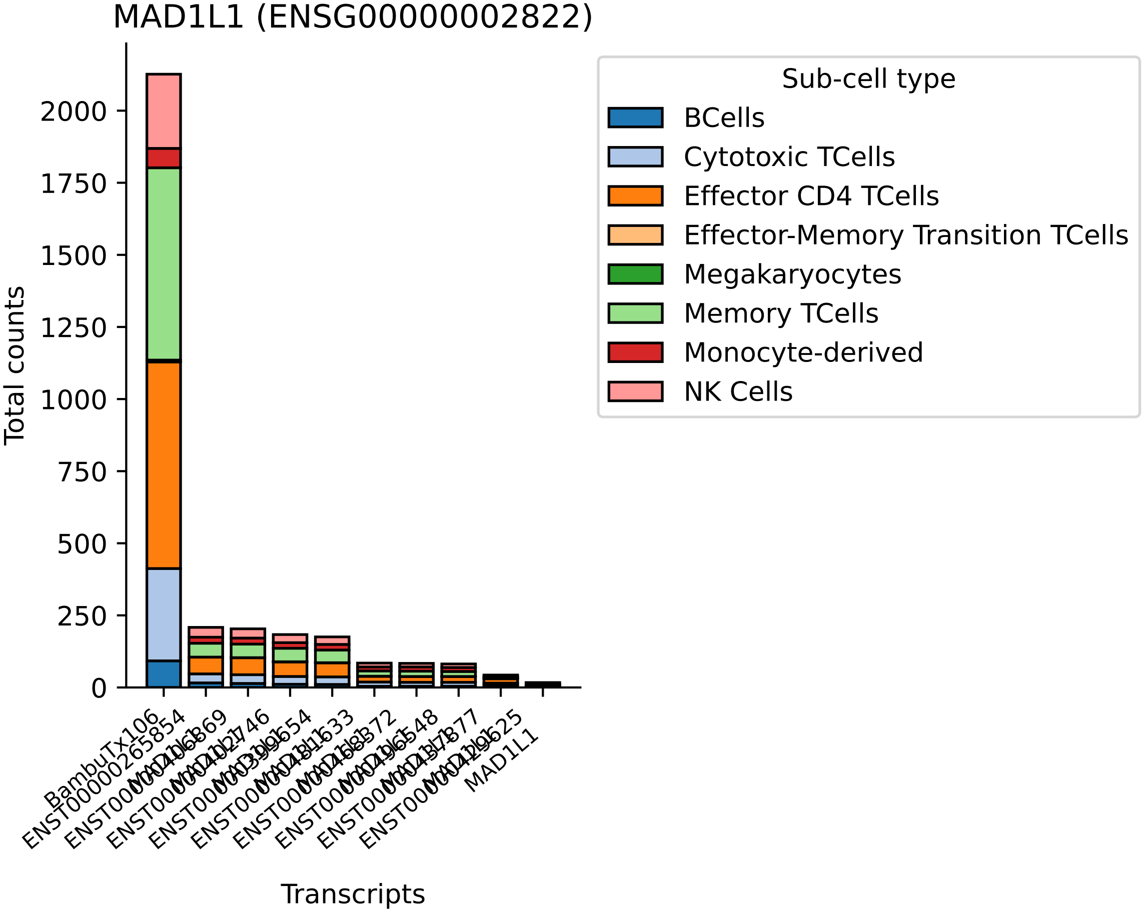

MAD1L1 (ENSG00000002822)
Total counts
0
250
500
750
1000
1250
1500
1750
2000
Transcripts
BambuTx106
MAD1L1
Sub-cell type
BCells
Cytotoxic TCells
Effector CD4 TCells
Effector-Memory Transition TCells
Megakaryocytes
Memory TCells
Monocyte-derived
NK Cells

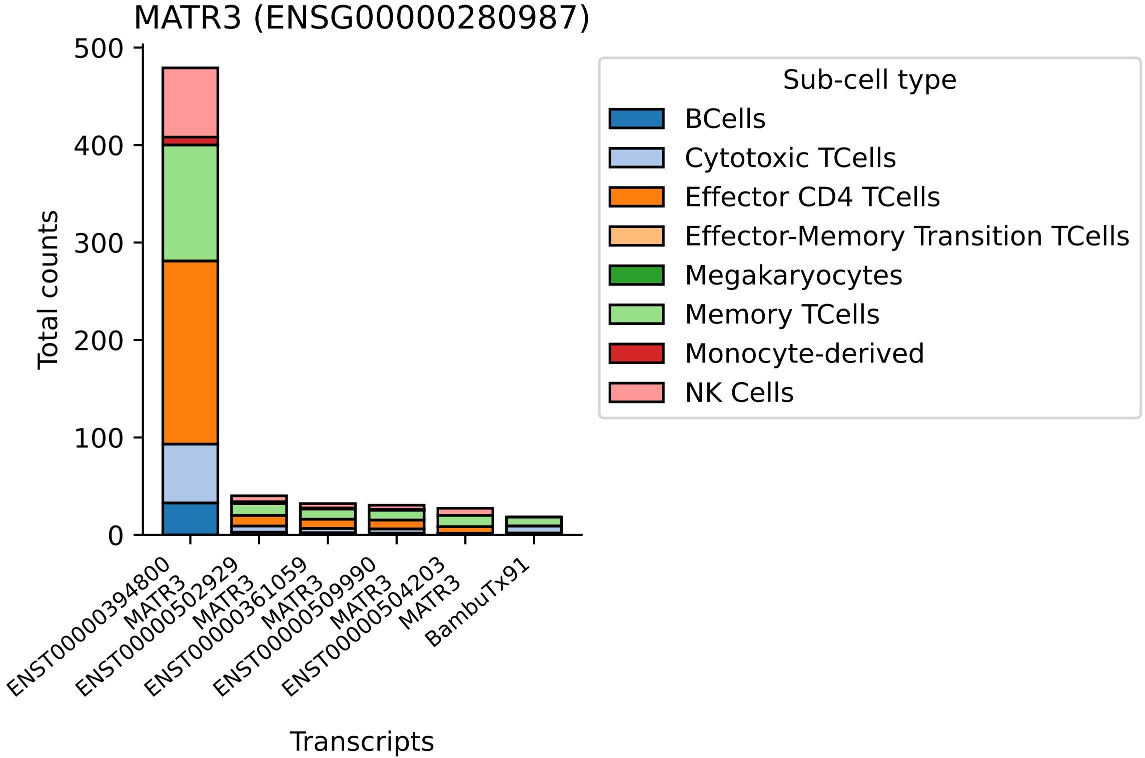

MATR3 (ENSG00000280987)
Total counts
0
100
200
300
400
500
ENST00000394800
MATR3
ENST00000502929
MATR3
ENST00000361059
MATR3
ENST00000509990
MATR3
ENST00000504203
MATR3
BambuTx91
Transcripts
Sub-cell type
BCells
Cytotoxic TCells
Effector CD4 TCells
Effector-Memory Transition TCells
Megakaryocytes
Memory TCells
Monocyte-derived
NK Cells

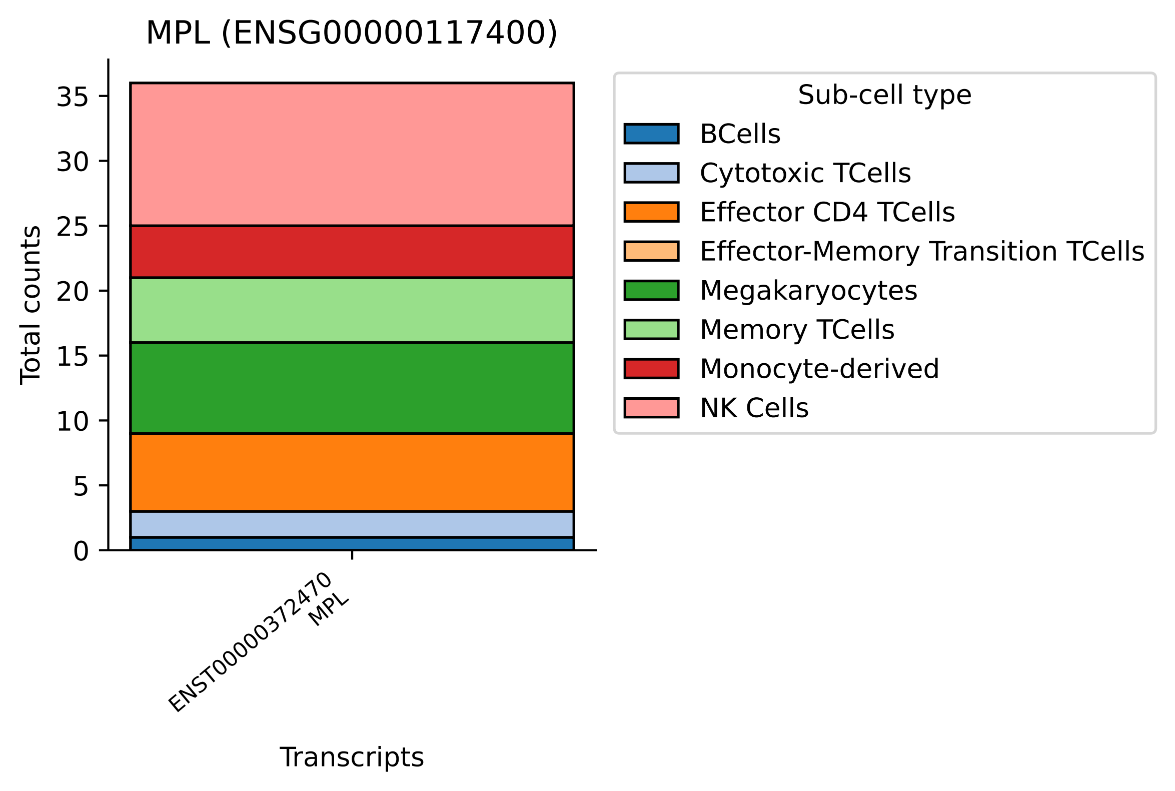
MPL (ENSG00000117400)
Total counts
0
5
10
15
20
25
30
35
ENST00000372470
MPL
Transcripts
Sub-cell type
BCells
Cytotoxic TCells
Effector CD4 TCells
Effector-Memory Transition TCells
Megakaryocytes
Memory TCells
Monocyte-derived
NK Cells

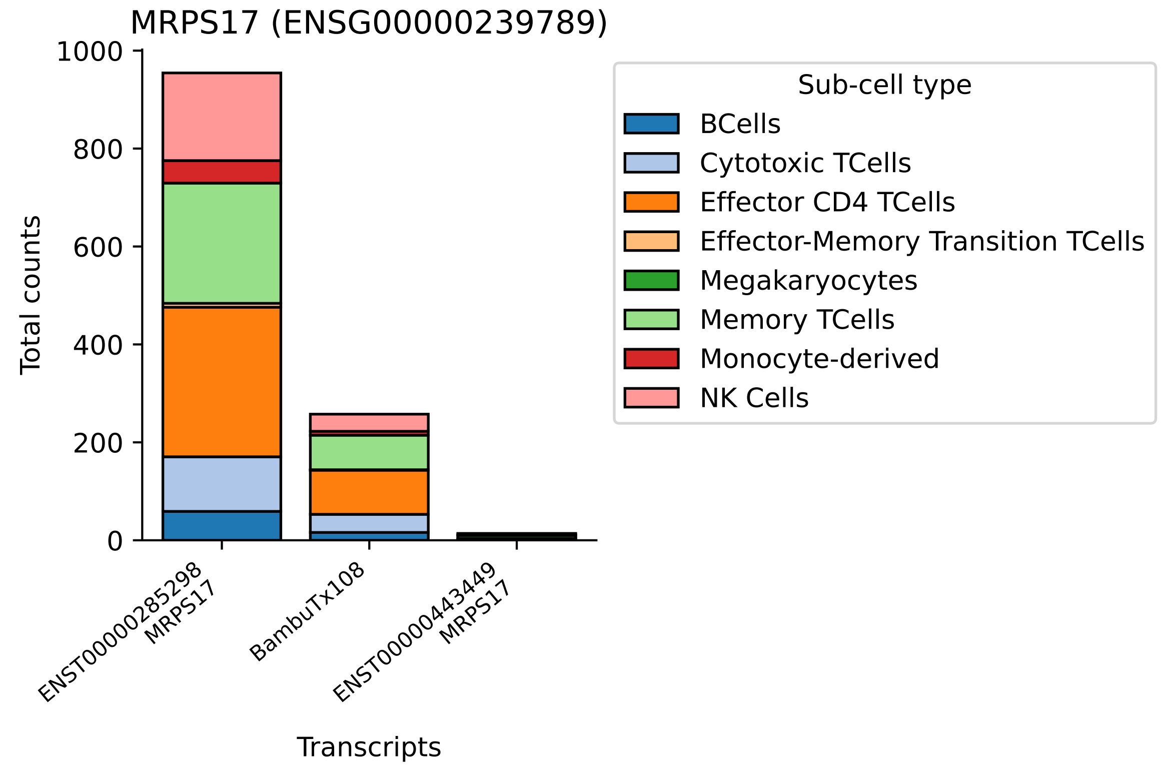

MRPS17 (ENSG00000239789)
Total counts
1000
800
600
400
200
0
ENST00000285298
MRPS17
BambuTx108
ENST00000443449
MRPS17
Transcripts
Sub-cell type
BCells
Cytotoxic TCells
Effector CD4 TCells
Effector-Memory Transition TCells
Megakaryocytes
Memory TCells
Monocyte-derived
NK Cells

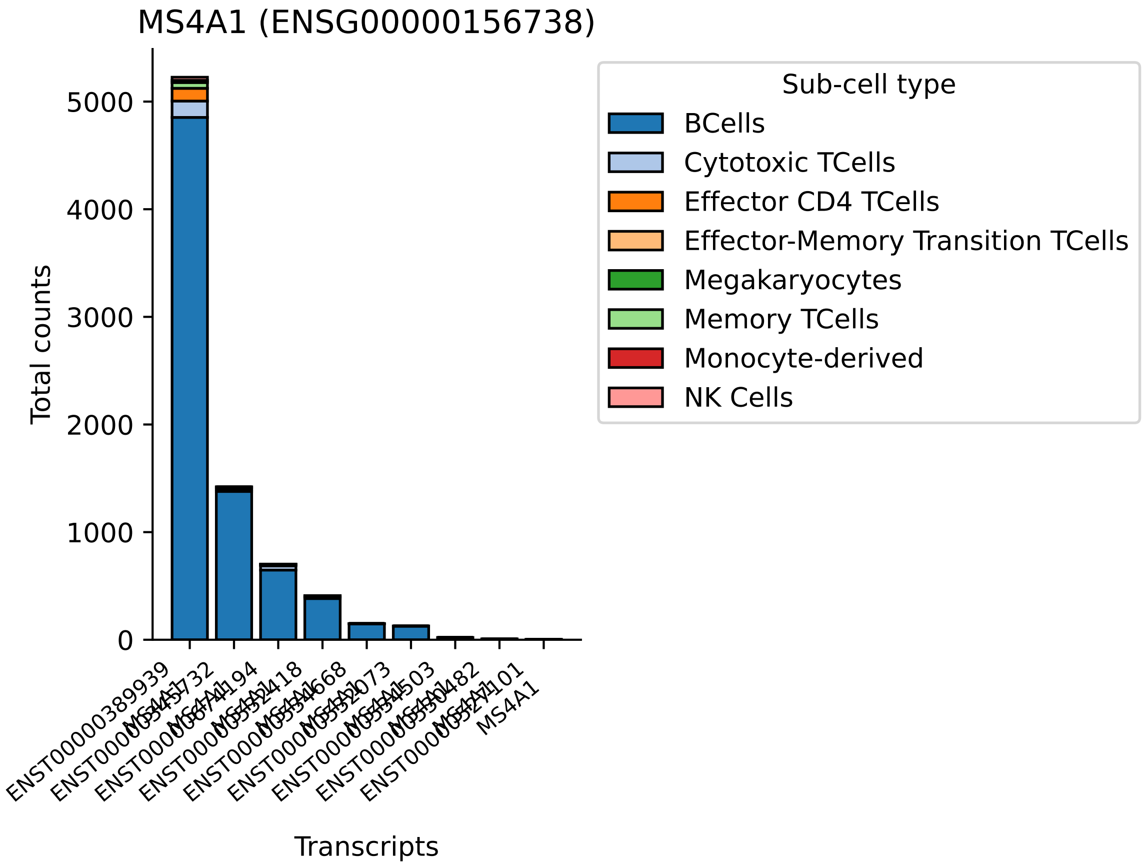
MS4A1 (ENSG00000156738)
Total counts
0
1000
2000
3000
4000
5000
ENST00000389939
MS4A1
Transcripts
Sub-cell type
BCells
Cytotoxic TCells
Effector CD4 TCells
Effector-Memory Transition TCells
Megakaryocytes
Memory TCells
Monocyte-derived
NK Cells

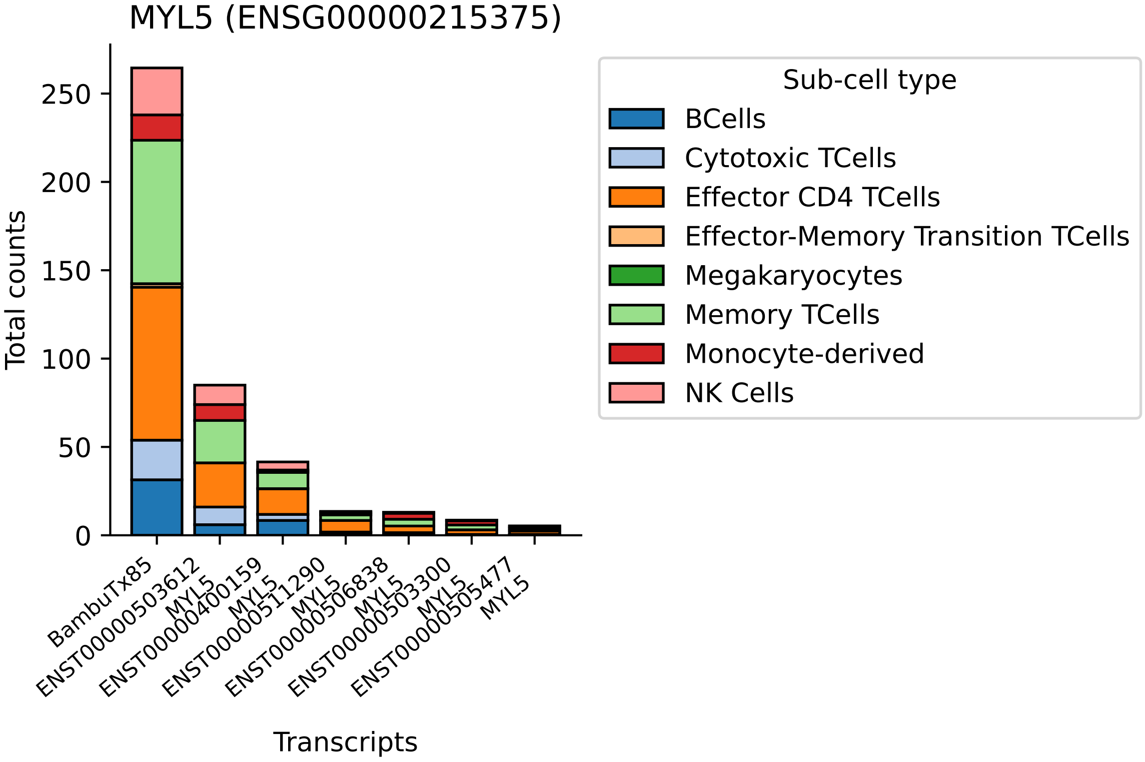

MYL5 (ENSG00000215375)
Total counts
0
50
100
150
200
250
Transcripts
BambuTx85
ENST00000503612
MYL5
ENST00000400159
MYL5
ENST00000511290
MYL5
ENST00000506838
MYL5
ENST00000503300
MYL5
ENST00000505477
MYL5
Sub-cell type
BCells
Cytotoxic TCells
Effector CD4 TCells
Effector-Memory Transition TCells
Megakaryocytes
Memory TCells
Monocyte-derived
NK Cells

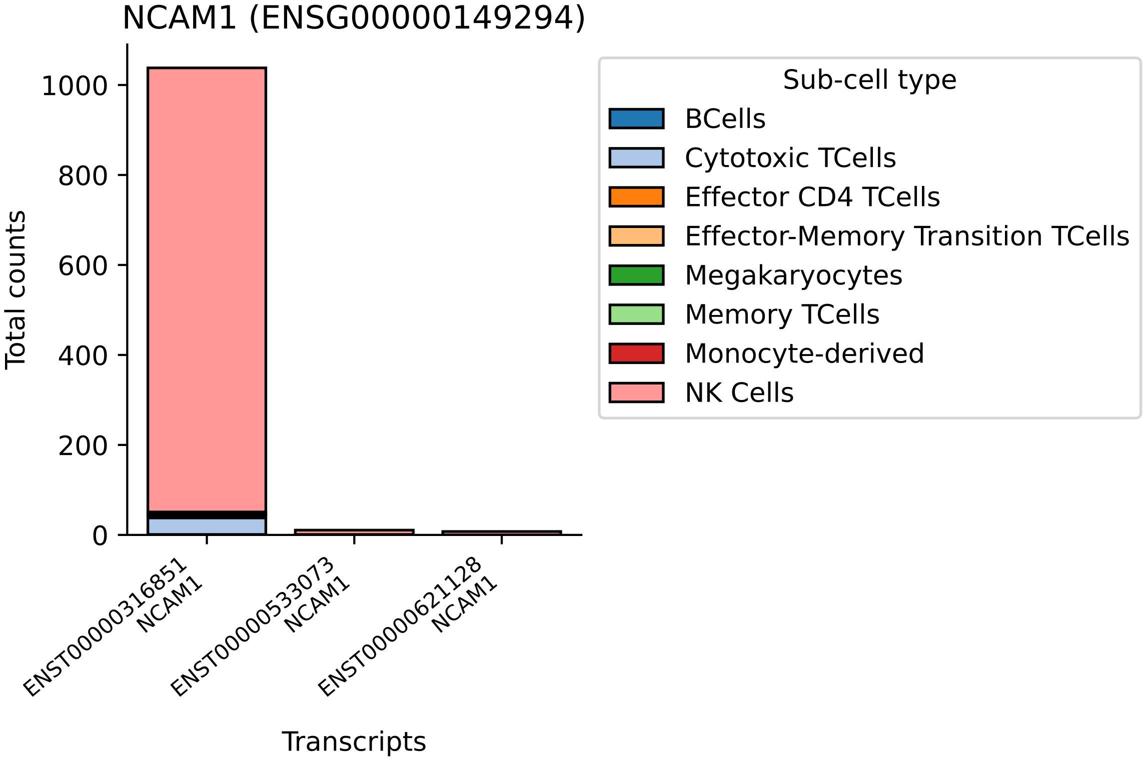
NCAM1 (ENSG00000149294)
Total counts
0
200
400
600
800
1000
ENST00000316851
NCAM1
ENST00000533073
NCAM1
ENST00000621128
NCAM1
Transcripts
Sub-cell type
BCells
Cytotoxic TCells
Effector CD4 TCells
Effector-Memory Transition TCells
Megakaryocytes
Memory TCells
Monocyte-derived
NK Cells

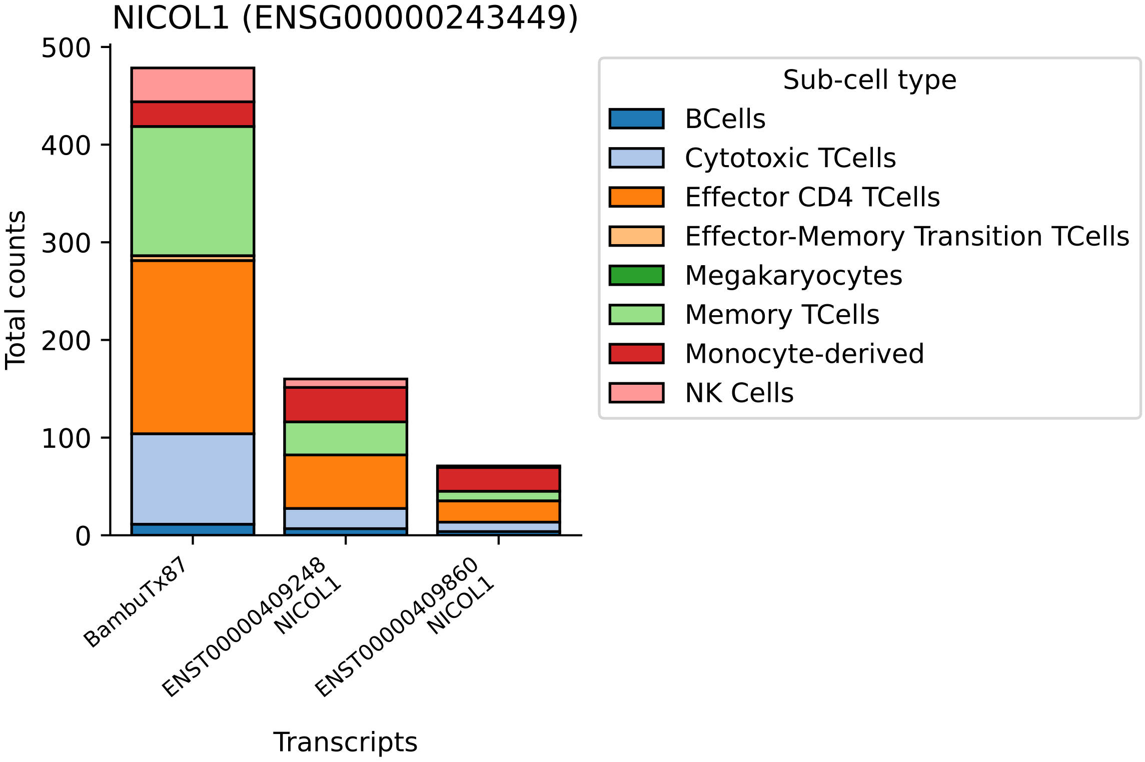
NICOL1 (ENSG00000243449)
Total counts
500
400
300
200
100
0
BambuTx87
ENST00000409248
NICOL1
ENST00000409860
NICOL1
Transcripts
Sub-cell type
BCells
Cytotoxic TCells
Effector CD4 TCells
Effector-Memory Transition TCells
Megakaryocytes
Memory TCells
Monocyte-derived
NK Cells

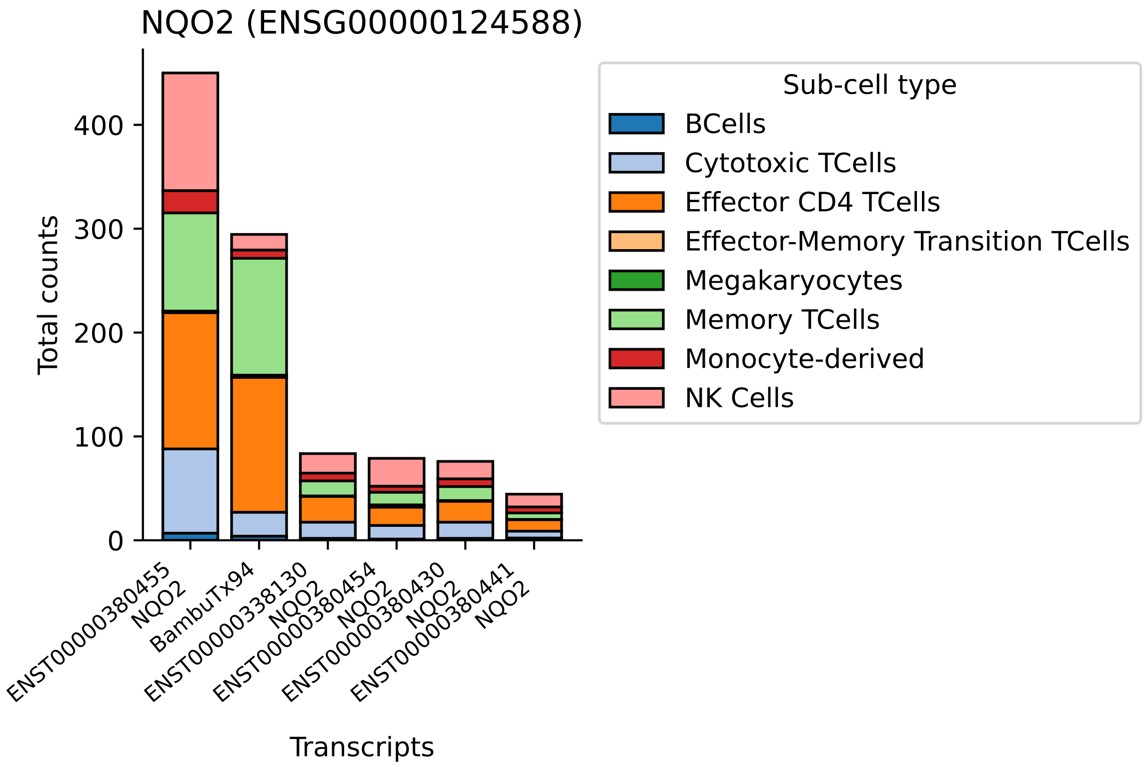
NQO2 (ENSG00000124588)
Total counts
0
100
200
300
400
ENST00000380455
NQO2
BambuTx94
ENST00000338130
NQO2
ENST00000380454
NQO2
ENST00000380430
NQO2
ENST00000380441
NQO2
Transcripts
Sub-cell type
BCells
Cytotoxic TCells
Effector CD4 TCells
Effector-Memory Transition TCells
Megakaryocytes
Memory TCells
Monocyte-derived
NK Cells

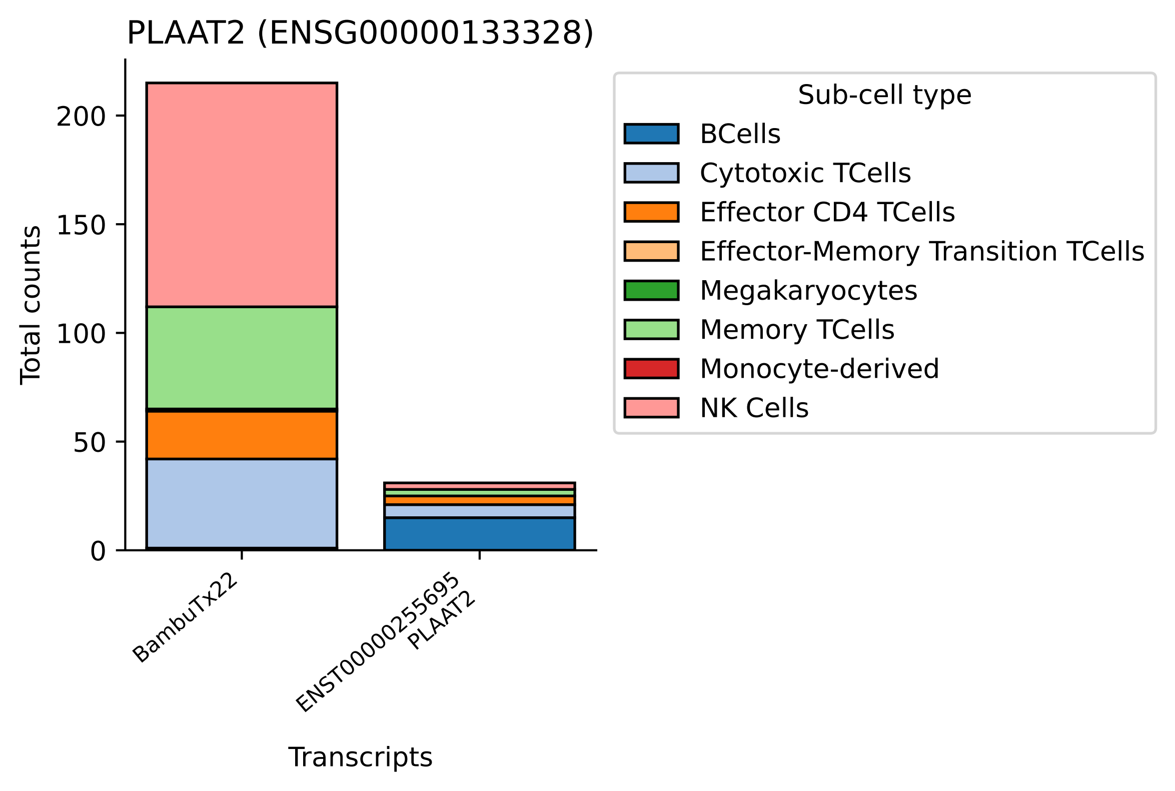

PLAAT2 (ENSG00000133328)
Total counts
0
50
100
150
200
BambuTx22
ENST00000255695
PLAAT2
Transcripts
Sub-cell type
BCells
Cytotoxic TCells
Effector CD4 TCells
Effector-Memory Transition TCells
Megakaryocytes
Memory TCells
Monocyte-derived
NK Cells

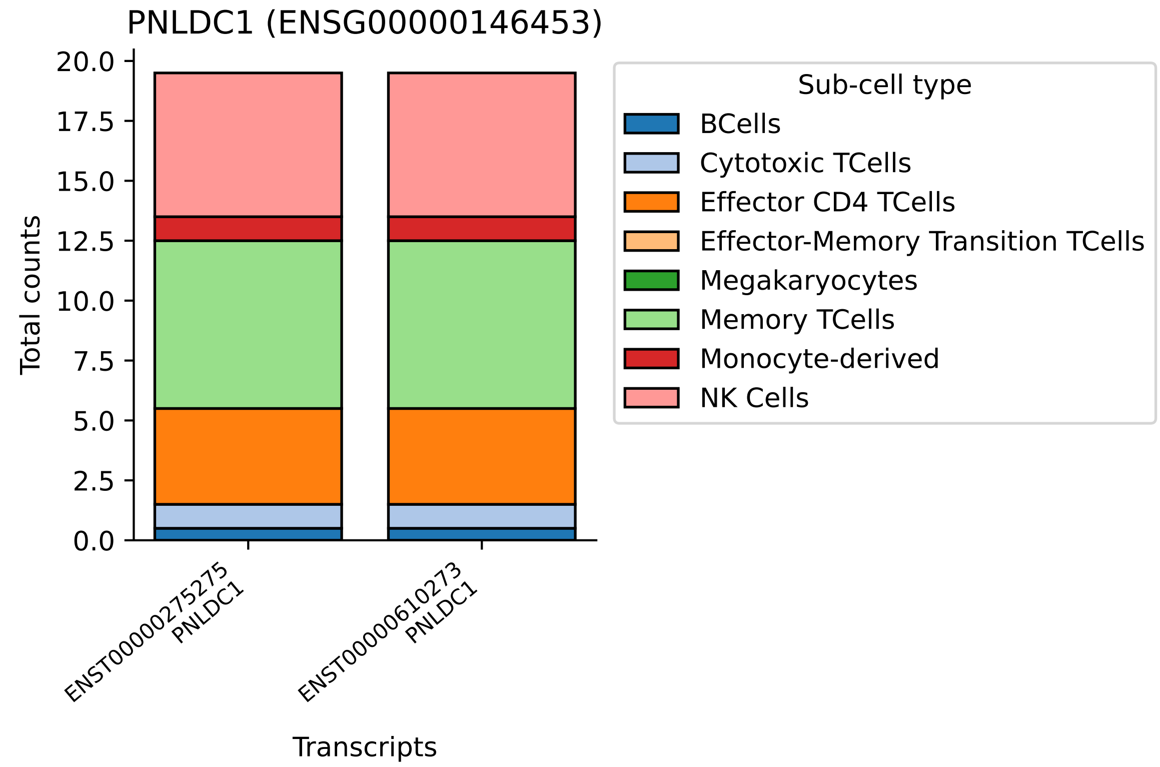

PNLDC1 (ENSG00000146453)
Total counts
20.0
17.5
15.0
12.5
10.0
7.5
5.0
2.5
0.0
ENST00000275275
PNLDC1
ENST00000610273
PNLDC1
Transcripts
Sub-cell type
BCells
Cytotoxic TCells
Effector CD4 TCells
Effector-Memory Transition TCells
Megakaryocytes
Memory TCells
Monocyte-derived
NK Cells

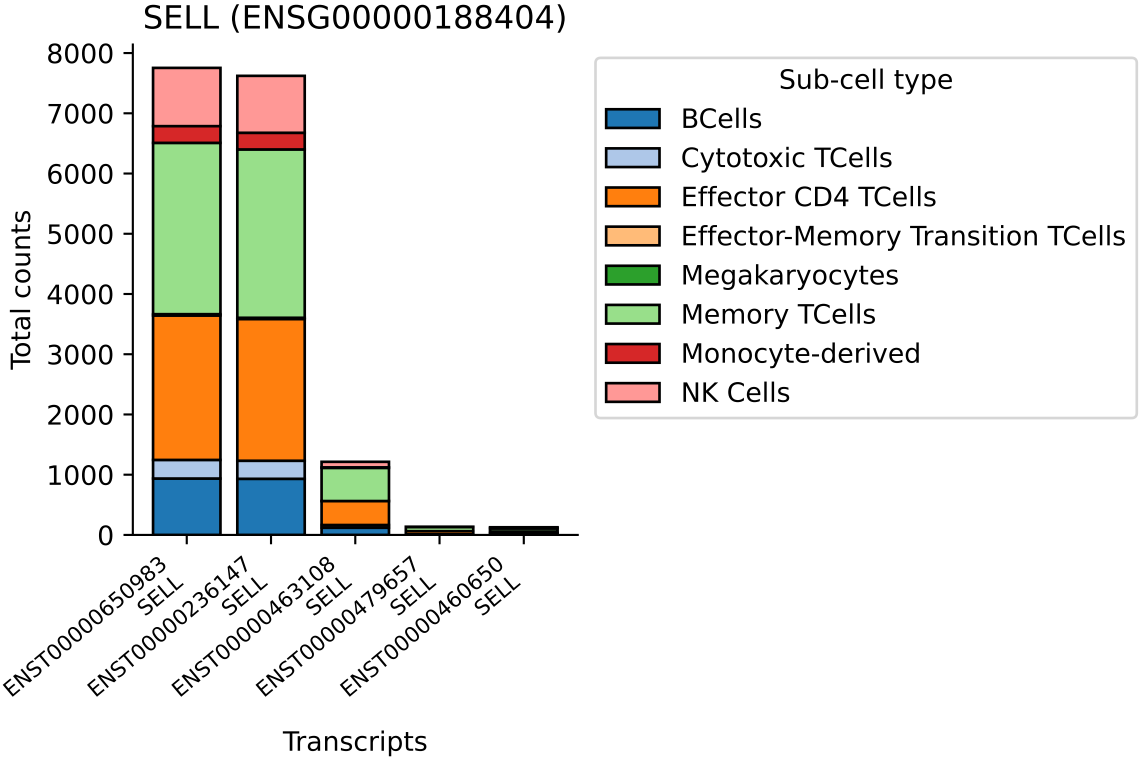

SELL (ENSG00000188404)
Total counts
0
1000
2000
3000
4000
5000
6000
7000
8000
ENST00000650983 SELL
ENST00000236147 SELL
ENST00000463108 SELL
ENST00000479657 SELL
ENST00000460650 SELL
Transcripts
Sub-cell type
BCells
Cytotoxic TCells
Effector CD4 TCells
Effector-Memory Transition TCells
Megakaryocytes
Memory TCells
Monocyte-derived
NK Cells

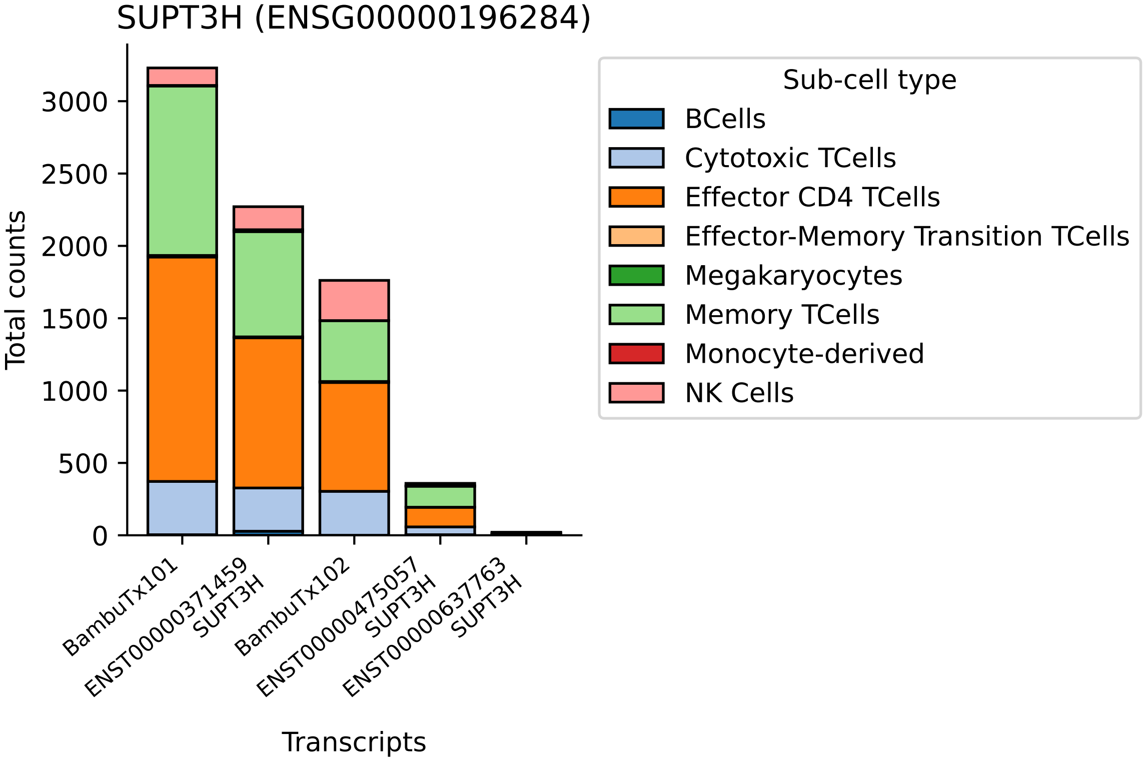

SUPT3H (ENSG00000196284)
Total counts
0
500
1000
1500
2000
2500
3000
BambuTx101
ENST00000371459 SUPT3H
BambuTx102
ENST00000475057 SUPT3H
ENST00000637763 SUPT3H
Transcripts
Sub-cell type
BCells
Cytotoxic TCells
Effector CD4 TCells
Effector-Memory Transition TCells
Megakaryocytes
Memory TCells
Monocyte-derived
NK Cells

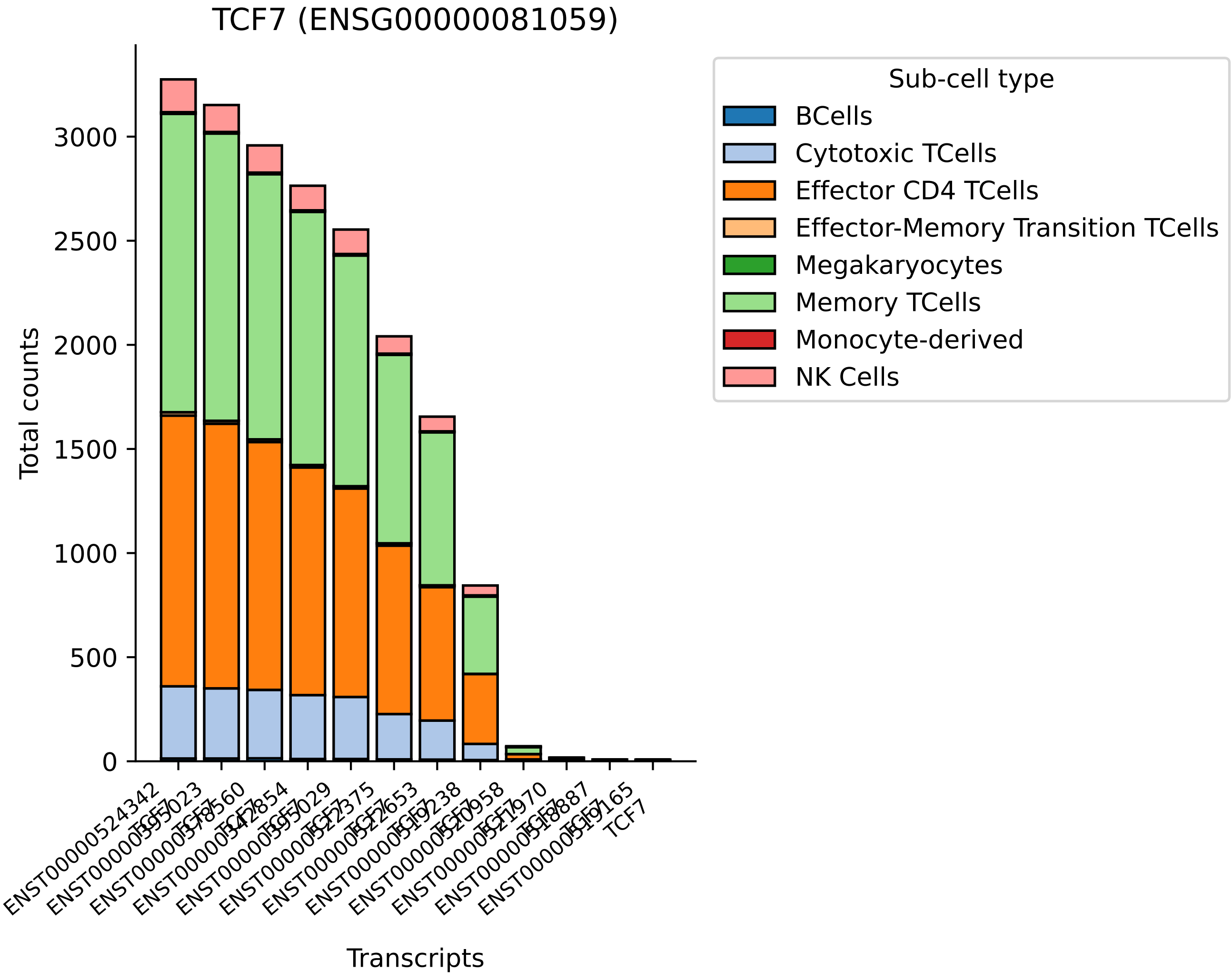

TCF7 (ENSG00000081059)
Total counts
Transcripts
0
500
1000
1500
2000
2500
3000
Sub-cell type
BCells
Cytotoxic TCells
Effector CD4 TCells
Effector-Memory Transition TCells
Megakaryocytes
Memory TCells
Monocyte-derived
NK Cells

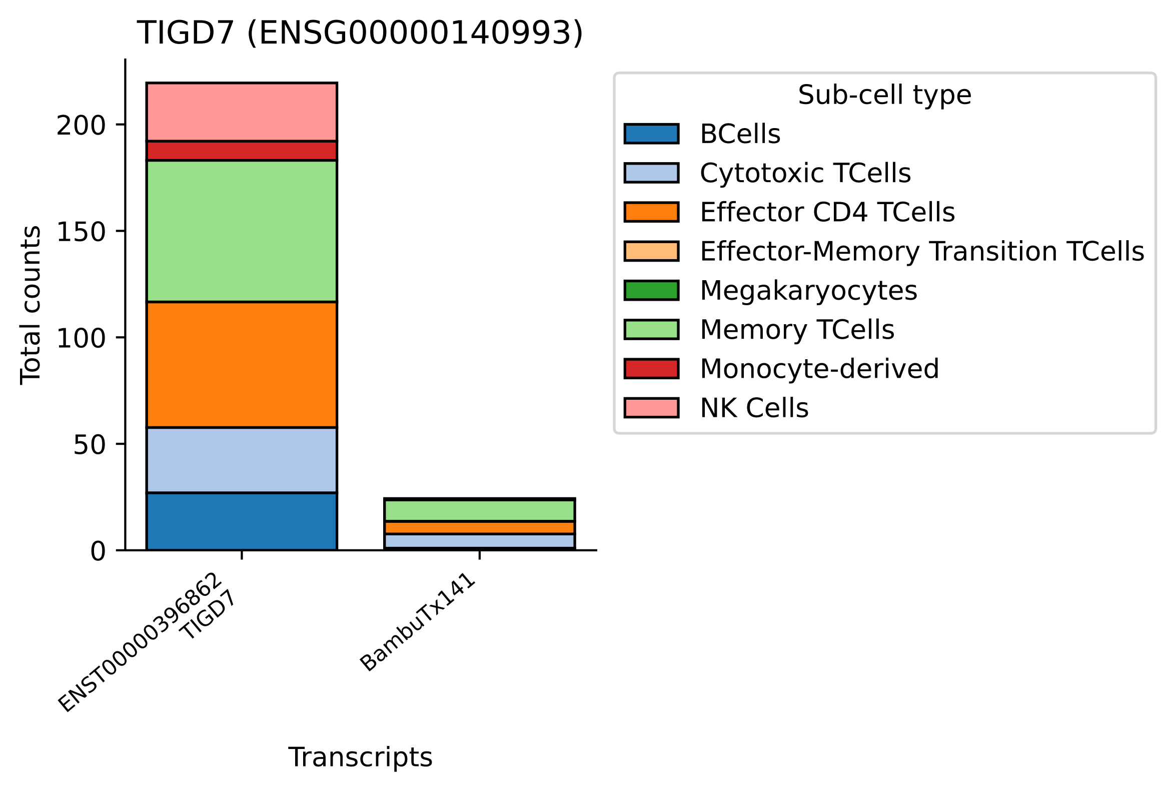
TIGD7 (ENSG00000140993)
Total counts
0
50
100
150
200
ENST00000396862
TIGD7
BambuTx141
Transcripts
Sub-cell type
BCells
Cytotoxic TCells
Effector CD4 TCells
Effector-Memory Transition TCells
Megakaryocytes
Memory TCells
Monocyte-derived
NK Cells

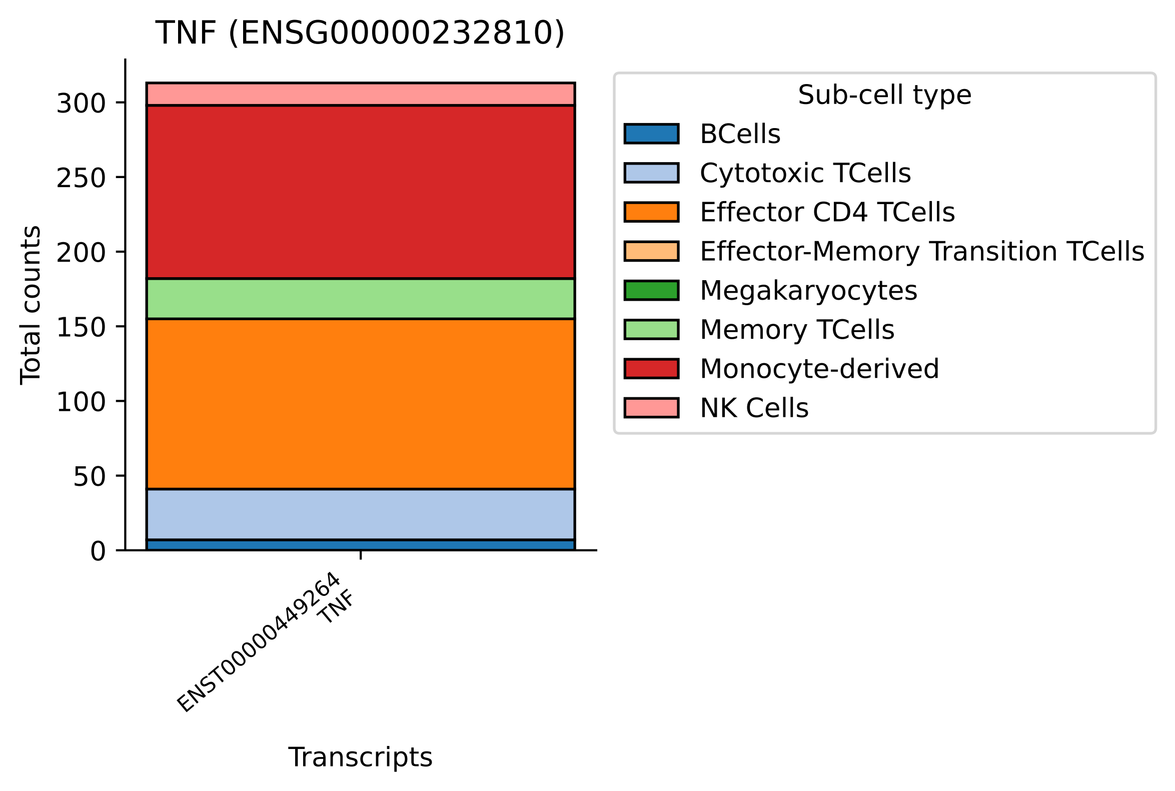

TNF (ENSG00000232810)
Total counts
0
50
100
150
200
250
300
ENST00000449264
TNF
Transcripts
Sub-cell type
BCells
Cytotoxic TCells
Effector CD4 TCells
Effector-Memory Transition TCells
Megakaryocytes
Memory TCells
Monocyte-derived
NK Cells

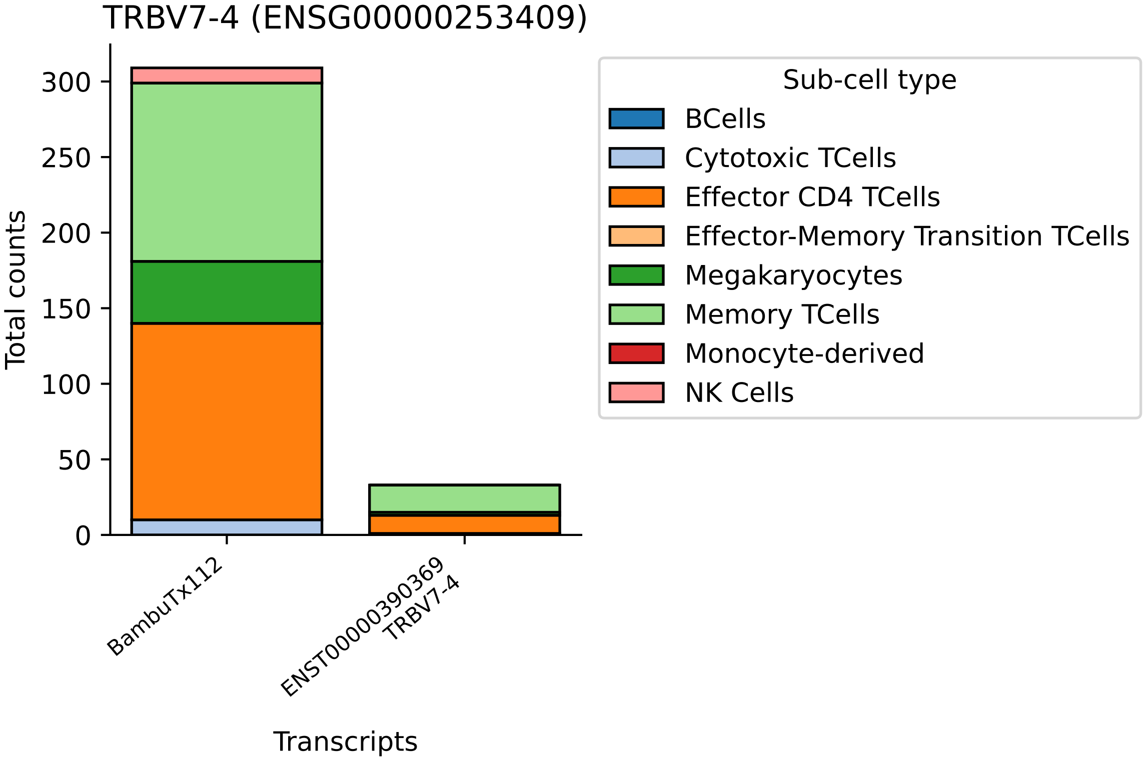

TRBV7-4 (ENSG00000253409)
Total counts
0
50
100
150
200
250
300
BambuTx112
ENST00000390369
TRBV7-4
Transcripts
Sub-cell type
BCells
Cytotoxic TCells
Effector CD4 TCells
Effector-Memory Transition TCells
Megakaryocytes
Memory TCells
Monocyte-derived
NK Cells

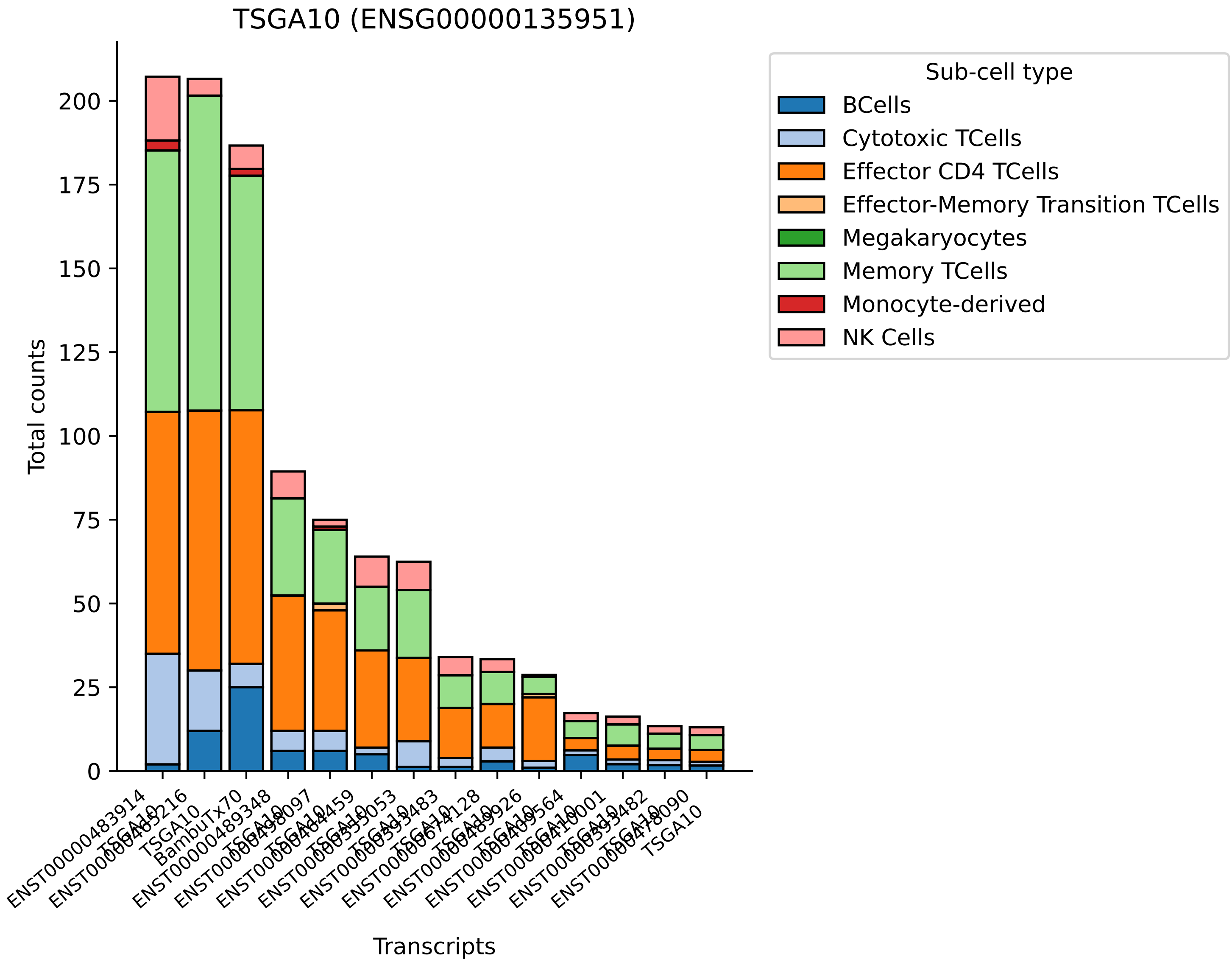
TSGA10 (ENSG00000135951)
Total counts
0
25
50
75
100
125
150
175
200
Transcripts
Sub-cell type
BCells
Cytotoxic TCells
Effector CD4 TCells
Effector-Memory Transition TCells
Megakaryocytes
Memory TCells
Monocyte-derived
NK Cells

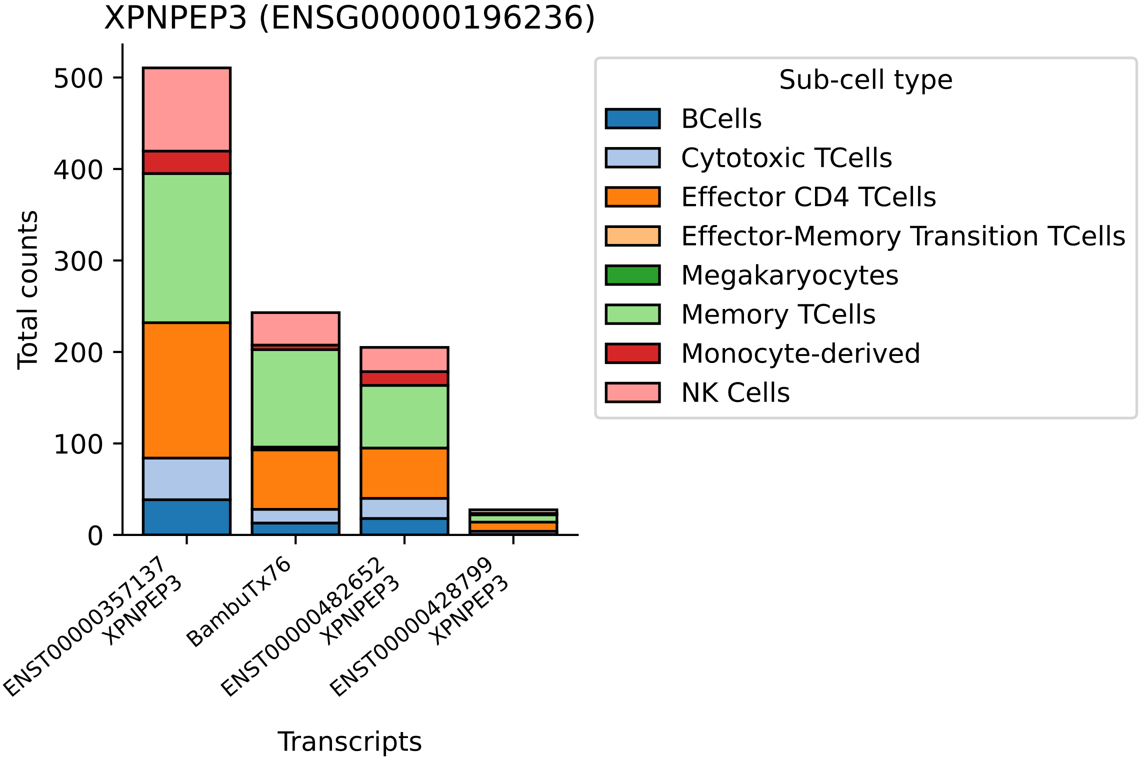

XPNPEP3 (ENSG00000196236)
Total counts
0
100
200
300
400
500
ENST00000357137
XPNPEP3
BambuTx76
ENST00000482652
XPNPEP3
ENST00000428799
XPNPEP3
Transcripts
Sub-cell type
BCells
Cytotoxic TCells
Effector CD4 TCells
Effector-Memory Transition TCells
Megakaryocytes
Memory TCells
Monocyte-derived
NK Cells

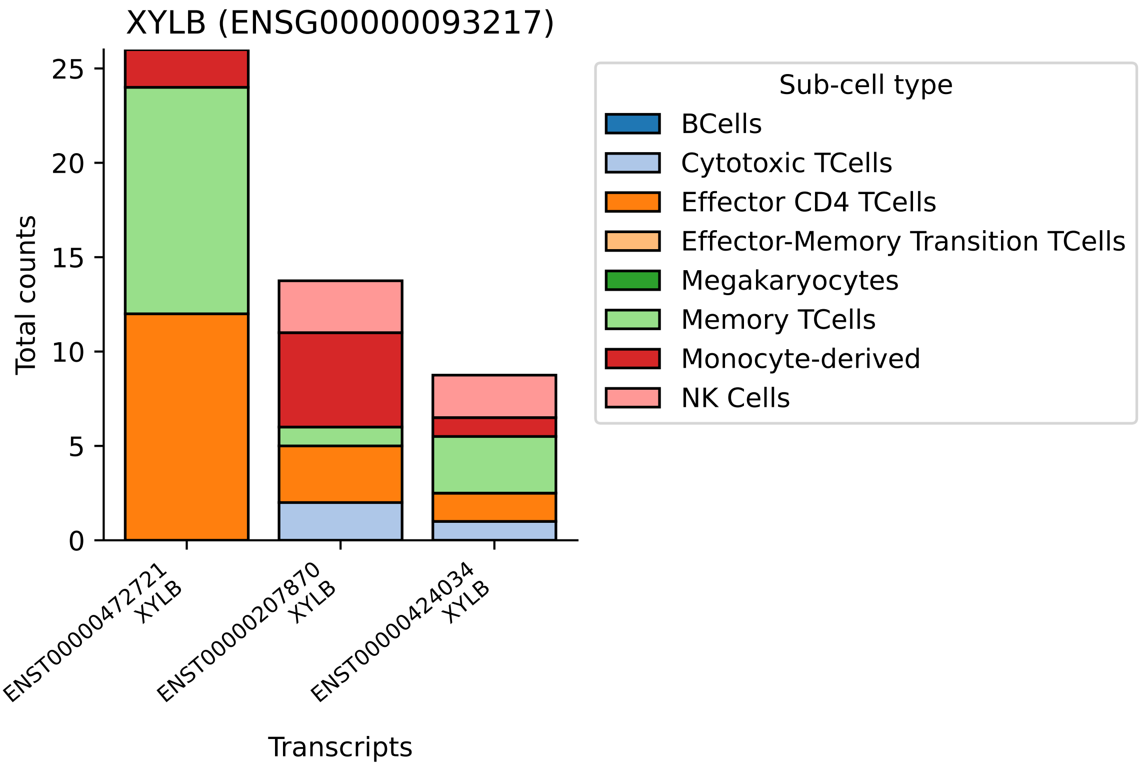
XYLB (ENSG00000093217)
Total counts
0
5
10
15
20
25
ENST00000472721
XYLB
ENST00000207870
XYLB
ENST00000424034
XYLB
Transcripts
Sub-cell type
BCells
Cytotoxic TCells
Effector CD4 TCells
Effector-Memory Transition TCells
Megakaryocytes
Memory TCells
Monocyte-derived
NK Cells

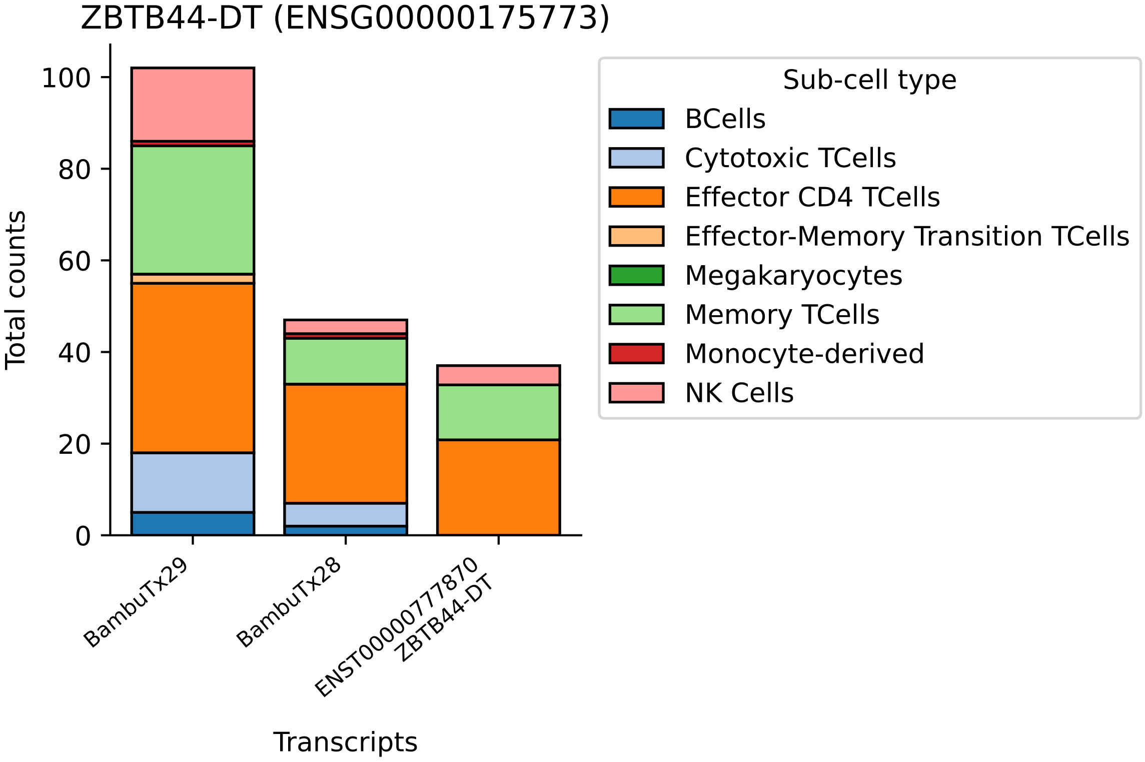

ZBTB44-DT (ENSG00000175773)
Total counts
0
20
40
60
80
100
BambuTx29
BambuTx28
ENST00000777870
ZBTB44-DT
Transcripts
Sub-cell type
BCells
Cytotoxic TCells
Effector CD4 TCells
Effector-Memory Transition TCells
Megakaryocytes
Memory TCells
Monocyte-derived
NK Cells

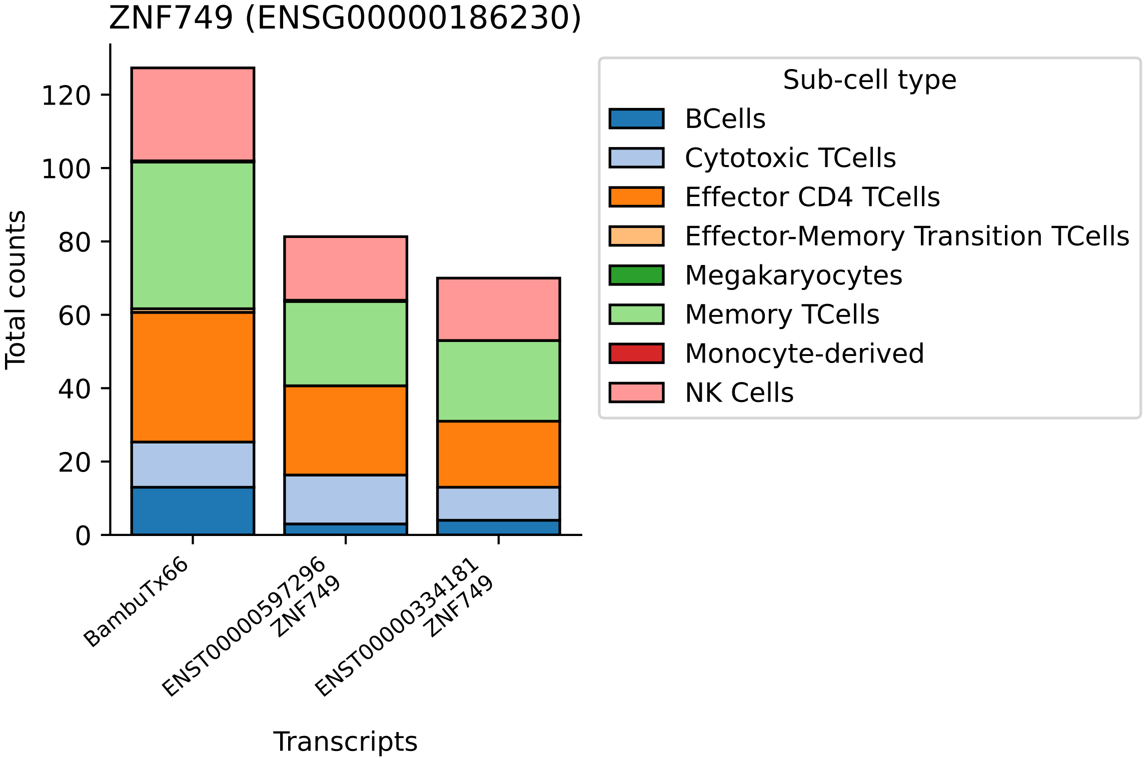

ZNF749 (ENSG00000186230)
Total counts
0
20
40
60
80
100
120
BambuTx66
ENST00000597296
ZNF749
ENST00000334181
ZNF749
Transcripts
Sub-cell type
BCells
Cytotoxic TCells
Effector CD4 TCells
Effector-Memory Transition TCells
Megakaryocytes
Memory TCells
Monocyte-derived
NK Cells
